# Epistasis decreases with increasing antibiotic pressure but not temperature

**DOI:** 10.1101/2022.09.01.506172

**Authors:** Ana-Hermina Ghenu, André Amado, Isabel Gordo, Claudia Bank

## Abstract

Predicting mutational effects is essential for the control of antibiotic resistance (ABR). Predictions are difficult when there are strong genotype-by-environment (G*×*E), gene-by-gene (G*×*G or epistatic), or gene-by-gene-by-environment (G*×*G*×*E) interactions. We quantified G*×*G*×*E effects in *Escherichia coli* across environmental gradients. We created intergenic fitness landscapes using gene knock-outs and single nucleotide ABR mutations previously identified to vary in the extent of G*×*E effects in our environments of interest. Then, we measured competitive fitness across a complete combinatorial set of temperature and antibiotic dosage gradients. In this way, we assessed the predictability of 15 fitness landscapes across 12 different but related environments. We found G*×*G interactions and rugged fitness landscapes in the absence of antibiotic, but as antibiotic concentration increased, the fitness effects of ABR genotypes quickly overshadowed those of gene knock-outs, and the landscapes became smoother. Our work reiterates that some single mutants, like those conferring resistance or susceptibility to antibiotics, have consistent effects across genetic backgrounds in stressful environments. Thus, although epistasis may reduce the predictability of evolution in benign environments, evolution may be more predictable in adverse environments.

## 1 Introduction

It has been debated for decades whether gene-by-gene interactions (G*×*G or epistasis) influence evolutionary processes [reviewed in 1, 2]. Compelling evidence for the successive fixation of epistatic substitutions comes from studies of amino-acid replacements over phylogenetic time scales [e.g., 3–5]. Moreover, experimental evolution has demonstrated that adaptive paths can be constrained by epistasis [e.g., 6]. Also, the environment can modulate mutational effects through gene-by-environment (G*×*E) interactions. Whereas multiple studies have demonstrated the existence of GxE interactions [e.g., 7–9], our knowledge of the extent and the consequences of GxGxE interactions, i.e. those in which epistasis interacts with the environment, are limited [10–13, reviewed in 2].

G*×*G, G*×*E, and G*×*G*×*E interactions complicate evolutionary predictions because they alter expected phenotypes or fitness of individuals [2, 14]. Studying their extent and incorporating their effect into evolutionary predictions is daunted by the complexity of the genotype space and the myriad of environments that could be tested [15]. However, our lack of quantitative knowledge of G*×*G and G*×*E effects poses dangers to fields in which genetic and evolutionary models are applied: they could lead to the failed genetic rescue of endangered species, or the unexpected spread or maintenance of antibiotic resistance (ABR).

At the same time, the existence of G*×*E interactions is a crucial assumption in ABR evolution. By definition, the original genotype grows well in the antibiotic-free environment and poorly in the antibiotic environment. Conversely, a resistant genotype grows well in the antibiotic environment and is usually assumed to have low fitness in the antibiotic-free environment, resulting in so-called costs of ABR [reviewed in 16, but see 17]. In addition, G*×*E interactions with other environmental variables, such as high temperature or minimal media, can facilitate the evolution of resistant genotypes in the absence of antibiotics [6, 18]. Although G×E interactions are central in ABR evolution, many questions about the role of G×E remain unexplored [but see 19–21]; for example, how does the G×E relationship between susceptible and resistant genotypes change across non-antimicrobial environmental axes, and how frequently is ABR modulated by the genetic background (i.e., through G×G effects).

ABR was first identified as a problem shortly after the introduction of antibiotics for clinical use [22], and it continues to be a leading cause of death worldwide [23]. For this reason, inferring potential resistances, cross-resistances, compensatory mutations, and, ultimately, predicting the evolutionary trajectories of bacterial genomes in the presence of antibiotics is an application of evolutionary biology that is important for the health of humans and the planet [24, 25]. Critically, G*×*G and G*×*E effects influencing ABR can complicate the identification of new resistances. For example, with G*×*G effects, resistance may depend on more than one easily identifiable allele. In addition, with G*×*E effects, laboratory evolution settings may not be informative of the fitness of genotypes in the wild.

Including G*×*G effects in the study of ABR requires the consideration of many potentially interacting genotypes. With epistasis, interactions can be specific between particular mutations due to mechanistic interactions of residues or proteins, for example [reviewed in 26], or global, where any combination of mutations ultimately shows a pattern of G*×*G [27, 28]. In contrast to increasing numbers of reports of ubiquitous epistasis [29, 12, 30] much existing work to date has considered single ABR genotypes at a time under the assumptions of additive effects [but see 31, 32]. The theoretical concept of a fitness landscape, which maps every possible genotype to its fitness, captures G*×*G interactions and enables the study of how such interactions affect evolutionary trajectories [reviewed in 2, 33]. Unless there is a known underlying pattern of interactions, epistasis makes evolution on the fitness landscape less predictable. Moreover, with added G*×*E interactions, the fitness landscape may change across environmental gradients, leading to changes in both single genotypes and epistasis [12, 8] and to different possible evolutionary trajectories [34, 13].

We here take a step towards quantifying G*×*G*×*E interactions by studying 15 small (i.e., two mutational-step) fitness landscapes across two environmental gradients and their combination, antibiotic concentration and temperature. In our fitness landscapes, we combine three known ABR mutations in the *rpoB* gene with five gene knock-outs, resulting in 24 genotypes that are screened for competitive fitness in 12 abiotic environments. Unlike previous studies, we study interactions of two different mutation types, single amino-acid substitutions (more specifically, single nucleotide polymorphisms, or SNPs) and whole gene deletions. By screening the fitness landscapes across a grid of environments, we quantify their change across environmental gradients, which is related to the robustness of fitness predictions under environmental change. Since the ABR mutations and gene knock-outs were selected based on their *not* exhibiting any direct, mechanistic interactions, one might *a priori* expect few *G × G* interactions and that the fitness of the double mutants would be additive as compared to the single mutants. On the other hand, non-specific epistasis (epistasis between mutations that do not directly interact mechanistically) has been observed ubiquitously [26]. Our fitness landscapes feature moderate epistasis in the benign environments, which decreases at higher antibiotic concentrations. Altogether, we conclude that our fitness landscapes are very predictable across environments.

## 2 Results

### 2.1 Choice of genotypes

We selected rifampicin-resistant single-nucleotide mutations that were *a priori* known to differ in their performance between different temperature environments in M9 minimal medium. The ABR mutations modify single amino-acids at the antibiotic target site in the gene RNA polymerase B (*rpoB*). Prior to our study, it was unknown how the mutants would perform in different combinations of antibiotic and high-temperature environments (i.e., we only knew about the effect of one environmental axis on its own). *RpoB* H526Y was established to show a trade-off between environment types: it confers high rifampicin resistance at 37*^◦^C* but grows poorly as compared to wild-type at higher temperatures in the absence of antibiotic [7, 35]. Conversely, *rpoB* S512F was established to have a synergy between environment types: it grows well both at high rifampicin concentrations and higher temperatures [7], separately. Finally, *rpoB* I572N is a weak rifampicin resistance mutation that was first identified during evolution to high temperature [6, 36].

We initially tried to select knock-out mutations similarly as we had done for *rpoB* mutants: we used data from a large phenotypic screen of *E. coli* (accessed from ecoliwiki.org/tools/chemgen/) to select knock-out mutations that were known to differ in their performance between environment types. However, neither the bacterial colony size s-scores [37] nor the bacterial colony opacity and density s-scores [38] were able to predict the growth effects of Keio collection [39] knock-out mutants in batch culture environments (data not shown). Ultimately, we selected knock-out mutations based on their *a priori* functional effects and the presence of segregating knock-out polymorphism in natural populations (accessed from the panX database at pangenome.org; [40]). We selected four knock-outs of functional genes, all involved in how the cell interacts with its environment as follows: *marR* (multiple antibiotic resistance regulator) detects and responds to chemical stressors, and its knock-out is sensitive to heat shock [41], conferring either resistance [42] or susceptibility [43] to some antibiotics but not rifampicin [37]; *nuoC* and *yidK* are both located on the plasma membrane and involved in transport [44, 45]; and *waaP* impacts outer membrane stability through lipopolysaccharide biosynthesis [46]. The fifth gene, *ybfG*, is a pseudogene and was selected as a control since its knock-out should perform similarly to the wild-type genotype. None of the knock-outs interact directly with *rpoB* or other genes involved in transcription, protein synthesis, or DNA supercoiling. Therefore, any epistasis that will be observed should be an instance of ‘non-specific epistasis’ [26]. Non-specific epistasis is attributable to factors *other than* mechanistic interactions: for example, non-linearities in the genotype-to-phenotype map.

Figure 1a shows a schematic of the resulting topology of the fitness landscape when combining the ABR and knock-out mutations.

**Figure 1:**
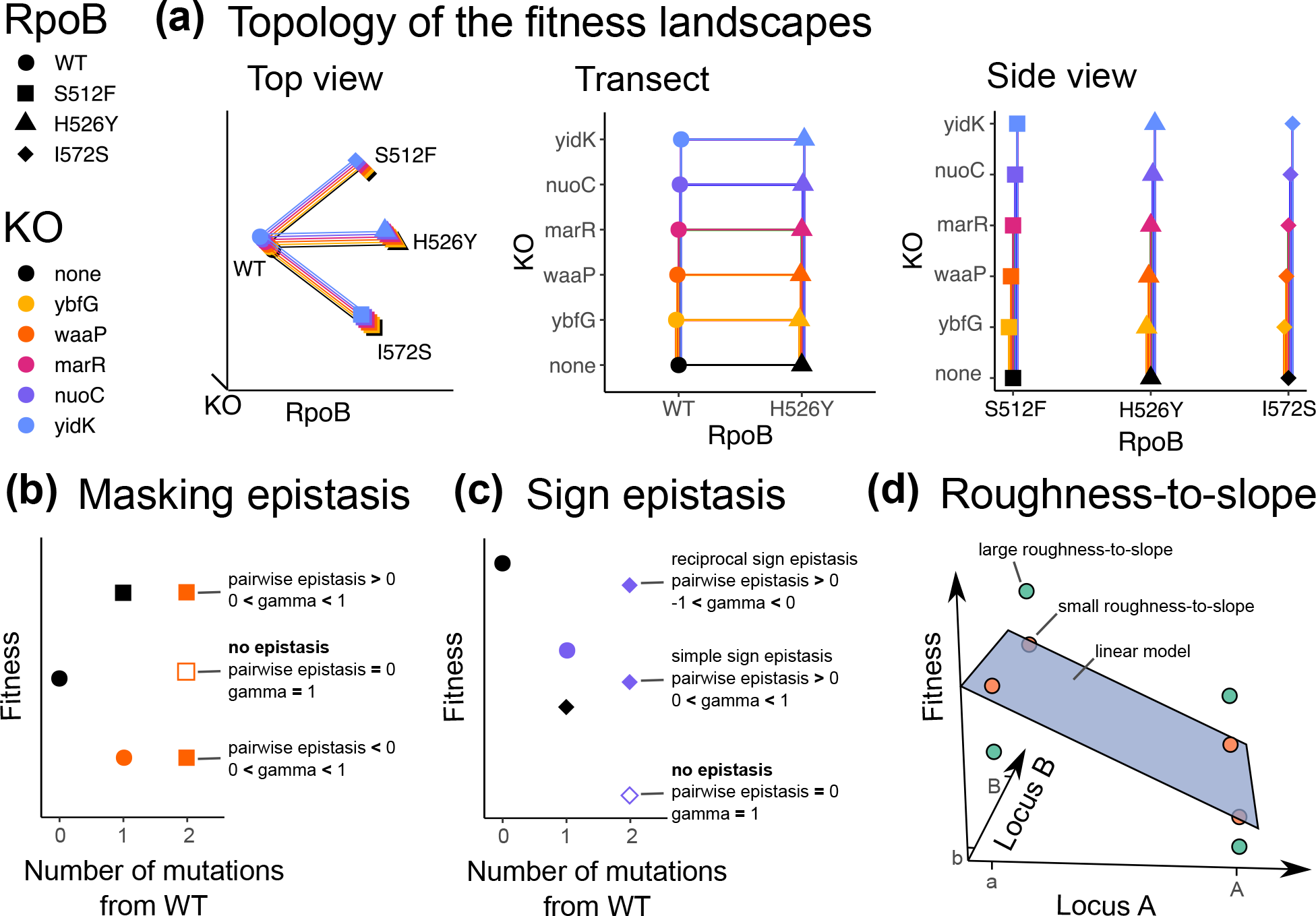
a) 3-Dimensional schematics depicting the topology of the studied fitness landscapes. The schematics show the same topology from different perspectives. All mutations are at unique, di-allelic loci. There are 15 fitness landscapes each composed of the wild-type, a *rpoB* single mutant, a knock-out single mutant, and the resultant double mutant. Lines connect genotypes that are one mutational step apart. **Depiction of ‘classical’ pairwise epistasis and gamma epistasis metrics for (b) masking and (c) sign epistasis.** Masking epistasis is when the fitness effect of a beneficial or deleterious mutation on the wild-type background is zero on a different background. Sign epistasis occurs when the beneficial or deleterious effect of a mutation reverses in the presence of another mutation to become deleterious or beneficial, respectively. Reciprocal sign epistasis is a special case of sign epistasis where both mutations change sign on the other’s background. The gamma statistic measures the correlation of fitness effects among the different genetic backgrounds where a mutation appears. The white-filled shape shows the expected fitness value without epistasis. For panels (a)-(c) the shape indicates the *rpoB* point mutation and the colour indicates which gene has been knocked-out (KO); the black circle indicates the wild-type genotype without any mutations on *rpoB* and without any knock-outs. **(d) Fitness landscapes exhibiting large (teal points) and small (orange points) roughness-to-slope ratios.** The roughness-to-slope statistic measures how different the fitness landscape is from an additive, linear model (a purely additive landscape has a ratio of zero). The plane shows the fit of the multidimensional linear model to the fitness landscapes for two di-allelic loci. The vertical distance between the genotype (i.e., point) and the plane shows its residual. Epistasis is quantified as the ratio of the standard deviation of the residuals to the mean of the slopes of the plane. Panel (d) does not follow the colour and shape scheme as the other panels.

Illumina whole-genome re-sequencing was used to confirm the constructed mutants and identify any *de novo* mutations compared to the wild-type genotype (table S3). See the supplementary results for details.

### 2.2 Choice of environments

Since the wild-type and knock-out single mutants are sensitive to rifampicin, the screened antibiotic concentrations were selected based on the wild-type dose response (figure S14). The screened temperature environments were selected based on their physiological relevance for *E. coli* and its human host: 37*^◦^C* is the temperature of a healthy human host, 40*^◦^C* is the temperature of high fever, and 42*^◦^C* is near the maximum temperature that *E. coli* can withstand. Antibiotic-susceptible genotypes (i.e., genotypes that carry the wild-type allele at the *rpoB* locus) can be considered as ‘specialists’ of antibiotic-free environments because their growth deteriorates in the presence of the antibiotic, i.e. they show strong G*×*E interactions. Based on previous results, we expected that the *rpoB* S512F single mutant that grows well in the presence of antibiotic and at high temperature, separately, would exhibit ‘generalist’ growth in all environments with little G*×*E interaction.

### 2.3 Measure of competitive fitness

The GFP-labeled wild-type, single, and double mutants were competed against the same reference genotype, mCherry-labeled *rpoB* H526Y, in all environments by starting at approximately equal ratios and a fixed inoculum size. The competitive index (*w*^), a proxy for fitness, was estimated after 20 hours of batch culture growth as follows:

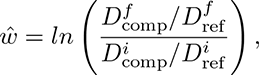

where *D^i^* represents the initial density and *D^f^* represents the final density as measured by flow cytometry. We found nearly identical *w*^ estimates when the fluorophores of the reference and competitor genotypes were swapped (*w*^_GFP_ = *β ∗ w*^_mCherry_ + *b*; adjusted *R*^2^ = 0.95; *β* = 0.96, *p <* 10*^−^*^15^; figure S15 and table S4); including the effect of environment in the regression was not significant (*F* (11, 127) = 1.27, *p* = 0.25, table S5 and figure S16), but adding the effect of genotype was significant (*F* (11, 127) = 4.00, *p <* 10*^−^*^4^, figure S17 and table S6), although genotype explains only about 1% more of the variation in the data (adjusted *R*^2^ = 0.96 for overall regression) as compared to not including this effect. Therefore we concluded that the fluorescence is a neutral marker.

The mean *w*^ for all genotypes and environments is shown in figure 2. As expected, genotypes that do not have an ABR mutation (figure 2 top row) perform worse than the ABR reference competitor (*w*^ *<* 0) when antibiotic is present. The Δ*waaP* single mutant (figure 2 top right facet) is the most susceptible to the antibiotic of all the investigated genotypes.

**Figure 2:**
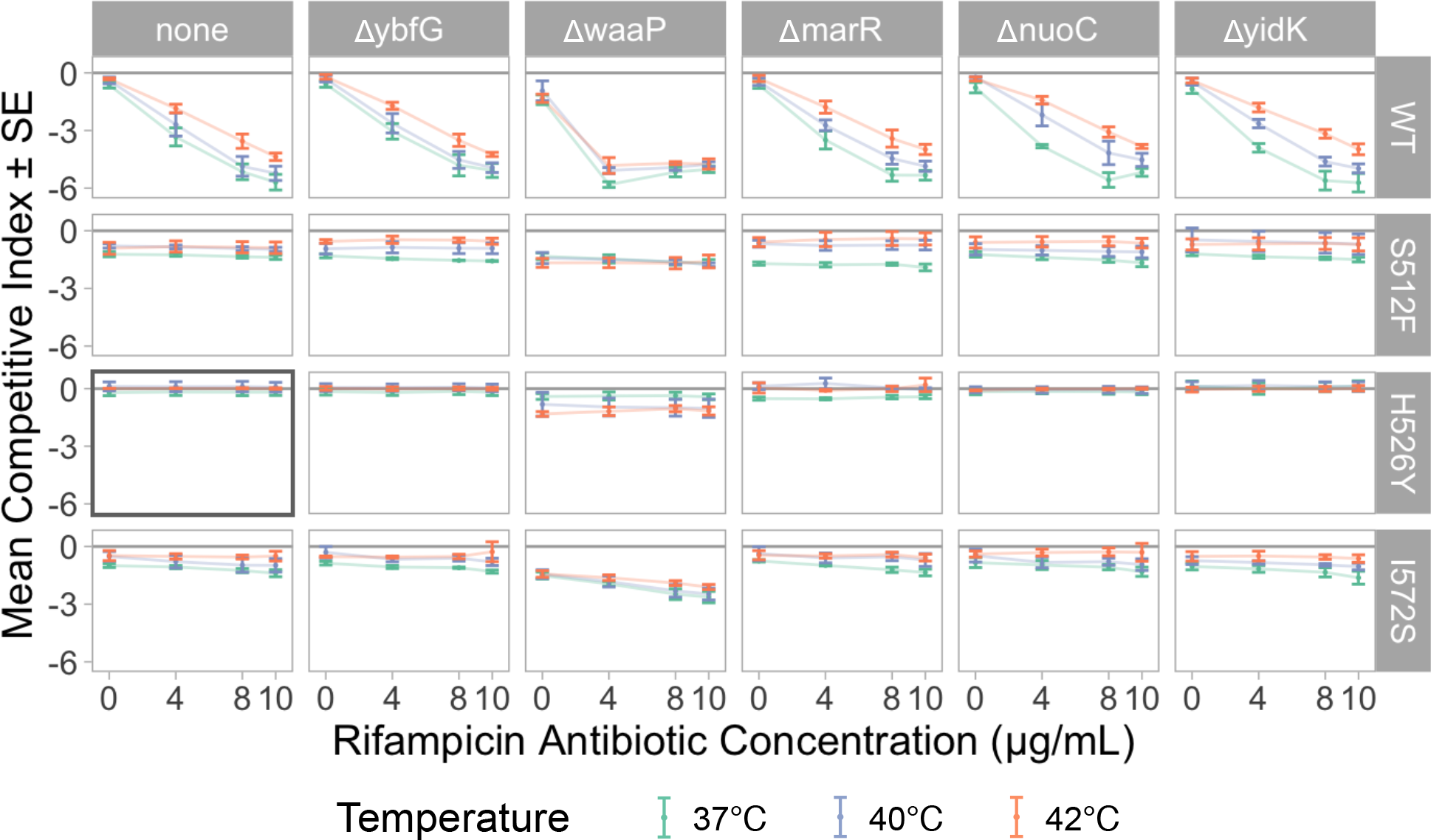
E**s**timated **competitive index (***w*^**) across all genotypes and environments.** The x-axis shows the four rifampicin antibiotic concentrations environments (*µg/mL*) and the y-axis shows the mean estimated competitive index with error-bars for standard errors (*n* = 3 *−* 4). Colours show the three temperature environments. Knock-out genotypes are shown in each column (with “none” indicating the wild-type like state of having no knock-outs) and *rpoB* antibiotic resistant mutants are shown in each row (with “WT” indicating the susceptible wild-type *rpoB* sequence); therefore the plot on the top left shows the wild-type without any mutations and, similarly, the left column shows all *rpoB* single mutants while the top row shows all knock-out single mutants. Finally, the grey box indicates the *rpoB* H526Y genotype used as the reference for all competitions. All experiments were performed in M9 liquid minimal media with 0.4% glucose. The MIC for rifampicin (*µg/mL*) in LB agar medium are as follows: WT *≈* 6 [79], 20 *<* I572S *<* 50, S512F *>* 100 [80], H526Y *>* 768 [79].

For all knock-out backgrounds, we compared the competitive fitness of the ABR mutations against the *rpoB* wild-type sequence in the absence of antibiotic to ascertain whether there was a cost of resistance. We observed no consistent cost of resistance across all backgrounds (figure S18) and (in contrast to previous studies [7, 36]) all three ABR mutations grew similarly to the wild-type at higher temperatures.

### 2.4 Multiple summary statistics found few G***×***G interactions

Several measures exist to determine the extent of G*×*G interactions (also termed epistasis). ‘Classical’ pairwise epistasis compares the observed mutational effect of the double-mutant against the additive expectation from each of the two single mutants (figure 1b-c). We inferred pairwise epistasis by using the fitness of the wild-type genotype as a baseline and subtracting the fitness effects of both ABR and knock-out single mutants from the fitness effect of the double mutant (figures S19 & S20 show examples without and with pairwise epistasis, respectively). We estimated pairwise epistasis for each of the 15 sets of double mutations in each of the 12 environments (figure S21). We found significant pairwise epistasis only for the Δ*waaP* mutant. Across all ABR backgrounds, the Δ*waaP* mutant achieved a maximum value of positive pairwise epistasis at the low antibiotic concentration (4*µ*g/mL). In the absence of antibiotic and at higher antibiotic concentrations (8 *−* 10*µ*g/mL), the Δ*waaP* mutant exhibited no significant pairwise epistasis on most of the ABR backgrounds (small negative pairwise epistasis was observed with some ABR backgrounds). This dose-dependent peak of the pairwise epistasis arises because the Δ*waaP* single mutant has a large deleterious fitness effect at 4*µ*g/mL and, when this knock-out is found in combination with an ABR mutation, the ABR mutation rescues bacterial growth (e.g., figure S20 second column). We detected almost no pairwise interactions of the Δ*waaP* mutant at higher antibiotic concentrations because the relative fitness effect of the Δ*waaP* mutation becomes much smaller as compared to the fitness effect of the ABR mutations (e.g., figure S20 third and fourth columns).

Second, we inferred epistasis over the whole landscape using the gamma statistic [47], which measures the correlation of fitness effects among the different genetic backgrounds where a mutation appears (figure 1b-c). Mutations that exhibit little G*×*G interaction have similar effects on different backgrounds; consequently, their effects will be strongly correlated across backgrounds (gamma near one). On the other hand, a weak or negative correlation of the effects of a mutation in different backgrounds (gamma near zero or negative) indicates that a mutation exhibits G*×*G interactions. Overall, we observed gamma epistasis values near one, which are indicative of very little G*×*G interaction. Antibiotic concentration was positively correlated with gamma epistasis (figure 3a; *F* (1, 10) = 52.7, *p <* 10*^−^*^4^, adjusted *R*^2^ = 0.825) regardless of temperature. This means that the amount of G*×*G interactions decreases as antibiotic concentration increases and that it is independent of temperature. Focusing solely on the gamma epistasis of ABR mutants or knock-outs, respectively (figure S22), we observed different qualitative but non-significant trends between the mutational classes and between the different temperatures. ABR mutations exhibited the strongest epistasis (i.e., gamma near zero) at the intermediate antibiotic concentration and increased with temperature. Conversely, knock-out mutations exhibited the strongest epistasis (again, gamma near zero) at the two highest antibiotic concentrations and decreased with temperature. Independent of the focal mutational class, there was less gamma epistasis at 42*^◦^C* than at 37*^◦^C*.

**Figure 3:**
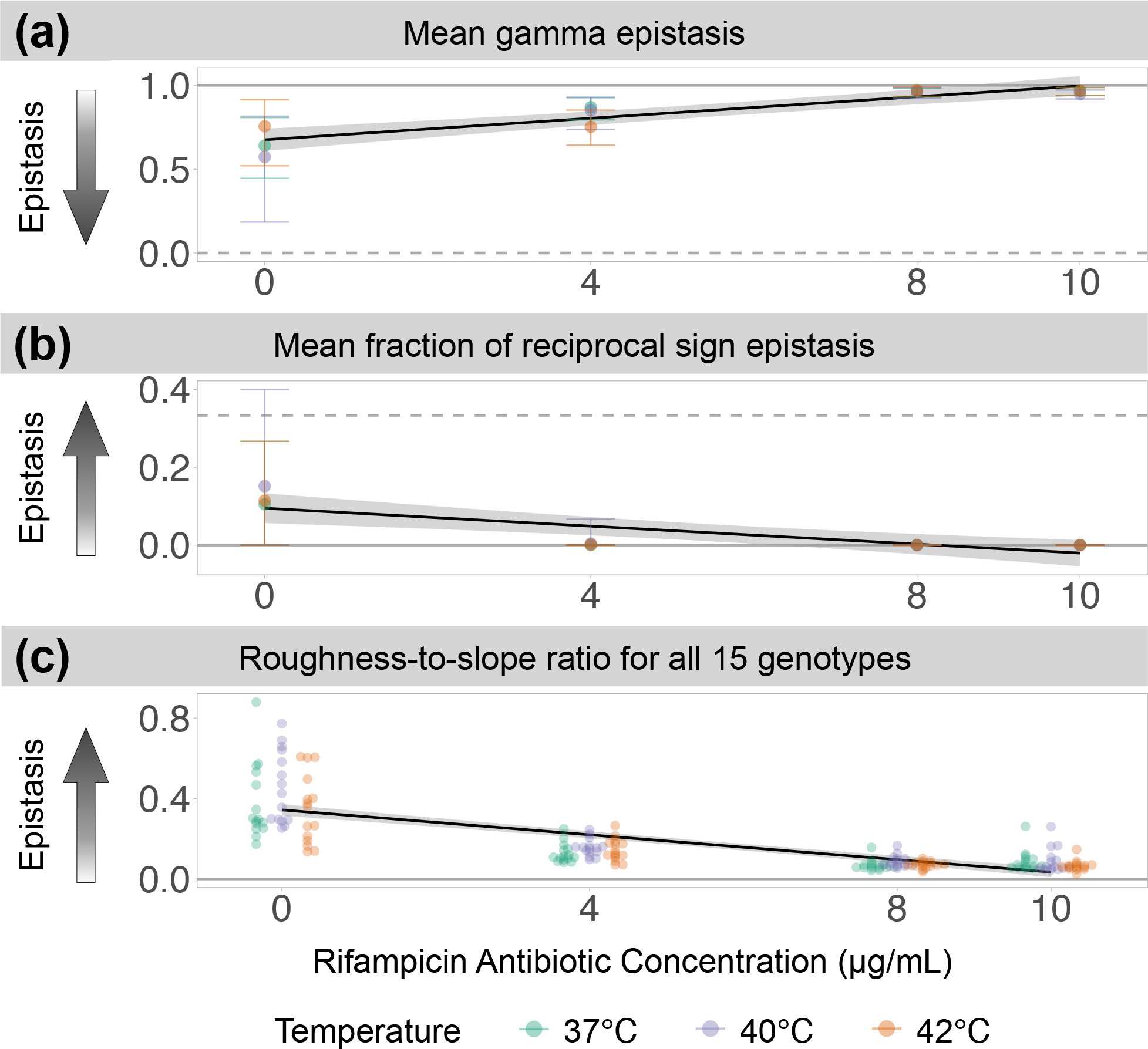
A**n**tibiotic **concentration has a significant negative trend with three measures of epistasis: (a)** the mean gamma epistasis across all loci, **(b)** the mean fraction of loci exhibiting reciprocal sign epistasis, and **(c)** the roughness-to-slope ratios by genotype. For each panel the x-axis shows the rifampicin antibiotic concentration in *µg/mL*, the colours show the temperature, and the y-axis shows the value of the epistasis measure. Arrows indicate the direction of stronger epistasis, solid horizontal lines show values of no epistasis, and dashed lines show the values of the maximum possible epistasis for each metric. The black trend-lines show the significant negative correlations between antibiotic concentration and each epistasis measure, and the grey shaded regions show the 95% confidence interval of the corresponding least-squares estimated linear regression. For panels (a)-(b), the error bars show the 95% parametric bootstrap confidence intervals. For figure clarity, bootstrap confidence intervals are omitted in panel (c) but can be found in figure S24, which shows the roughness-to-slope ratios for individual genotypes and environments. Significance is reported at an *α* = 0.01.

Third, we inferred the presence of simple sign and reciprocal sign epistasis (figure 1c). We identified simple sign epistasis as instances where the sign of the fitness effect of a mutation changed depending on its background in over five per cent of bootstrap samples. Then we identified reciprocal sign epistasis as instances where two mutations show sign epistasis on each other’s background. Importantly, the presence of reciprocal sign epistasis is a prerequisite for multiple peaks in the fitness landscape [48]. In our study, significant reciprocal sign epistasis occurred only in antibiotic-free environments but across all temperatures. This trend can be summarised as a negative correlation between antibiotic concentration and the mean fraction of reciprocal sign epistasis (figure 3b; *F* (1, 10) = 20.3, *p <* 0.005, adjusted *R*^2^ = 0.637). Reciprocal sign epistasis was observed between the weak ABR mutation *rpoB* I572S and most of the knock-out mutations (table S7). Apart from that, the fraction of simple sign epistasis was significant in all environments and showed no trend across environments (*F* (1, 10) = 2.15, *p* = 0.17, figure S23).

Next, we computed the roughness-to-slope ratio. The roughness-to-slope ratio fits a linear model to the fitness landscape and then quantifies epistasis as the ratio of the standard deviation of the residuals to the linear component of the fit (figure 1d). Thus, completely additive landscapes exhibit a roughness-to-slope ratio of zero [48]. In our data, the mean estimated roughness-to-slope ratio was negatively correlated with antibiotic concentration (figure 3c; *F* (1, 178) = 206.9, *p <* 10*^−^*^15^, adjusted *R*^2^ = 0.535). Roughness-to-slope ratios were very small for all genotypes in the presence of the antibiotic; nevertheless the 95% bootstrap confidence intervals were significantly different from zero for all values.

Finally, we inferred whether the studied fitness landscape exhibited diminishing-returns epistasis. Diminishing returns epistasis measures whether the effect size of a beneficial mutation decreases as the fitness of the genetic background increases and is one of the most frequently observed patterns of epistasis in fitness landscapes [2]. We observed that the fitness effects of the knock-out mutations were not correlated with the overall fitness of the background, regardless of the environment (figures 4a & c). In contrast, for most environments, the ABR mutations exhibited a negative correlation with the fitness of the background that the mutation is on, indicating a pattern of diminishing-returns epistasis (figures 4b & c). The observed trend of diminishing returns for the knock-out and ABR mutants did not depend on the low fitness of the Δ*waaP* mutants. Indeed, when the Δ*waaP* backgrounds were removed from the analysis, the trend became more pronounced as significant diminishing-returns epistasis was observed in all but one of the 12 environments (figure S25).

**Figure 4:**
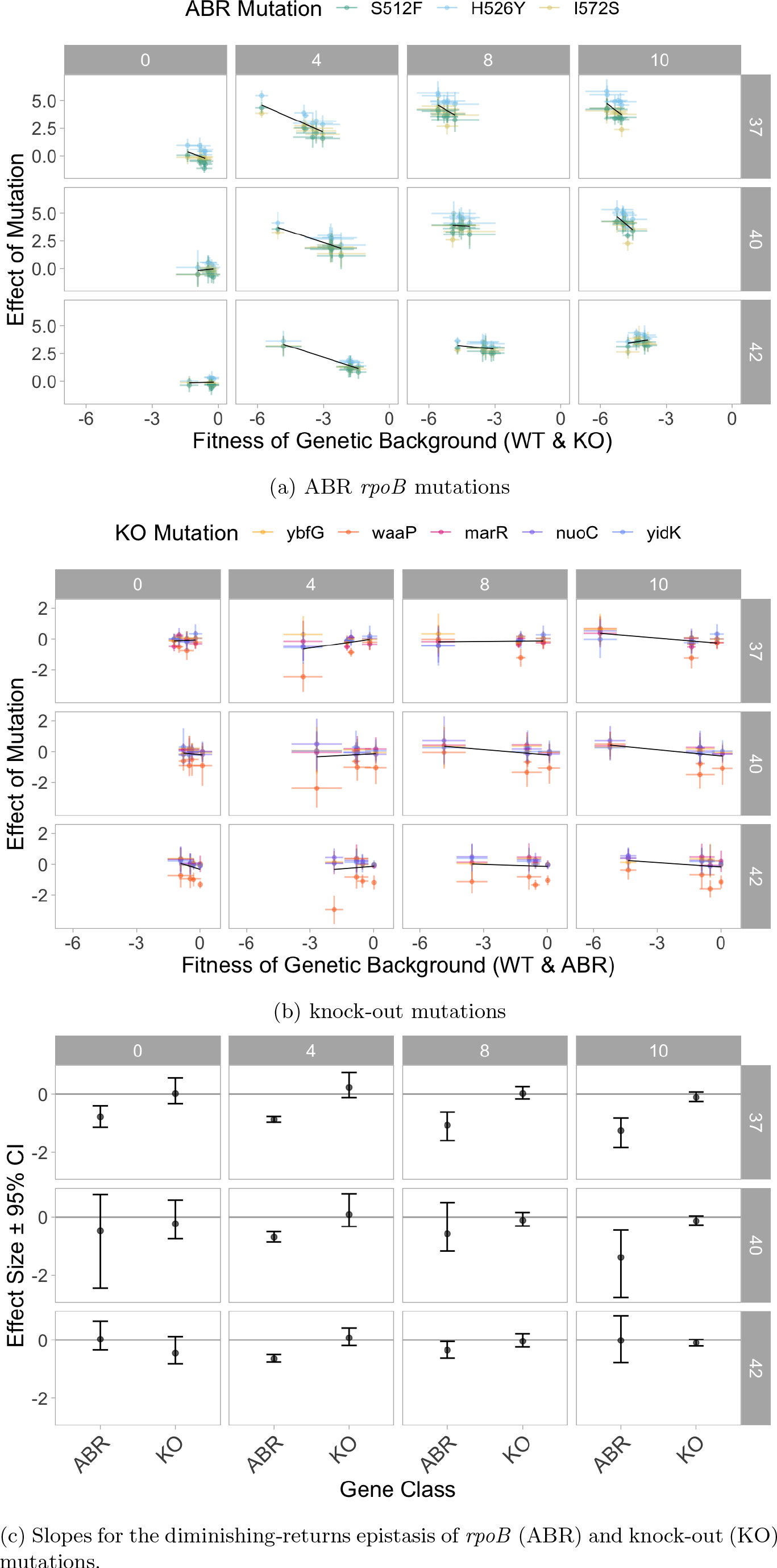
ABR *rpoB* mutations exhibit diminishing-returns epistasis in most environments whereas knock-out mutations never exhibit diminishing-returns epistasis. **In each panel, the** columns show different rifampicin antibiotic concentrations (in units of *µg/mL*) and the rows show different temperature environments (in *^◦^C*). **(a)** ABR *rpoB* mutations exhibit diminishing-returns epistasis in eight of the 12 environments. The x-axis shows the fitness *w*^ of the background (either wild-type or one of the five knock-out mutations) and the y-axis shows the effect size of the ABR mutation on that background. **(b)** Knock-out mutations do not exhibit diminishing-returns epistasis in any of the environments. The x-axis shows the fitness *w*^ of the background (either wild-type or one of the three ABR *rpoB* mutations) and the y-axis shows the effect size of the knock-out mutation on that background. For panels (a)-(b), the points show mean values and are coloured corresponding to the mutation; the error bars show 95% parametric bootstrap confidence intervals for each value. The black lines show the means of the regressions across all bootstrap samples. **(c)** Quantification of the mean slope effect sizes (*±* 95% confidence intervals) of the diminishing-returns epistasis for the *rpoB* mutations (ABR) and the knock-out mutations (KO). The mean slopes are shown as black trend lines in panels (a) & (b). Confidence intervals are determined from linear regressions of each of the parametric bootstrap samples. The confidence intervals that overlap with zero indicate slopes that are *not* significantly different from zero.

### 2.5 Competitive fitness is predicted by antibiotic concentration

We used multiple linear regression to study the quantitative effects of antibiotic concentration and temperature environments on the competitive index, *w*^. We found that the ‘null model’ regression with *w*^ as the dependent variable and only additive effects of genotype and environment variables was able to explain a considerable amount of the variation observed in the data (adjusted *R*^2^ = 0.68, *F* (13, 982) = 162.5, *p <* 10*^−^*^15^, table S8). Most of the residual variation was attributable to the wild-type *rpoB* genotypes (figure S26). Contrary to resistant genotypes, the antibiotic-susceptible genotypes interacted with the rifampicin environments by having higher fitness in the absence of rifampicin but very low fitness at higher rifampicin concentrations. Next, we quantified the interaction between the *rpoB* antibiotic susceptible and resistant genotypes with the antibiotic concentration by fitting a model with the additive effects and the *G_rpoB_ ×E*_AB_ interaction. This model fitted the data better than the additive null model, explaining almost all of the variation (adjusted *R*^2^ = 0.881, *F* (9, 973) = 186.0, *p <* 10*^−^*^15^, figure S27). Polynomial contrasts indicated that higher temperature was correlated with increased *w*^ (*t* = 15.0, *p <* 10*^−^*^15^, figures S28-S29). The fixed effect of antibiotic concentration alone on *w*^ was not significant (figure S28). Instead, the effect of antibiotic concentration was only significant as an interaction term with the susceptible *rpoB* wild-type genotype or the weak ABR genotype I572S. Then, we compared the above model with a slightly more complex model that included an additional *G_rpoB_ × G*_KO_ interaction between genotypes (i.e., ‘classical’ pairwise epistasis between *rpoB* and knock-out mutations),

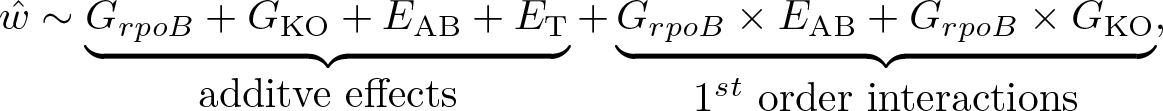

Although this model with G*×*E and G*×*G effects was significantly different from the model with only G*×*E effects (*F* (15, 958) = 2.60, *p <* 0.001, figure S30), none of the additional coefficients were significant (figure S31), and the model predictions were very similar to those of the model with only G*×*E. Indeed, the amount of variation explained by the more complex model (adjusted *R*^2^ = 0.883) showed only a 0.2 percentage-point improvement in the total variation explained as compared to the simpler model with only G*×*E effects. This finding confirms our previous results of small but statistically significant pairwise epistasis in our data.

Despite our relatively large dataset (n=996), we did not have sufficient data to systematically study second-order interactions without overfitting. We fitted models with second-order interactions for descriptive purposes (see supplementary results 8.6.3). In brief, we found that there may be a higher-order interaction between epistasis and temperature that our data has insufficient power to reveal statistically.

### 2.6 Environmental quality is less important for predicting G*×*G than antibiotic concentration alone

To study the interactions between epistasis and environment (G*×*G*×*E) without the problem of over- fitting, we summarised the temperature and antibiotic environments as a single, continuous environmental quality metric that we used to regress the different epistasis measures [49]. We compared three metrics of environmental quality (mean growth of the reference strain, mean growth of all competitor strains, and mean combined growth of reference and competitor strains) and found that the mean growth of all competitor strains in each environment was the best metric of environmental quality (supplementary results 8.7.1). This metric of environmental quality was able to distinguish environments from each other (figure S32) and correctly distinguish specialist from generalist genotypes (figure S33). As expected, the environmental quality metric was correlated with antibiotic concentration (adjusted *R*^2^ = 0.559) and somewhat correlated with temperature (adjusted *R*^2^ = 0.325).

We computed the correlation of environmental quality with all of the above measures of G*×*G by fitting the environmental quality metric as a linear predictor of pairwise epistasis (figures S37-S38), gamma epistasis (figure S39), the fraction of sign epistasis (figure S40), and the roughness-to-slope ratio (figure S41). Most of the regressions were either not significant to *α* = 0.01 (gamma epistasis for ABR mutations, simple sign epistasis, reciprocal sign epistasis, and pairwise epistasis excluding Δ*waaP*) or explained very little of the variation in the data (pairwise epistasis including Δ*waaP* : adjusted *R*^2^ = 0.036). For metrics that exhibited a significant effect of environmental quality on G*×*G (total gamma epistasis, figure S39, and roughness-to-slope ratio, figure S41), antibiotic concentration alone regressed as a dependent variable explained more of the variation in the G*×*G metric than was explained by environmental quality. Moreover, the results of the regression on environmental quality were broadly in agreement with our above findings that G*×*G decreases as antibiotic concentration increases and environmental quality deteriorates (i.e., positive correlations were found between the environmental quality metric and the G*×*G summary statistics). The only exception was the gamma epistasis for knock-out mutations (figure S39), which exhibited the opposite trend as other G*×*G metrics (i.e., G*×*G was found to increase as environmental quality deteriorated).

## 3 Discussion

Whether mutational effects and epistasis vary across environments, and how this matters for evolution, has been debated for decades [reviewed in 1, 2]. Here, we created 15 small (i.e., two mutational-step) fitness landscapes composed of three single amino-acid substitutions at a gene involved in ABR and whole gene knock-out mutations of five different genes. We screened these fitness landscapes at increasing gradients of temperature and an antibiotic. We found relatively little epistasis for four of the five different epistasis metrics. The benign environments exhibited the most epistasis, yet adding antibiotic decreased epistasis (figure 3). ABR mutations, but not gene knock-outs, exhibited diminishing-returns epistasis in most environments (figure 4). These results suggest that, while epistatic interactions may confound predictions in the absence of an antibiotic, the effects of ABR mutations are predictable in the presence of an antibiotic.

### 3.1 Biological significance of our experiment

We combined ABR mutations with gene knock-out mutations because the presence or absence of accessory genes is the most common standing genetic variation in natural populations of *E. coli* [50]. To our knowledge, this is the first study to systematically investigate how G*×*G interactions between SNPs and gene deletions change across different environments. We selected the gene knock-outs especially to avoid direct, mechanistic interactions with *rpoB* or other genes involved in transcription, protein synthesis, or DNA supercoiling. Therefore our null expectation was that there would not be any G*×*G interactions and that fitness would be additive throughout. If there was specific epistasis as a result of direct, mechanistic interactions between the ABR mutations and gene knock-outs, we expected to observe masking epistasis where the effect of the complete gene deletion is dominant over all of the ABR mutations. We did observe masking epistasis, but in the opposite direction than expected under specific epistasis: the ABR mutations dominated over the knock-outs. This demonstrates that the effects of single amino-acid substitutions at the *rpoB* locus are robust to non-core gene knock-out mutations. This result is an example of non-specific epistasis. Given its large population sizes and diverse pangenome composition, the core genes of *E. coli*, like *rpoB*, may have evolved to tolerate common polymorphisms like gene gain and loss events. We hypothesise that low pairwise epistasis between single amino-acid substitutions in core genes and knock-out mutations of non-core genes may generally be expected for *E. coli* and other prokaryotes with large pan-genomes [51].

We studied ABR mutations at the *rpoB* locus because this locus is clinically important and has been shown to exhibit G*×*G interactions. Amino-acid mutations at the *rpoB* locus (e.g., at site H526 included in our study) are the main, evolutionarily conserved mechanism of rifampicin resistance relevant for many bacterial pathogens [52, 53]. This includes the clinically relevant *Mycobacterium tuberculosis*, against which rifampicin treatment remains an important first-line antibiotic [52]. Previous studies have found that the fitness of *rpoB* mutants depends on the presence of mutations in other mechanistically interacting proteins, like other subunits of the RNA polymerase complex [54] and other protein complexes involved in protein synthesis and DNA stability [31, 55]. This type of ‘specific epistasis’ is easily explained by the close physical and functional proximity of these proteins to *rpoB*. ‘Nonspecific epistasis’, however, has been observed between genes and proteins that do not interact directly [26]. *RpoB* is an excellent candidate to investigate nonspecific epistasis as it was already shown to be highly pleiotropic: *rpoB* mutations impact the expression of hundreds of genes and many cell phenotypes [36, 56–58]. The demonstrated functional effect of *rpoB* upon many other genes implies that mutations at other loci could impact the fitness of ABR *rpoB* mutations. Therefore, the quantification of G*×*G interactions at the *rpoB* locus is important to know how the danger of ABR depends on the genetic background.

We detected a signal of pairwise epistasis in only one of five gene knock-outs, Δ*waaP* (figure S21). We had expected Δ*marR* to be the most likely knock-out mutation to exhibit pairwise epistasis, given its functional role in responding to chemical stressors and antibiotics [59, 42, 43] and its previously observed epistatic interactions with ABR mutations at genes involved in DNA stability [60]. Δ*marR* did not display significant pairwise epistasis in any environment; only Δ*waaP* did. The Δ*waaP* genotype on the susceptible *rpoB* wild-type background was more sensitive to low antibiotic concentration than the wild-type and other knock-out single mutants, but this sensitivity was masked in combination with any of the three ABR mutations. Gene deletions of other lipopolysaccharide biosynthesis genes were shown to exhibit negative pairwise epistasis with ABR mutations [61]. Δ*waaP* could make cells more sensitive to rifampicin by increasing the permeability of the outer membrane [62, 63]. Among the four functional genes studied, the *waaP* gene has the highest gene diversity (*π_waaP_* =0.034 vs *π_nuoC_* =0.015, *π_marR_*=0.014, *π_yidK_* =0.013) and is most frequently found knocked-out (*waaP* is present in 78.5% of strains as compared to 99.5%, 99.3%, and 90.8% for *nuoC, marR*, and *yidK*, respectively) in natural populations, according to the panX database. Moreover, *waaP* impacts bacterial virulence and, so, is relevant in host-pathogen interactions [64]. *E. coli* strains from which *waaP* is absent may be more sensitive to low doses of rifampicin and other antibiotics than *waaP* -carrying genotypes. Our observed pattern of epistasis implies that this sensitivity is masked when the genotype carries a resistance mutation. Thus, the G*×*G interaction of *waaP* and *rpoB* erases selection against Δ*waaP* mutations that would occur at low doses of rifampicin.

### 3.2 Measurement challenges of G***×***G interactions

One methodological challenge for detecting G*×*G interactions in our study was that, in the presence of antibiotics, the fitness effects of the ABR mutants quickly overshadowed those of the knock-outs. In this case, methods that involve a relative comparison of direct fitness effects to epistatic effects are bound to infer less epistasis when there are stronger direct effects. The negative correlation of the roughness-to-slope ratio as a function of antibiotic concentration could be explained by this phenomenon: we observed mostly neutral fitness effects and moderate epistasis in the absence of the antibiotic but large beneficial fitness effects and no epistasis in the presence of the antibiotic. However, the negative correlation between antibiotic concentration and epistasis was also exhibited by other epistasis summary statistics that are independent of the size of direct fitness effects (i.e., gamma epistasis and the mean fraction of reciprocal sign epistasis, figure 3). Therefore, the observed decrease of epistasis with antibiotic concentration is likely a real biological phenomenon.

Our estimates of gamma epistasis for different gene classes suggest that the fitness landscape as a whole was less epistatic than when only ABR mutations or knock-out mutations were considered. This could reflect a biological pattern (as supported by the low pairwise epistasis results) where the epistatic interactions between amino-acid mutations and gene knock-outs are smaller than within those mutational classes, perhaps due to evolved mutational robustness for gene knock-outs in *E. coli*. Nevertheless, the different trends observed between the gamma epistasis of the different mutational classes are possibly impacted by the differences in sample size. The gamma statistic relies on averaging over multiple mutational effects, which leads to a dilution of the epistatic signal for larger fitness landscapes. The total dataset has a larger sample size than subsets of the data, and the knock-out mutational class has a larger sample size than the ABR class. Future work should explore how the gamma epistasis measure can be used more appropriately for comparing epistasis between gene classes and differently sized fitness landscapes, for example by randomly subsampling the data to an equal size for all categories.

### 3.3 Environmental interactions & their importance

It has long been established that there is a G*×*E interaction between rifampicin resistance mutations at the *rpoB* locus and temperature [65], among other environmental factors [66, 18, 67]. In a previous experiment, [36] observed that *E. coli* adapted to high temperature evolved rifampicin resistance despite no rifampicin treatment. Therefore, we hypothesised that there would be an E*×*E interaction between high-temperature environments and rifampicin environments, which, if mediated by ABR mutations at the *rpoB* locus, could result in *G_rpoB_ × E_AB_ × E_T_* interactions as well. Overall we observed no cost of rifampicin resistance in minimal media, which is consistent with [68] and could be attributed to ABR *rpoB* mutations mimicking the stringent response [54]. There was a modest positive effect of high temperatures on the competitive fitness of all genotypes and a weak E*×*E interaction between temperature and antibiotic. However, contrary to previous studies that also used batch cultures with glucose and minimal medium [7, 36], the *rpoB* mutations S512F and I572S did not grow better and H526Y did not grow worse than the wild type at higher temperatures. (In fact, H526Y was observed to grow *better* than the wild type at 42*^◦^C*.) The discrepancy for I572S could be attributed to G*×*G*×*E with the genetic background: [36] found a strong effect of *E. coli* genetic background on the fitness at site I572. On the other hand, the discrepancies for S512F and H526Y could be that, unlike in [7], our batch cultures were not acclimatised to the environment in which the competitions occurred. This underscores the sensitivity of organisms to their environments and the importance of complete reporting of experimental methods.

We used the overall environmental quality as a regressor to quantify the interaction between epistasis and environment, G*×*G*×*E. We found that the antibiotic concentration alone was a better regressor of epistasis than the environmental quality, implying that temperature did not have an effect on epistasis. We had expected that the overall environmental quality would be a better regressor because it is highly correlated with antibiotic concentration while also taking into account the effects of temperature. On the other hand, fitting of linear models with higher-order interactions suggested that temperature exhibited a stronger interaction with *G_rpoB_ × G_KO_* than antibiotic concentration. Unfortunately, we have limited confidence in the linear regression results due to the low sample size and the high order of the investigated interactions. However, this apparent contradiction raises new questions about G*×*G*×*E interactions: for example, is it possible that one environmental variable affects direct G*×*E interactions, whereas the other acts on the G*×*G*×*E level? We are not aware of any previous work that has extracted such systematic patterns empirically, inferred their functional underpinnings, or developed models to account for such interaction. Certainly, the consequences of different interaction levels will be important to study in the future to predict potential evolutionary trajectories in varying environments.

### 3.4 Why is there so little epistasis in the studied ABR fitness landscapes?

Overall, we found little G*×*G and G*×*G*×*E interactions in our data. This result is in stark contrast to various recent experimental studies, which have presented strong evidence that both fitness effects of single mutations and their epistatic fitness interactions may vary greatly between environments [10, 19, 9, reviewed in 2]. One reason for this discrepancy may be the choice of mutations for the fitness landscapes. Previous studies focused on SNPs between or within genes that had been found in an adaptive walk [11, 4, 5], indirectly inferred strong G*×*G(*×*E) by observing that adaptations were unique to the genetic background [30, 69, 21], or measured epistasis as the different effect of ABR mutations across vastly different genetic backgrounds [70, 71]. We here quantified epistatic interactions between ABR SNPs and (non-ABR) gene knock-out mutations that occur in natural populations of *E. coli*. The studied combinations of mutations thus had no immediate relationship except the *rpoB* mutations’ previously characterised G*×*E interaction for fitness under higher temperature or antibiotic. However, multiple studies in systems or molecular biology have shown that epistasis is common [reviewed in 26]. In particular, when two or more beneficial mutations are combined, negative epistasis in the shape of diminishing returns (i.e., where the fitness effect of a mutation is less beneficial when it occurs on a fitter background) has been observed ubiquitously [72, 29, 61, 28], including for *rpoB* mutations conferring ABR to rifampicin [73]. In our study, we observed diminishing-returns epistasis only for the *rpoB* ABR mutations but no diminishing-returns epistasis for the knock-out mutations. Moreover, we did not observe any qualitative environmental trend in the strength of the diminishing-returns epistasis for the ABR mutations. However, the mutations shifted from neutral to beneficial as a function of antibiotic. Although there have been studies that derived null expectations of epistasis between random mutations of the same type [74], between mutations characterised by their fitness effect [75], or between SNPs in a pair of genes in a metabolic pathway [76], we are not aware of any general models or empirical studies that have proposed or quantified a null distribution of epistatic effects between SNPs and different structural mutation types (except for the assumption of no epistasis). Therefore, it is not clear whether the low G*×*G and G*×*G*×*E interactions observed in our data are to be expected.

Despite a lack of general epistatic null models, theoretical and empirical works have proposed a few hypotheses on how much epistasis to expect between mutations that confer ABR. For example, [34] found ubiquitous sign epistasis, in all 30 environments assayed, for a fitness landscape with all possible combinations of four ABR mutations. [31] found that epistasis was rare between combinations of ABR mutations. Our finding that *∼* 20% of randomly selected combinations of mutations exhibit pairwise epistasis is in good quantitative agreement with the results of [31]. A critical difference between our work and most previous work on ABR epistasis is that the knock-out mutations we investigated neither interacted directly with *rpoB* (as required for specific epistasis) nor, except Δ*waaP*, displayed any significant effects on fitness (figure S28). Epistasis expressed by seemingly neutral mutations is termed ‘cryptic epistasis’, and has been previously observed between mutations that were fixed during an adaptive walk [3, 5]. We have uncovered new instances of cryptic epistasis that depend on the environment. Using a theoretical model, [77] related the expected epistasis to the extent of G*×*E interactions of the involved mutations: they proposed that when there is no cost to the ABR mutations, there should be no epistasis between them. Moreover, a study by [13], considering only ABR mutations with a cost of resistance predicted, using mathematical models, and empirically observed the strongest epistasis at intermediate antibiotic concentrations. Our experiment screened a range of intermediate antibiotic concentrations, with 4 *µg/mL* below and 10 *µg/mL* near the minimum inhibitory concentration (MIC) of the wild-type *rpoB* (figure S14), and ABR mutations without a cost of resistance (although a cost had been expected according to previous studies, see above) in combination with non-ABR mutations. Our finding of very little epistasis in the presence of antibiotic is thus most consistent with the prediction of [77].

### 3.5 Do fitness landscapes become smoother as the concentration of an environmental stressor increases?

How generalisable are our findings that fitness landscapes become smoother in more stressful environments? To discuss this question, we compare our findings with those of [8], who measured fitness under different cadmium concentrations for all combinations of knock-out mutations that had evolved in response to those toxic environments. Contrary to our results, their co-selected mutations conferring resistance to increasing cadmium exhibited increasingly strong selective effects and positive pairwise epistatic effects as the heavy metal concentration increased [8]. Interestingly, both our results and those of [8] contradict theoretical predictions. According to Fisher’s geometric model [78], the average pairwise epistasis should be unchanged with environmental stress for random mutations, like the combinations of mutations used in our study, whereas it should decrease for co-selected mutations, like those studied in [8]. Also, our results and those of [8] differ from the prediction and empirical results of [13] that epistasis should be strongest at an intermediate concentration of an antibiotic stressor. However, the model of [13] was specific to increasing antibiotic (and not cadmium) stress and assumed a cost of resistance which was not observed in our study or in [8]. These conflicting results and predictions call for the study of additional empirical fitness landscapes under increasing concentrations of an environmental stressor, and new theoretical fitness landscape models that incorporate mechanistic details of biological phenomena.

Our main conclusion is that epistasis decreases with increasing antibiotic concentration. Only in the absence of the antibiotic, we observed several instances of reciprocal sign epistasis for all three ABR mutations and at all temperature environments (figure 3b, table S7). The presence of reciprocal sign epistasis implies that the fitness landscape has multiple peaks and that evolution may be less predictable in the absence of antibiotic. Overall, our results are consistent with the conclusion that the underlying fitness landscapes of ABR mutations and gene knock-outs are more rugged in the absence of antibiotic than in the presence of higher concentrations of antibiotic. Extrapolated to the larger sequence space, our results would imply that evolution is more predictable in the presence than in the absence of antibiotic, because ABR mutations have such strong beneficial effects in the presence of high antibiotic concentrations. Here, the strong effect of the ABR mutations potentially overrides any effects of the genetic background.

## 4 Data Availability

All flow cytometry data, competitive fitness estimates, and the annotated code used to generate the analyses are publicly available at the following Git repository: https://gitlab.com/evoldynamics/epistasis-decreases-with-increasing-antibiotic-pressure and will be archived on Zenodo upon publication of the manuscript. The whole-genome sequencing is archived on NCBI with the BioProject accession: PRJNA910115.

## Supporting information

electronic supplement

## Acknowledgements

We thank the Flow Cytometry Facility of Instituto Gulbenkian de Cîencia and the Next Generation Sequencing (NGS) Platform of UBern for their services and assistance. AHG acknowledges funding from FCT PhD funding grant PD/BD/138215/2018. CB acknowledges funding from ERC Starting Grant 804569 (FIT2GO), HFSP Young Investigator Grant RGY0081/2020, and SNSF Project Grant 315230 204838/1 (MiCo4Sys). IG and CB acknowledge funding from FCT PREPARE project (JPIAMR/0001/2016-ERA-NET). We are grateful to the members of the Evolutionary Biology group for *E. coli* strains (R. Balbontín & Durão), training in microbiology (D. Guleresi), and consultation on methods, and to the members of the Evolutionary Dynamics/Theoretical Ecology and Evolution group for help with preliminary environmental screenings (C. Diwo), genetic engineering (M. Schmitz), consultation on WGS (A. Kapopoulou), and discussions. We thank T. Batallion, S. Yeaman, F. Blanquart, and A. Wagner for feedback on results; and the PREPARE consortium for ideas and help with experimental design.

## 8 Supplementary results

### 8.1 Sample confirmation by whole genome re-sequencing

The genotypes were verified by Illumina whole-genome sequencing to just over 115-fold read-depth on average and consistent coverage throughout the reference genome/sequences (see tables S1-S2 and representative figures S1-S9, S11). Sequencing results suggested that the double-mutant *rpoB* S512F Δ*ybfG* (AHG100) was contaminated by the preceding double-mutant sample from our library, *rpoB* S512F Δ*waaP* (AHG098; evidence shown in figures S7, S9d, S10), at a near 50% population frequency based on the relative read depth of the new junction evidences (figure S12). PCR of the potentially contaminated *rpoB* S512F Δ*ybfG−*80*^◦^C* stock culture and the DNA extraction sent for sequencing failed to amplify the contaminating Δ*waaP*construct (figure S13). Therefore we assume that the contamination happened during sequencing and results from *rpoB* S512F Δ*ybfG* are included in all analyses. The complete list of mutations identified by whole-genome re-sequencing as compared to the wild-type genotype (sample AHG015) is given in table S3. Three genotypes (the double-mutant *rpoB* I572S Δ*nuoC*, the single-mutant *rpoB* S512F, and the single-mutant Δ*yidK*) had mutations in coding sequence regions as compared to the wild-type.

Of the 25 genotypes, three were found to have *de novo* mutations that may impact our results. The anticipated effects of these *de novo* mutations are as follows: *I.* The double-mutant *rpoB* I572S Δ*nuoC* had two non-conservative mutations, N135K in the phage superinfection exclusion protein *sieB* and D263Y in the respiratory enzyme *glpD*. We anticipate that the first mutation would not have any functional effects in our study as phages are dormant in the MG1655 strain. Similarly, we anticipate little functional effects from the second mutation since the gene is involved in glycerol metabolism and glycerol is absent from our experimental media. *II.* The single-mutant *rpoB* S512F had a very conservative amino-acid mutation (A32G) in a gene of unknown function (*yaeC*); we anticipate little functional effects for this mutation. *III.* Finally, the single-mutant Δ*yidK* had an in-frame 9 bp mobile element insertion in a putative endonuclease involved in DNA recombination and Type VI secretion (*yhhZ*). We cannot anticipate what the effect of this mutation would be in our experiment.

It is unlikely that the *de novo* mutations of the aforementioned three genotypes impact the main results of our experiment. If these *de novo* mutations had significant effects on the fitnesses of the genotypes they are found on, we might expect to detect this as pairwise epistasis because each *de novo* mutation occurred on a different background. However, none of the genotypes above was found to have a different pattern of pairwise epistasis as compared to other backgrounds.

**Table S1:**
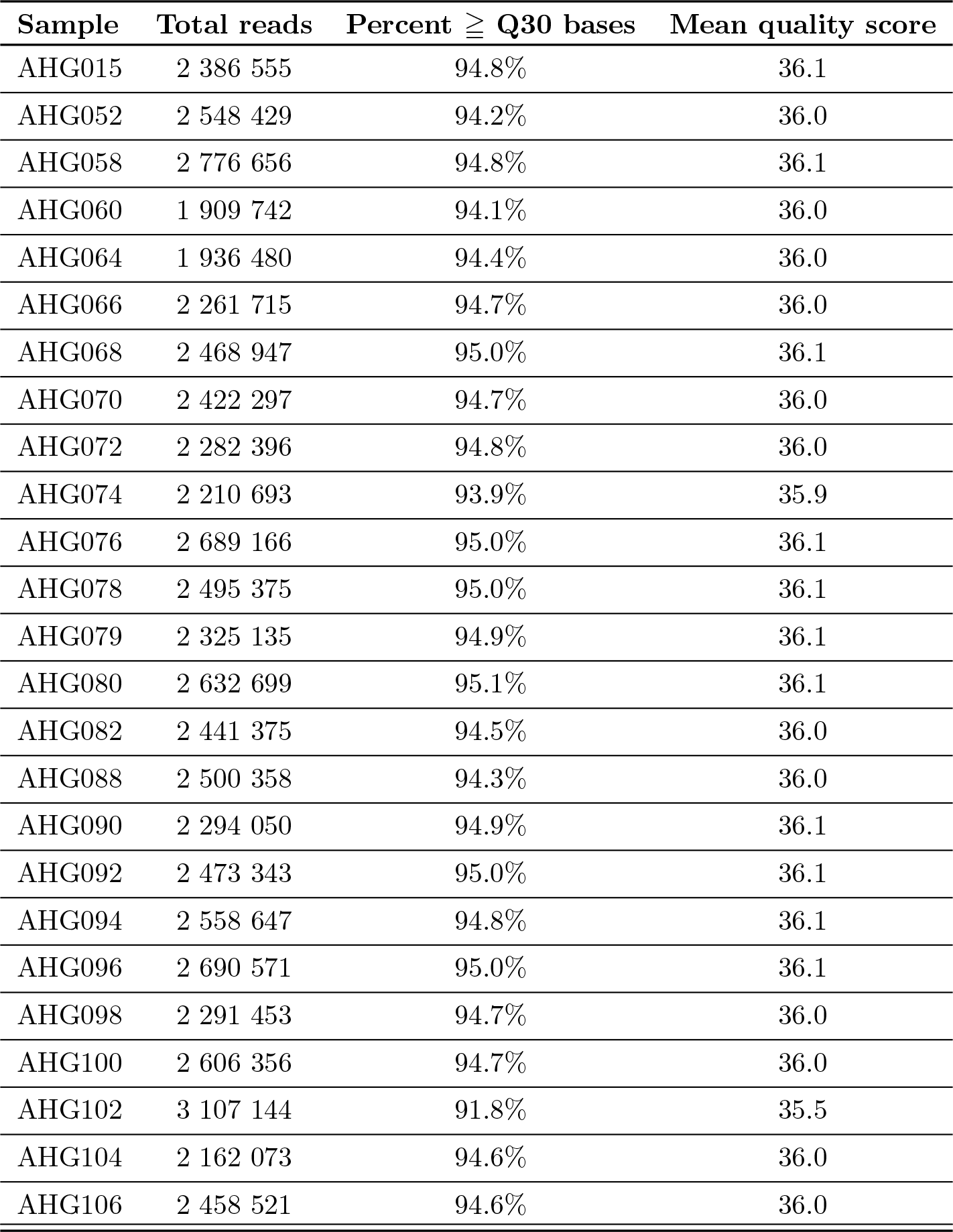
Summary of Illumina sequencing quality for all 25 genotypes used in competitions. (Engineered mutations are listed in table S12.)

**Table S2:**
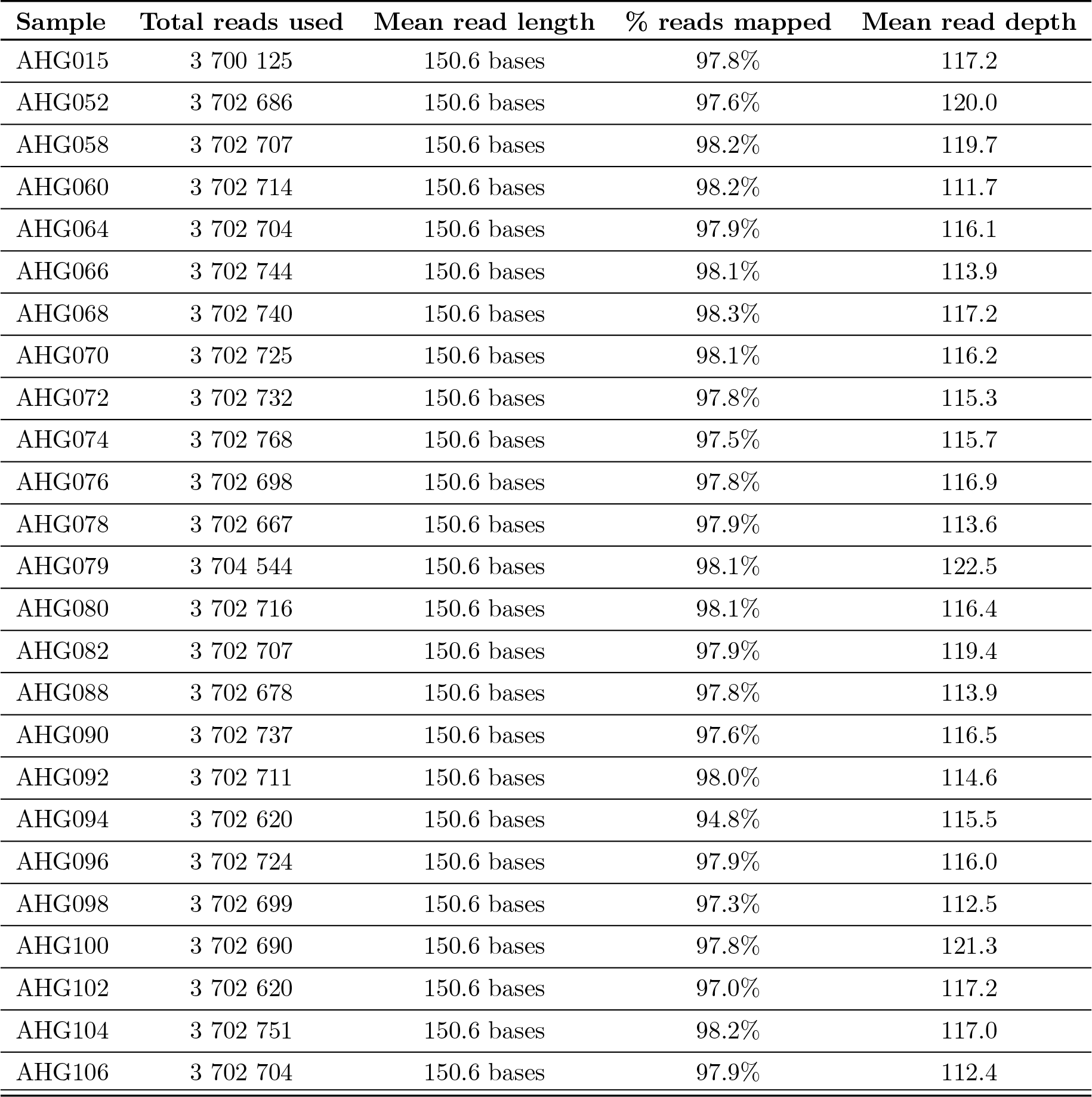
Summary of breseq sequence mapping quality for all 25 genotypes used in competitions. Total reads used is less than total reads sequenced from table S1 since an average read depth limit of 120x was set, as recommended in the breseq manual. Mean read depth is the average across all reference sequences (not weighted by reference sequence length). Engineered mutations are listed in table S12.

**Figure S1:**
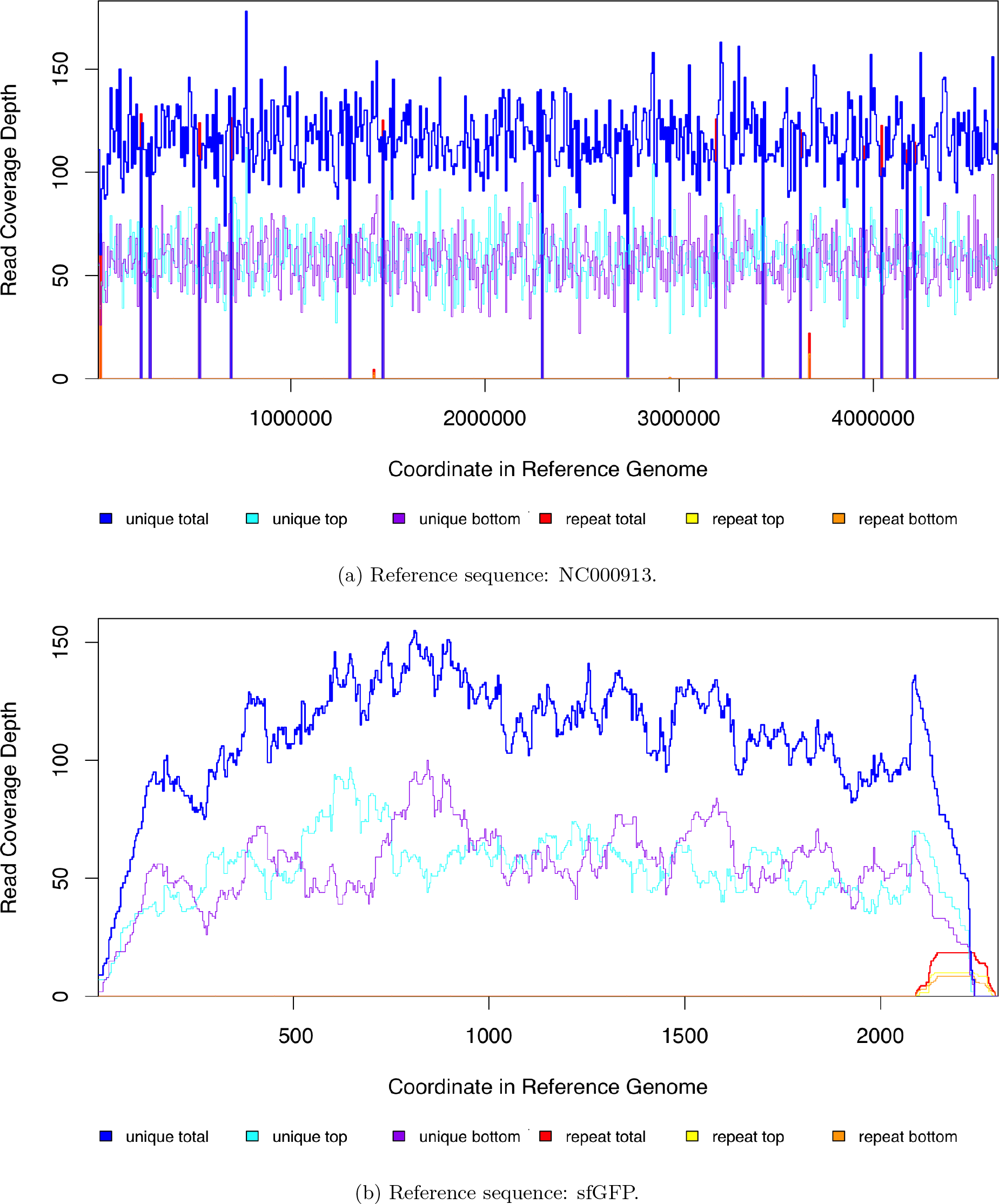
Breseq read coverage plots of all reference genomes used for wild-type sample (AHG015). a) ‘NC000913’ is the MG1655 reference sequence from GenBank (NC 000913.3) and b) the sequence for the sfGFP construct is included in the supplementary methods.

**Figure S2:**
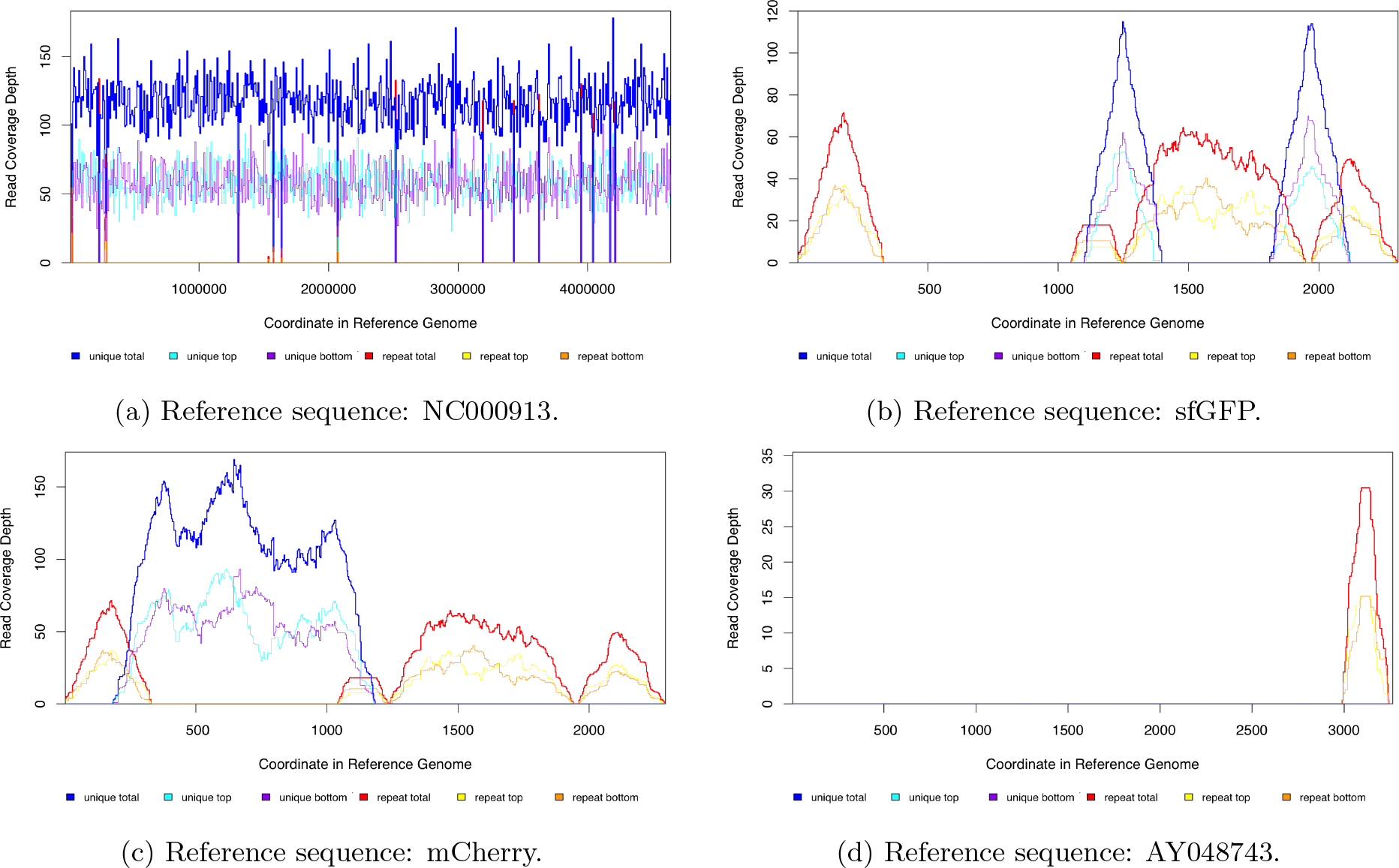
Breseq read coverage plots of all reference genomes used for the *rpoB* H526Y sample that was used as the common, mCherry-labeled reference in all competitions (AHG079). a) ‘NC000913’ is the MG1655 reference sequence from GenBank (NC 000913.3). b) There are no unique reads mapped to the sfGFP coding sequence on the construct because sample AHG079 has c) the mCherry construct instead (both reference sequences are included in the supplementary methods). d) AY048743 is the pDK4 plasmid sequence from GenBank (AY048743.1). Unique reads do not map to this sequence because there is no engineered gene knock-out (nor kanamyacin cassette) present.

**Figure S3:**
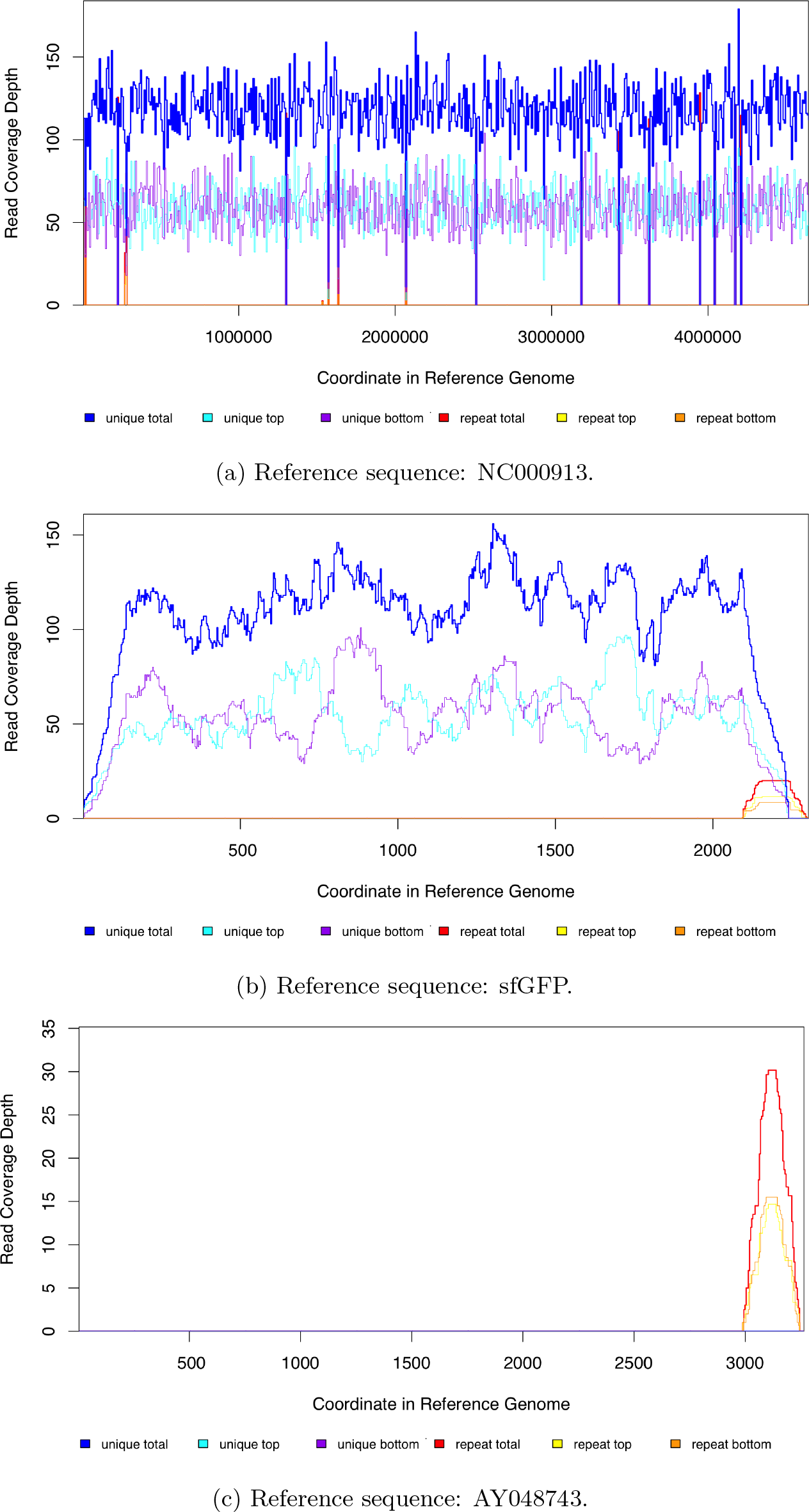
Breseq read coverage plots of all reference genomes used for sample AHG068. a) ‘NC000913’ is the MG1655 reference sequence from GenBank (NC 000913.3). b) The sequence for the sfGFP construct is included in the supplementary methods. c) AY048743 is the pDK4 plasmid sequence from GenBank (AY048743.1). Unique reads do not map to this sequence because there is no engineered gene knock-out (nor kanamyacin cassette) present.

**Figure S4:**
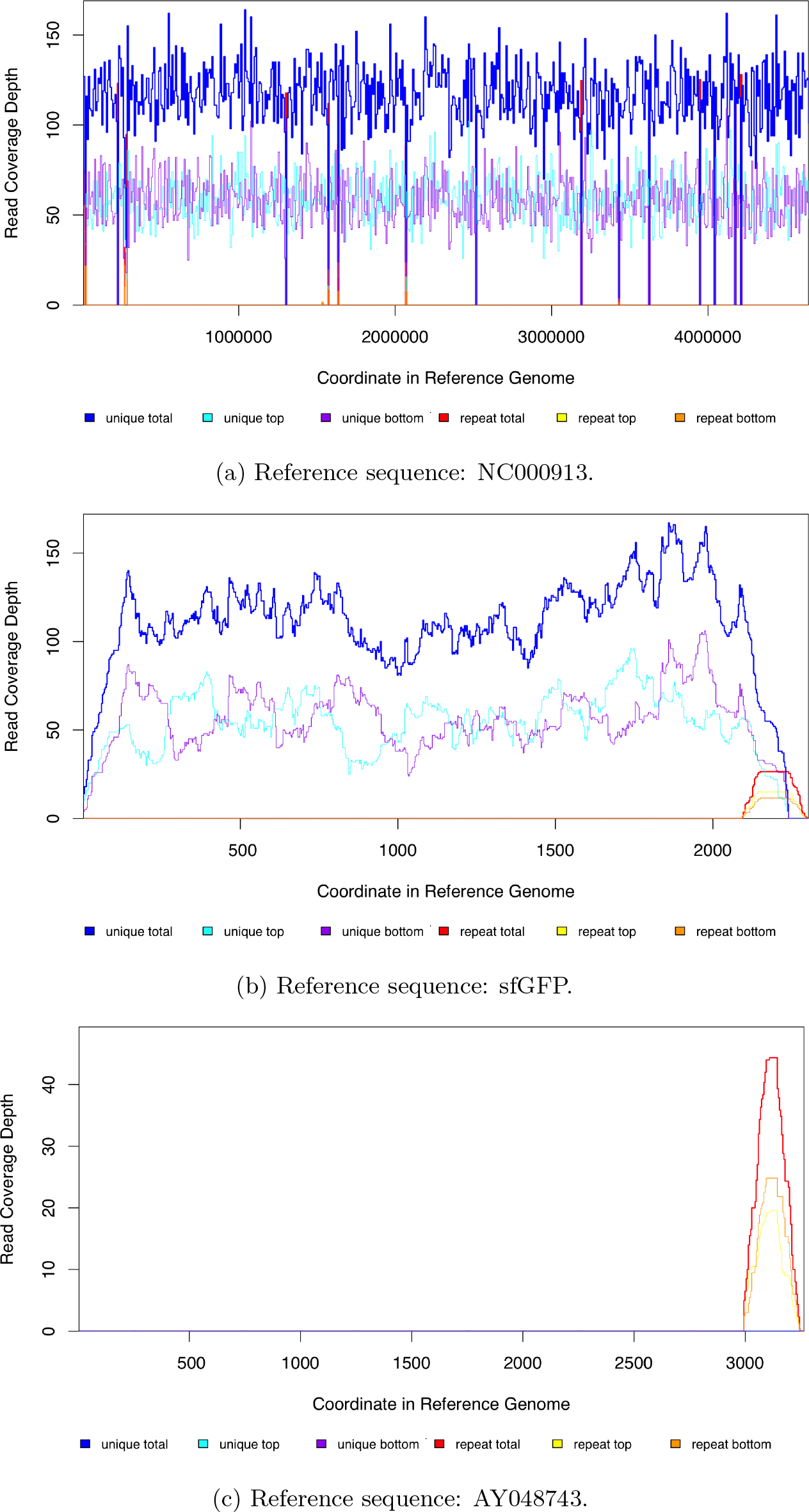
Breseq read coverage plots of all reference genomes used for sample AHG080. a) ‘NC000913’ is the MG1655 reference sequence from GenBank (NC 000913.3). b) The sequence for the sfGFP construct is included in the supplementary methods. c) AY048743 is the pDK4 plasmid sequence from GenBank (AY048743.1). Unique reads do not map to this sequence because there is no engineered gene knock-out (nor kanamyacin cassette) present.

**Figure S5:**
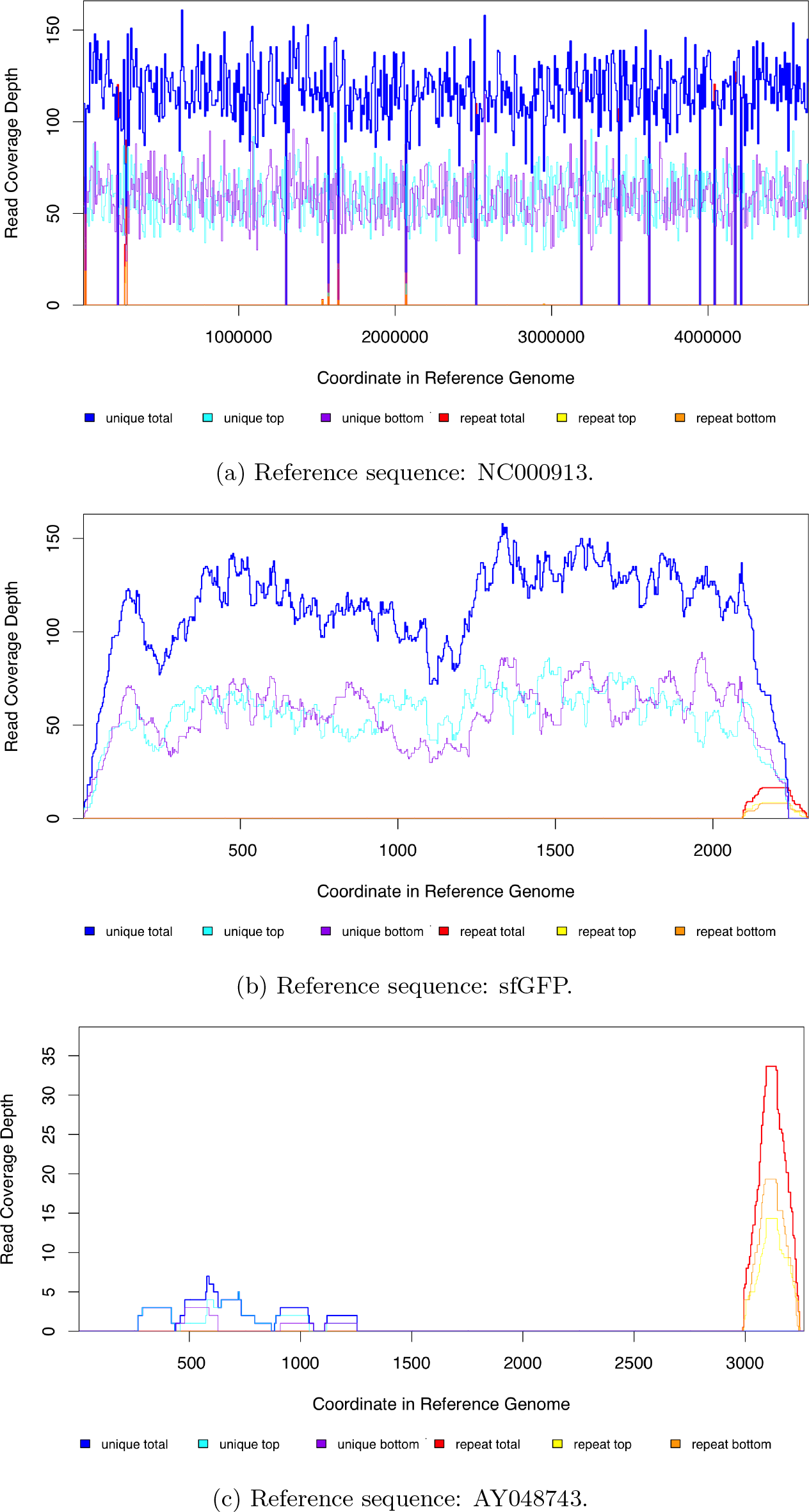
Breseq read coverage plots of all reference genomes used for sample AHG082. a) ‘NC000913’ is the MG1655 reference sequence from GenBank (NC 000913.3). b) The sequence for the sfGFP construct is included in the supplementary methods. c) AY048743 is the pDK4 plasmid sequence from GenBank (AY048743.1). Unique reads do not map to this sequence because there is no engineered gene knock-out (nor kanamyacin cassette) present.

**Figure S6:**
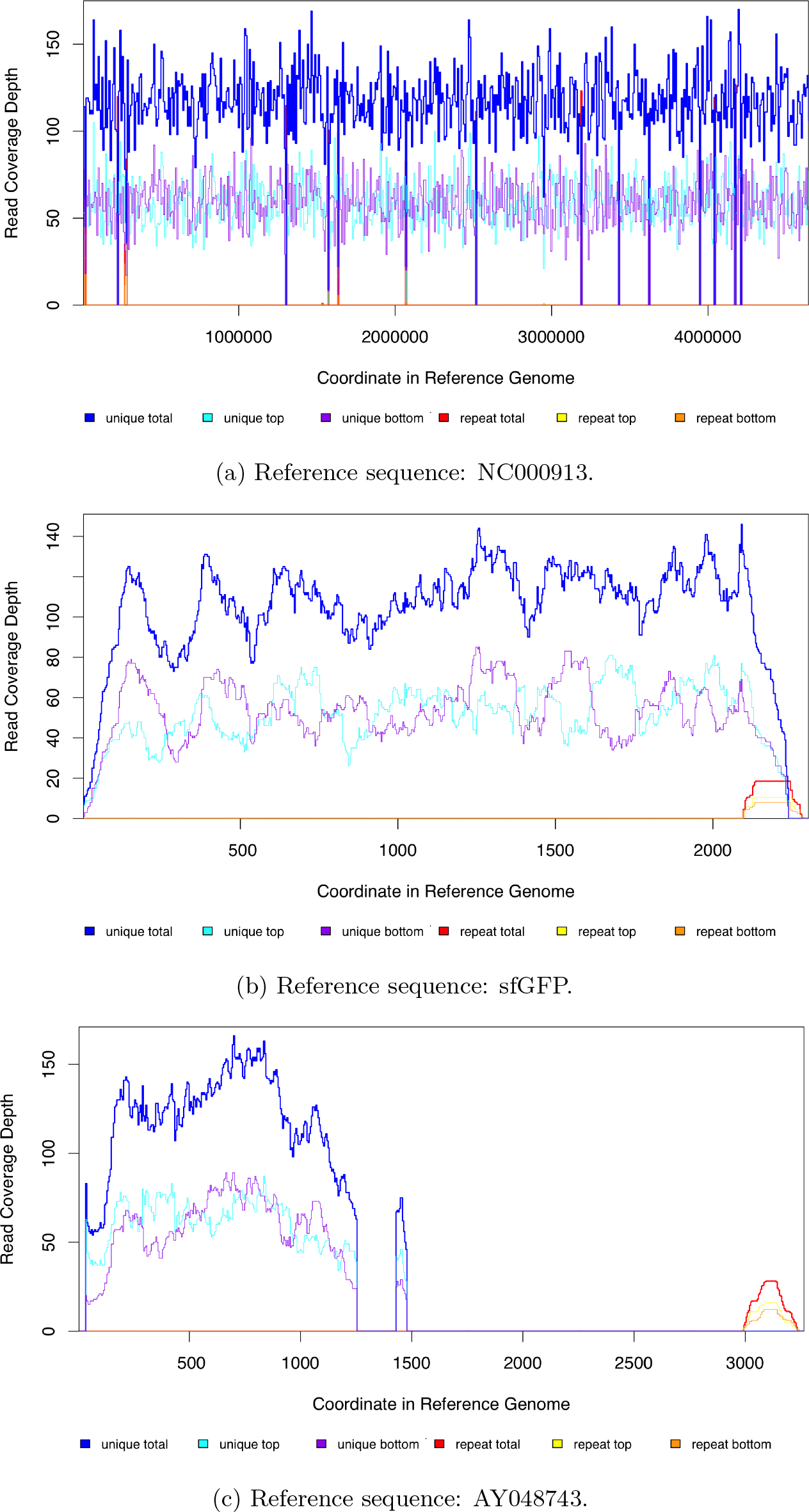
Breseq read coverage plots of all reference genomes used for sample AHG052. This sample is represenative of all coverage plots. a) ‘NC000913’ is the MG1655 reference sequence from GenBank (NC 000913.3). b) The sequence for the sfGFP construct is included in the supplementary methods. c) AY048743 is the pDK4 plasmid sequence from GenBank (AY048743.1). Reads map to this sequence because there is an engineered gene knock-out created by swapping the gene of interest with a kanamyacin cassette.

**Figure S7:**
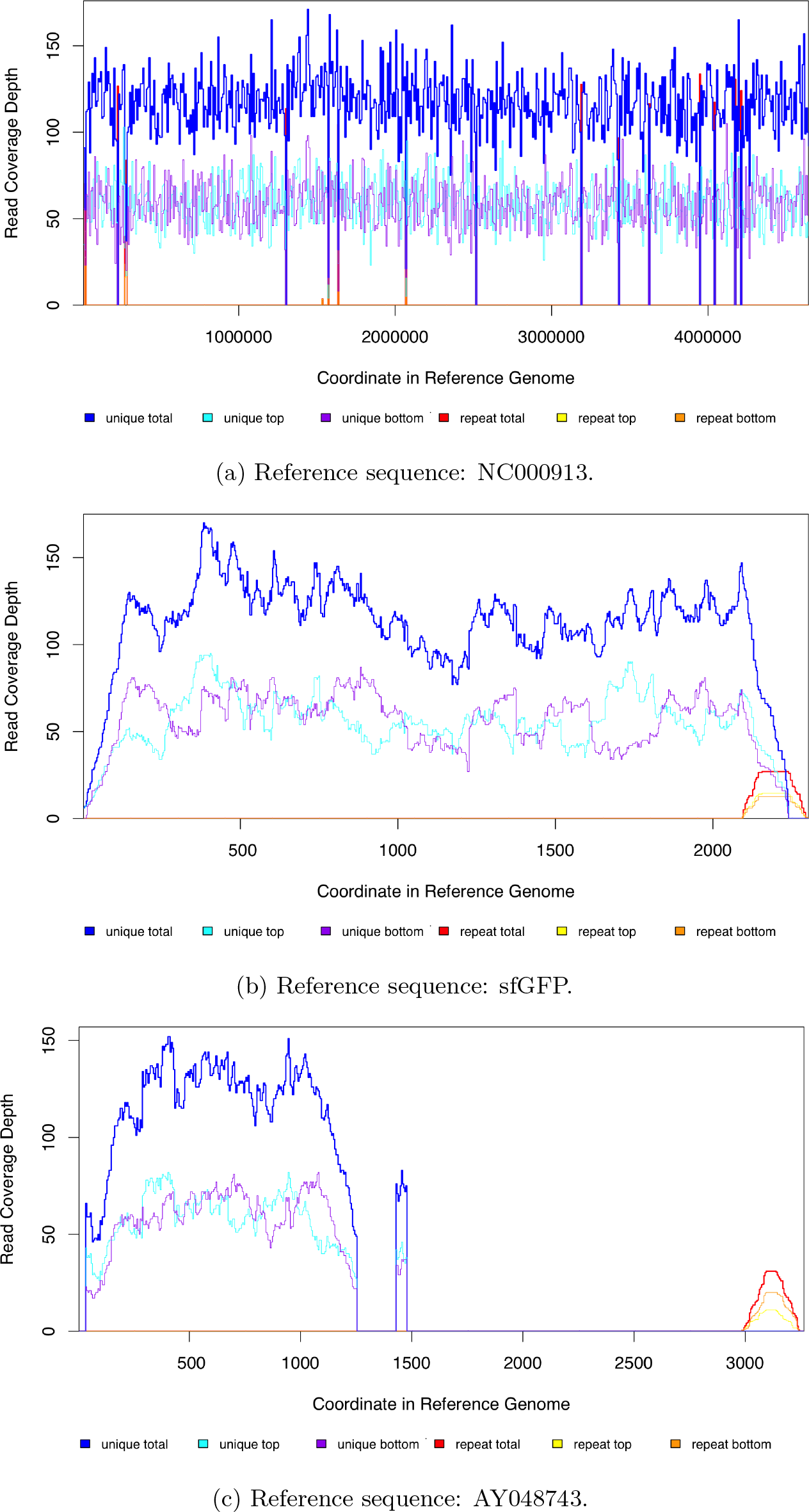
Breseq read coverage plots of all reference genomes used for sample AHG100. This sample is represenative of all coverage plots. a) ‘NC000913’ is the MG1655 reference sequence from GenBank (NC 000913.3). b) The sequence for the sfGFP construct is included in the supplementary methods. c) AY048743 is the pDK4 plasmid sequence from GenBank (AY048743.1). Reads map to this sequence because there is an engineered gene knock-out created by swapping the gene of interest with a kanamyacin cassette.

**Figure S8:**
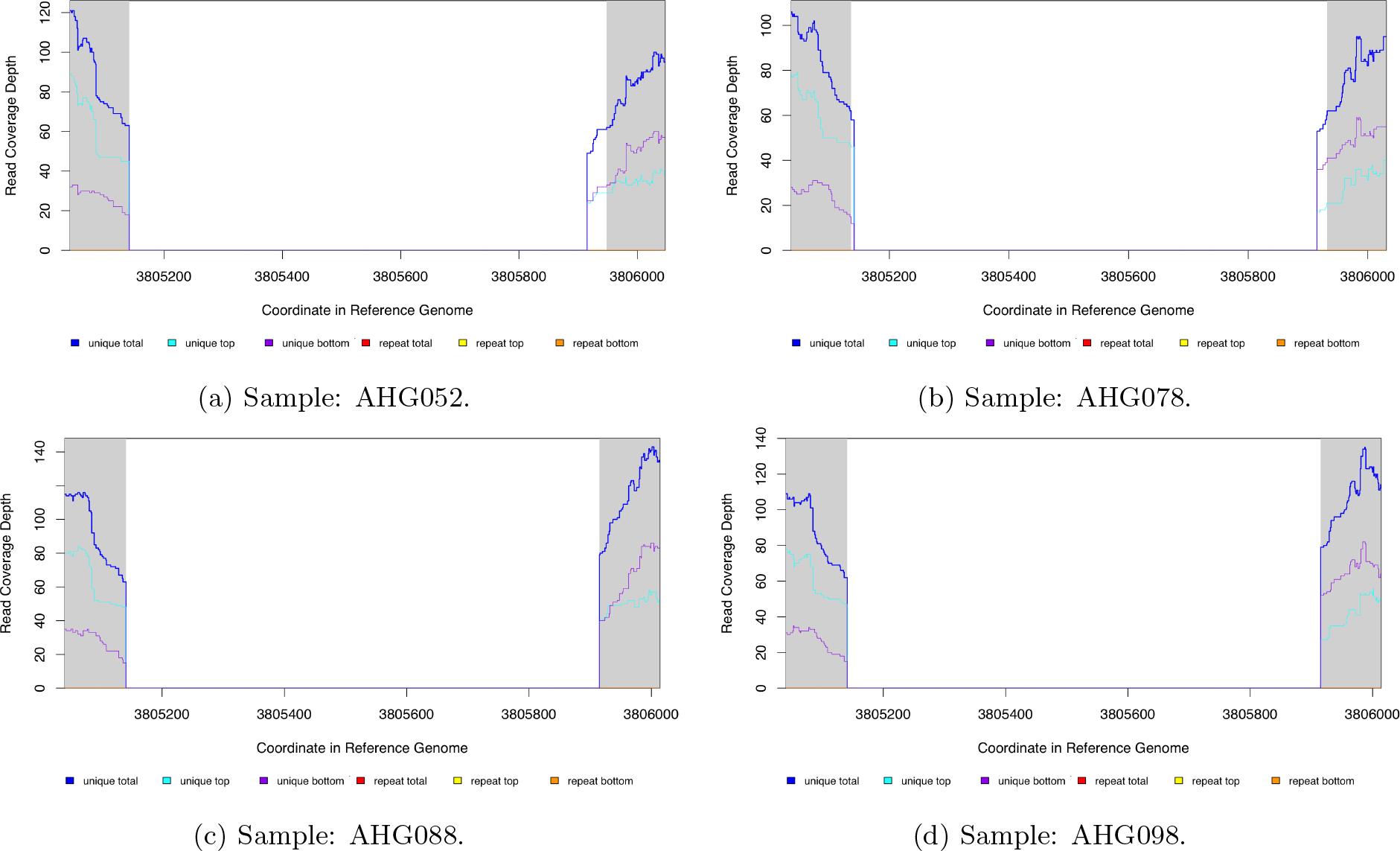
Breseq read coverage plots for the region of the NC 000913.3 reference sequence around the gene *waaP* for the four samples engineered to have a knock-out at this gene: a) AHG052, b)AHG078, c) AHG088, and d) AHG098. The white background on the plot shows the region detected by breseq as having a reduced read coverage as compared to the regions with grey background.. No reads map to the *waaP* coding sequence because a knock-out has been engineered at this region by replacing the gene coding sequence with a kanamyacin cassette. Only samples with Δ*waaP* are included as representative since the coverage plots at the knock-out sites for the other genes and samples are similar.

**Figure S9:**
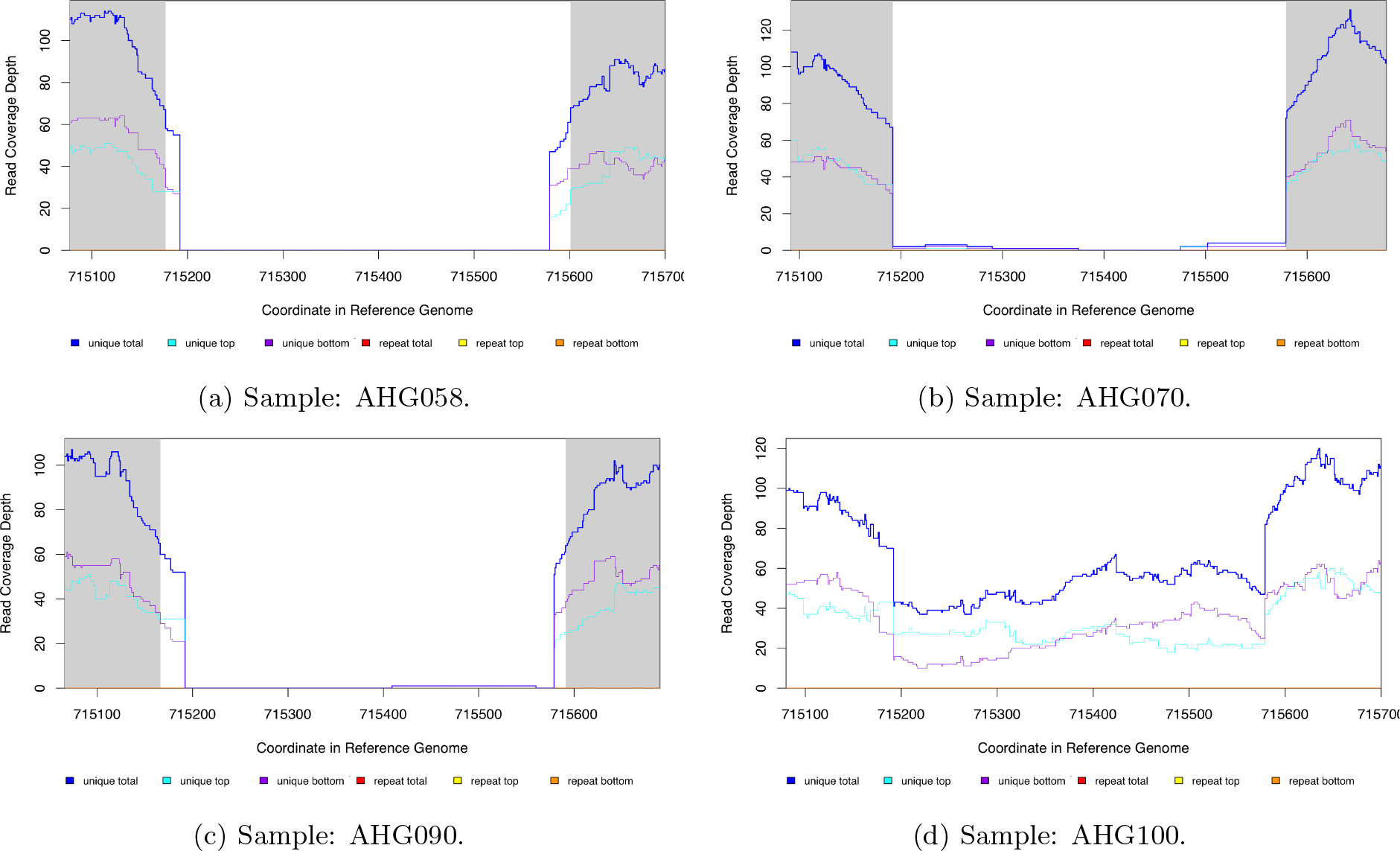
Breseq read coverage plots for the region of the NC 000913.3 reference sequence around the pseudogene *ybfG* for the four samples engineered to have a knock-out at this locus: a) AHG058, b)AHG070, AHG090, and d) AHG100. In a-c), no reads map to the *ybfG* putative coding sequence because a knock-out has been engineered at this region by replacing the pseudogene sequence with a kanamyacin cassette. However, for d) there has likely been contamination by sample AHG098 (Δ*waaP*, *rpoB* S512F) into sample AHG100 (Δ*ybfG*, *rpoB* S512F) and so reads are mapping to the region around *ybfG* (see also read coverage plot in figure S10 and read junction evidence in figure S12). The white background on the plots for panels a-c) shows the region detected by breseq as having a reduced read coverage as compared to the regions with grey background. Panel d) was created manually at the *ybfG* locus using the breseq BAM2COV command and so there is no background shading to the plot.

**Figure S10:**
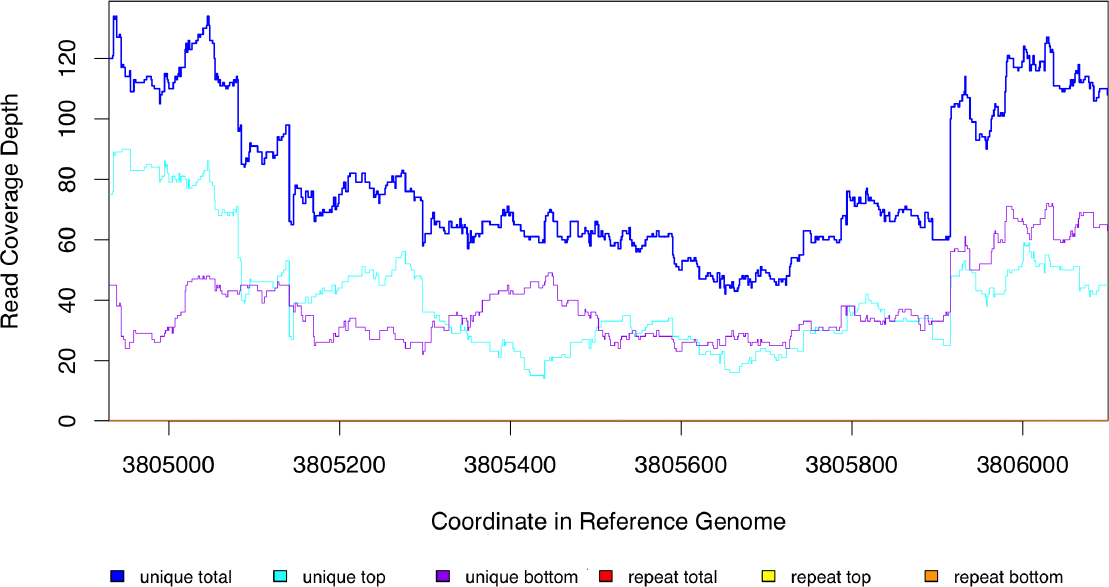
Breseq read coverage plots for the region of the NC 000913.3 reference sequence around the gene *waaP* (3805140 - 3805915) for the AHG100 samples that is **not** engineered to have a knock-out at this site. There has likely been contamination by sample AHG098 (Δ*waaP*, *rpoB* S512F) into sample AHG100 (Δ*ybfG*, *rpoB* S512F) and so there is decreased read coverage at the *waaP* locus as a result (see also read coverage plot in figure S9d and read junction evidence in figure S12).

**Figure S11:**
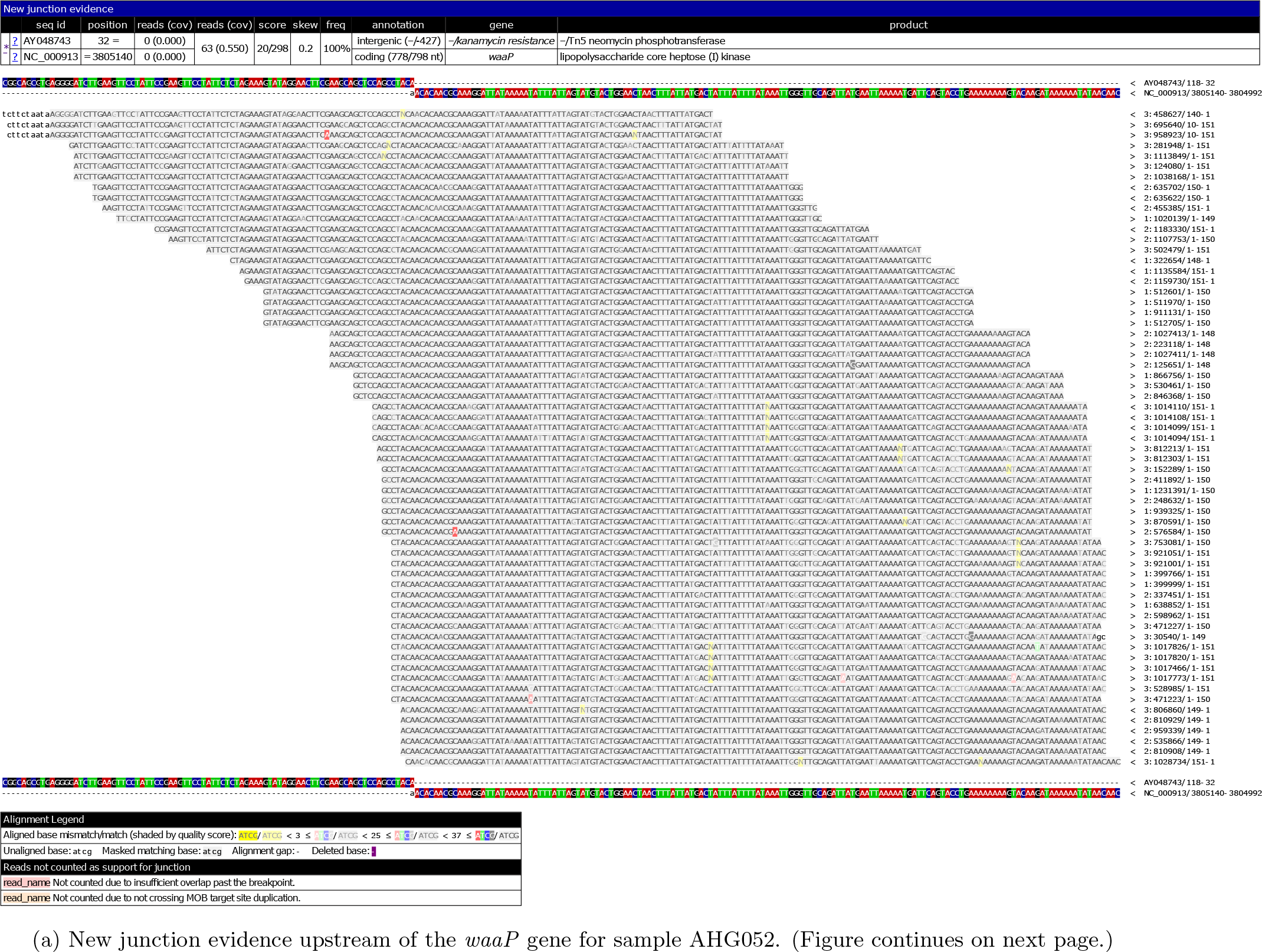
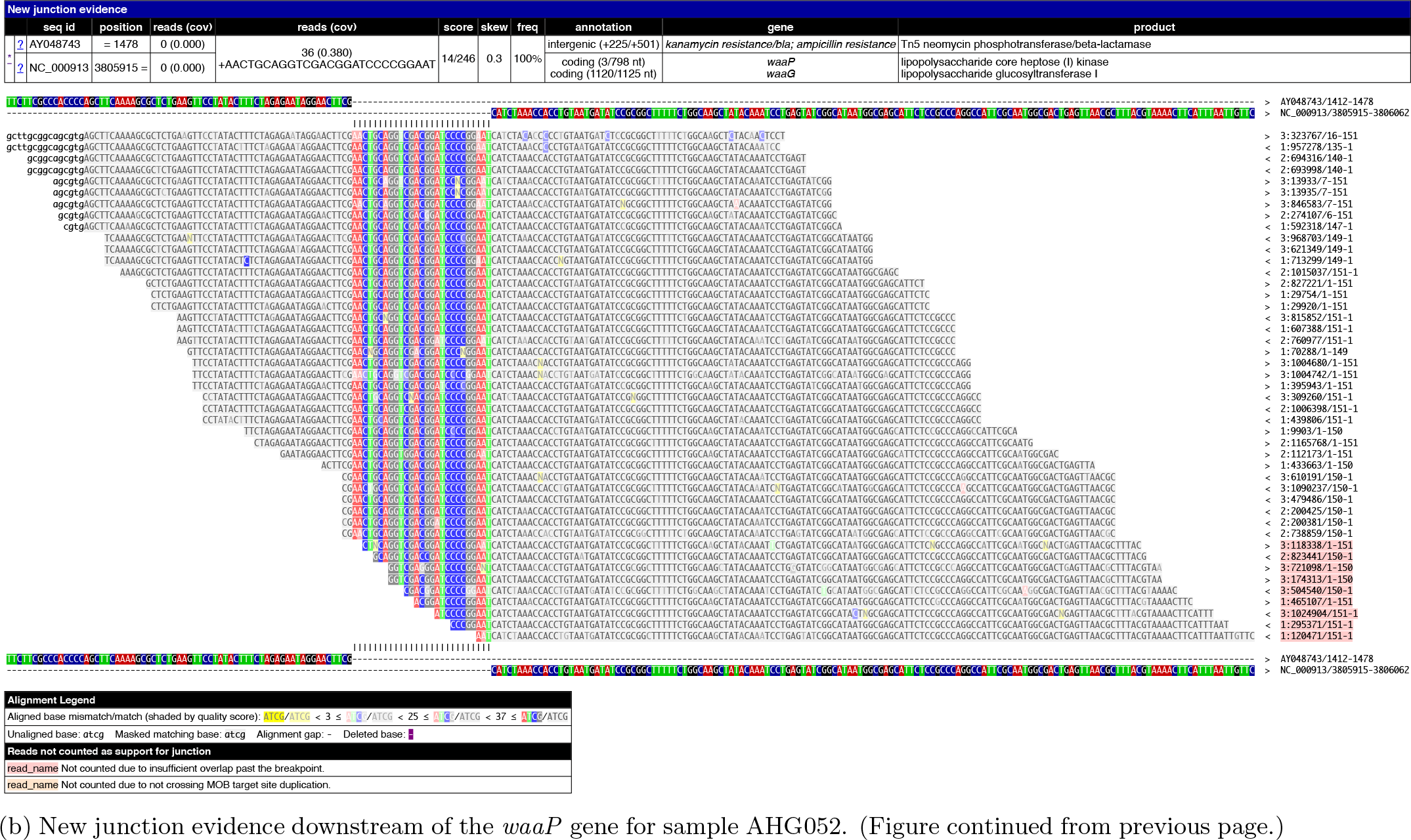
Breseq new junction evidence supporting the insertion of the kanamyacin cassette from the pKD4 plasmid (AY048743.1) at the region of the NC 000913.3 reference sequence around the *waaP* gene for sample AHG052. Panel a), which is on the previous page, shows evidence in support of a new junction at position 3 805 150 of the NC 000913.3 reference sequence (upstream of the *waaP* gene). Panel b) shows evidence of a second junction around position 3 805 915 (downstream of the *waaP* gene). These positions correspond well with the lack of read coverage for this sample at this region shown in figure S8a. Only the new junction evidence from AHG052 is shown as a representative since new junction evidence for other samples and knocked-out genes look similar. All plots are from Breseq.

**Figure S12:**
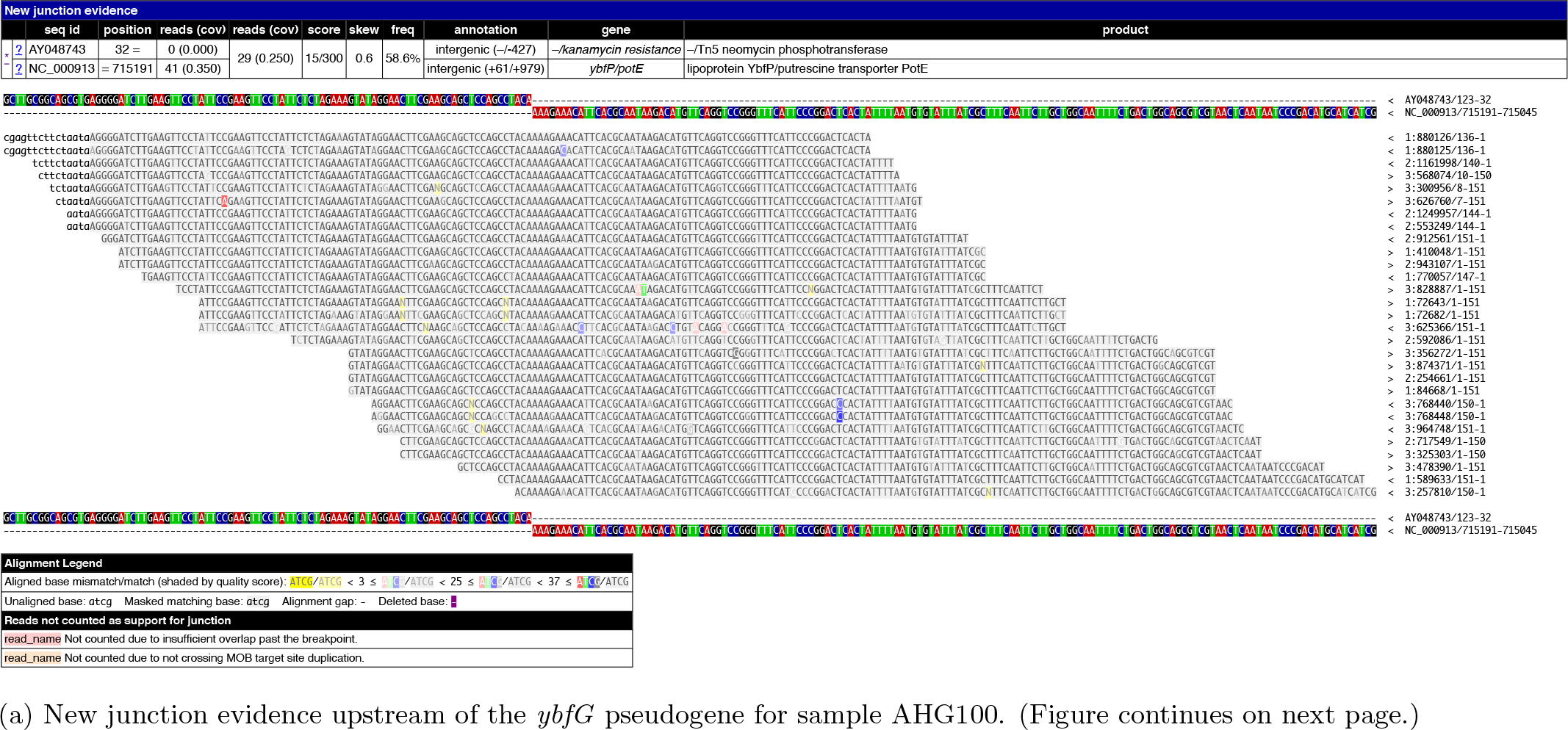
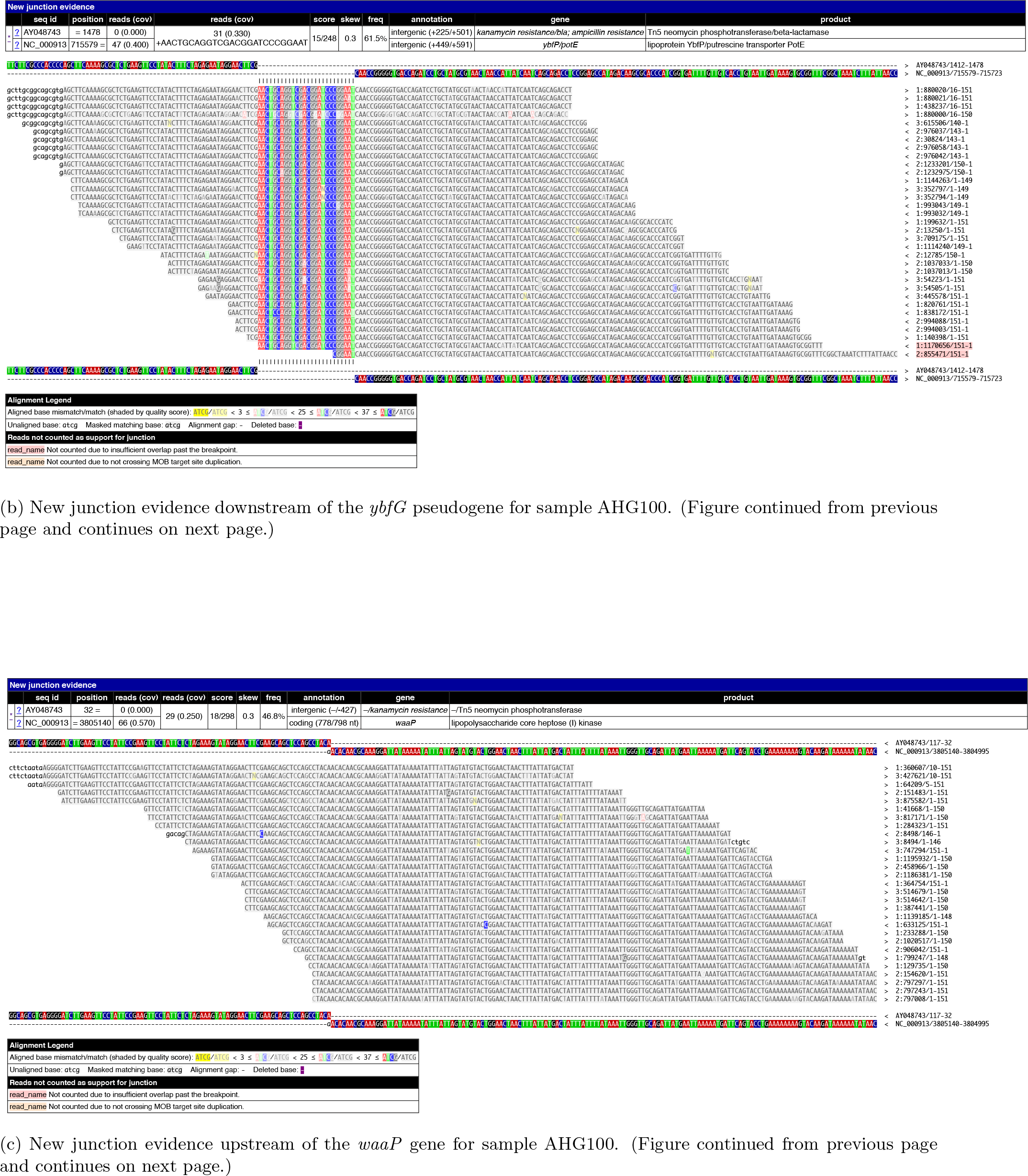
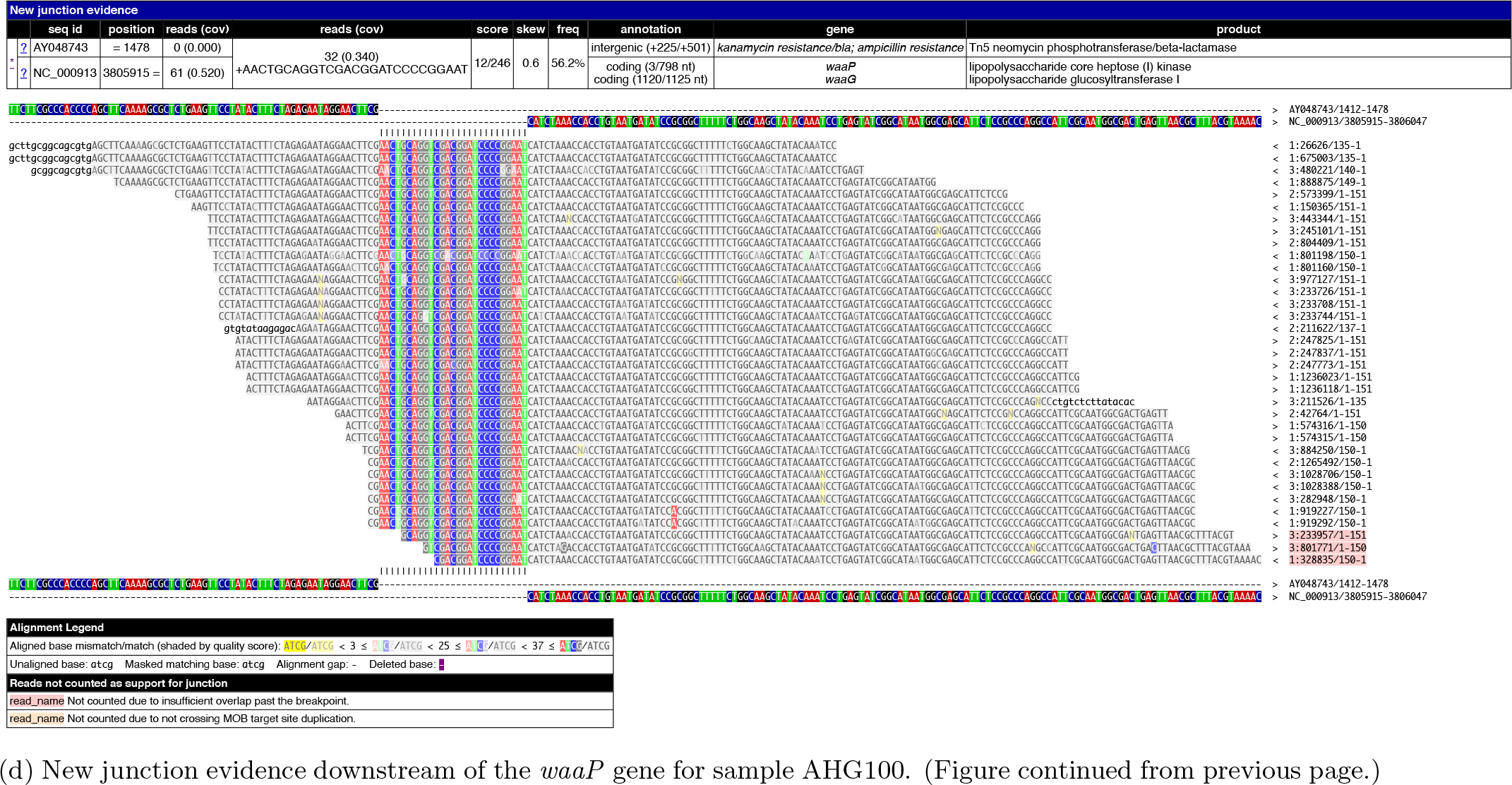
Breseq new junction evidence supporting the hypothesis of likely contamination by sample AHG098 (Δ*waaP*, *rpoB* S512F) into sample AHG100 (Δ*ybfG*, *rpoB* S512F); see also read coverage plots in figures S9d and S10. The new junction evidence for sample AHG100 shows existence of insertion of the kanamyacin cassette from the pKD4 plasmid (AY048743.1) at two different loci of the NC 000913.3 reference sequence. Panels a-b) show the new junctions around the *ybfG* pseudogene and panels c-d) show the new junctions around the *waaP* gene. All plots are from Breseq.

**Figure S13:**
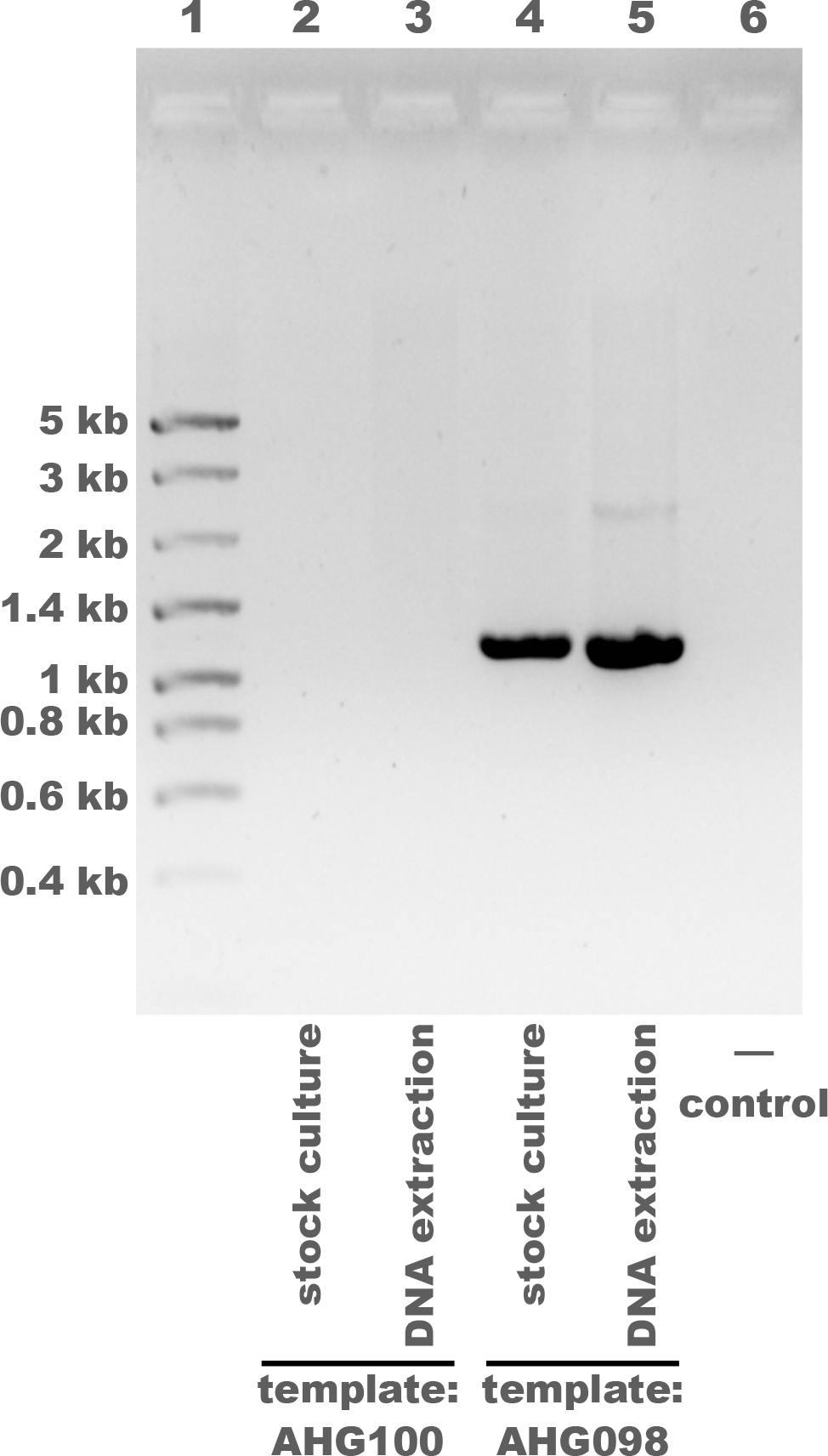
PCR suggests that contamination of sample AHG100 (*rpoB* S512F Δ*ybfG*) occurred after creation of the stock culture and after DNA extraction, perhaps during DNA sequencing. PCR was done using a Δ*waaP* -specific primer (waaP 76747 For in table S13) and the Datsenko-Wanner K1 primer specific to the inserted kanamyacin cassette. Therefore amplification is expected only when the Δ*waaP* knock-out DNA sequence is present in the PCR template. The test sample, AHG100: *rpoB* S512F Δ*ybfG*, was used as a PCR template in lanes 2-3; lanes 4-5 show the positive control with AHG098: *rpoB* S512F Δ*waaP* and lane 6 shows the negative control without any PCR template. No amplification is observed for the test sample AHG100, this suggests that the sample was not contaminated prior to being sent for DNA sequencing. The frozen stock cultures (lanes 2, 4) and the DNA extractions that were sent for sequencing (lanes 3, 5) were used as PCR templates for both samples. For the stock cultures, 1mL of LB was inoculated directly with frozen stock, incubated for 1.5 hours at 37*^◦^C*, then 1*µL* was added to the PCR reaction. A 25*µL* total PCR reaction volume was used with DreamTaq DNA polymerase, annealing temperature of 55*^◦^C*, and 2 minute elongation time. Gel electrophoresis was done in 1% agarose. Lane 1 shows the NZYDNA Ladder VIII, with band sizes labeled at left.

**Table S3:**
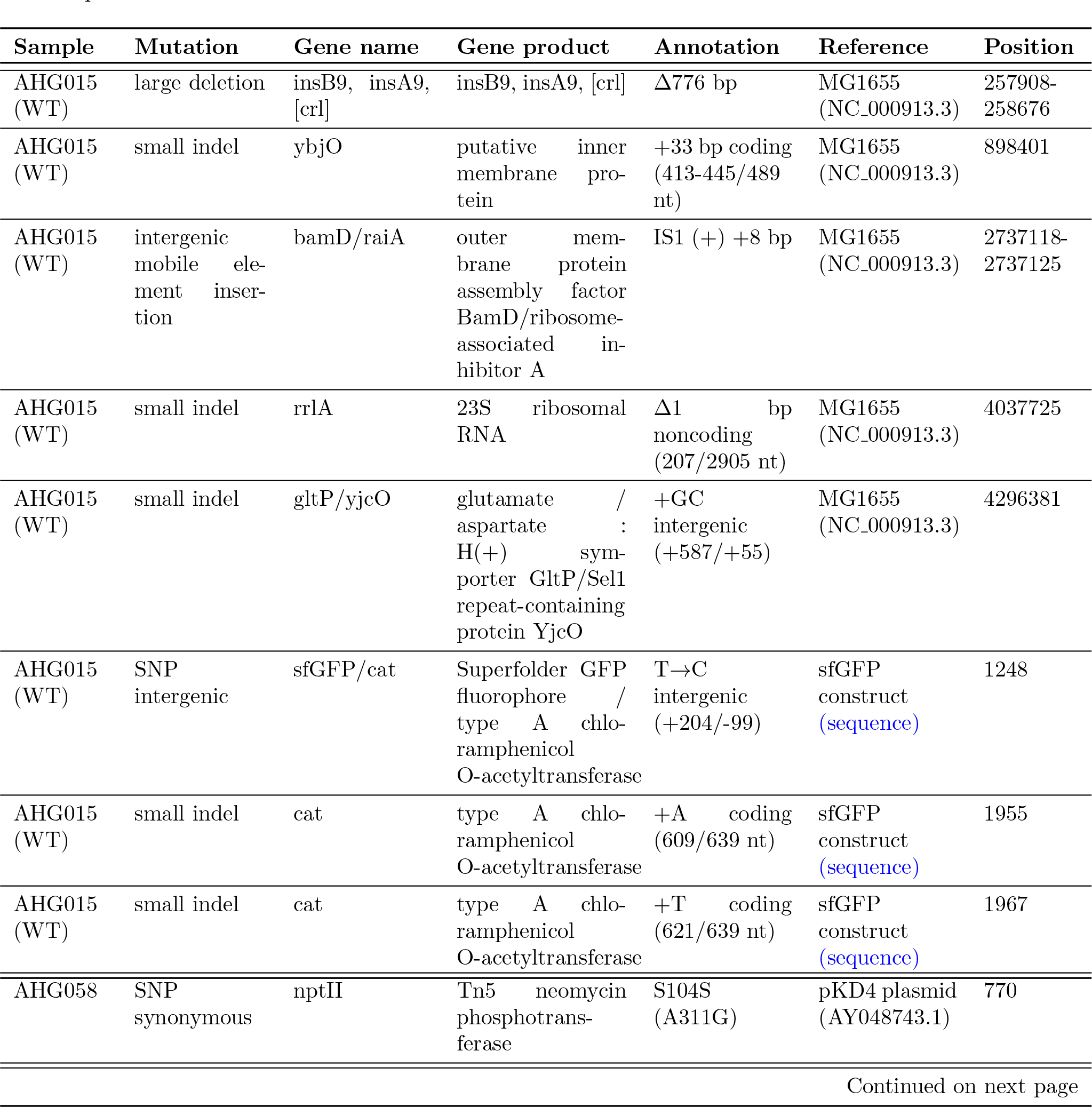
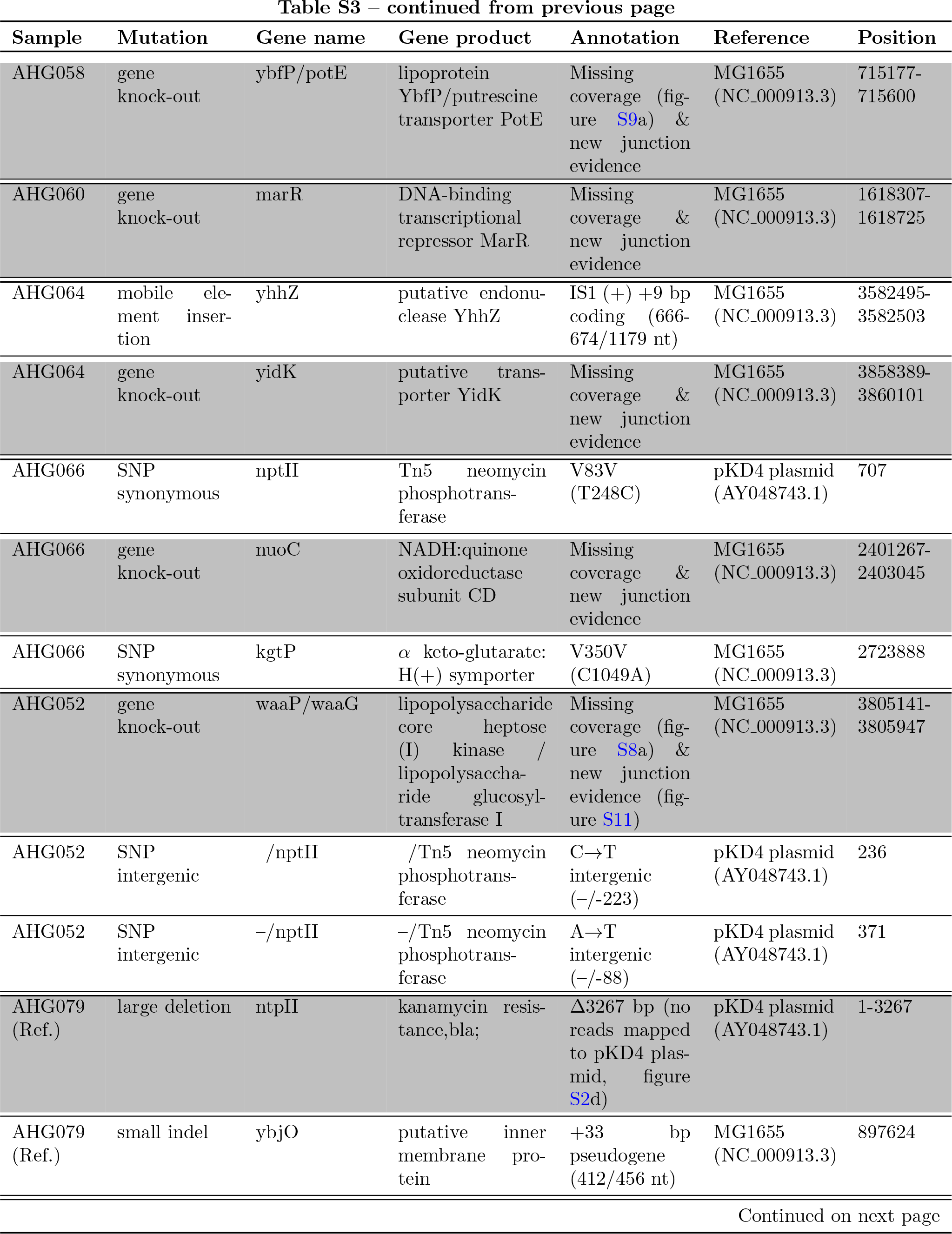
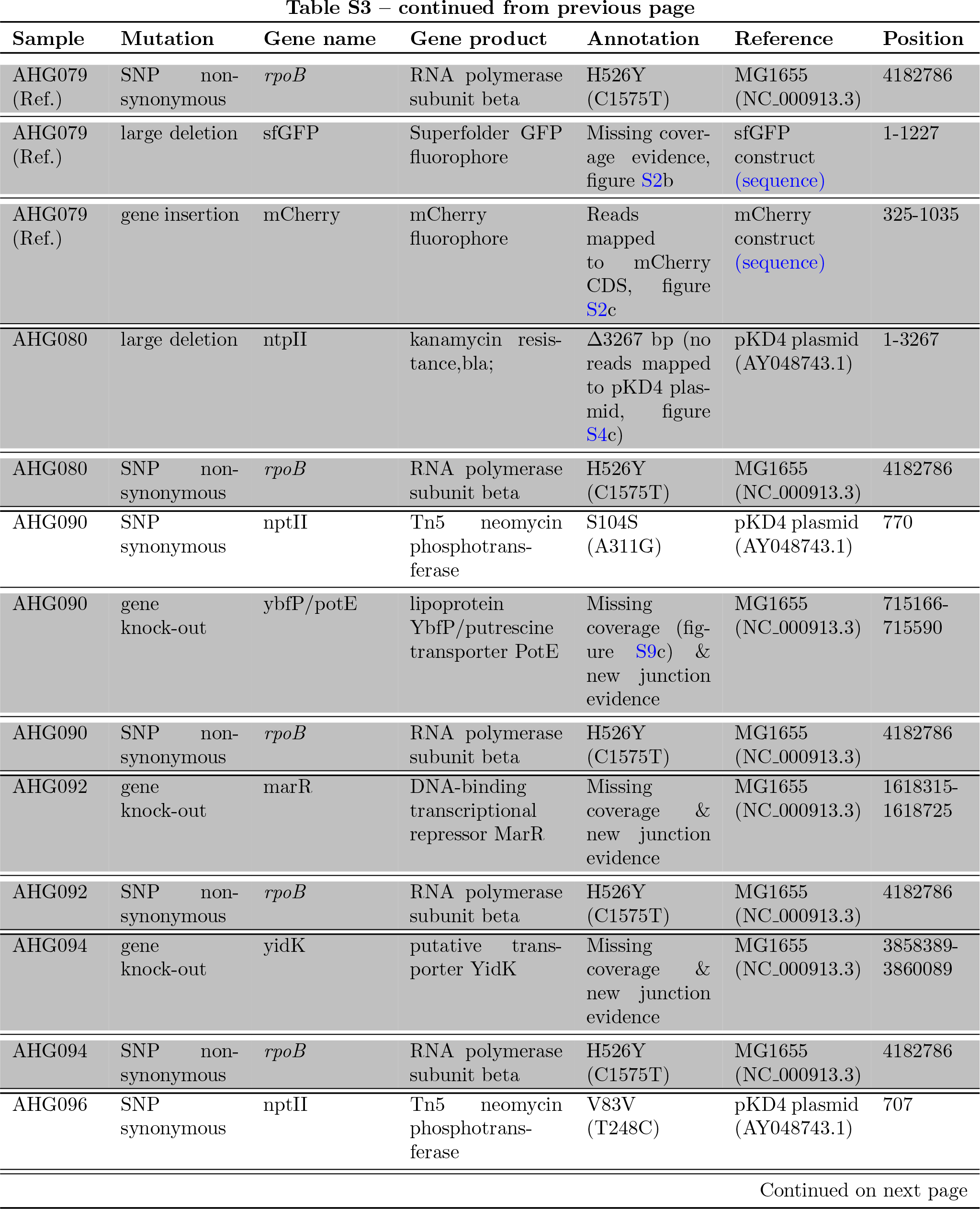
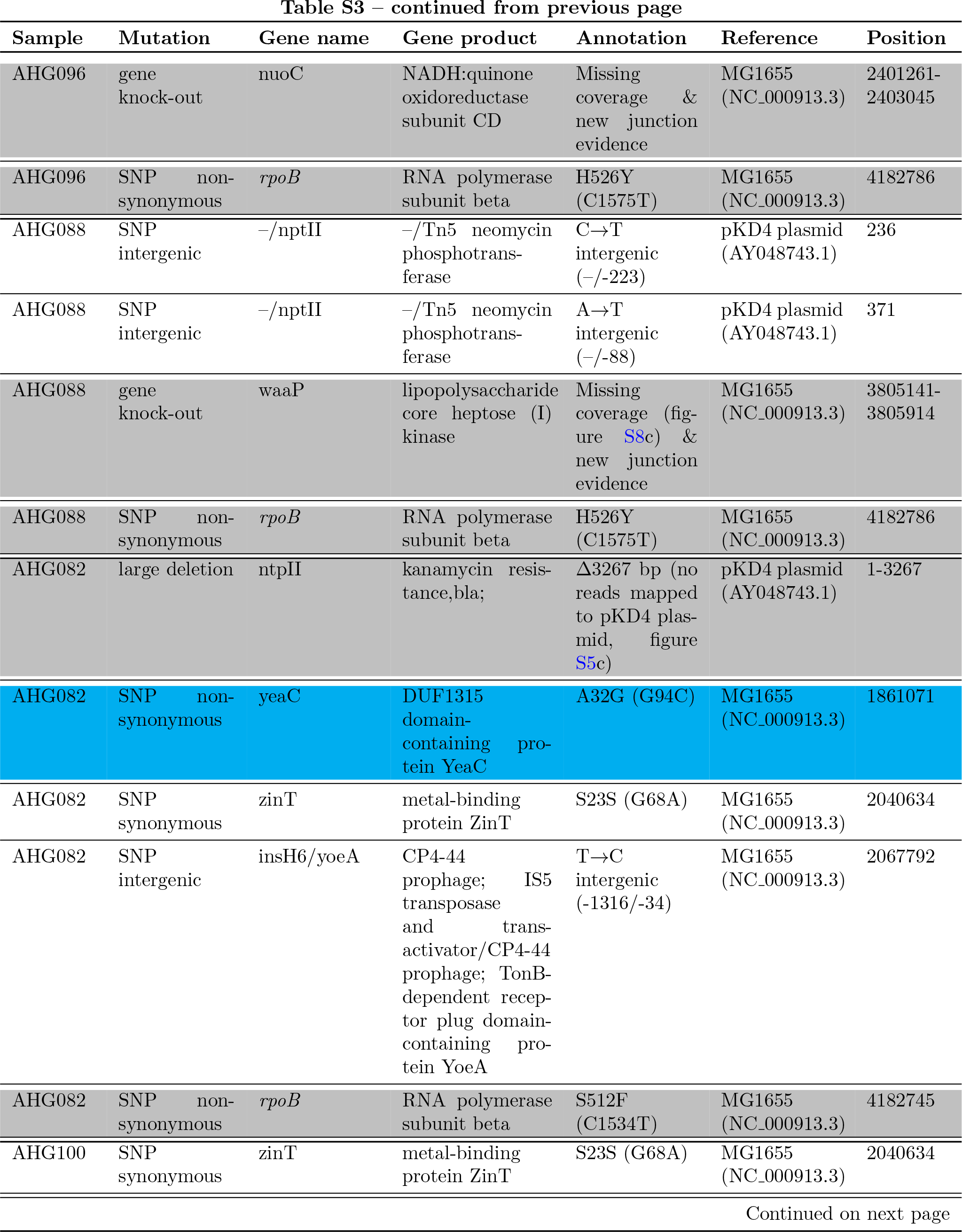
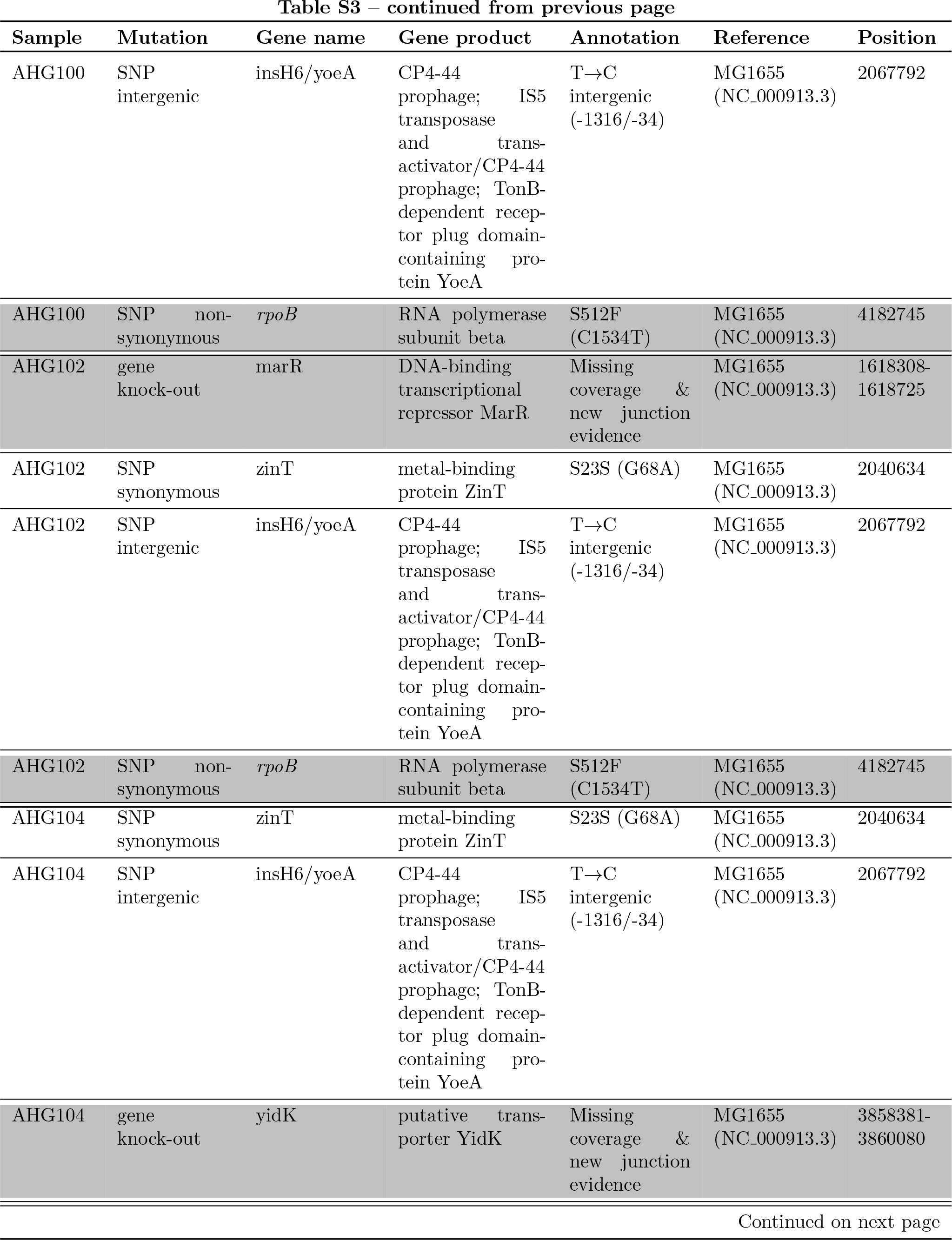
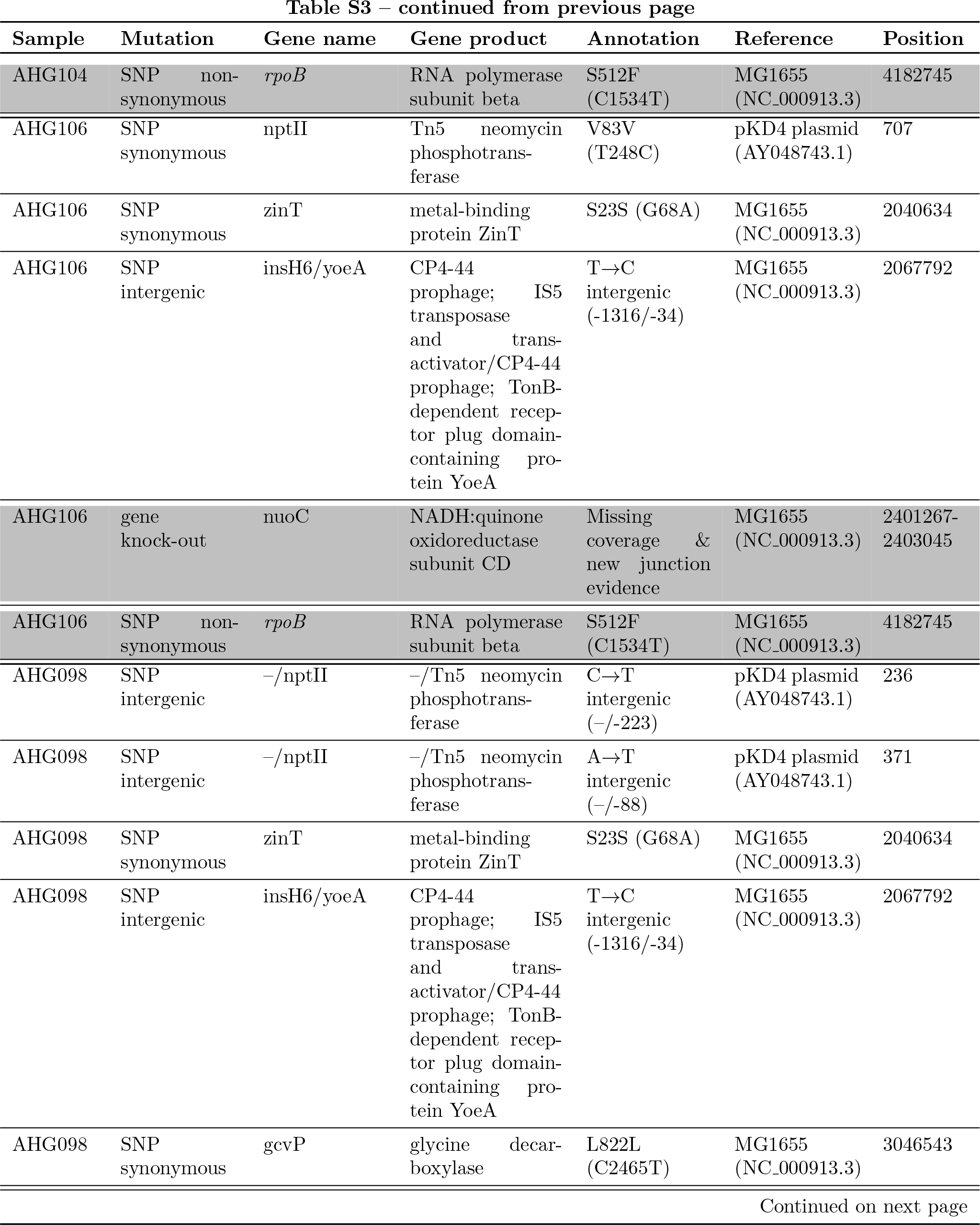
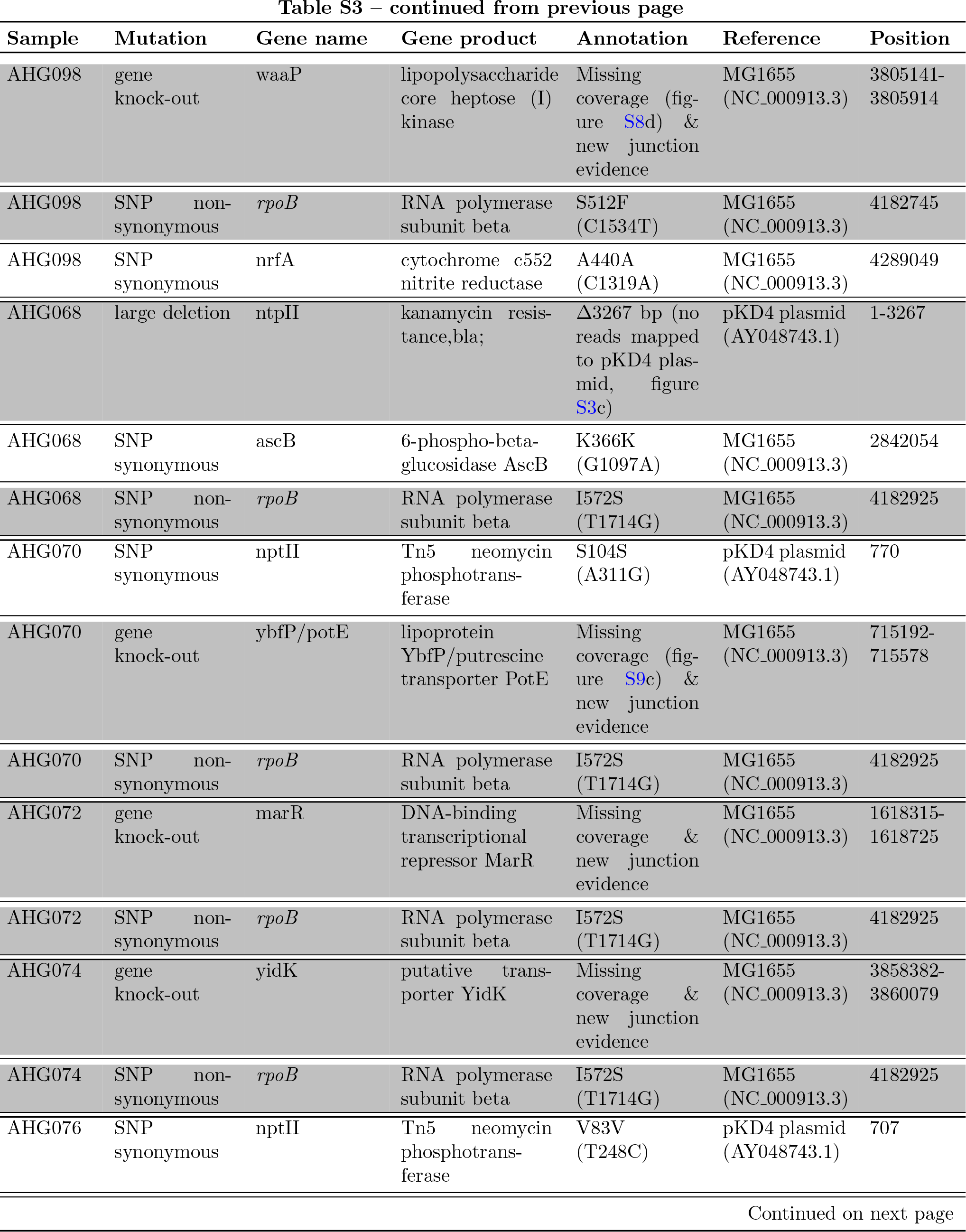
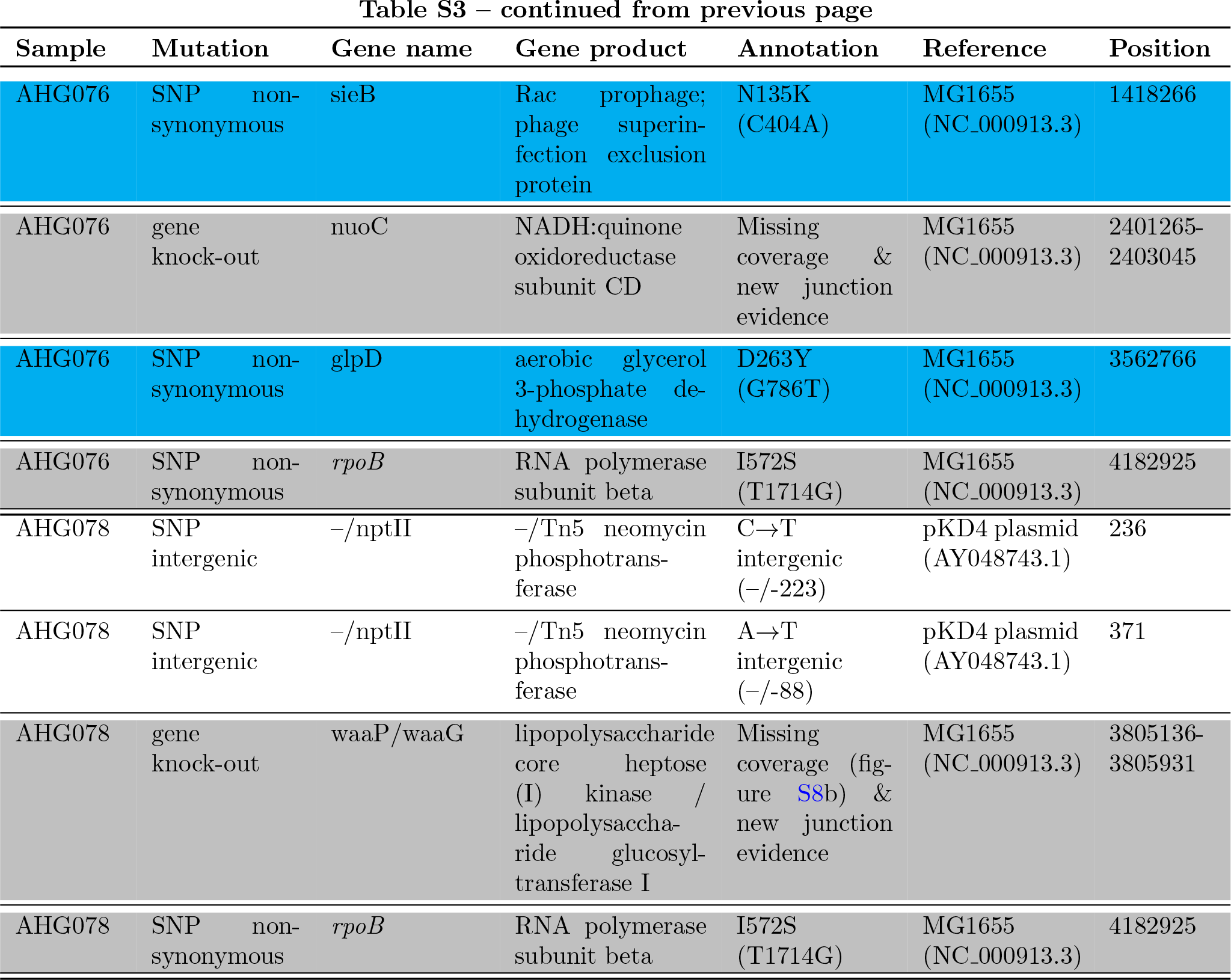
Complete list of mutations as identified by breseq for each of the 25 genotypes. (Engineered mutations listed in table S12.) Rows coloured in gray highlight mutations identified by breseq that correspond with the genetic constructs created for this study. Rows coloured in blue highlight non-synonymous nucleotide substitutions that were not engineered and differ from the wild-type strain (AHG015). All other types of mutations (synonymous nucleotide substitutions, intergenic nucleotide substitutions, small indels, and mobile element insertions) that were not engineered and mutations present on the wild-type strain are listed on white background. Sample AHG079 is labeled with “(Ref.)” to highlight that this was the reference genotype used in competitions.

### 8.2 **Antibiotic dose response curves of the wild-type genotype**

**Figure S14:**
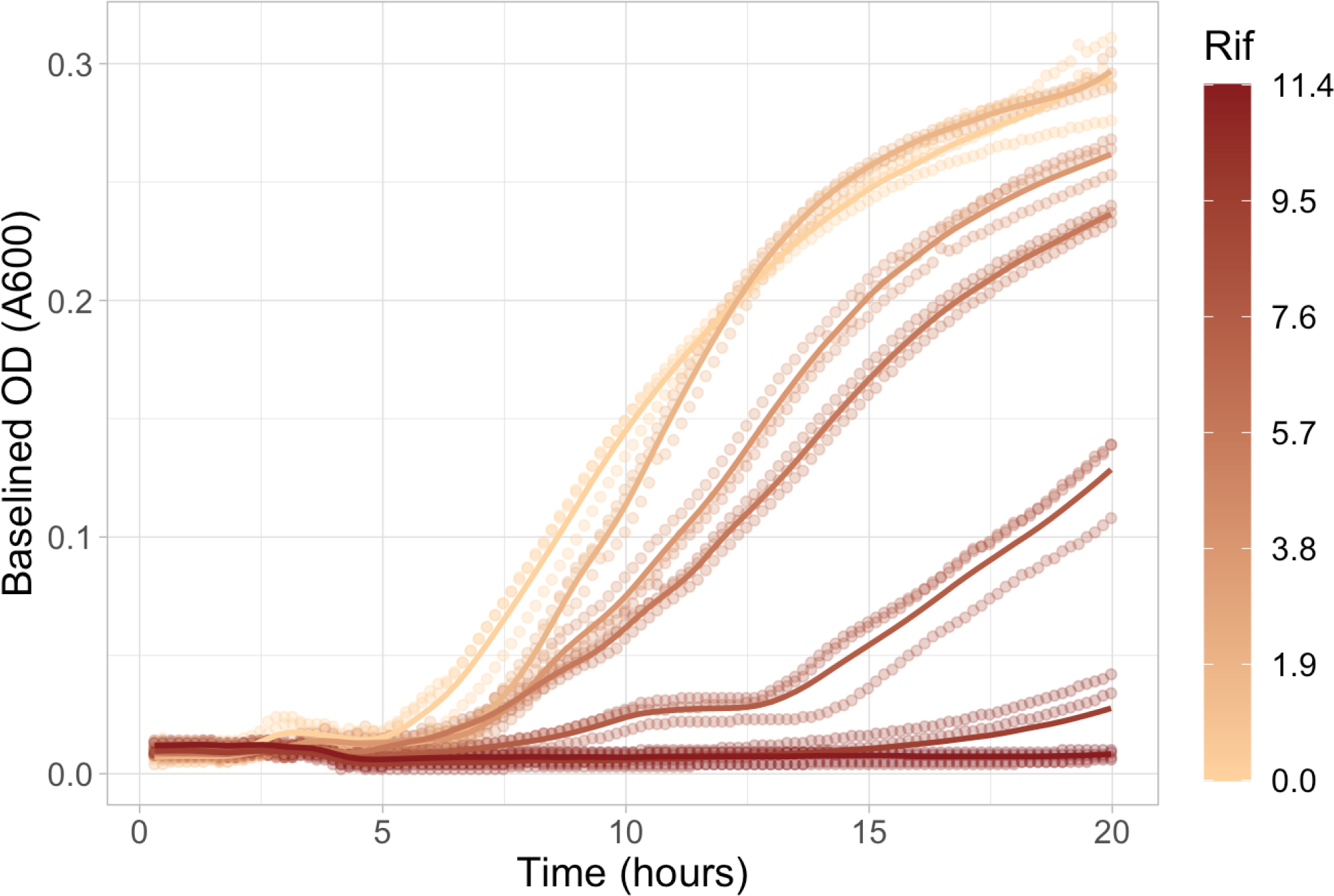
Dose response curves of the antibiotic-susceptible, wild-type genotype (sample AHG015) to different rifampicin concentrations. The y-axis shows the time in hours, the x-axis shows the baseline-subtracted optical density (OD) at 600nm, and the colours indicate different rifampicin concentrations with the legend on the right in units of *µg/mL*. Each line shows the average of three growth curve replicates (loess smoothing was used to interpolate) and points show the individual replicates. Baselining was done by subtracting the smallest observed OD value (including blank wells) at each time point.

### 8.3 Ascertaining flow cytometry estimates of competitive fitness

**Figure S15:**
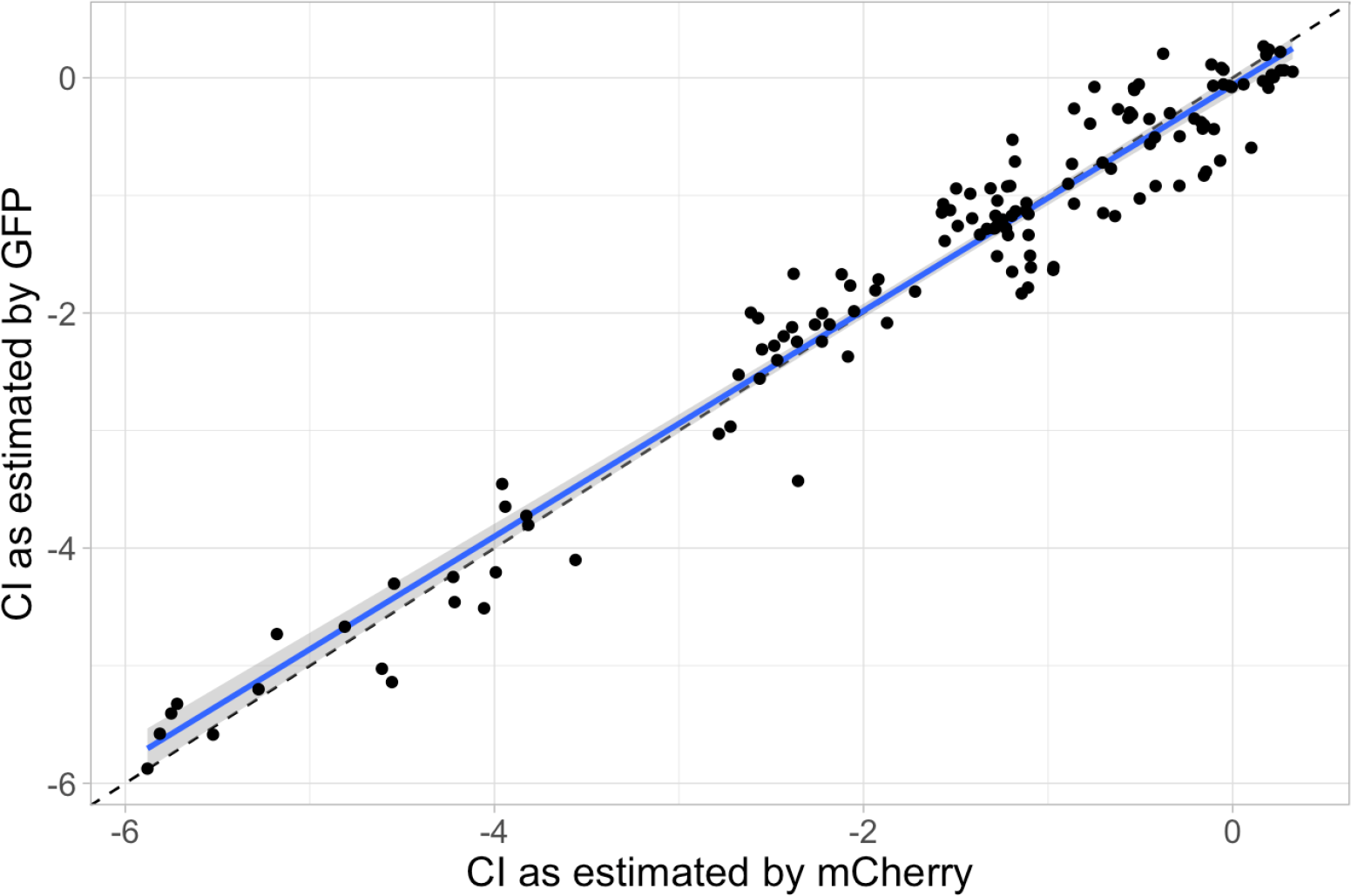
N**e**arly **identical** *w*^ estimates were found when the fluorophores of the reference **and competitor genotypes were swapped.** The competitive index (*w*^) as estimated by using mCherry-labeled competitor genotypes and a GFP-labeled reference genotype (shown in x-axis) corresponds well with the same metric estimated by using GFP-labeled competitor genotypes and an mCherry-labeled reference genotype (shown in y-axis). Data corresponds to a sample of genotypes representing all knock-outs and *rpoB* mutants as assayed once in all environments. The dashed, black line shows *y* = *x*. The blue line shows the mean and the blue shaded region shows the 95% confidence interval of the mean for the least-squares estimated linear regression. The coefficients of this regression are given in table S4; the overall regression is statistically significant and explains most of the variation in the response variable (adjusted *R*^2^ = 0.949, *F* (1, 138) = 2586, *p <* 10*^−^*^15^).

**Table S4:**
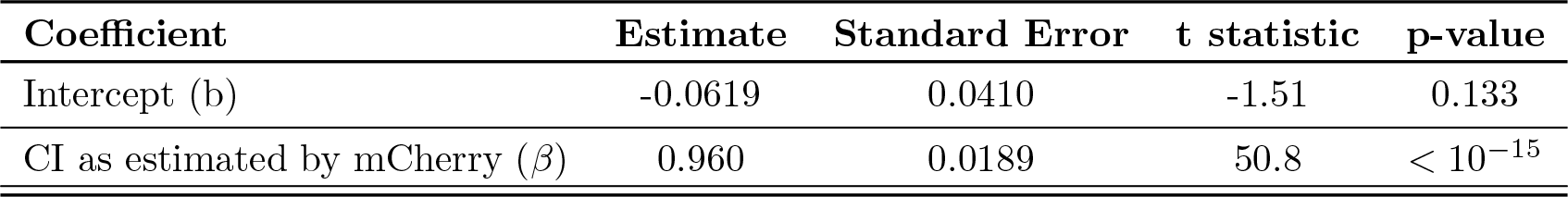
Least-squares estimated coefficients of the linear regression shown in figure S15.

**Figure S16:**
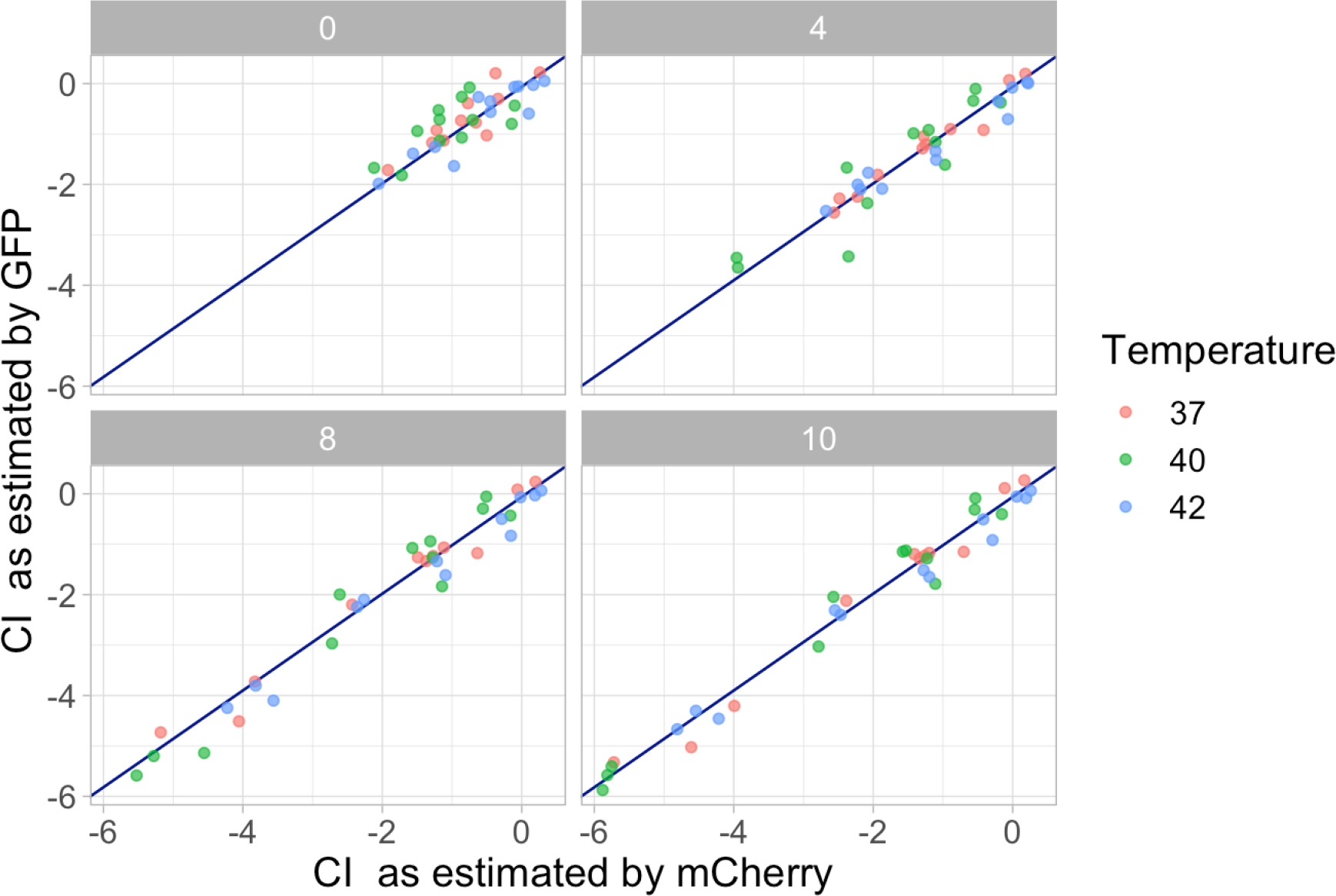
The plot shows environment-specific effects of swapping fluorophores. The same data is plotted as in figure S15 but here panels show the different antibiotic concentrations and points are coloured by temperature. The blue line shows the simple linear regression on CI as estimated by mCherry, and whose coefficient estimates are given in table S4. As shown in table S5, the effect of environment is not significant.

**Table S5:**
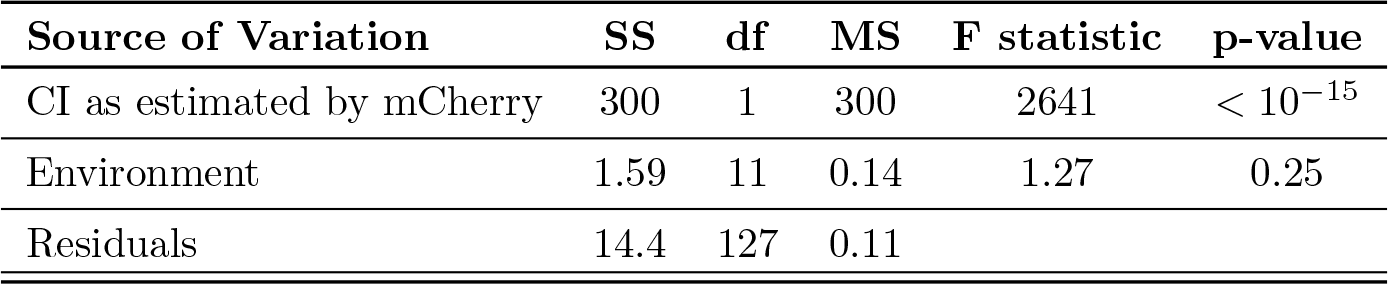
Results of ANOVA show that the environment-specific effect of swapping fluorophores is not statistically significant.

**Figure S17:**
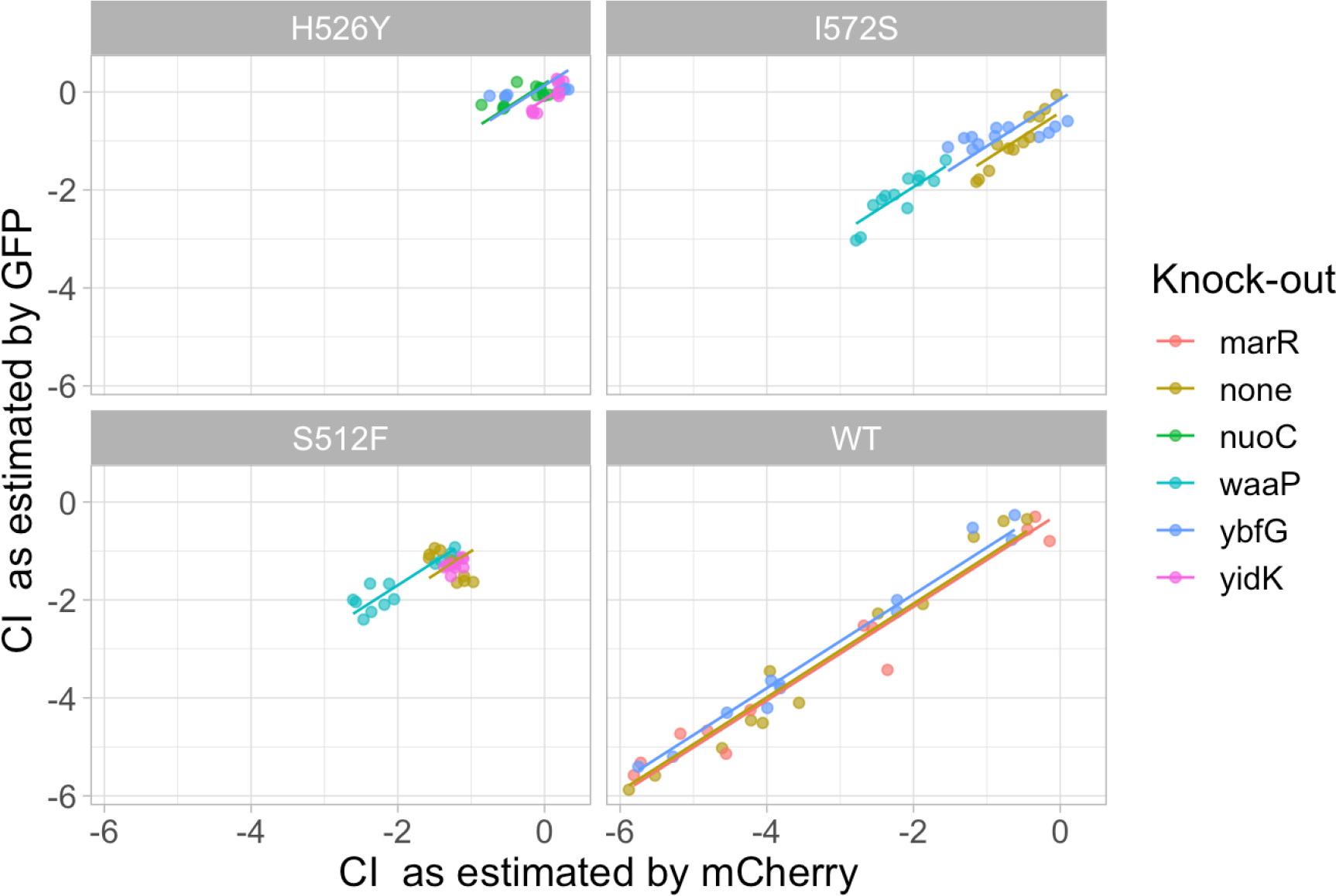
The plot shows genotype-specific effects of swapping fluorophores. The same data is plotted as in figure S15 but here panels show the different *rpoB* substitutions and points are coloured by genotype. Lines show the multiple regression on genotype as detailed in table S6. This effect is statistically significant; however it explains only about 1% more of the variation in the data (adjusted *R*^2^ = 0.9588 for overall regression) as compared to not including this effect.

**Table S6:**
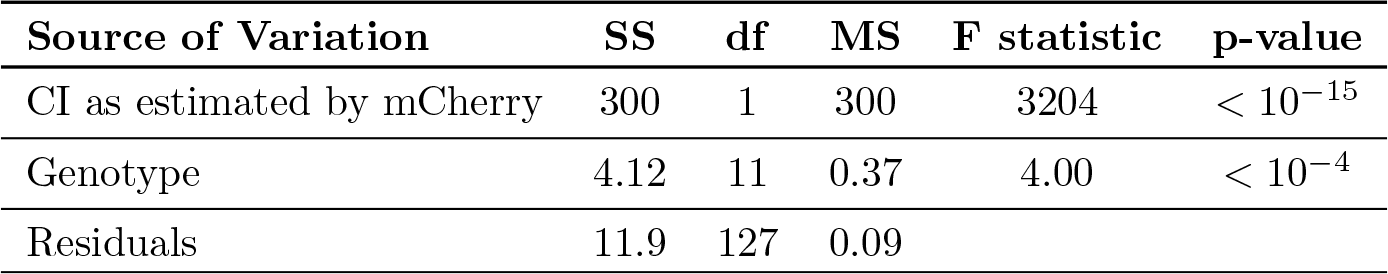
Results of ANOVA show that the genotype-specific effect of swapping fluorophores is statistically significant but small.

### 8.4 Cost of resistance

**Figure S18:**
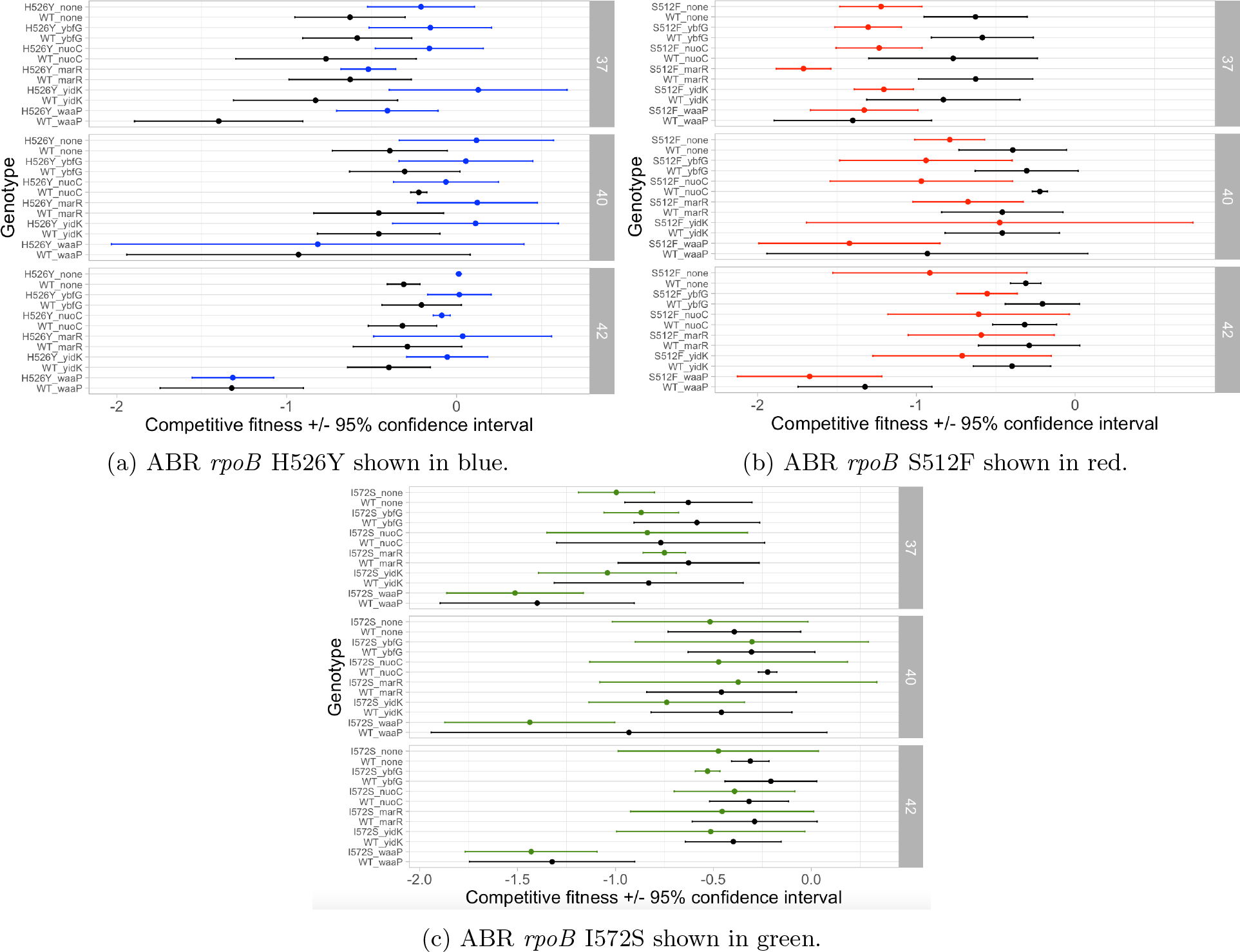
Competitive fitness for ABR mutations in the absence of antibiotic (0*µg/mL* rifampicin) indicates the cost of resistance. Each panel shows a different ABR *rpoB* mutation (colours) as compared to the wild-type *rpoB* (black). The x-axis indicates the competitive index, with points showing mean values and error bars showing the 95% confidence interval on the mean values. The y-axis shows the genotype, with first the *rpoB* mutation then the knock-out (e.g., “H526Y nuoC” indicates the H526Y mutation at *rpoB* on the Δ*nuoc* background); “WT” refers to the wild-type *rpoB* sequence and “none” indicates that there is no knock-out. In each panel, the facets show the three different temperature environments. **(a)** Contrary to previous studies [7, 35], the H526Y mutation has no significant fitness cost – in fact, it exhibits a significantly higher fitness than wild-type at 42*^◦^C* and has a fitness advantage on the Δ*waaP* background at 37*^◦^C*. **(b)** The S512F mutation has no significant fitness cost on most backgrounds except two at 37*^◦^C* (Δ*ybfG* and Δ*marR*, but not on the wild-type *rpoB* background) and one (Δ*nuoC*) at 40*^◦^C*. This is contrary to previous studies that have found S512F to have a higher fitness than wild-type at 40*^◦^C* [7]. **(c)** The I572S mutation has no significant fitness costs, except on the Δ*ybfG* background at 42*^◦^C*.

### 8.5 GxG summary statistics

#### 8.5.1 ‘Classical’ pairwise epistasis

**Figure S19:**
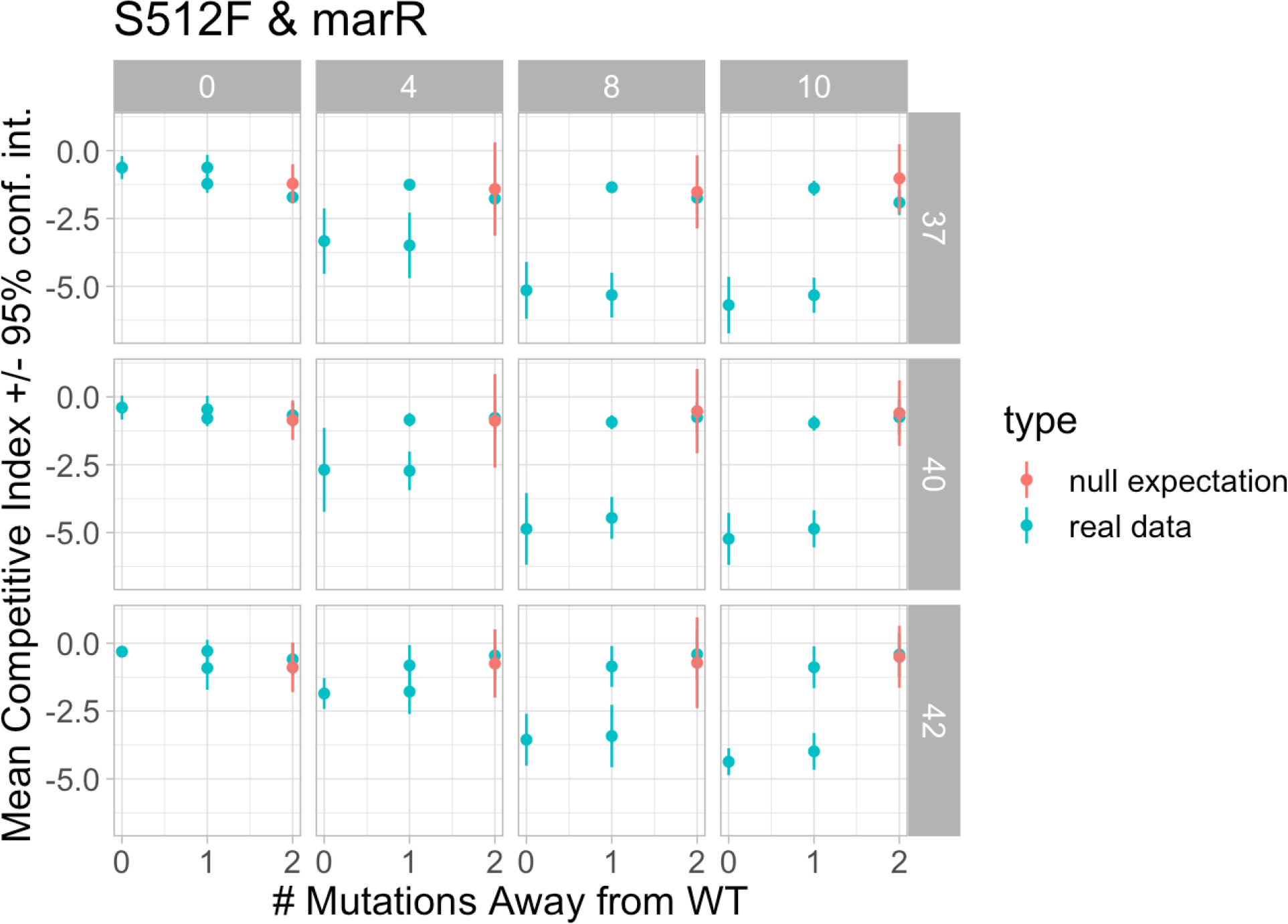
An illustration of how pairwise epistasis was calculated, in this case for the combination of the S512F *rpoB* ABR and the Δ*marR* knock-out mutations. The plot shows the observed mean competitive index (*±* the 95% confidence interval as determined from parametric boostrapping) in blue on the y-axis as a dependent-variable of the number of mutations away from wild-type (x-axis) in each of the 12 environments. Different antibiotic concentrations are shown in each column and different temperatures are shown on each row. The red points show the mean competitive index expected under the null hypothesis of no epistasis. In this example there are no instances where the epistasis is significantly different from zero as shown by the consistent overlap between the 95% confidence interval error bars and the points indicating mean values. Figure S21 summarizes the pairwise epistasis estimates for all genotypes combinations in all environments.

**Figure S20:**
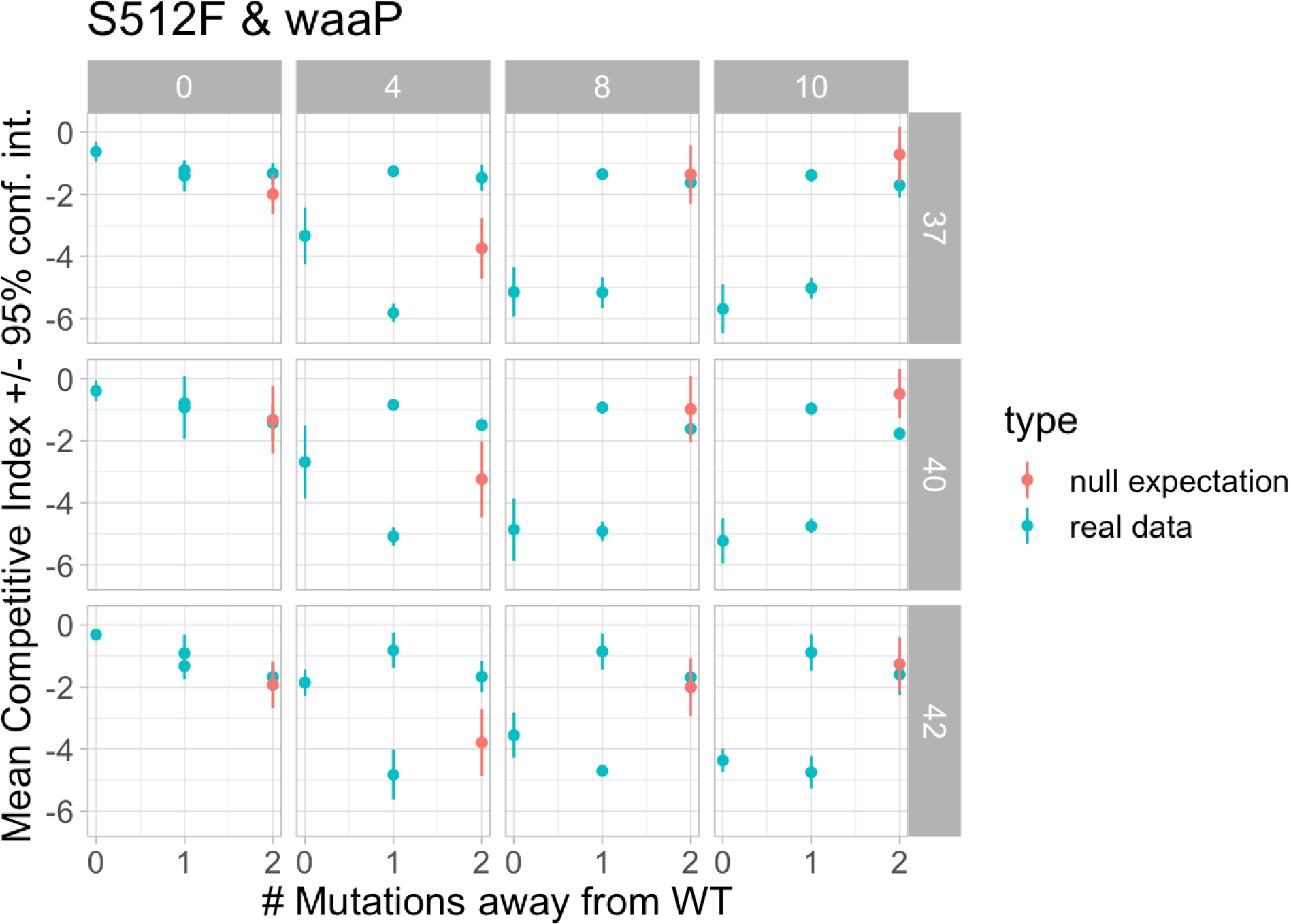
An example of significant pairwise epistasis, in this case for the combination of the S512F *rpoB* ABR and the Δ*waaP* knock-out mutations. The data is plotted in the same style as described in figure S19. Positive epistasis is apparent in all of the temperature environments with 4*µ*g/mL rifampicin, as seen by the larger fitness values of the real data (blue points) as compared to the null expectation of no epistasis (red points). No overlap between the 95% confidence intervals and the mean indicates that the effect is statistically significant. The equivalent plots for Δ*waaP* in combination with the other ABR mutants, H526Y and I572S, show a similar trend of positive epistasis at 4*µ*g/mL rifampicin, as summarized in figure S21. The 40*^◦^C* and 10*µ*g/mL environment displays significant negative pairwise epistasis.

**Figure S21:**
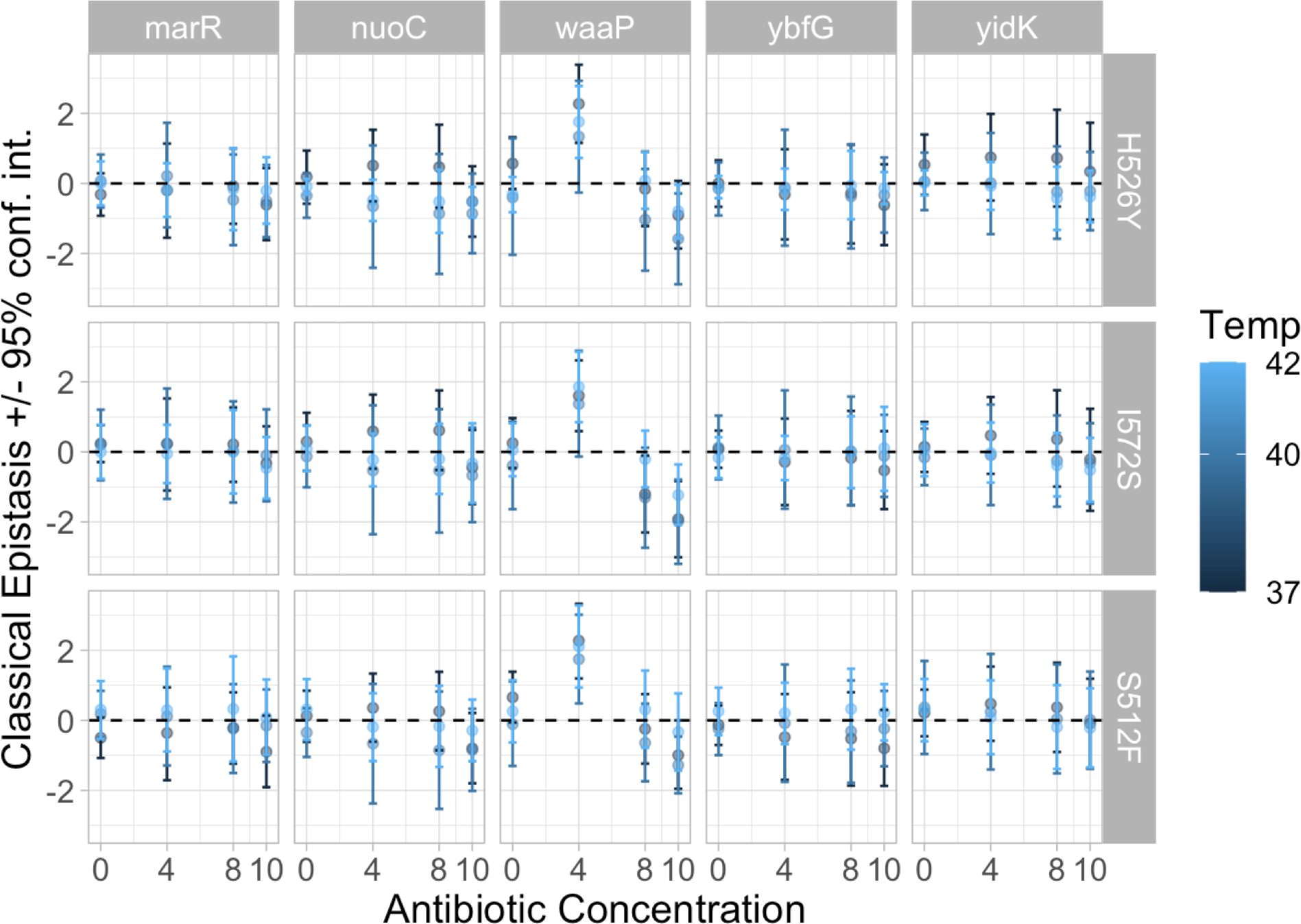
E**s**timated **pairwise epistasis across all genotypes and environments.** The plots are laid out in the same manner as figure 2 except that the y-axis here shows the mean estimated ‘classical’ pairwise epistasis and error bars show the 95% confidence interval as determined from parametric bootstrapping.

#### 8.5.2 Gamma epistasis

**Figure S22:**
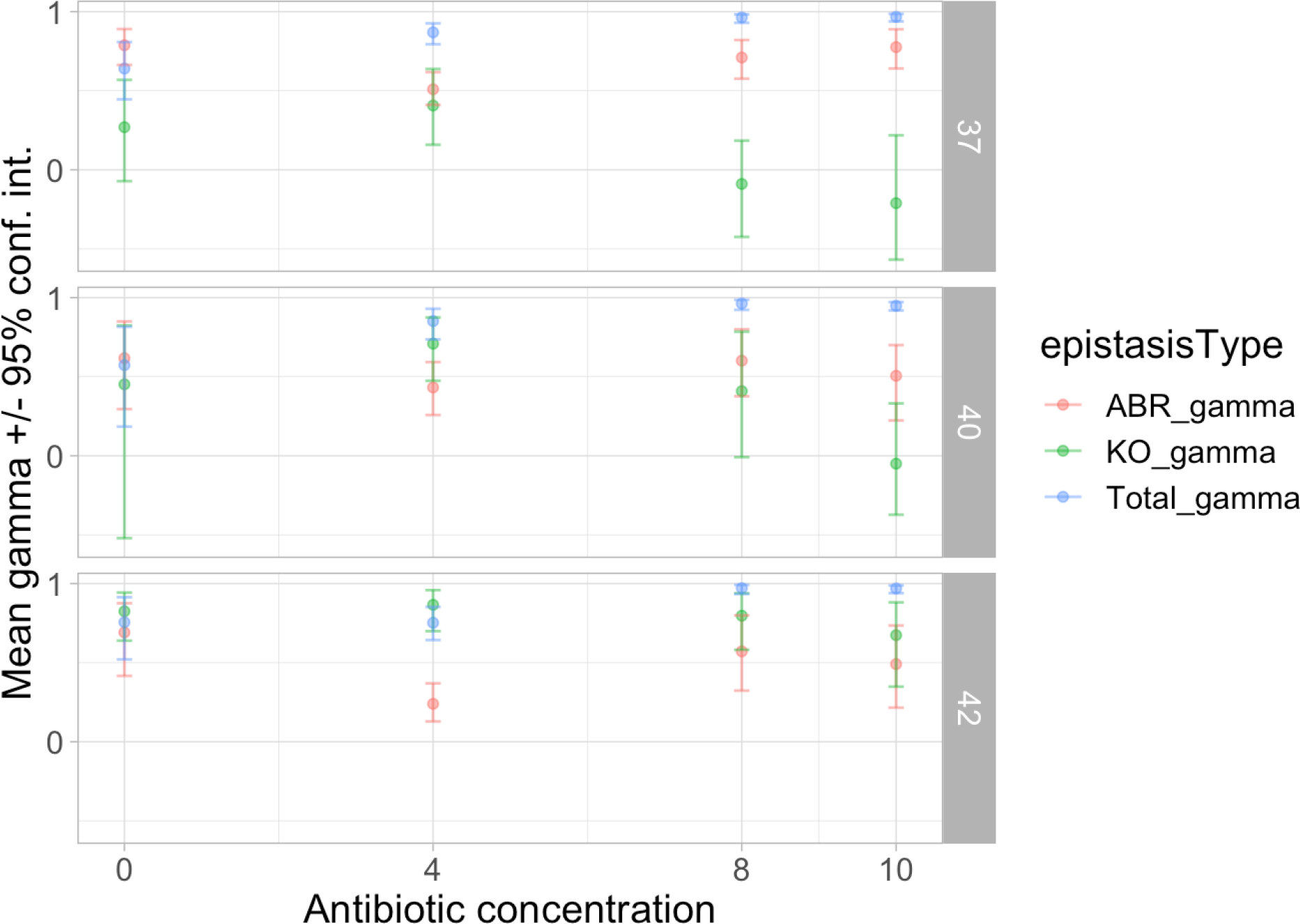
The amount of gamma epistasis is different for different gene classes and temperature environments. Points show the mean gamma epistasis for different gene classes as indicated by the colour (red for ABR mutations at *rpoB*, green for gene knock-out mutations, and blue for the total as already shown in figure 3a. The x-axis shows the rifampicin antibiotic concentration in *µg/mL* and the rows show the three different temperature environments. The error bars show the 95% parametric bootstrap confidence interval. No regression lines are shown as the correlations are not significant at *α* = 0.01.

#### 8.5.3 Fraction of sign epistasis

**Table S7:**
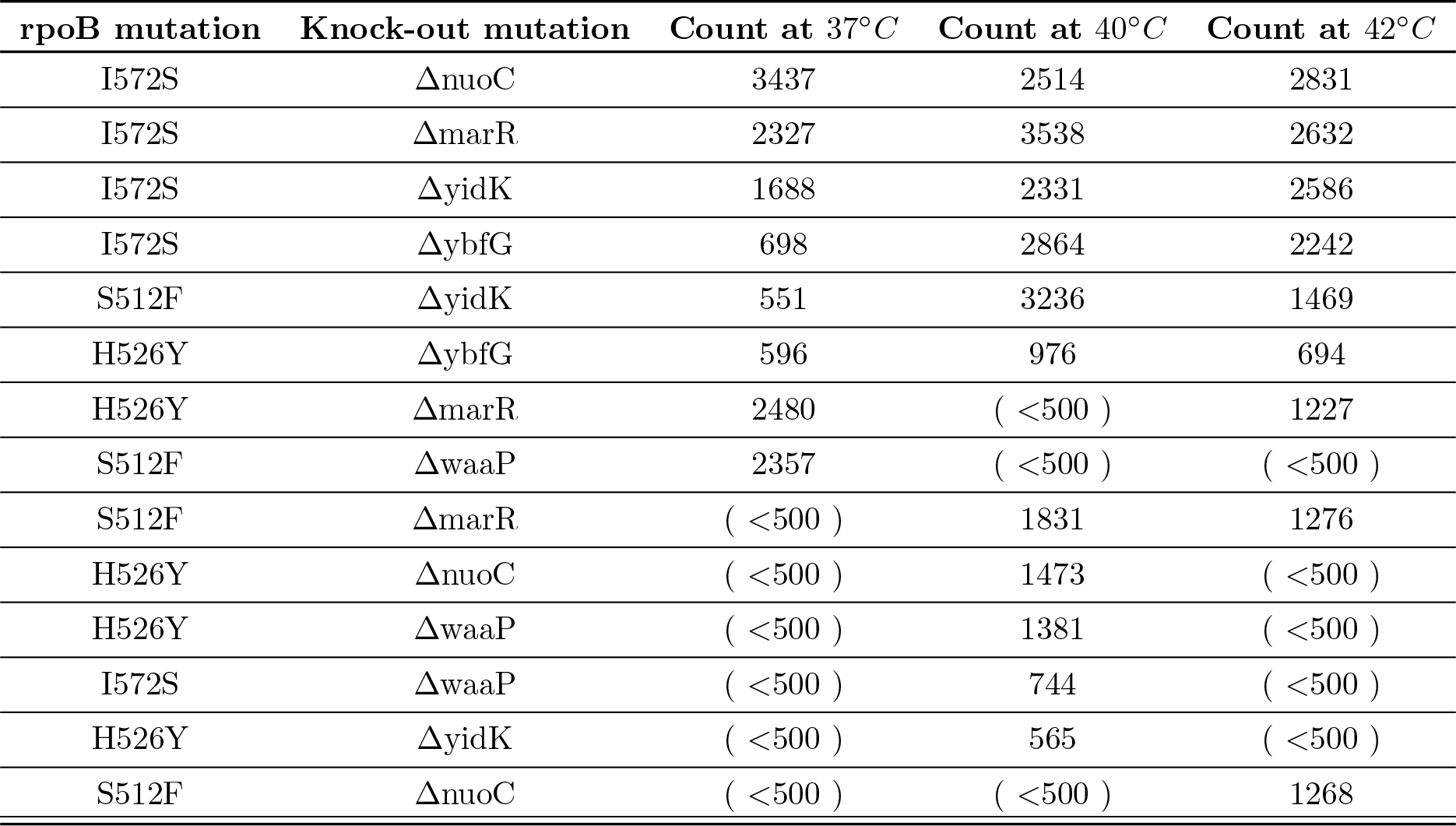
List of the genotypes that exhibited reciprocal sign epistasis in more than 5% of the 10 000 parametric bootstrap samples. The counts of observed reciprocal sign epistasis for all bootstrap samples is given for each of the three temperature environments. Only the environment without anitbiotic is shown because the other environments did not exhibit reciprocal sign epistasis in more than 5% of bootstrap samples.

| **rpoB mutation** | **Knock-out mutation** | **Count at 37°C** | **Count at 40°C** | **Count at 42°C** |
| --- | --- | --- | --- | --- |
| I572S | ΔnuoC | 3437 | 2514 | 2831 |
| I572S | ΔmarR | 2327 | 3538 | 2632 |
| I572S | ΔyidK | 1688 | 2331 | 2586 |
| I572S | ΔybfG | 698 | 2864 | 2242 |
| S512F | ΔyidK | 551 | 3236 | 1469 |
| H526Y | ΔybfG | 596 | 976 | 694 |
| H526Y | ΔmarR | 2480 | ( <500 ) | 1227 |
| S512F | ΔwaaP | 2357 | ( <500 ) | ( <500 ) |
| S512F | ΔmarR | ( <500 ) | 1831 | 1276 |
| H526Y | ΔnuoC | ( <500 ) | 1473 | ( <500 ) |
| H526Y | ΔwaaP | ( <500 ) | 1381 | ( <500 ) |
| I572S | ΔwaaP | ( <500 ) | 744 | ( <500 ) |
| H526Y | ΔyidK | ( <500 ) | 565 | ( <500 ) |
| S512F | ΔnuoC | ( <500 ) | ( <500 ) | 1268 |

**Figure S23:**
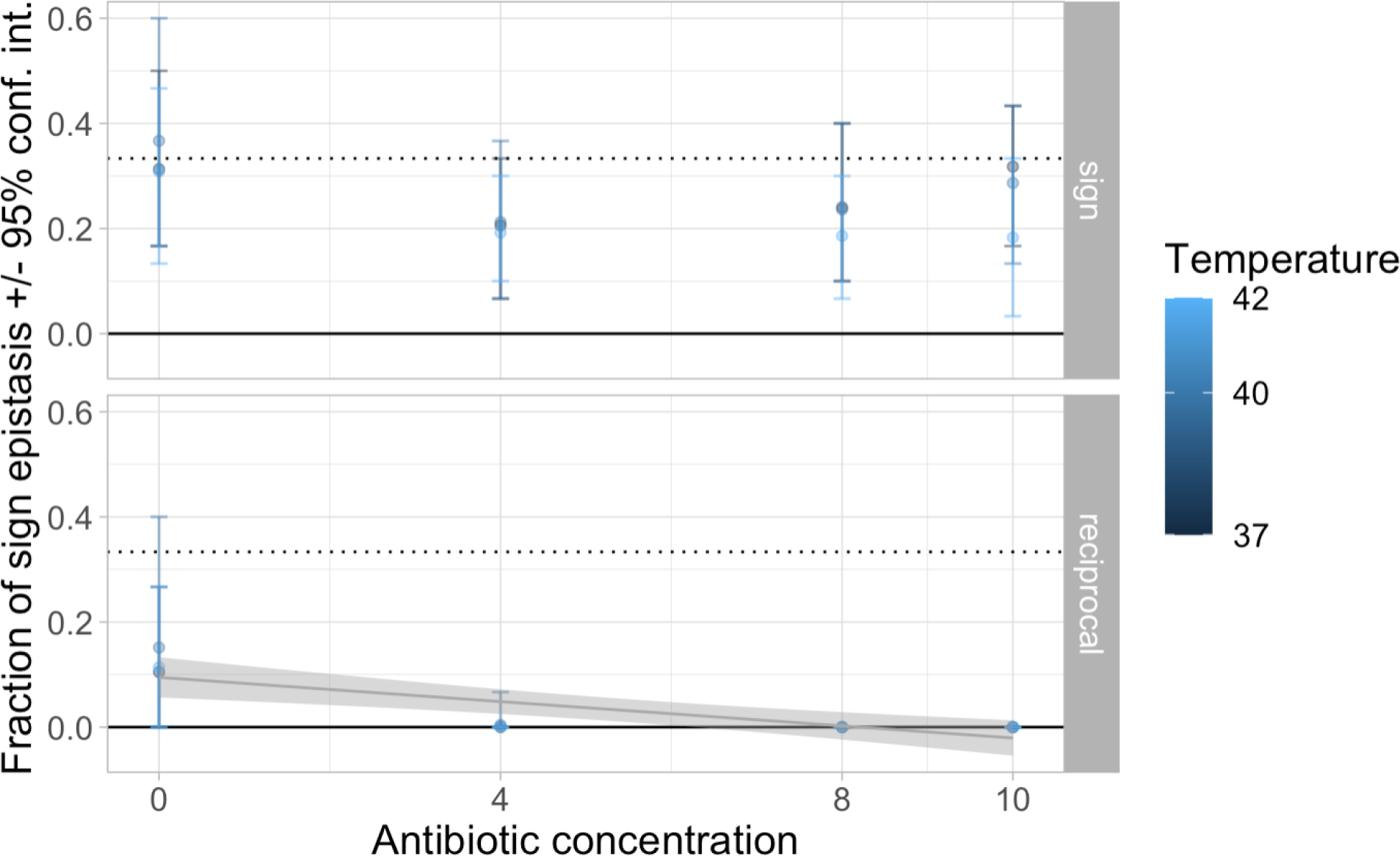
The fraction of simple and reciprocal sign epistasis for different environments. Reciprocal sign epistasis (bottom facet) is mostly observed in the absence of antibiotic while the fraction of simple sign epistasis (top facet) shows no apparent trend with different environments. The points show the mean fraction and the error bars show the 95% parametric bootstrap confidence interval. The x-axis shows the antibiotic concentration and the colour shows the temperature. The solid line at *y* = 0 highlights the fraction of simple and reciprocal sign epistasis expected in a purely additive landscape without any epistasis. The dotted line at *y* = ^1^ highlights the fraction of simple and reciprocal sign epistasis expected in a completely epistatic house-of-cards landscape. There is no significant correlation between the antibiotic concentration and the fraction of simple sign epistasis (*F* (1, 10) = 2.15, *p* = 0.17). The grey trend-line shows the significant negative correlation of antibiotic concentration on the fraction of reciprocal sign epistasis (*F* (1, 10) = 20.3, *p <* 0.005, adjusted *R*^2^ = 0.637) and the grey shaded region shows the 95% confidence interval of the mean for the least-squares estimated linear regression. Antibiotic concentration explains more of the variation in the fraction of reciprocal sign epistasis than environmental quality (figure S40).

#### 8.5.4 Roughness-to-slope ratio

**Figure S24:**
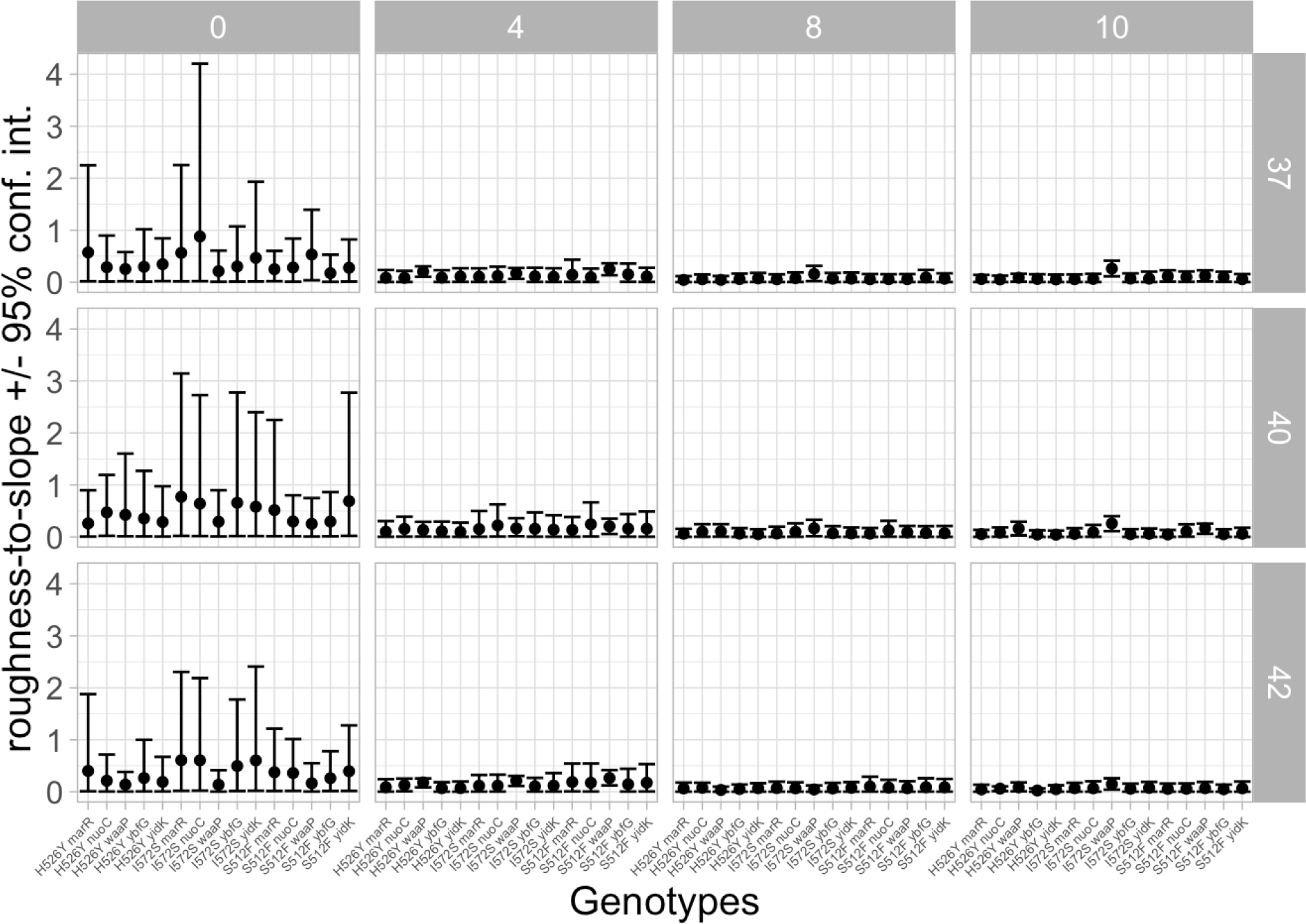
The roughness-to-slope ratio (y-axis) for each genotype (x-axis) in each environment. The dots show the mean roughness-to-slope ratios and the error bars show the 95% parametric bootstrap confidence interval. The columns show different antibiotic concentrations and the rows show different temperature environments.

#### 8.5.5 Diminishing-returns epistasis

**Figure S25:**
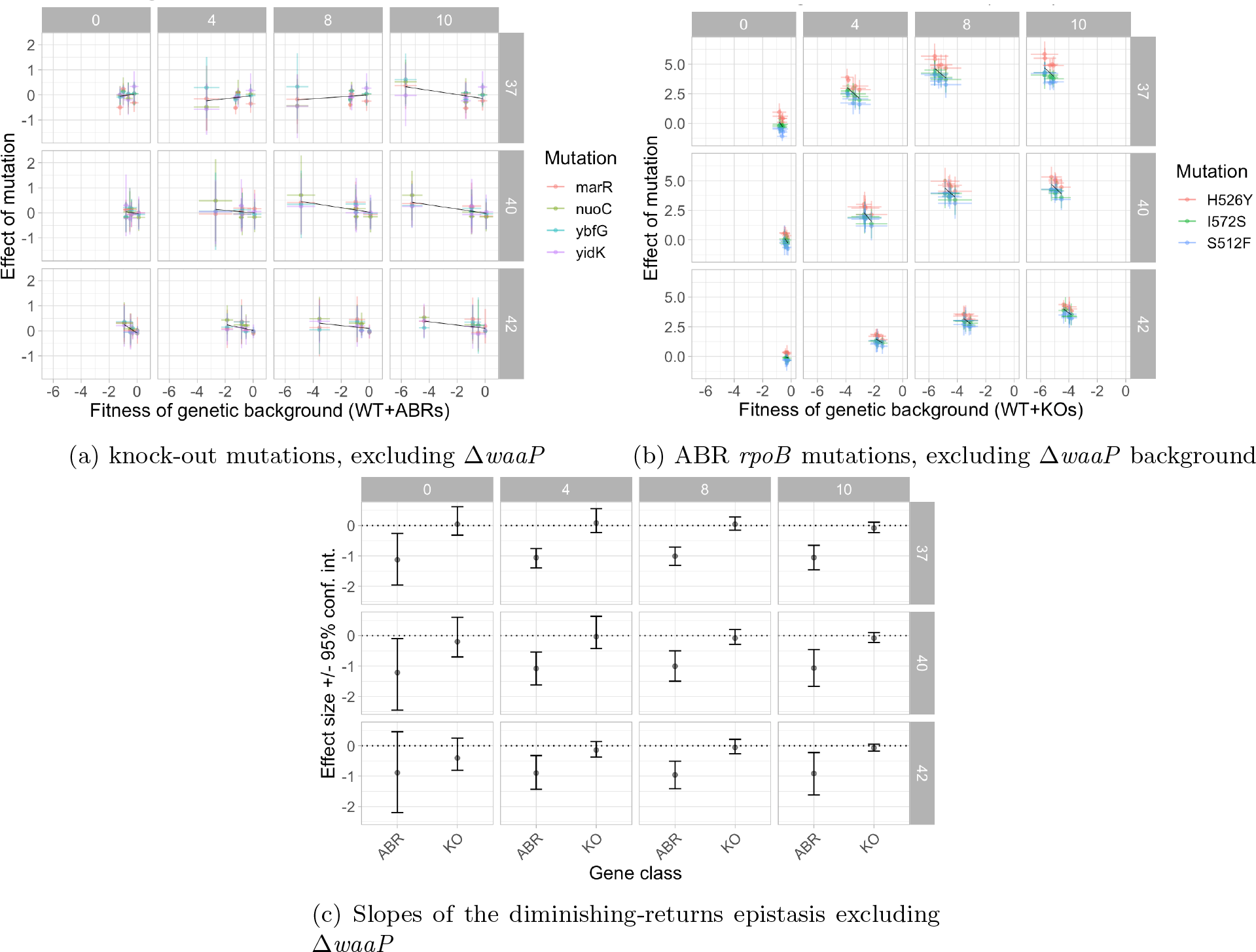
When the Δ*waaP* mutation and background are removed from the analysis, knock- outs continue to display no diminishing-returns epistasis and ABR *rpoB* mutations exhibit diminishing-returns epistasis in all but one of the environments. The plots are in the same style as figure 4 except that the effect of the Δ*waaP* mutation and mutations on the Δ*waaP* background have been removed to confirm that Δ*waaP* is not responsible for the overall trend.

### 8.6 Multiple regression for competitive fitness

#### 8.6.1 Additive (‘null’) model

**Table S8:**
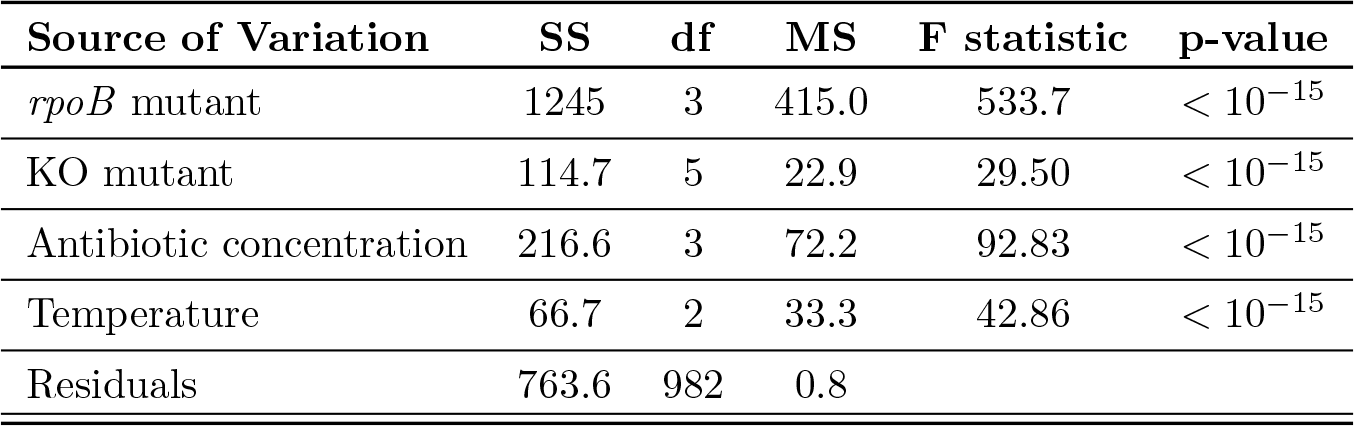
ANOVA shows that the additive effects of rpoB mutants, KO mutants, antibiotic concentration, and temperature are all significant.

**Figure S26:**
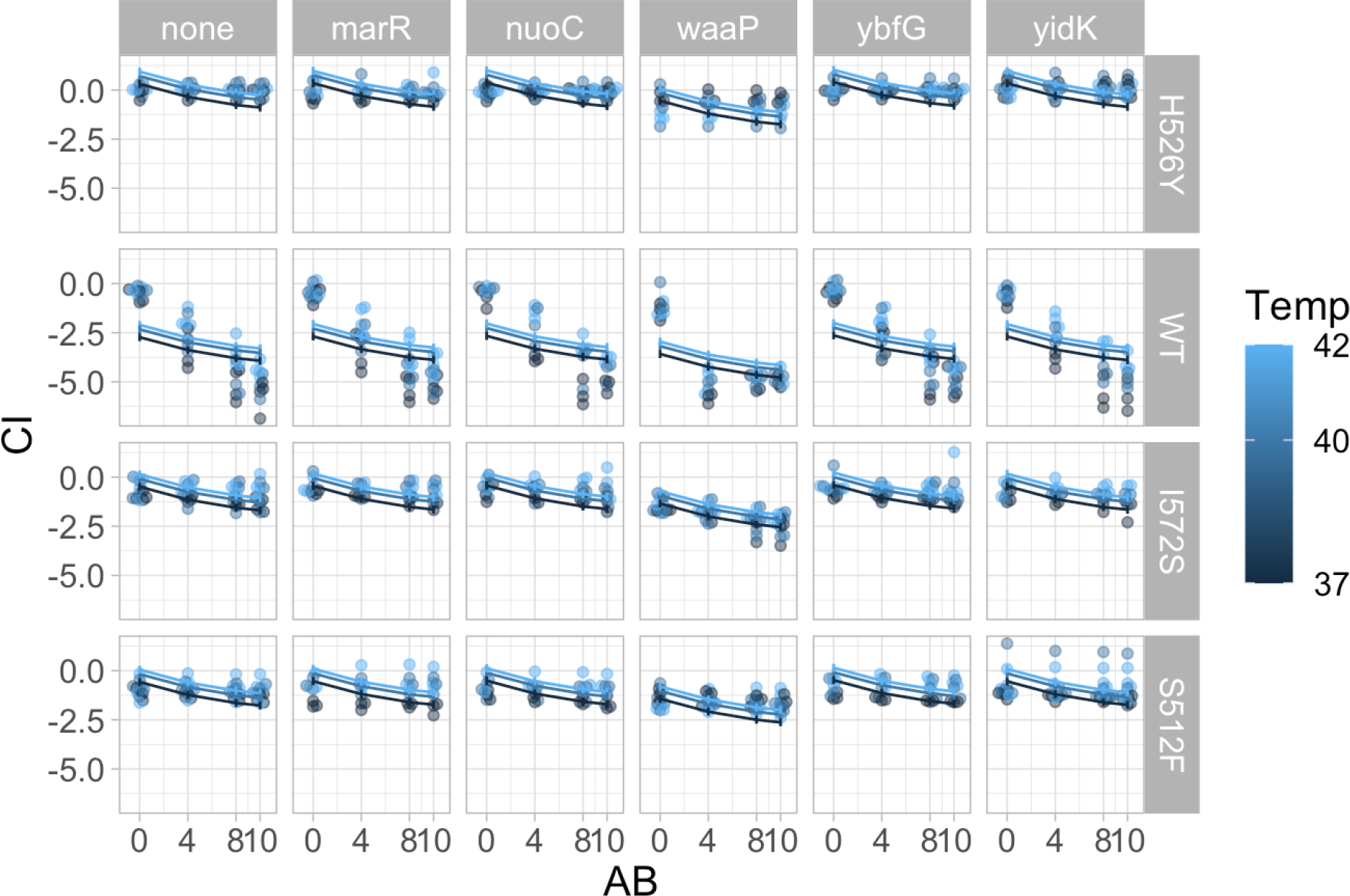
A linear regression with only additive effects explains almost 70% of the variation observed in the competition data. The x-axis shows the rifampicin antibiotic concentration (AB) in *µg/mL* and the y-axis shows the *w*^ competitive index (CI). Points show the data and lines show the mean predictions with error- bars depicting the 95% confidence intervals for the means of the least-squares estimated linear regression. Therefore the space between the data points and the mean lines indicates the residuals of the model fitting. This plot shows us that much of the residual variation can be attributed to wild-type, antibiotic-susceptible genotypes (WT, second row) since this predictor has the biggest deviation between the data points and the model. This model was found to explain less of the variation than models with an interaction between *rpoB* genotypes and antibiotic environment.

#### 8.6.2 Models with first order interactions

**Figure S27:**
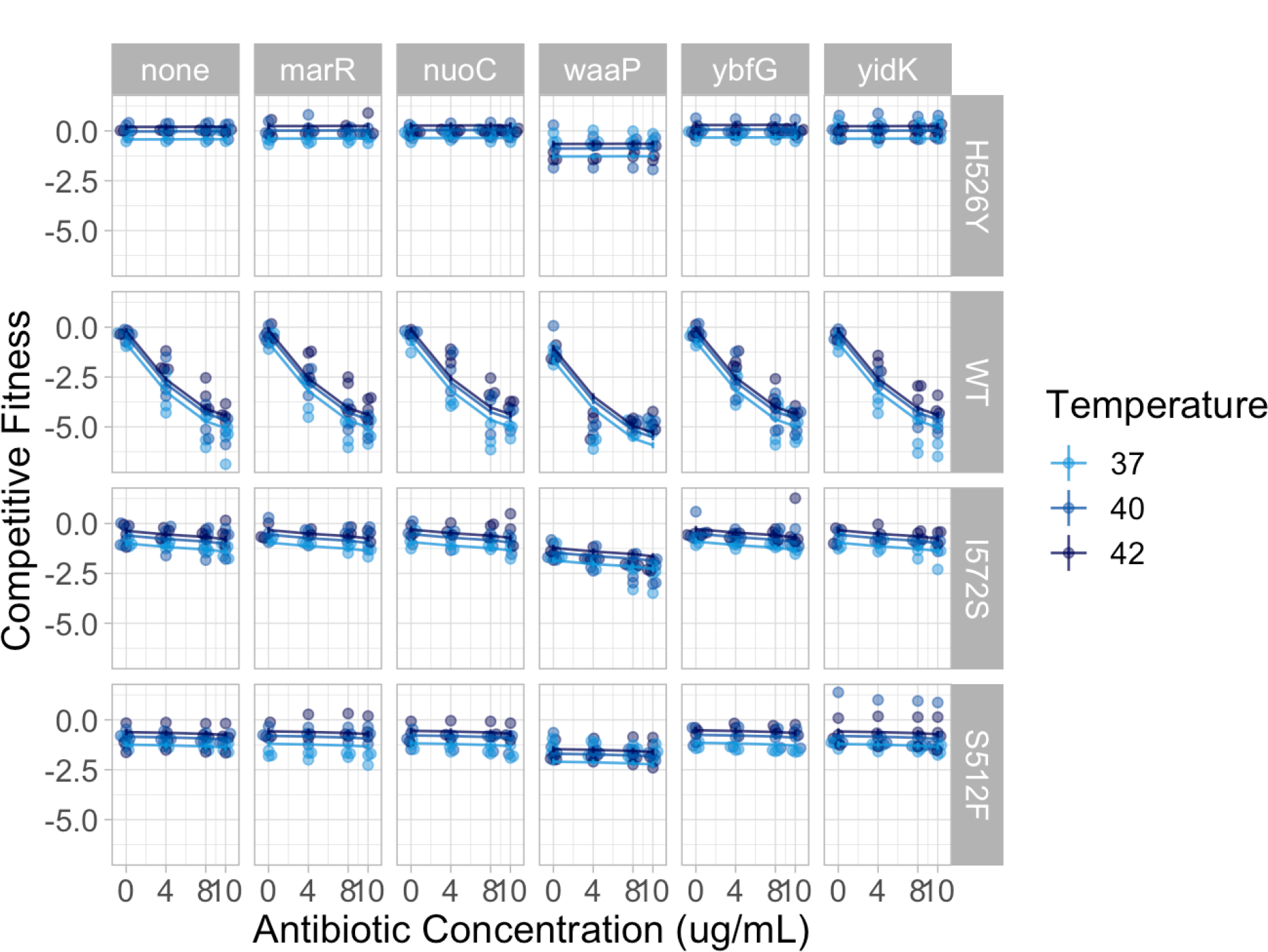
A linear regression with additive effects and *G_rpoB_ × E*_AB_, an interaction between *rpoB* genotype and antibiotic environment, explains much of the variation observed in the competition data. The x-axis shows the rifampicin antibiotic concentration in *µg/mL* and the y-axis shows the *w*^ competitive index. Points show the data and lines show the mean predictions with error-bars depicting the 95% confidence intervals for the means of the least-squares estimated linear regression. Therefore the space between the data points and the mean lines indicates the residuals of the model fitting. This plot shows that most of the systematic variation in the data is explained by this model.

**Figure S28:**
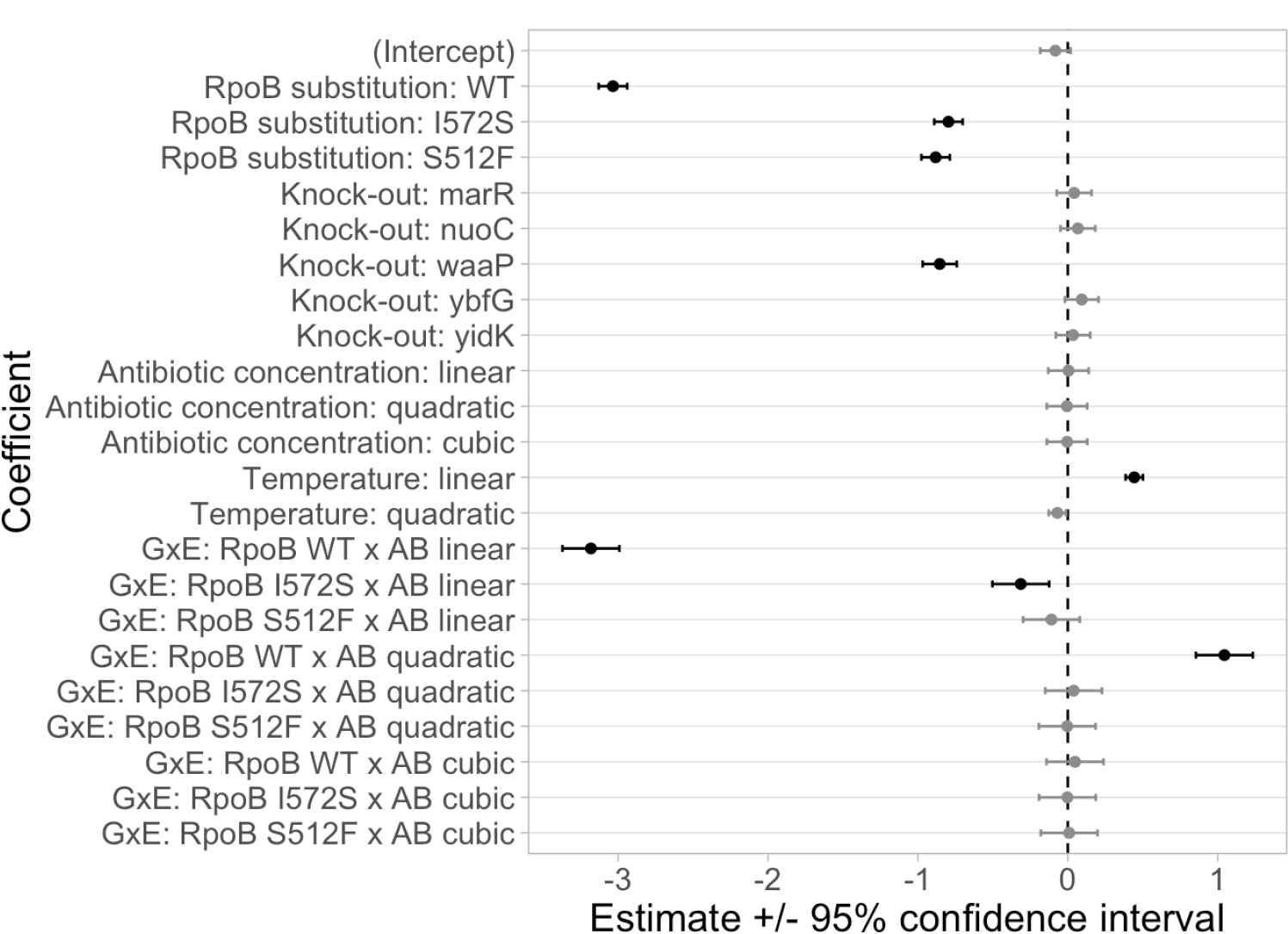
Estimated mean values for the coefficients in the regression model with additive effects and *G_rpoB_ × E*_AB_, an interaction between *rpoB* genotype and antibiotic environment (error bars show 95% confidence intervals). The intercept is for *rpoB* with the H526Y substitution and without any knock-out mutation (i.e., the reference genotype for all competitions) as grown in the 37*^◦^*C environment without any rifampicin antibiotic. Statistically significant coefficients (*α* = 0.05) are shown in black and have confidence intervals that do not overlap with zero. Coefficients whose estimates are not statistically significant are shown in gray and have confidence intervals that overlap with zero (dashed vertical line).

**Figure S29:**
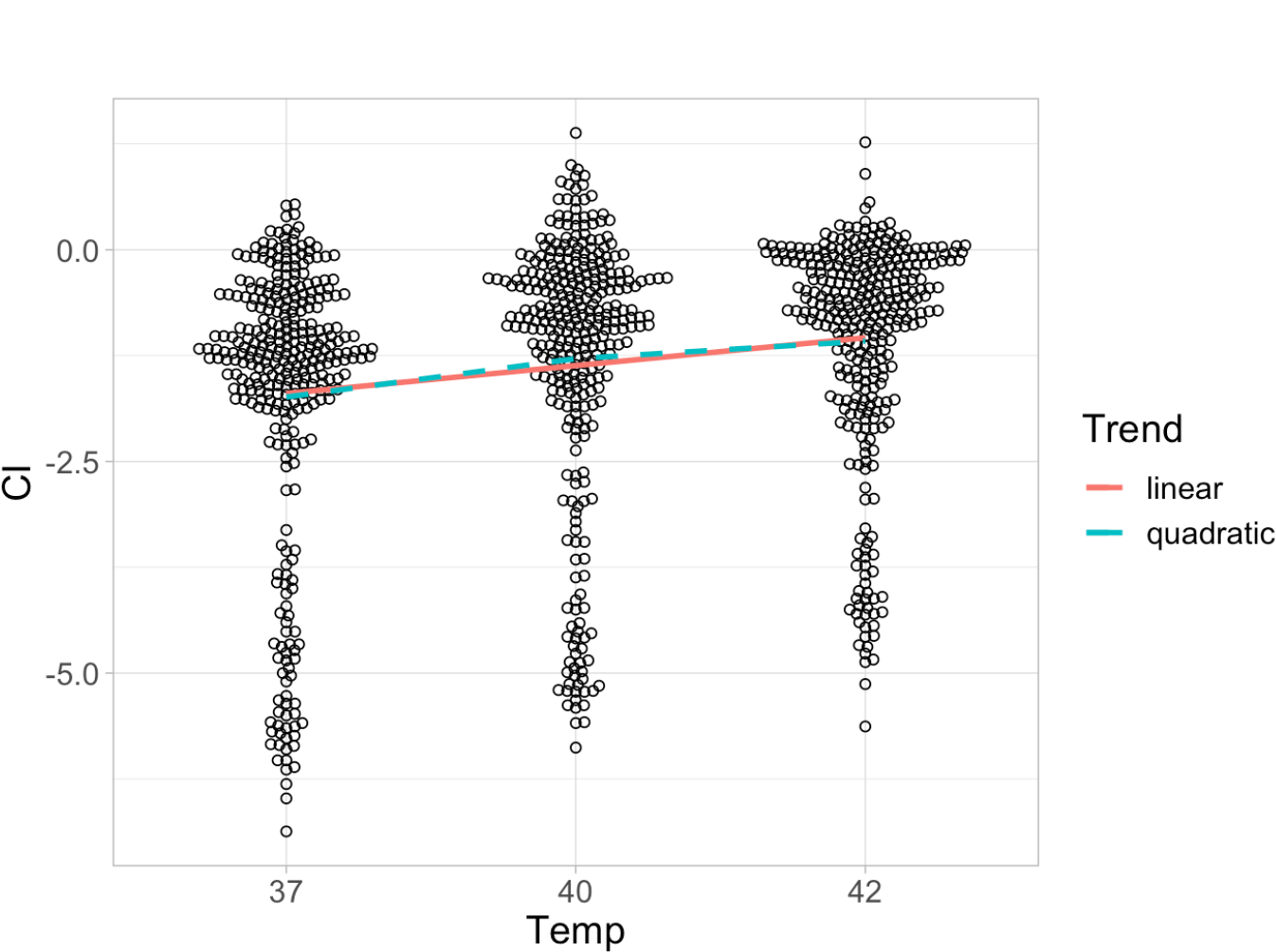
A beeswarm plot of the average effect of temperature (categorical x-axis) on estimated competitive index (quantitative y-axis) across all genotypes and antibiotic environments. Circles show individual estimates, but deviations along the x-axis away from the category labels are not quantitatively meaningful: it’s merely to prevent overlapping of dense data points. The solid line shows the average linear (statistically significant) trend, and the dashed line shows the average quadratic (but **not** statistically significant) trend (see estimated coefficients in figures S28 and S31).

**Figure S30:**
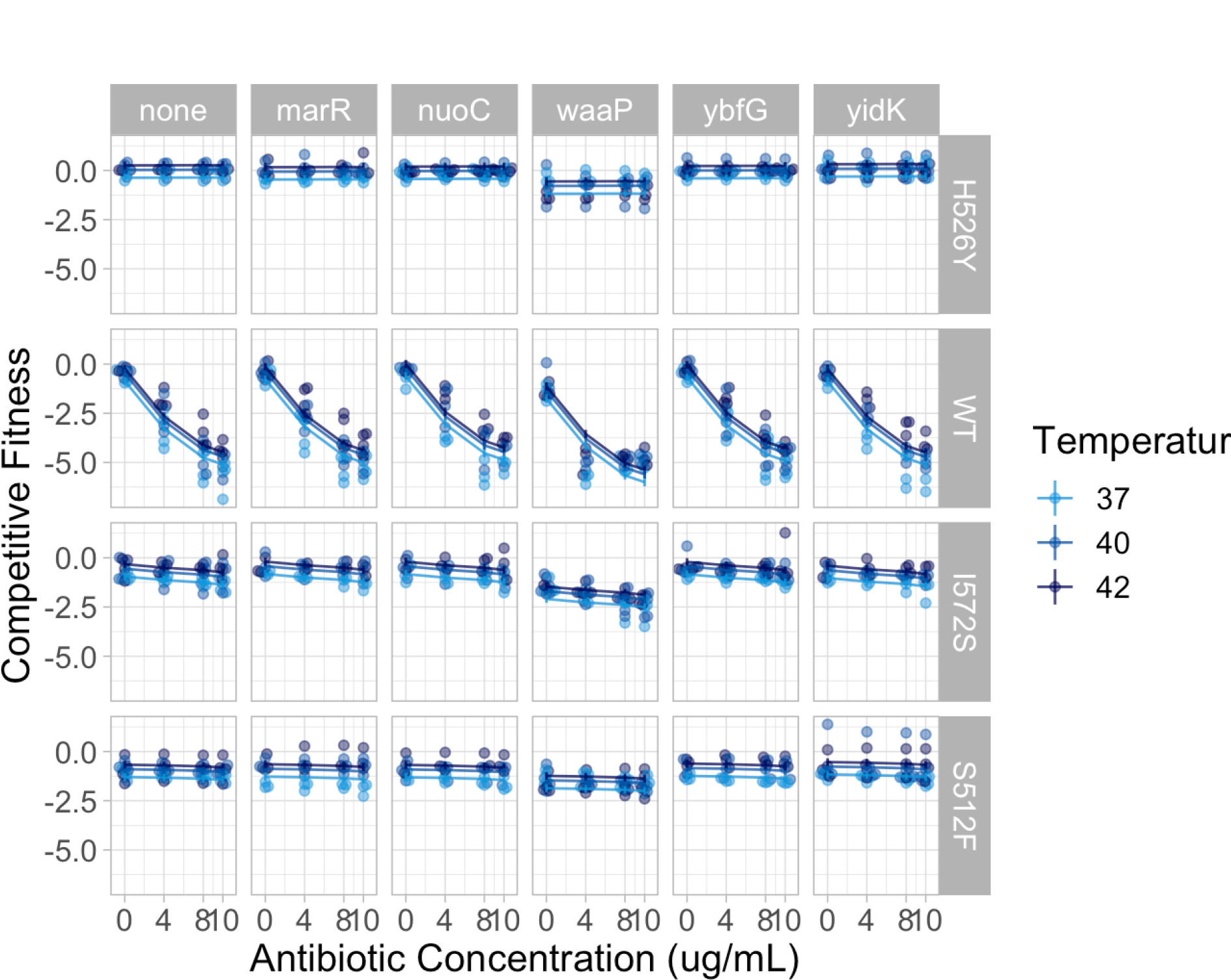
A linear regression with additive effects, an interaction between *rpoB* genotype and antibiotic environment (*G_rpoB_ × E*_AB_), and a pairwise epistatic interaction between *rpoB* and knock-out genotypes (*G_rpoB_ × G*_KO_) explains much of the variation observed in the competition data. The x-axis shows the rifampicin antibiotic concentration in *µg/mL* and the y-axis shows the *w*^ competitive index. Points show the data and lines show the mean predictions with error-bars depicting the 95% confidence intervals for the means of the least-squares linear regression. Therefore the space between the data points and the mean lines indicates the residuals of the model fitting. This plot shows that most of the systematic variation in the data is explained by this model.

**Figure S31:**
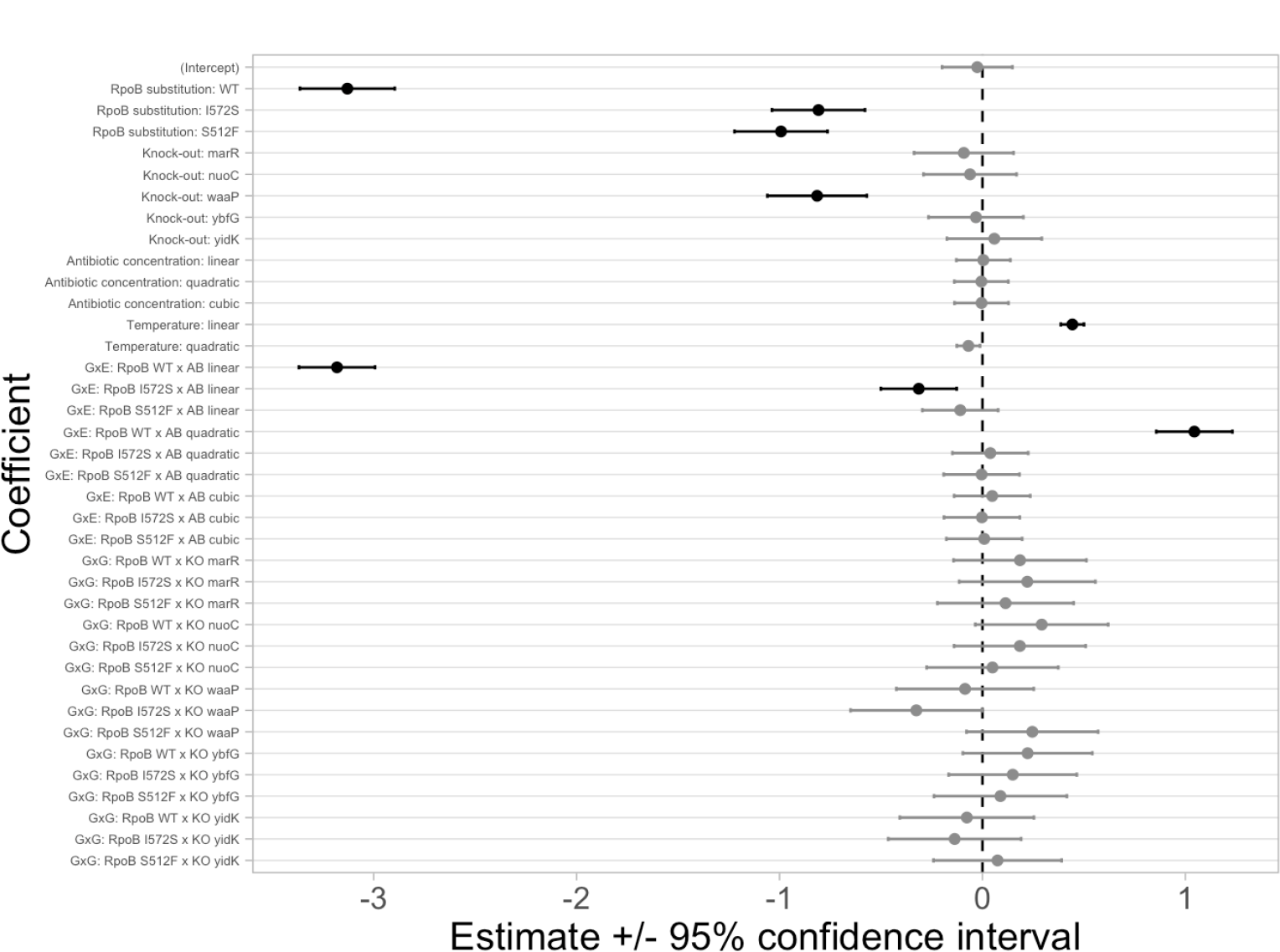
Estimated mean values for the coefficients in the regression model with additive effects and an interaction between *rpoB* genotype and antibiotic environment (*G_rpoB_ × E*_AB_) as well as epistasis between *rpoB* and knock-out genotypes (*G_rpoB_ ×G*_KO_). Error bars show 95% confidence intervals. The intercept is for *rpoB* with the H526Y substitution and without any knock-out mutation (i.e., the reference genotype for all competitions) as grown in the 37*^◦^*C environment without any rifampicin antibiotic. Statistically significant coefficients are shown in black and have confidence intervals that do not overlap with zero. Coefficients whose estimates are not statistically significant are shown in gray and have confidence intervals that overlap with zero (dashed vertical line). Notice that none of the coefficients associated with the epistatic interactions of *rpoB* and knock-out genotypes are significant. The interaction of *rpoB* I572S with the Δ*waaP* knock-out is barely significant (*p* = 0.0492).

#### 8.6.3 Models with second order interactions

We used multiple linear regression to look at the second-order interactions of the predictor variables on competitive fitness. In order to investigate second order interactions despite the risk of overfitting the data (n=996 whereas there are *>* 150 model parameters), we fitted the following complex model without reporting the p-values:

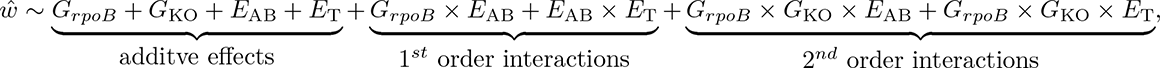

This model with 157 coefficients (i.e., just over 6 data points per estimated coefficient) resulted in an adjusted *R*^2^ = 0.916, a modest improvement as compared to the simpler models with only first order interactions.

This suggests that second order interactions may be present. Therefore, we fitted a model with the additive and first order interactions but only the second-order interaction between epistasis and temperature (*G_rpoB_ × G*_KO_ *× E*_T_). This model had a similar explanatory power as the most complex model (adjusted *R*^2^ = 0.906).

Conversely, a model with the additive and first order interactions but only the second-order interaction between epistasis and antibiotic had a lower explanatory power (*G_rpoB_ × G*_KO_ *× E*_AB_: adjusted *R*^2^ = 0.894). Thus, there may be higher-order interactions, for example between epistasis and temperature, that our data has insufficient power to reveal statistically.

### 8.7 Finlay-Wilkinson regression on overall environmental quality

#### 8.7.1 Selecting the metric of environmental quality

**Results of 3 metrics for environmental quality as part of Finlay-Wilkinson regression:**

**the mean growth of all competitor strains in each environment**: This metric of environmental quality is the best since the environment explains more of that variation in the metric than the replicate block (table S9), it is able to distinguish environments from one another (figure S32), and it is able to distinguish specialist from generalist genotypes (figure S33).

**the mean growth of the reference strain, H526Y, in each environment**: This metric of environmental quality was not found to be good since more of the variation in the metric is explained by the replicate block than by the environment itself (figure S34 and table S10).

**the mean total growth in the well (reference + competitor) in each environment**: This metric of environmental quality is not as good as the mean growth of all competitor strains. The environment explains slightly less of the variation in the metric as the replicate block (table S11) and so environments are not well distinguished from one another (figure S35). As well, this metric does not correctly distinguish specialist from generalist genotypes (figure S36)

**Figure S32:**
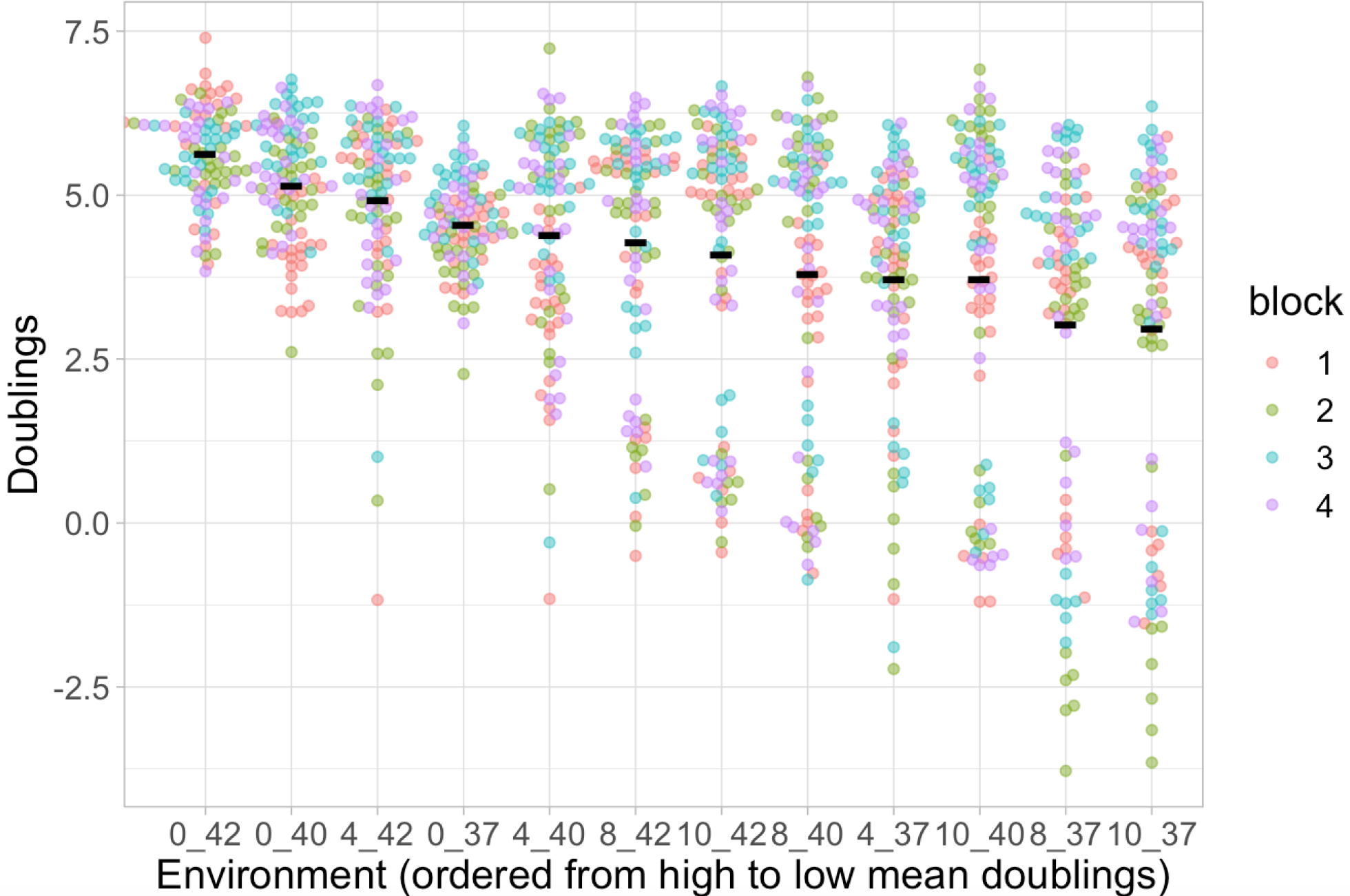
This plot shows that the environmental quality metric ‘mean growth of all competitor strains in each environment’ is able to distinguish between environments. The x-axis shows each environment in order from highest to lowest quality and the y-axis shows the total amount of growth estimated by the number of doublings observed between 0 and 20 hours (negative values suggest cell death). The black horizontal lines show the mean doublings across all replicates and genotypes: these values are used below as the metric of environmental quality. The coloured dots show the doublings of individual competitor genotypes and their replicates. Colours indicate the experimental replicate block and, ideally, should not show any trend. Both the environment and the replicate block explain a significant amount of the variation in the data (table S9). The genotypes that perform poorly in the low quality environments correspond to antibiotic susceptible *rpoB* wild-type genotypes as shown in figure S33.

**Table S9:**
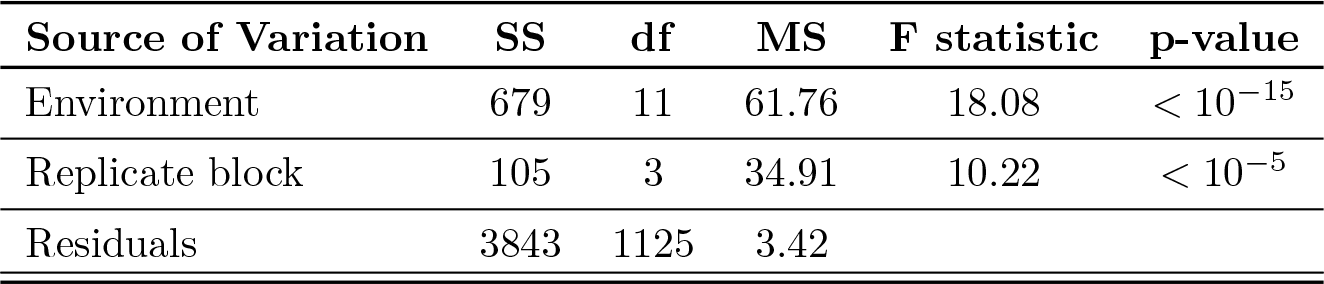
results of ANOVA for environmental quality metric: the mean growth of all competitor strains in each environment.

**Figure S33:**
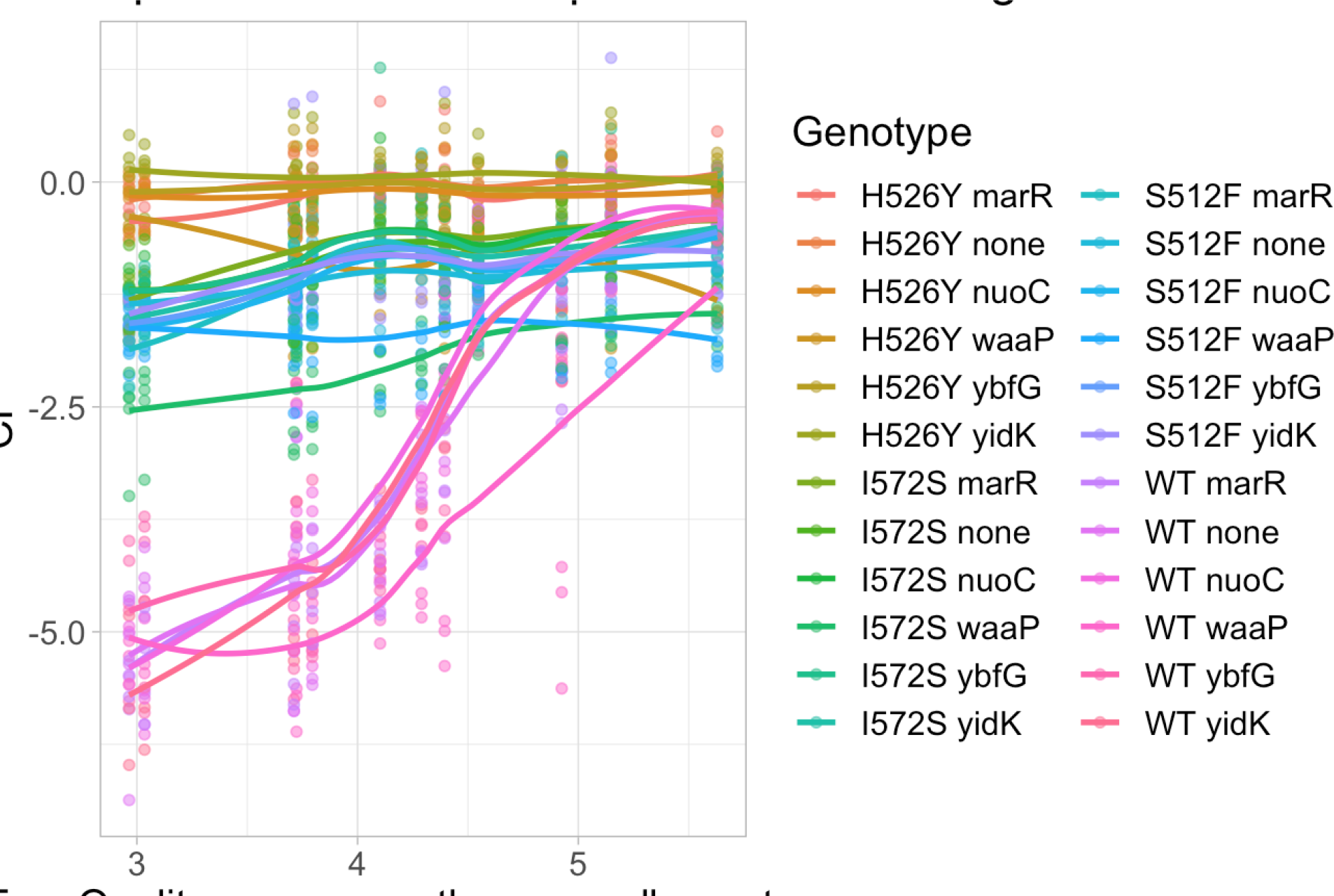
This plot shows that the environmental quality metric ‘mean growth of all competitor strains in each environment’ is able to distinguish specialist from generalist genotypes. The x-axis shows the 12 different abiotic environments as quantified by the environmental quality metric and ordered from lowest to highest quality; the y-axis shows the competitive index (*w*^) for each genotype, with values of individual replicates shown as dots (*n* = 3*−*4) and the mean value of each genotype shown as lines connected between environments.

**Figure S34:**
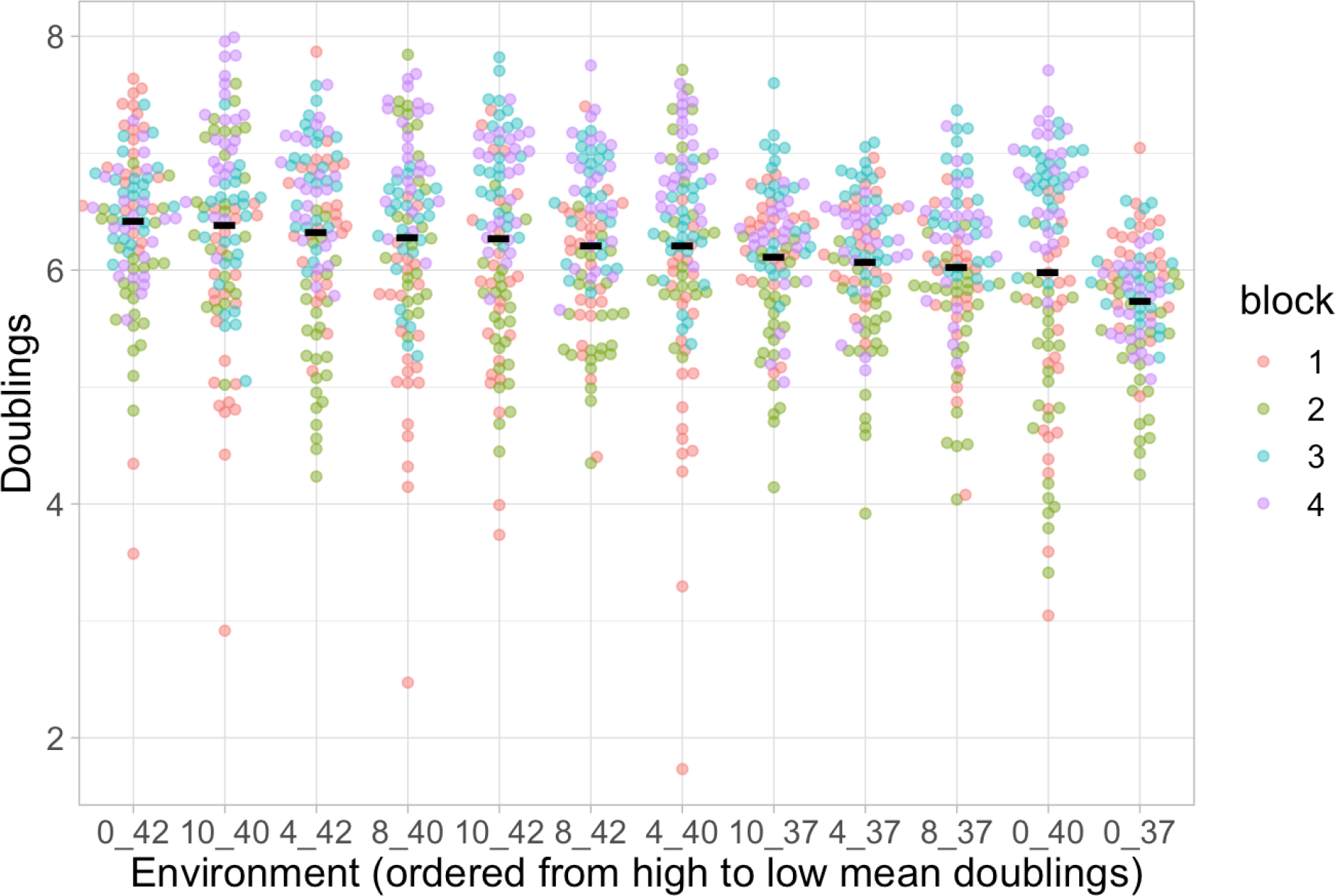
This plot shows that the environmental quality metric ‘the mean growth of the reference strain, H526Y, in each environment’ is not able to distinguish between environments. The data is plotted in the style of figure S32. However, unlike that figure, here there is a stronger effect of the replicate blocks as compared to environment (quantified in table S10).

**Table S10:**
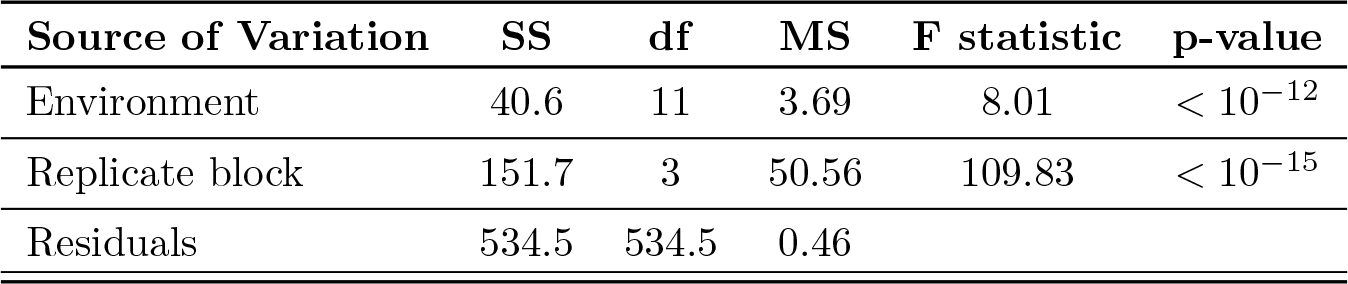
results of ANOVA for environmental quality metric: the mean growth of the reference strain, H526Y, in each environment.

**Figure S35:**
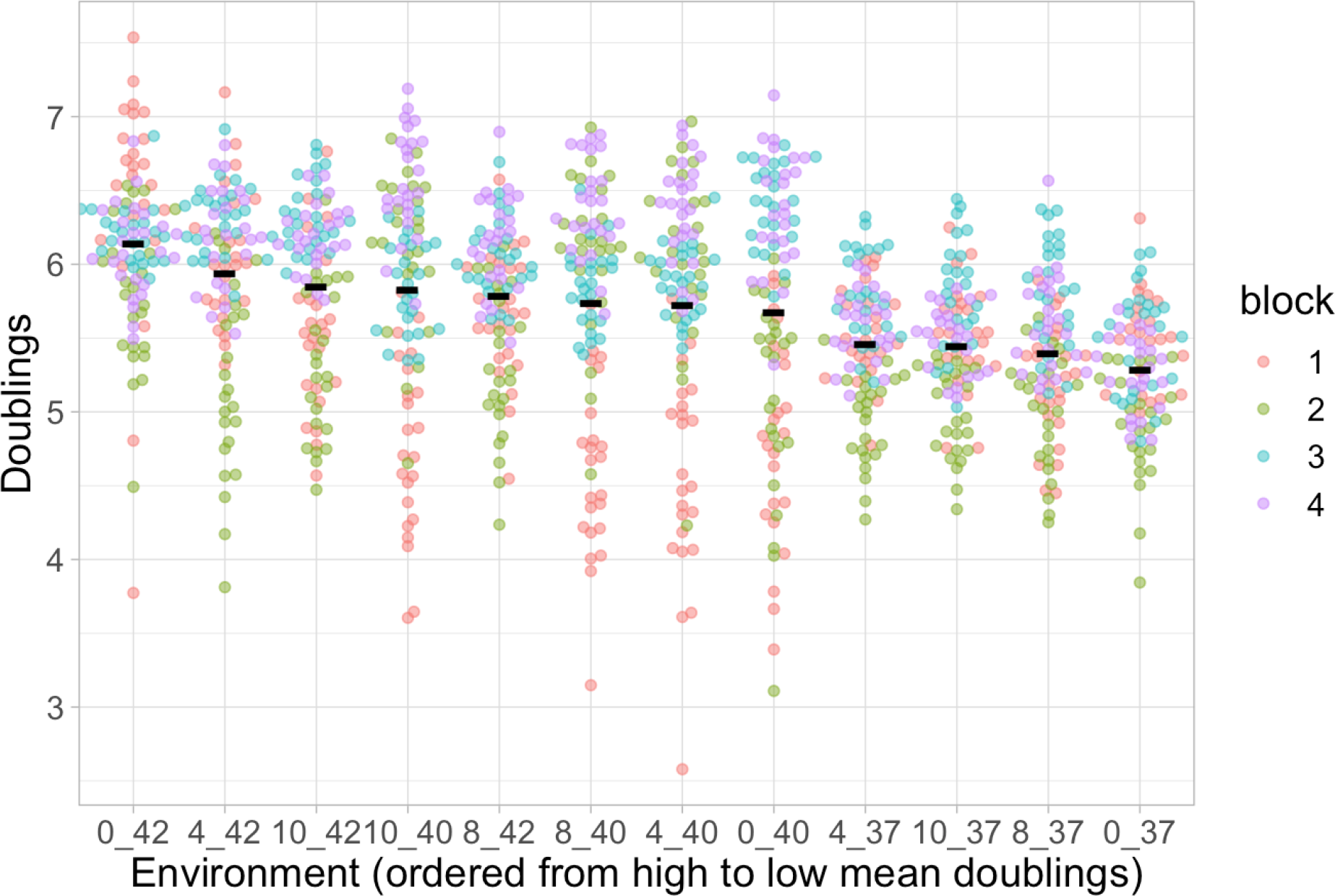
This plot shows that the environmental quality metric ‘the mean total growth in the well (reference + competitor) in each environment’ does not distinguish well between environments. The data is plotted in the style of figure S32. However, unlike that figure, here there is a stronger effect of the replicate blocks as compared to environment (quantified in table S11).

**Figure S36:**
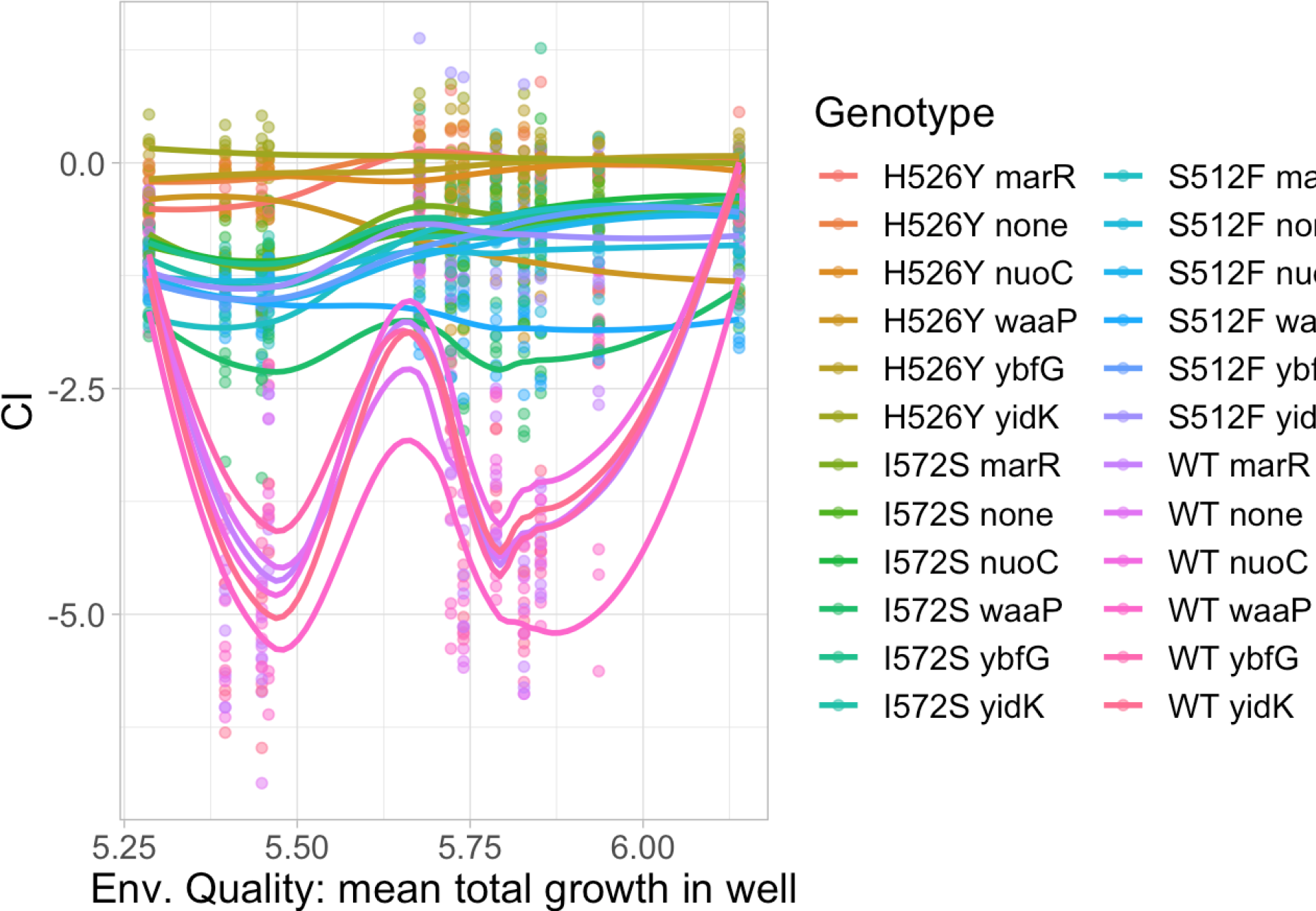
This plot shows that the environmental quality metric ‘mean total growth in the well (reference + competitor) in each environment’ is not able to distinguish specialist and generalist genotypes. The data is plotted in the style of figure S33. However, unlike that figure, here the antibiotic environments are not grouped together as low quality environments: that’s why the specialist genotypes make a ‘w’ shape in the plot.

**Figure S37:**
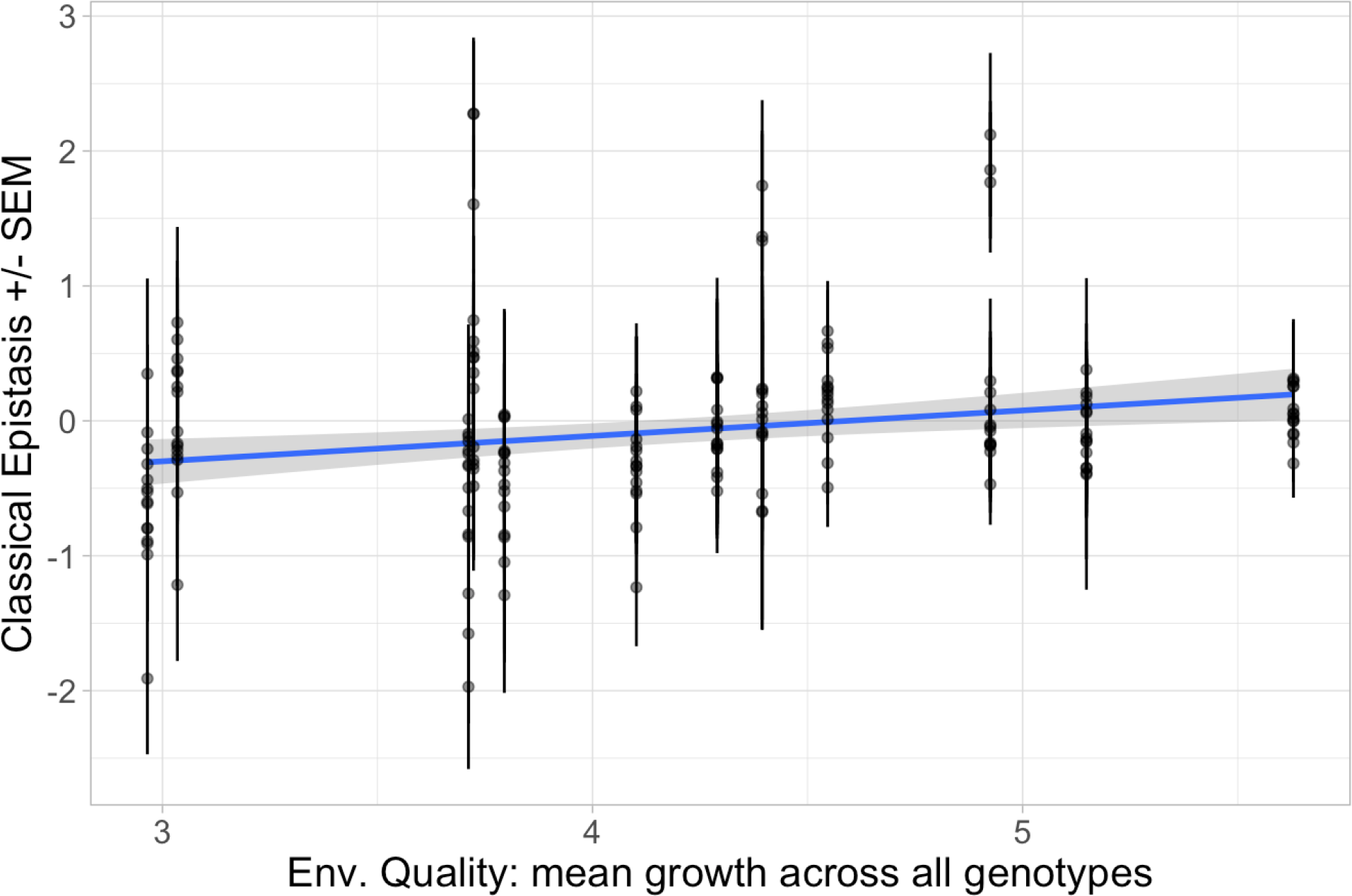
Plot of pairwise epistasis as a function of environmental quality across all genotypes and environments. The x-axis shows the environmental quality and the y-axis shows the mean pairwise epistasis for each of the 15 combinations of knock-outs and ABR *rpoB* mutations, with error bars for the standard error. The blue line shows the mean and the blue shaded region shows the 95% confidence interval of the least-squares estimated linear regression. The linear regression is significant (*F* (1, 178) = 10.16, *p <* 0.005) but does not explain much of the variation in the data (adjusted *R*^2^ = 0.0487).

**Figure S38:**
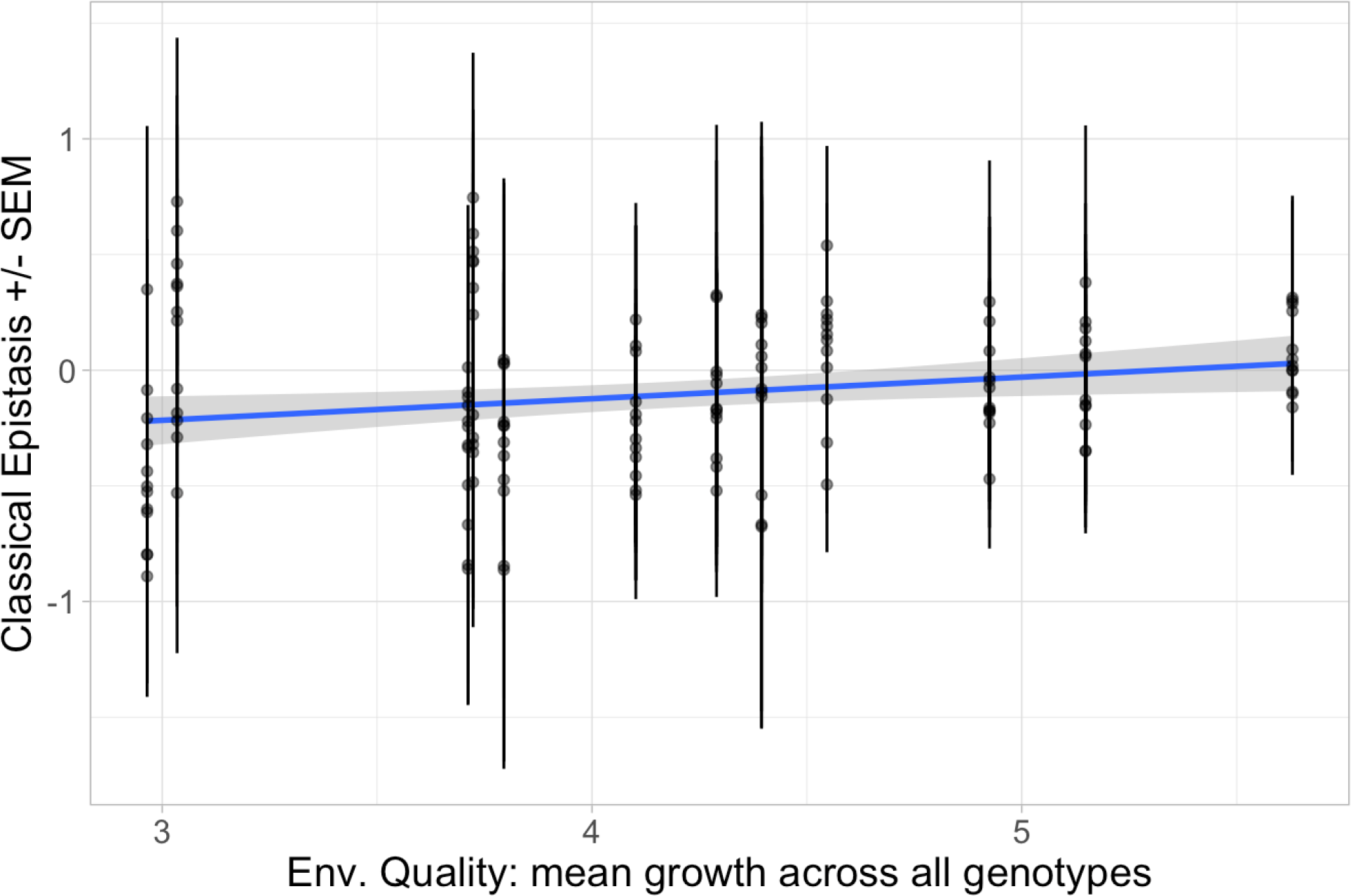
Plot of pairwise epistasis as a function of environmental quality for all gentoypes except the Δ*waaP* genotypes, as these have unusually high pairwise epistasis. This plot is similar to figure S37 except that only 12 points are shown (i.e., three Δ*waaP* genotypes were excluded) for each environment. The linear regression is marginally significant (*F* (1, 142) = 6.357, *p* = 0.0128) but it does not explain much of the variation in the data (adjusted *R*^2^ = 0.0361).

**Figure S39:**
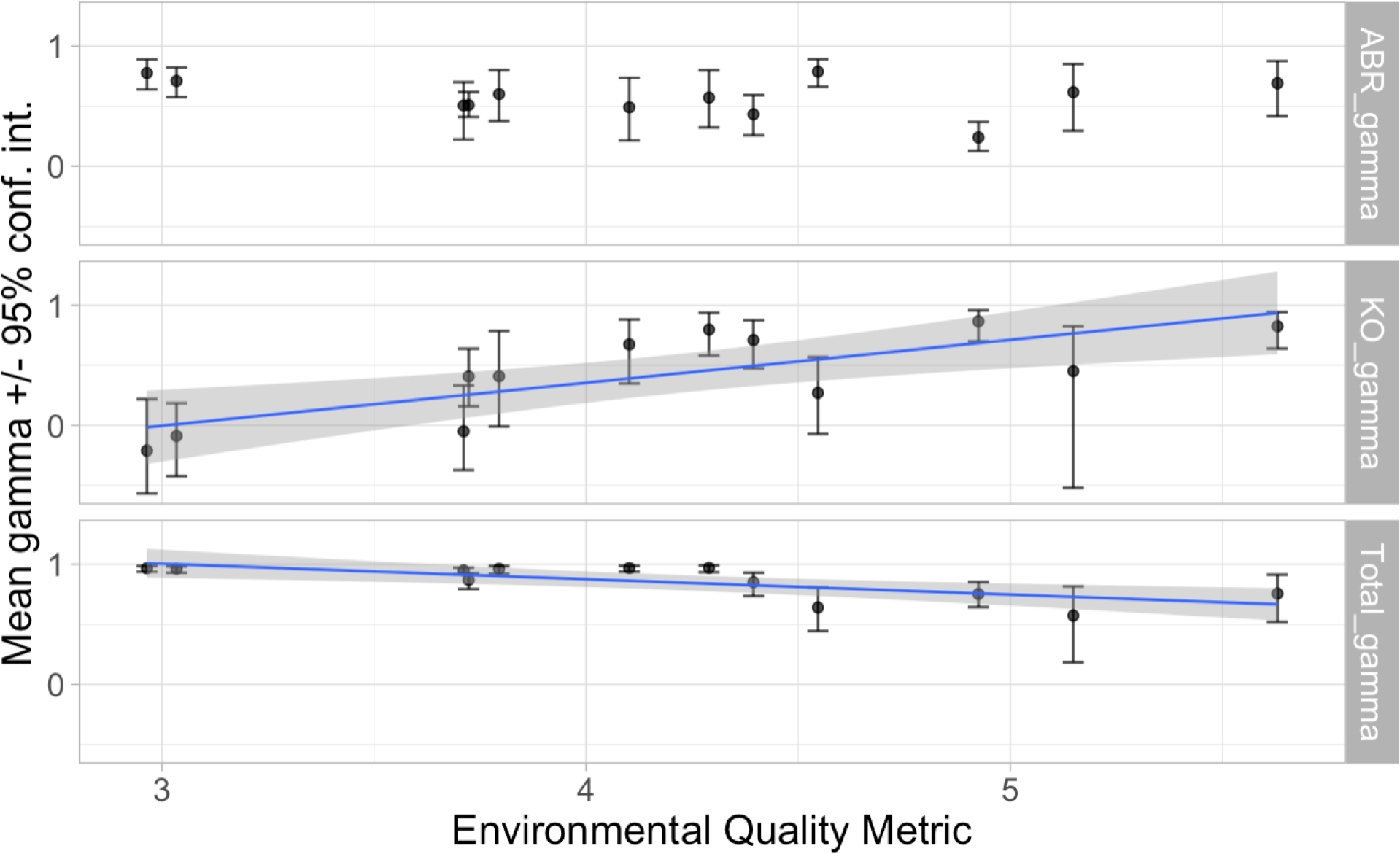
The mean gamma epistasis (y-axis) as a function of environmental quality (x-axis). The points show the mean gamma epistasis, with facets indicating the gene classes. The error bars show the 95% parametric bootstrap confidence intervals. If shown, the blue line indicates the mean of the significant, least-squares estimated linear regression and the grey shaded region shows its 95% confidence interval. Remember that gamma epistasis values near one indicate *low* G*×*G interactions and values near zero or negative one indicate *high* G*×*G interactions. **Top facet:** The gamma epistasis exhibited only by ABR *rpoB* mutations (ABR gamma) is not correlated with either environmental quality (*F* (1, 10) = 0.614, *p* = 0.45) nor antibiotic concentration alone (*F* (1, 10) = 0.0803, *p* = 0.78; see figure S22). **Middle facet:** The gamma epistasis exhibited only by knock-out mutations (KO gamma) is significantly correlated with environmental quality (*F* (1, 10) = 14.3, *p <* 0.005) and explains more of the variation in the data (adjusted *R*^2^ = 0.547) than when regressed against antibiotic concentration alone (adjusted *R*^2^ = 0.086; the regression is not significant: *F* (1, 10) = 2.04, *p* = 0.184, see figure S22). Note that this trend is opposite to that found for other measures: G*×*G is highest in the lowest quality environments and lowest in the highest quality environments. **Bottom facet:** For the total gamma epistasis, the regression on environmental quality is significant (*F* (1, 10) = 12.0, *p <* 0.01) but explains about half the variation in the data (adjusted *R*^2^ = 0.5) as compared to *∼* ^4^ of the variation in the data for the regression on antibiotic concentration alone (see main text and figure 3a).

**Figure S40:**
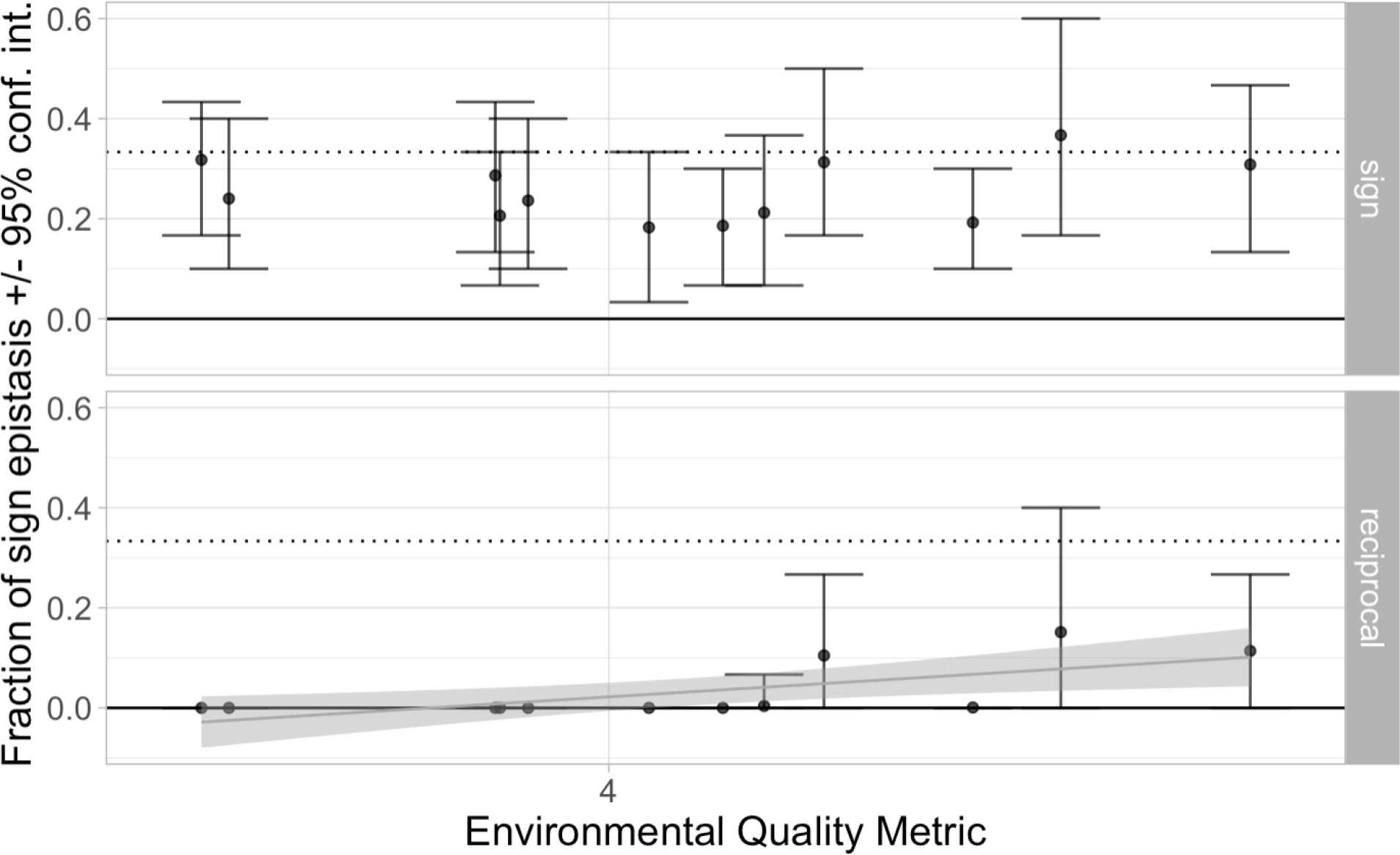
The mean fraction of simple sign (top facet) and reciprocal sign epistasis (bottom facet) as a function of environmental quality (x-axis) shows no specific trend that is an improvement over the effect of antibiotic concentration (figure S23). The points show the mean fraction and the error bars show the 95% parametric bootstrap confidence interval. There is no significant correlation between the environmental quality metric and the fraction of simple sign epistasis (*F* (1, 10) = 0.375, *p* = 0.55). For the reciprocal sign epistasis, the regression on environmental quality is marginally significant (*F* (1, 10) = 9.30, *p* = 0.0122, grey trend-line shows the mean and the shaded region shows the 95% confidence interval of the least-squares estimated linear regression). The regression of reciprocal sign epistasis explains less than half of the variation in the data (adjusted *R*^2^ = 0.430) as compared to over 60% of the variation in the data for the regression on antibiotic concentration alone (see figures 3b & S23).

**Figure S41:**
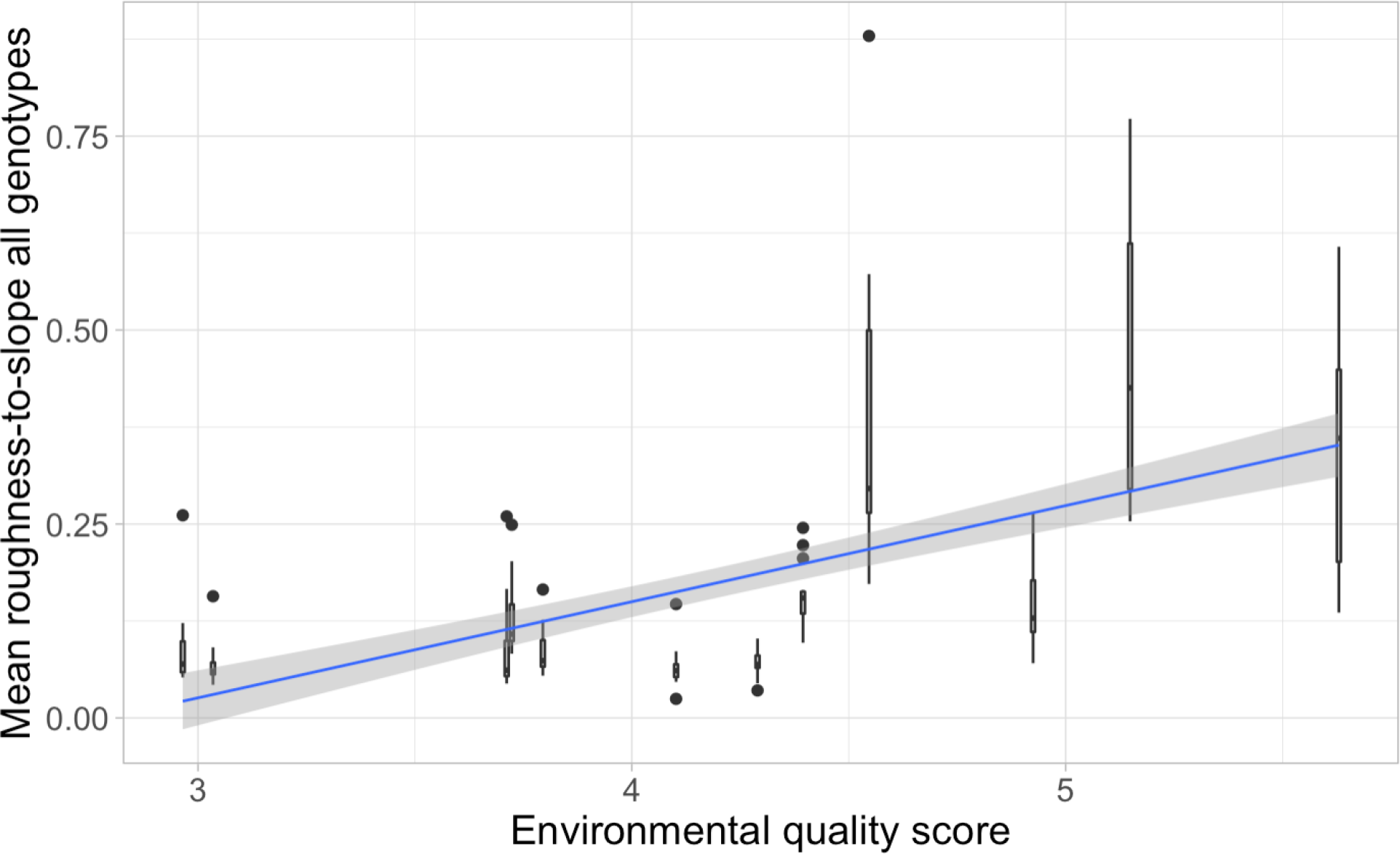
Boxplots of the roughness to slope ratio for all genes (y-axis) as a function of environmental quality (x-axis) shows a significant correlation (*F* (1, 178) = 95.7, *p <* 10*^−^*^15^) but the amount of variation explained by the environmental quality score (adjusted *R*^2^ = 0.346) is less than than explained by antibiotic concentration alone (*∼* 50%, see figure 3c).

The colours indicate the genotype. All genotypes starting with “WT” are susceptible to rifampicin and therefore considered specialists to environments with no or low antibiotic concentrations. Conversely, all other genotypes are antibiotic resistant and can be considered generalists across all environments. Environments with higher antibiotic concentrations are grouped together as low quality and therefore susceptible genotypes perform poorly at lower values of the environmental quality metric (exhibiting a ‘/’ or ‘S’ shape on the plot) and generalist genotypes perform about the same across all environments (exhibiting a straight line on the plot).

#### 8.7.2 Finlay-Wilkinson environmental quality regression for GxG summary statistics

## 9 Materials & Methods

### 9.1 Genotypes

All the strains used in this study (Table S12) derive from strains RB323 or RB324, which are derivatives of *E. coli* K12 MG1655 [81]. The strains are marked with constitutively expressed Superfolder GFP or mCherry inserted in the *ysaCD* pseudogene locus and have the entire lac operon deleted (to make constitutive the expression of the fluorescent proteins).

#### 9.1.1 Obtaining the ABR Mutations

*RpoB* H526Y and S512F mutants have strong resistance to rifampicin (*>* 200*µ*g/mL) and were already available in the Gordo lab [80]. *RpoB* I572N happened to evolve *de novo* by screening for spontaneous mutations with intermediate rifampicin resistance (resistant at 20 *µg/mL* but susceptible at 100 *µg/mL*) in rich media (Luria broth) batch culture and then using Sanger sequencing to confirm the SNP location on the *rpoB* locus.

#### 9.1.2 Genetic engineering

P1*vir* phage transduction [82] with rifampicin selective plates was used to move each *rpoB* mutant onto two different fluorescently labeled backgrounds, one with Superfolder GFP and the other with mCherry, both with a chloramphenicol resistance cassette.

**Table S11:**
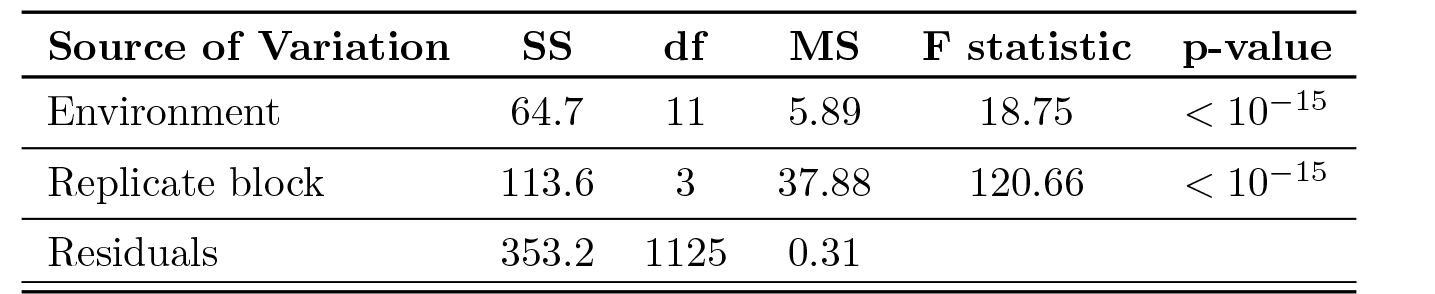
results of ANOVA for environmental quality metric: the mean total growth in the well (reference + competitor) in each environment.

**Table S12:**
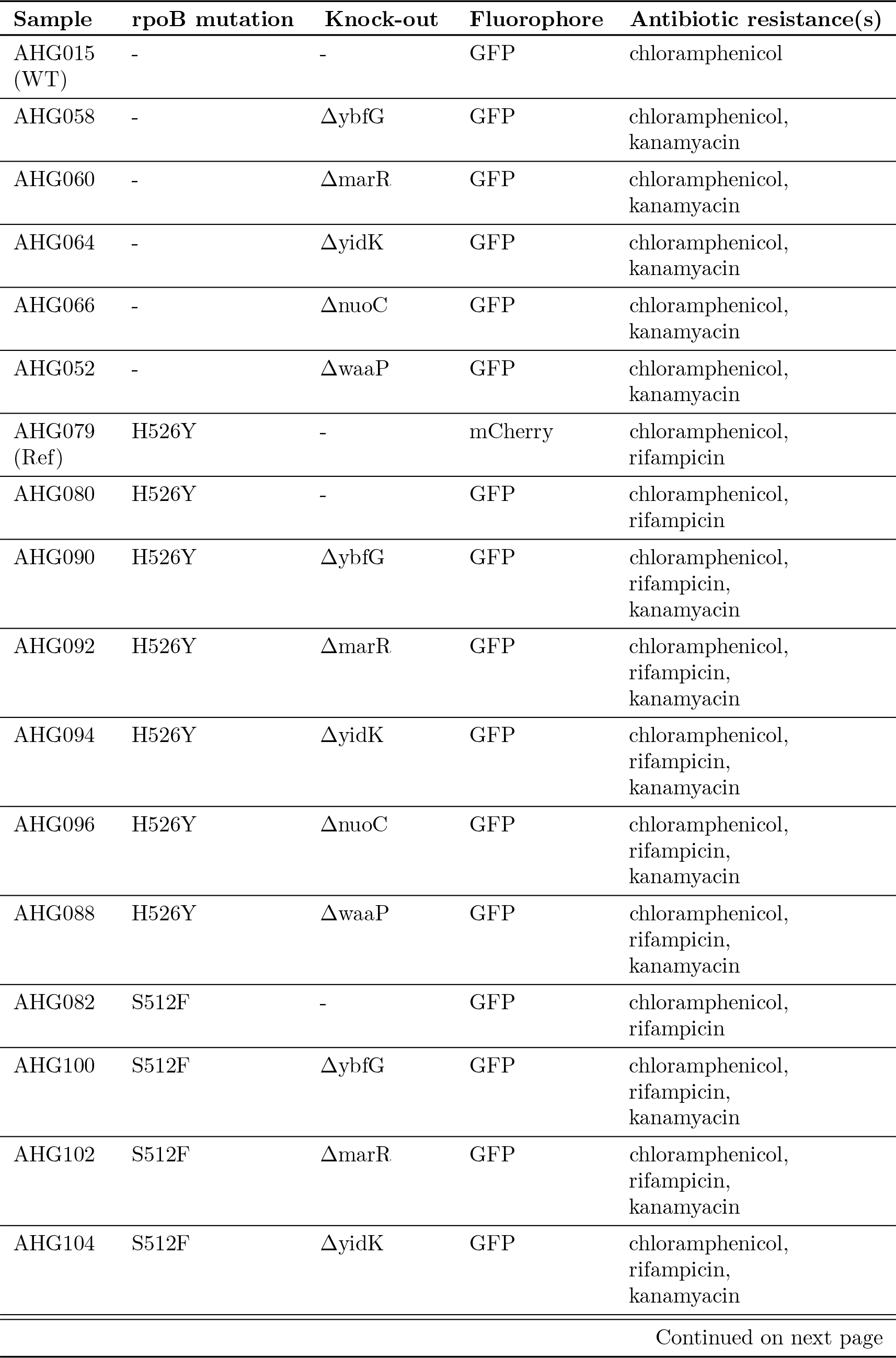
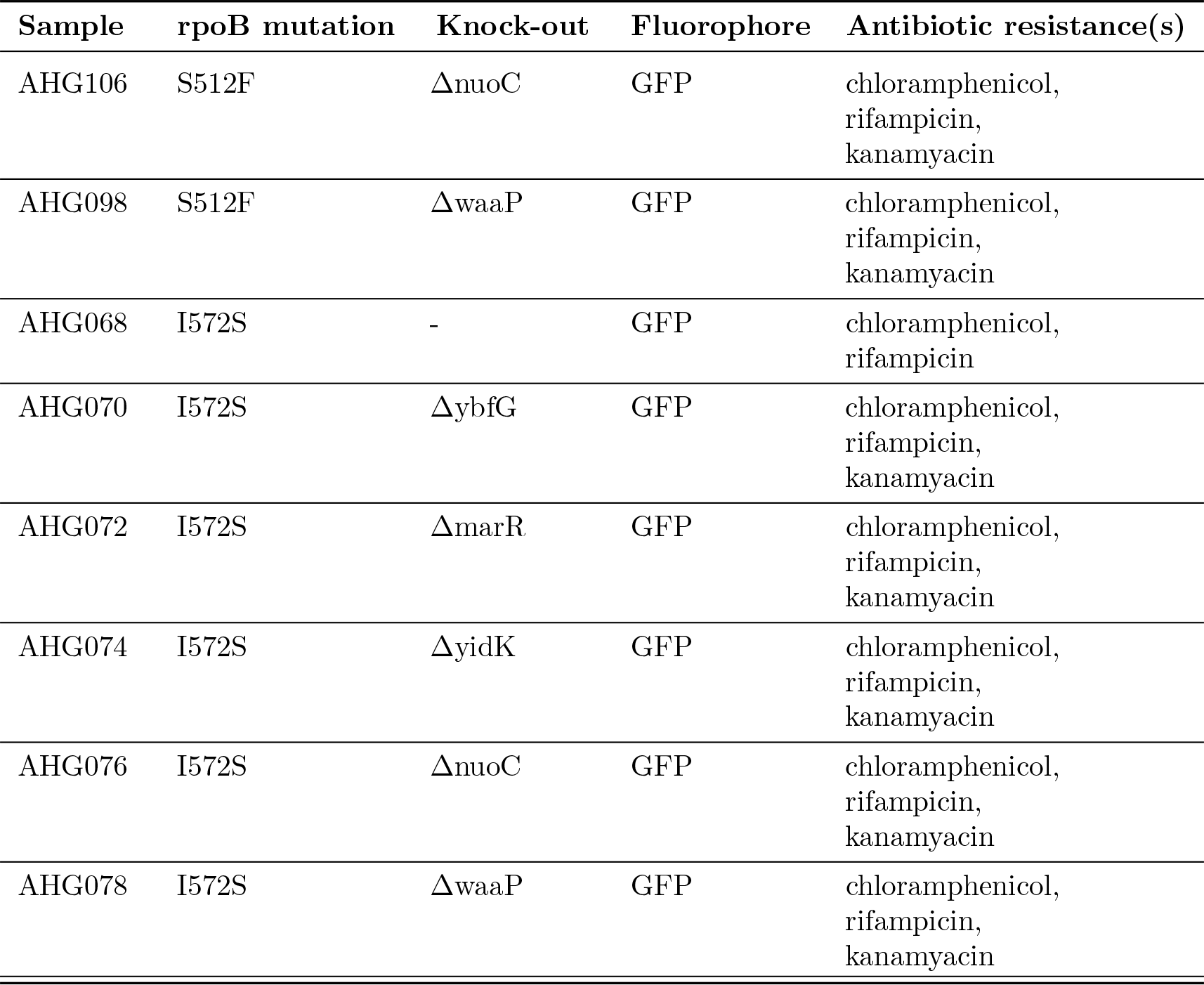
Sample names and their corresponding engineered mutations.

**Table S13:**
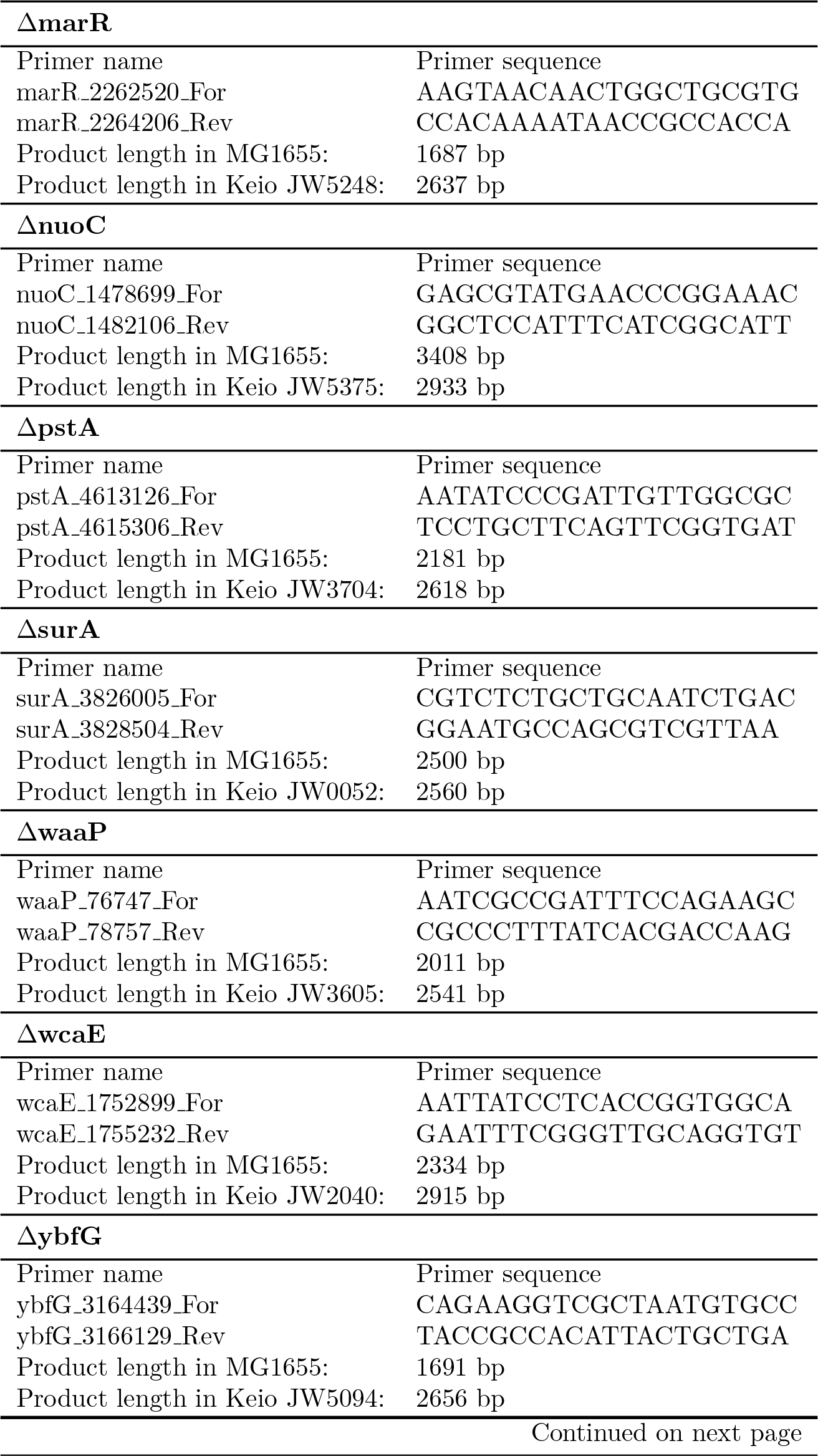
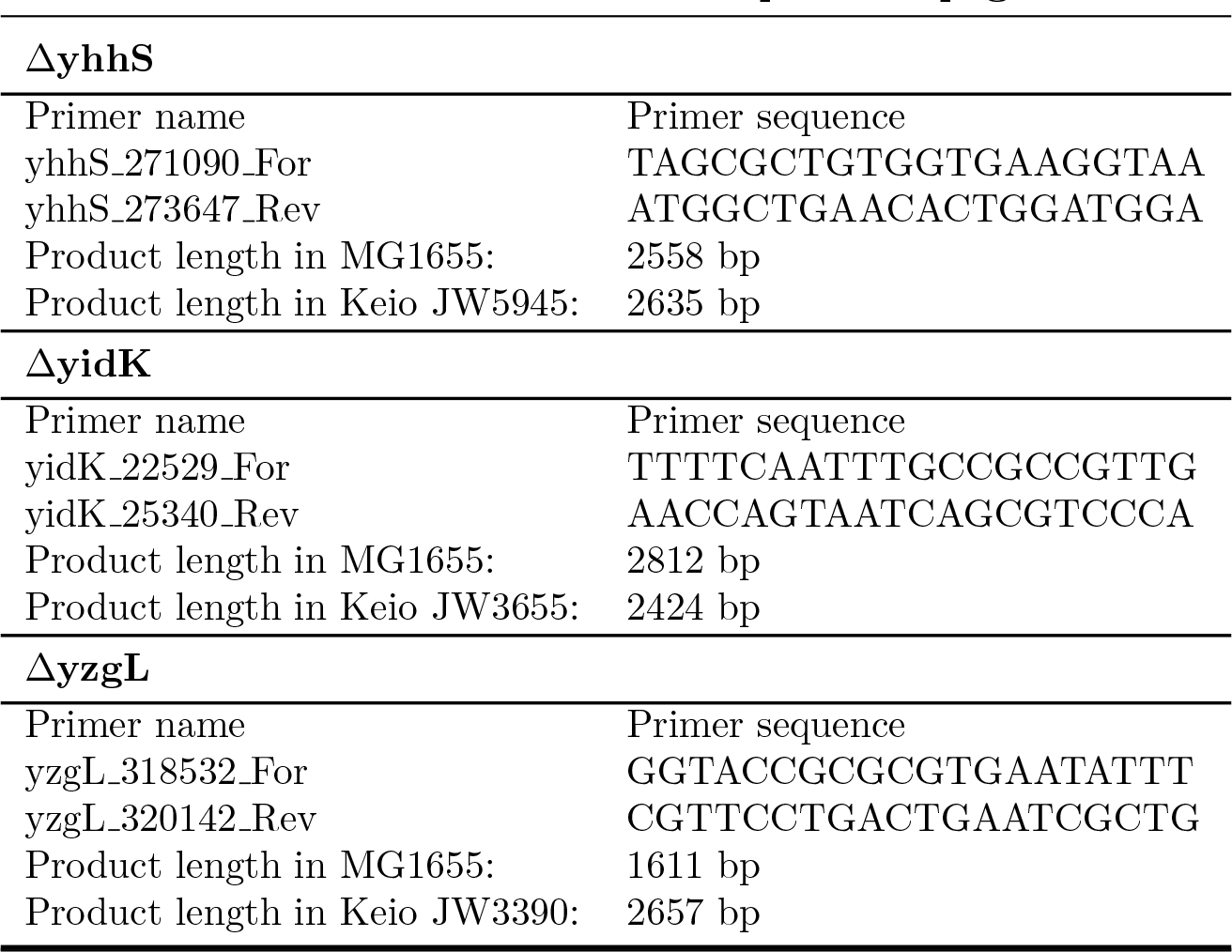
The sequences of the homology primers used to PCR amplify the in-frame, single gene deletion mutants from the listed Keio collection samples, respectively. Product lengths are given for when the gene is present in MG1655 and for when the gene is knocked-out in the Keio sample. The names of the primers indicate their positions in the MG1655 reference genome (GenBank accession CP032667.1). PCR for the Datsenko-Wanner recombineering was done with New England Biolabs Phusion High-Fidelity DNA Poly- merase (M0530) by following the manufacturer’s guidelines with a 2.5 minute elongation at 67*^◦^*C.

Knock-out single mutants were built onto the *rpoB* wild-type GFP-labeled and mCherry-labeled backgrounds by *λ* Red-mediated recombination using PCR products [83]. The homology primers listed in table S13 were used to PCR amplify the in-frame, single gene deletion mutants from the Keio collection samples, then these PCR products were used as templates for recombineering. P1 phage transduction with selection for the inserted kanamycin cassette was then used to clean-up any off-target mutations that may have been introduced by the *λ* Red recombinase. However, we did not counterselect to eliminate the kanamycin resistance cassette from knock-outs – meaning that all knock-outs have the gene-of-interest replaced with a kanamycin cassette.

Double-mutants were built by moving the knock-outs onto the *rpoB* mutant backgrounds, again using P1 phage transduction. All sample codes and their corresponding engineered mutations are listed in table S12.

All genotypes were phenotypically screened for their respective markers: resistance to chloramphenicol, susceptibility/resistance to rifampicin and kanamycin, and GFP or mCherry fluorescence. Fluorescence was screened using a BioTek H1 microplate reader with GFP (ex: 469 nm / em: 525 nm) and RFP (ex: 531 nm / em: 593 nm) filters.

#### 9.1.3 Whole genome sequencing and analysis

Single colonies for each of the 25 competed genotypes were grown up in 10 mL of rich media (Luria broth) batch culture so that DNA could be extracted and purified using the standard phenol/chloroform method. All samples were sent for whole genome sequencing to the Next Generation Sequencing (NGS) Platform of UBern in Bern, Switzerland. Library preparation was done using the Illumina DNA Prep kit and sequencing was done by Illumina NovaSeq6000 flow cell, 2 x 150 bp run on two lanes.

**Details about WGS analysis:** The breseq pipeline (0.36.1), with bowtie2 (2.3.3) and R (4.1.1), was used to map reads to reference sequences in order to confirm the genetically engineered mutations-of-interest and discover any new, off-target mutations. The MG1655 reference sequence from GenBank (NC 000913.3) and the inserted GFP fluorescent construct sequence (given in supplementary methods) were used as reference sequences for the wild-type assigned GFP-labelled genotype (sample AHG015). Then, all subsequent single and double mutant genotypes were mapped against this wild-type genotype, so that any mutations common to all genotypes could be ignored in down-stream analysis. In addition to the wild-type reference, reads from single and double mutant genotypes also used the GenBank sequence for the pDK4 plasmid (AY048743.1) as an additional reference in order to identify knock-outs where target genes have been swapped with the kanamycin resistance cassette. Finally, for the mCherry-labelled reference genotype (sample AHG079), the inserted mCherry fluorescent construct sequence (given in supplementary methods) was used as a reference in order to identify the correct fluorescent tag on this genotype. An average read depth limit of 120x was set for breseq as recommended in the manual for efficient completion times: this means that up to a quarter of the read data was discarded per sample, regardless of the actual quality of that data (see tables S1-S2). The different *rpoB* substitutions that were engineered for all samples were readily detected by breseq’s mutation prediction algorithm (table S3). The presence of engineered gene knock-outs was confirmed manually by looking for consistent results between three types of evidence: 1) high coverage of the kanamyacin coding sequence from the pKD4 plastmid (figure S6), 2) unassigned missing coverage evidence consistent with a knock-out of the target gene (figure S8), and 3) unassigned new junction evidence both upstream and downstream of the target gene (figure S11).

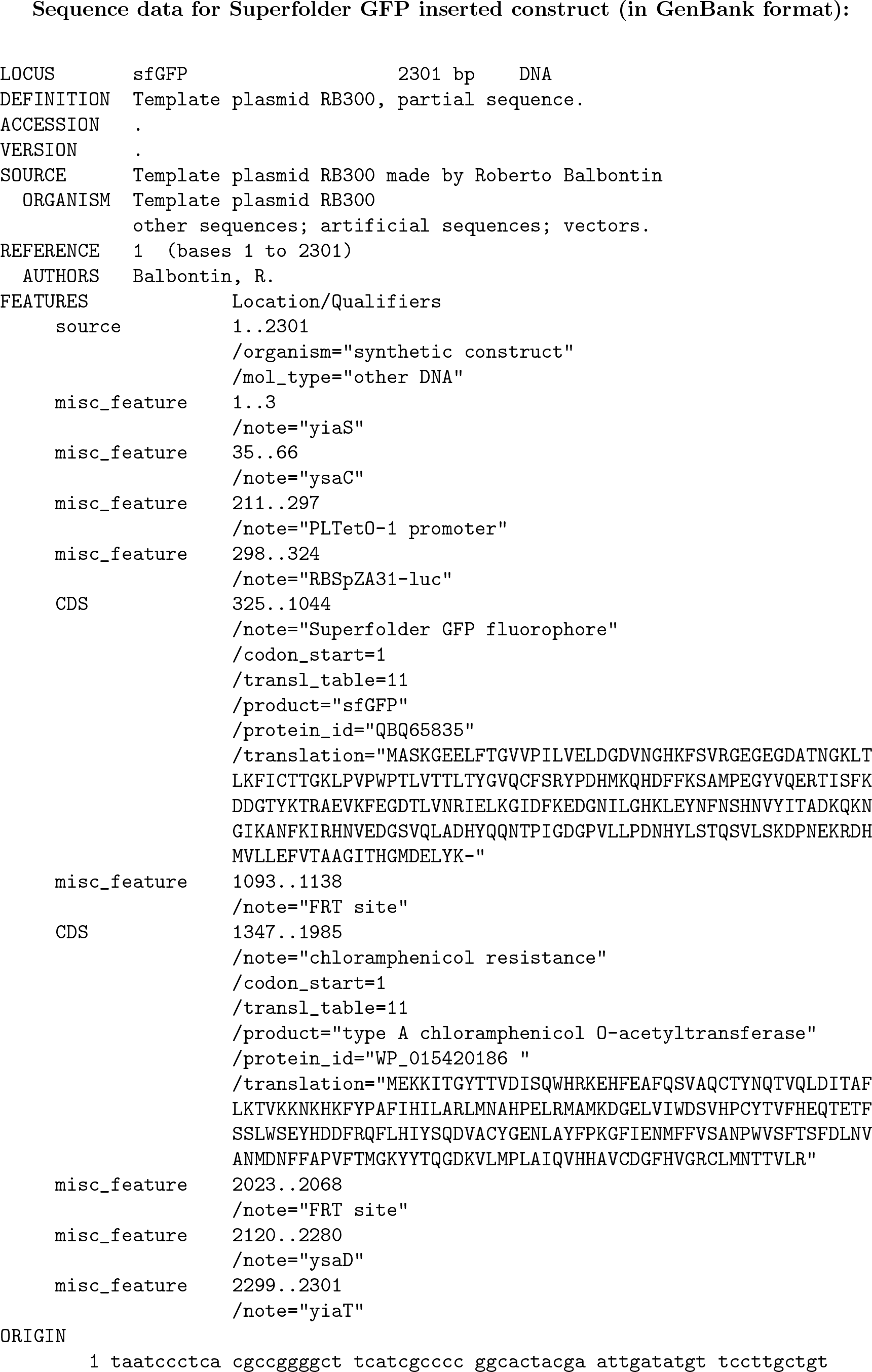
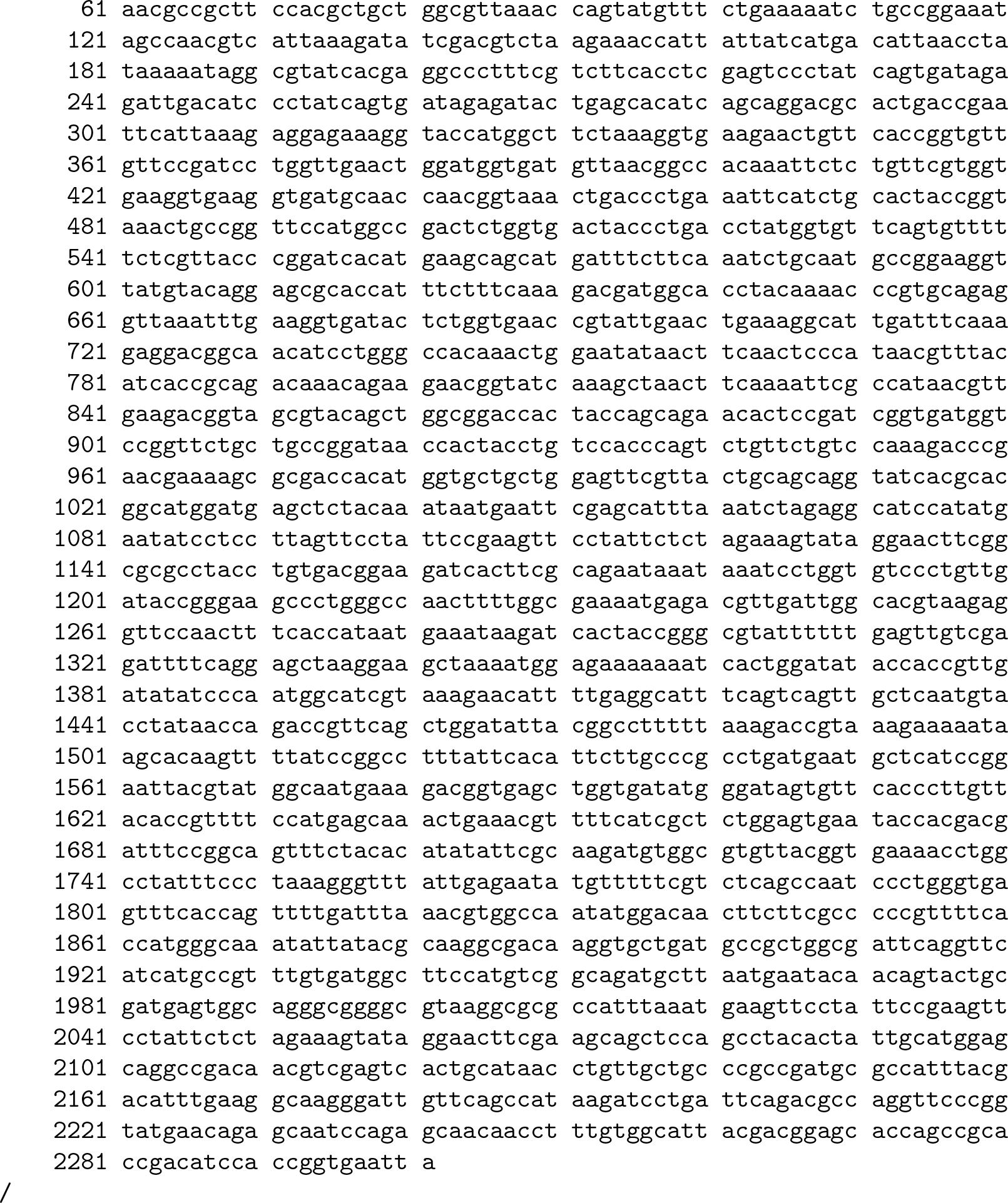
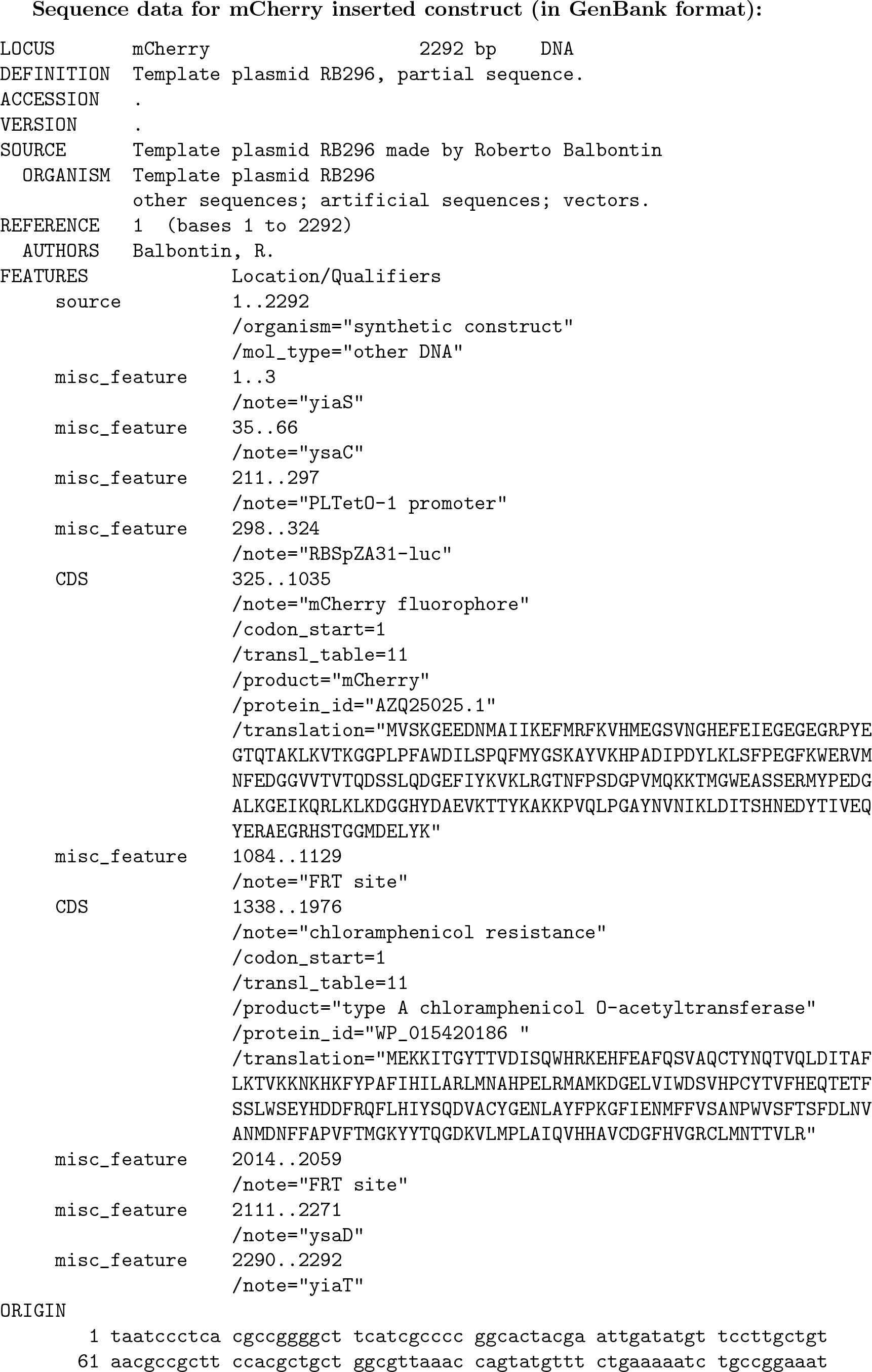
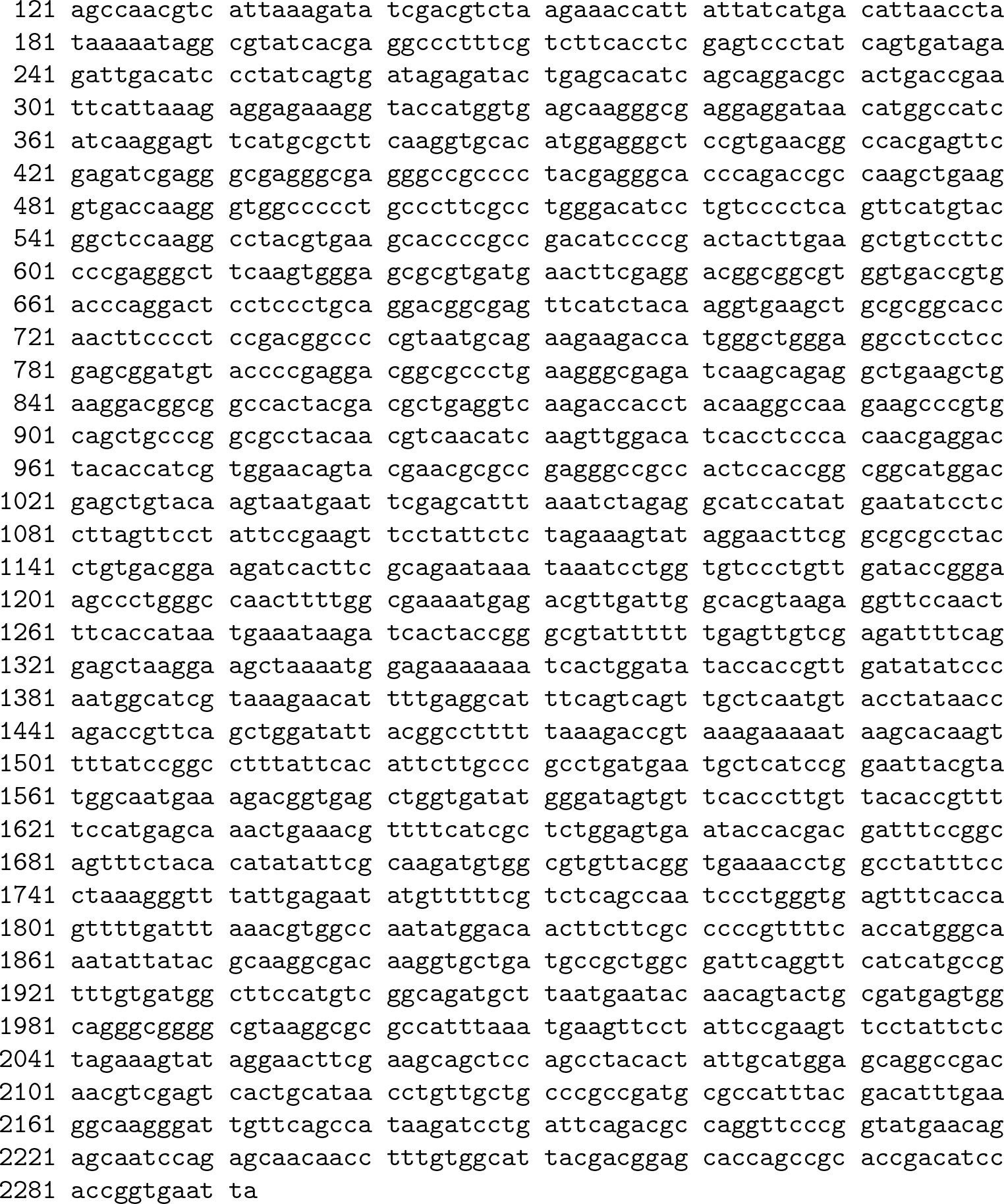

### 9.2 Estimating the competitive fitnesses

#### 9.2.1 Cell growth protocol

The protocol is summarized in figure S42. Freezer stocks contain 15% glycerol and LB. Chloramphenicol antibiotic (CM) was used to prevent contamination but only with agar plates, not at any later steps. M9 liquid minimal media with 0.4% glucose was used for acclimitization and fitness measures. Glass inoculum tubes were used for acclimitization as these were found to provide superior culture aeration as compared to plastic well plates. Batch culture competitions were done in flat-bottom wells sealed with breathable film and incubated in a BioTek H1 microplate reader with continuous shaking in order to provide a homogeneous environment that minimized clumping.

Each 96-well plate contained 24 genotypes competing at each of the four rifampicin antibiotic concentrations. Media with different concentrations of rifampicin was aliquoted to the plate in randomly ordered blocks of two rows. As well, the locations of the samples on the plate was randomized to prevent bias from possible plate edge effects.

Only one plate was incubated at a time in the microplate reader. Therefore sample measures are grouped into four blocks, each with three temperature environments. The order of temperature environments was randomized to prevent any systematic bias that may arise due to the duration of time that cells were stored at 4*^◦^*C.

All competitions began at a fixed inoculum size of 10^5^ cells and equal (1:1) ratios of the competitor and reference genotypes. Flow cytometry was for three measures. First, it was used on the early stationary phase cultures to determine the precise dilution factor required for each sample in order to arrive at an inoculum size of 10^5^ cells. Flow cytometry was also performed on the inocula after they were used for aliquoting to the 96-well plate: this way, the precise initial frequencies of competitor vs. reference are known, not assumed. Finally, flow cytometry was used to measure the final frequencies of competitor vs. reference after 20 hours of growth.

#### 9.2.2 Flow cytometry protocol

Flow cytometry with fluorescent beads was used to determine the standardized relative cell counts. All flow cytometry experiments were performed on a BD LSR Fortessa SORP machine running the BD FACSDiva software at the Flow Cytometry Facility of Instituto Gulbenkian de Cîencia. Cell counts were extracted from flow cytometry data using FlowJo (10.7.1). Bacterial cells were separated from fluorescent beads and debris using side-scatter and forward-scatter gating, next doublets were excluded using side-scatter area and width, then, finally, GFP-only expressing cells were distinguished from mCherry-only expressing cells using rectangular subset gating. The raw flow cytometry data, FlowJo quantification files, and cell counts data are publicly available in a GitLab repository, https://gitlab.com/evoldynamics/epistasis-decreases-with-increasing-antibiotic-pressure.

**Figure S42:**
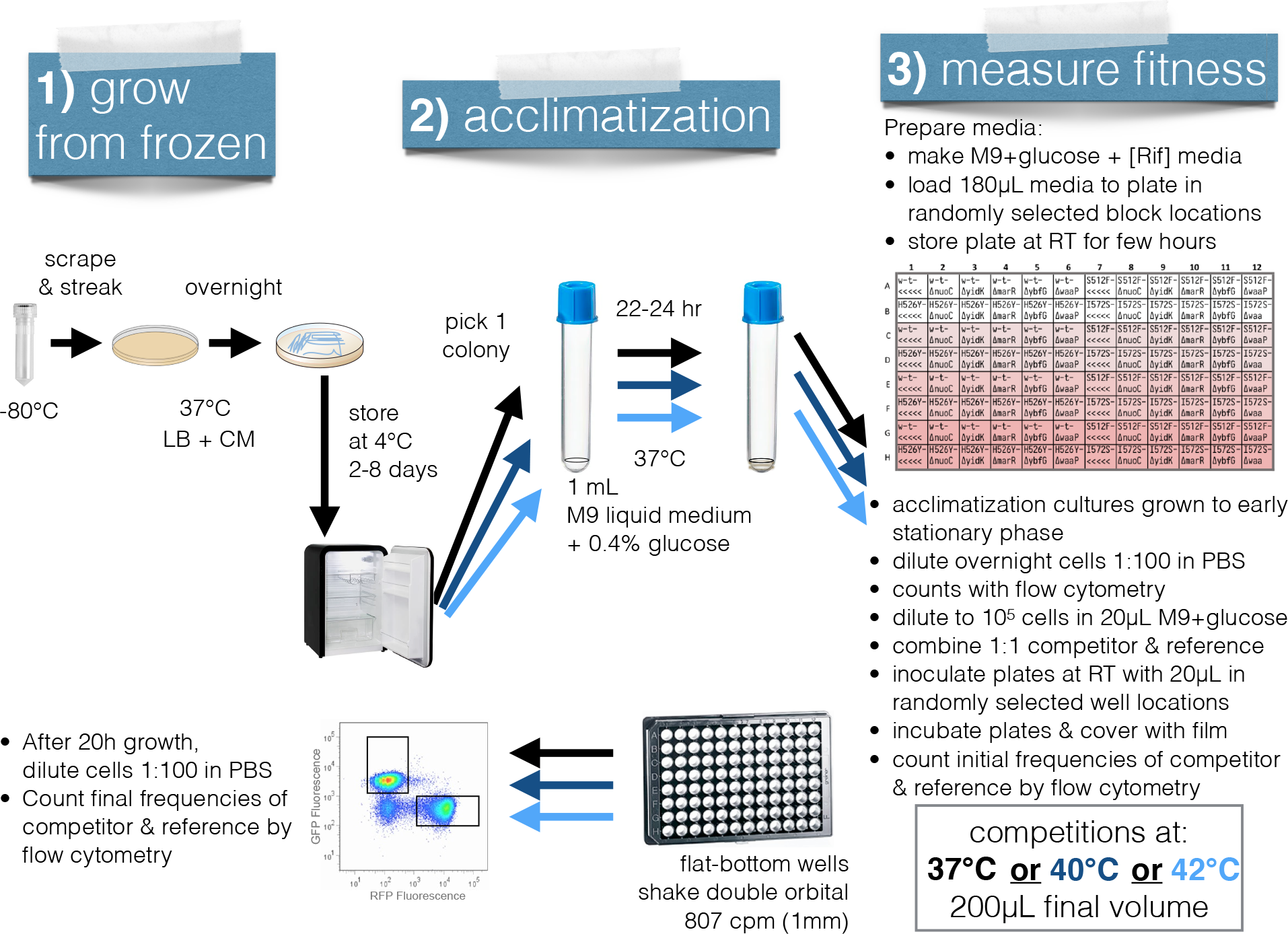
S**u**mmary **of the protocol used to estimate fitness in different antibiotic concentration and temperature environments. 1)** To ameliorate the possibility of *de novo* mutations, frozen glycerol stocks were bottle-necked to single colonies by streaking on agar plates. **2)** After streaking and growth on agar plates, all cells experienced the same temperature acclimitization of at least 40 hours and at most 8 days at 4*^◦^*C followed by batch culture growth at 37*^◦^*C for 22-24 hours. **3)** Batch culture competitions took place in 96-well plates with randomized positions of the four blocks of antibiotic concentrations and the 24 samples. Only one plate was incubated at a time, therefore triple arrows show steps in the protocol that had to be repeated for each temperature environment in order to sample a complete block of all samples in all environments. Regardless of the temperature environment where competition occurred, batch culture acclimitization always occurred at 37*^◦^*C.

#### 9.2.3 Estimating competitive index

We used R (4.1.1) as run using RStudio (2021.09.0+351) to calculate the competitive index using the equation indicated in the results. The GitLab repository for this manuscript, https://gitlab.com/evoldynamics/epistasis-decreases-with-increasing-antibiotic-pressure, contains the competitive index estimates and R code for calculating the competitive index from the cell counts data, calculating selection.Rmd.

### 9.3 Analyses

Parametric bootstrapping (10 000 bootstrap replicates) was used to determine the 95% confidence intervals on all of the summary statistics of epistasis. For the diminishing-returns epistasis, linear regressions were run on each of the 10 000 bootstrap replicates (x = fitness of the background, y = fitness effect of the mutation) and the mean slope value was retained in order to determine the 95% confidence intervals across all linear regressions.

We fitted a linear model in R using the equation *w*^ *∼ G_rpoB_* + *G*_KO_ + *E*_T_ + *E*_AB_ and the lm command from the stats package (version 4.1.1). The genotypic effects of *rpoB* (*G_rpoB_*) and knock-out (*G*_KO_) mutations were treated as categorical variables (in R: unordered factors), while the environmental effects of temperature (*E*_T_) and antibiotic concentration (*E*_AB_) were treated as semi-quantitative variables (in R: ordered factors) using polynomial contrasts.

All analyses are publicly available on GitLab at https://gitlab.com/evoldynamics/epistasis-decreases-with-increasing-antibiotic-pressure. The epistasis calculations can be found in estimating epistasis.Rmd, the linear model fitting can be found in ANOVA etal.Rmd, and Finlay-Wilkinson regression can be found in FinleyWilkinson.Rmd.

