## Supplementary material for "Epistasis decreases with increasing antibiotic pressure but not temperature": electronic supplement

### Electronic Supplementary Material for: Epistasis decreases with increasing antibiotic pressure but not temperature

**3 Supplementary References** **71**

### 1 Supplementary results

#### 1.1 Sample confirmation by whole genome re-sequencing

Table S1: Summary of Illumina sequencing quality for all 25 genotypes used in competitions. (Engineered mutations are listed in table S12.)

| Sample | Total reads | Percent ≧ Q30 bases | Mean quality score |
|---|---|---|---|
| AHG015 | 2 386 555 | 94.8% | 36.1 |
| AHG052 | 2 548 429 | 94.2% | 36.0 |
| AHG058 | 2 776 656 | 94.8% | 36.1 |
| AHG060 | 1 909 742 | 94.1% | 36.0 |
| AHG064 | 1 936 480 | 94.4% | 36.0 |
| AHG066 | 2 261 715 | 94.7% | 36.0 |
| AHG068 | 2 468 947 | 95.0% | 36.1 |
| AHG070 | 2 422 297 | 94.7% | 36.0 |
| AHG072 | 2 282 396 | 94.8% | 36.0 |
| AHG074 | 2 210 693 | 93.9% | 35.9 |
| AHG076 | 2 689 166 | 95.0% | 36.1 |
| AHG078 | 2 495 375 | 95.0% | 36.1 |
| AHG079 | 2 325 135 | 94.9% | 36.1 |
| AHG080 | 2 632 699 | 95.1% | 36.1 |
| AHG082 | 2 441 375 | 94.5% | 36.0 |
| AHG088 | 2 500 358 | 94.3% | 36.0 |
| AHG090 | 2 294 050 | 94.9% | 36.1 |
| AHG092 | 2 473 343 | 95.0% | 36.1 |
| AHG094 | 2 558 647 | 94.8% | 36.1 |
| AHG096 | 2 690 571 | 95.0% | 36.1 |
| AHG098 | 2 291 453 | 94.7% | 36.0 |
| AHG100 | 2 606 356 | 94.7% | 36.0 |
| AHG102 | 3 107 144 | 91.8% | 35.5 |
| AHG104 | 2 162 073 | 94.6% | 36.0 |
| AHG106 | 2 458 521 | 94.6% | 36.0 |

Table S2: Summary of breseq sequence mapping quality for all 25 genotypes used in competitions. Total reads used is less than total reads sequenced from table S1 since an average read depth limit of 120x was set, as recommended in the breseq manual. Mean read depth is the average across all reference sequences (*not* weighted by reference sequence length). Engineered mutations are listed in table S12.

| Sample | Total reads used | Mean read length | % reads mapped | Mean read depth |
|---|---|---|---|---|
| AHG015 | 3 700 125 | 150.6 bases | 97.8% | 117.2 |
| AHG052 | 3 702 686 | 150.6 bases | 97.6% | 120.0 |
| AHG058 | 3 702 707 | 150.6 bases | 98.2% | 119.7 |
| AHG060 | 3 702 714 | 150.6 bases | 98.2% | 111.7 |
| AHG064 | 3 702 704 | 150.6 bases | 97.9% | 116.1 |
| AHG066 | 3 702 744 | 150.6 bases | 98.1% | 113.9 |
| AHG068 | 3 702 740 | 150.6 bases | 98.3% | 117.2 |
| AHG070 | 3 702 725 | 150.6 bases | 98.1% | 116.2 |
| AHG072 | 3 702 732 | 150.6 bases | 97.8% | 115.3 |
| AHG074 | 3 702 768 | 150.6 bases | 97.5% | 115.7 |
| AHG076 | 3 702 698 | 150.6 bases | 97.8% | 116.9 |
| AHG078 | 3 702 667 | 150.6 bases | 97.9% | 113.6 |
| AHG079 | 3 704 544 | 150.6 bases | 98.1% | 122.5 |
| AHG080 | 3 702 716 | 150.6 bases | 98.1% | 116.4 |
| AHG082 | 3 702 707 | 150.6 bases | 97.9% | 119.4 |
| AHG088 | 3 702 678 | 150.6 bases | 97.8% | 113.9 |
| AHG090 | 3 702 737 | 150.6 bases | 97.6% | 116.5 |
| AHG092 | 3 702 711 | 150.6 bases | 98.0% | 114.6 |
| AHG094 | 3 702 620 | 150.6 bases | 94.8% | 115.5 |
| AHG096 | 3 702 724 | 150.6 bases | 97.9% | 116.0 |
| AHG098 | 3 702 699 | 150.6 bases | 97.3% | 112.5 |
| AHG100 | 3 702 690 | 150.6 bases | 97.8% | 121.3 |
| AHG102 | 3 702 620 | 150.6 bases | 97.0% | 117.2 |
| AHG104 | 3 702 751 | 150.6 bases | 98.2% | 117.0 |
| AHG106 | 3 702 704 | 150.6 bases | 97.9% | 112.4 |

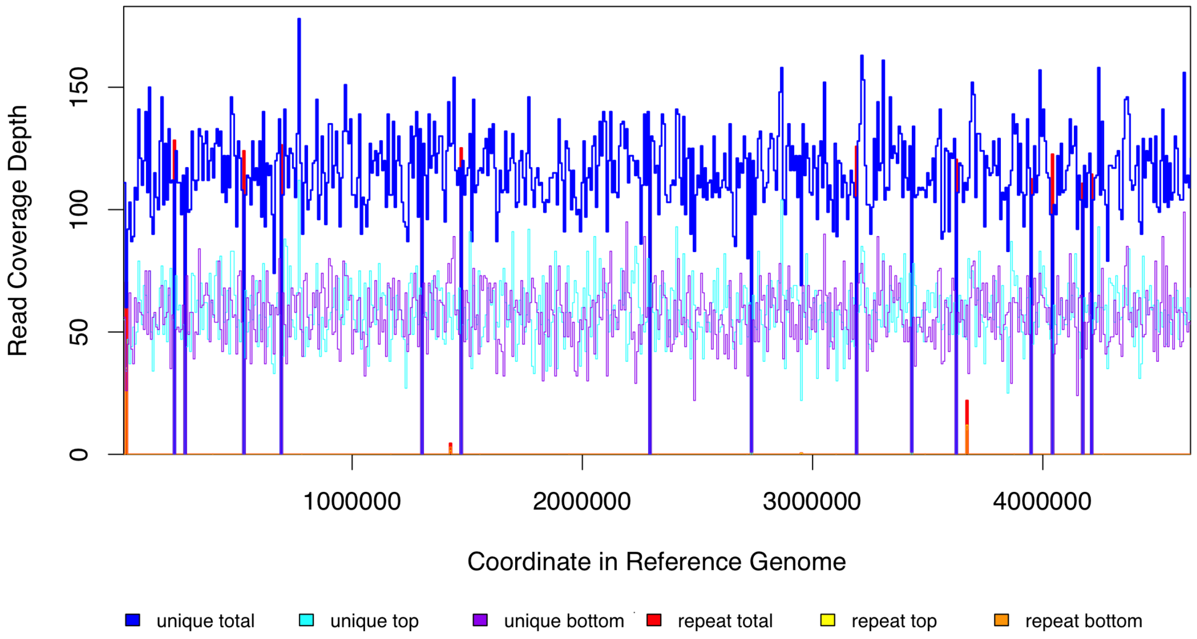

(a) Reference sequence: NC000913.

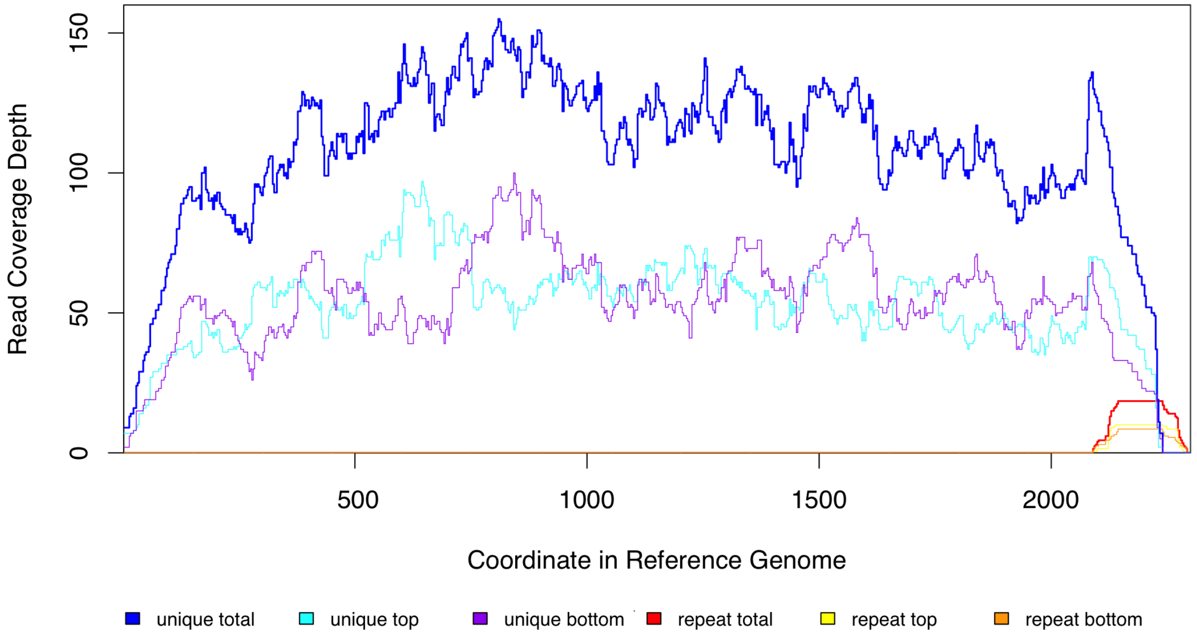

(b) Reference sequence: sfGFP.

Figure S1: Breseq read coverage plots of all reference genomes used for wild-type sample (AHG015). a) 'NC000913' is the MG1655 reference sequence from GenBank (NC_000913.3) and b) the sequence for the sfGFP construct is included in the supplementary methods.

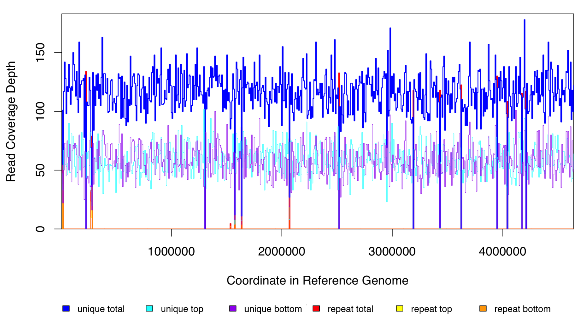

(a) Reference sequence: NC000913.

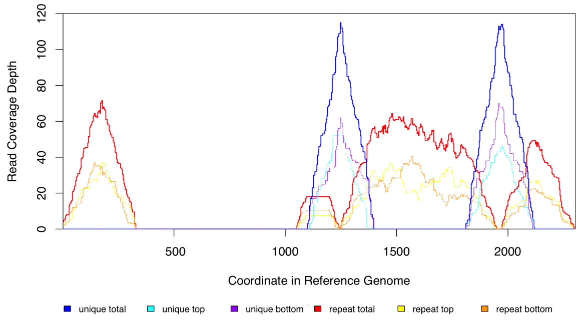

(b) Reference sequence: sfGFP.

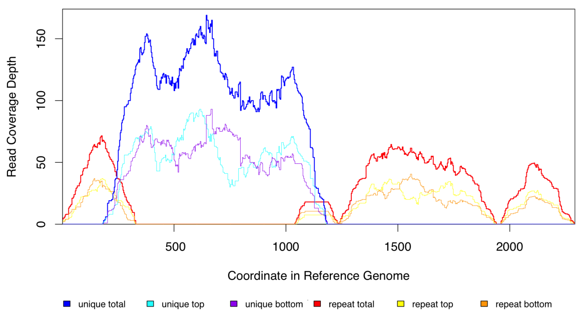

(c) Reference sequence: mCherry.

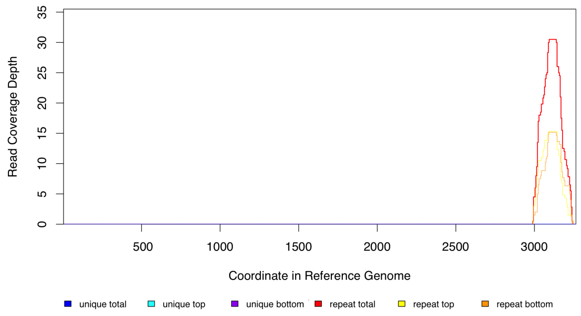

(d) Reference sequence: AY048743.

Figure S2: Breseq read coverage plots of all reference genomes used for the *rpoB* H526Y sample that was used as the common, mCherry-labeled reference in all competitions (AHG079). a) 'NC000913' is the MG1655 reference sequence from GenBank (NC_000913.3). b) There are no unique reads mapped to the sfGFP coding sequence on the construct because sample AHG079 has c) the mCherry construct instead (both reference sequences are included in the supplementary methods). d) AY048743 is the pDK4 plasmid sequence from GenBank (AY048743.1). Unique reads do not map to this sequence because there is no engineered gene knock-out (nor kanamyacin cassette) present.

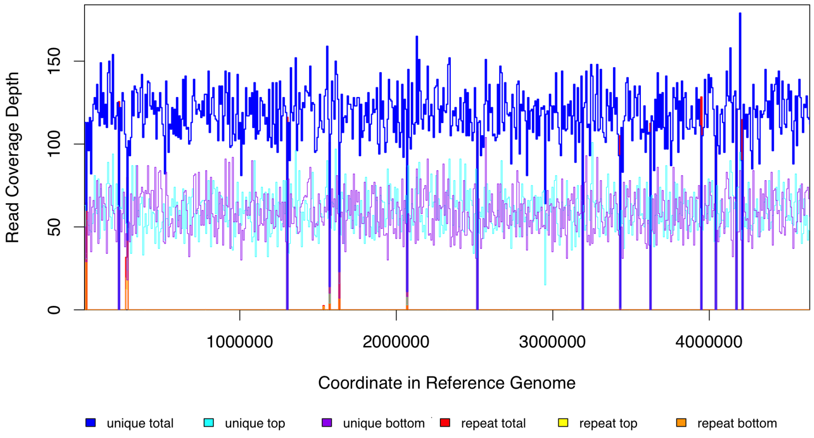

(a) Reference sequence: NC000913.

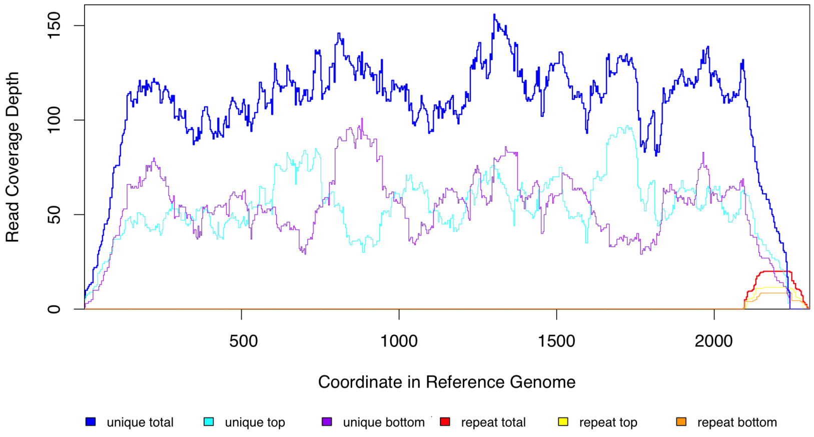

(b) Reference sequence: sfGFP.

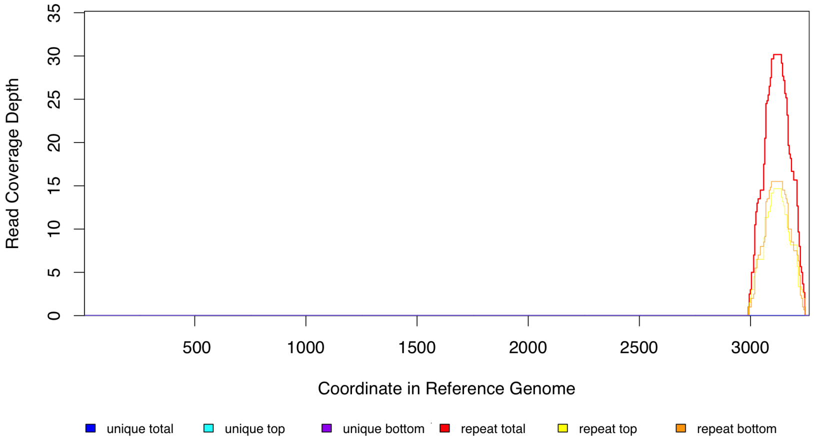

(c) Reference sequence: AY048743.

Figure S3: Breseq read coverage plots of all reference genomes used for sample AHG068. a) 'NC000913' is the MG1655 reference sequence from GenBank (NC_000913.3). b) The sequence for the sfGFP construct is included in the supplementary methods. c) AY048743 is the pDK4 plasmid sequence from GenBank (AY048743.1). Unique reads do not map to this sequence because there is no engineered gene knock-out (nor kanamyacin cassette) present.

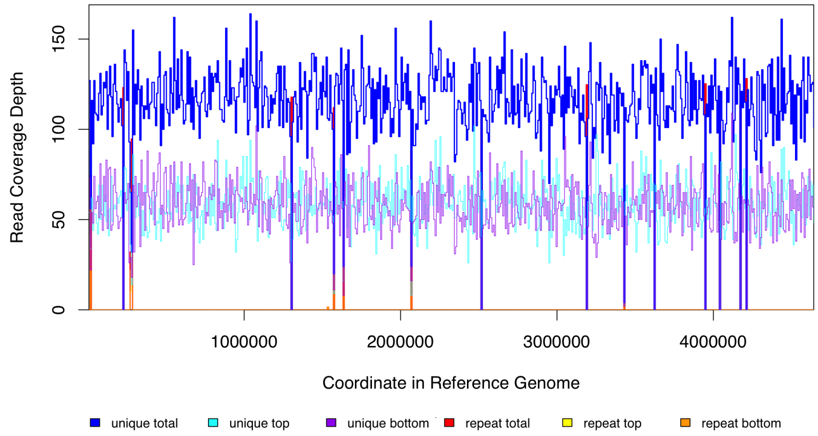

(a) Reference sequence: NC000913.

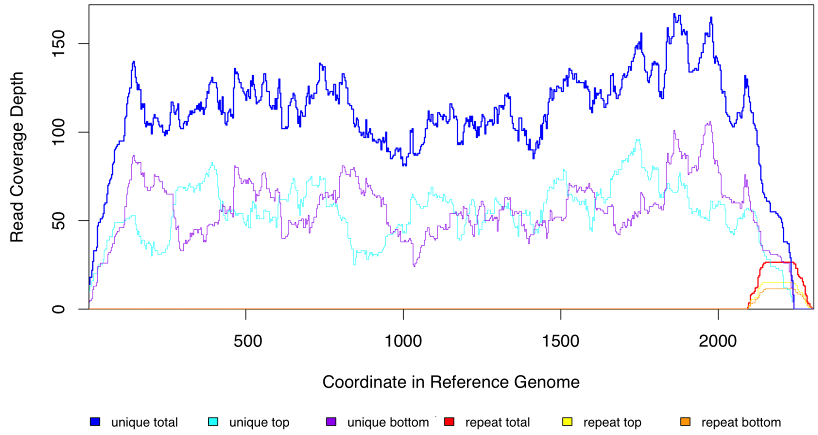

(b) Reference sequence: sfGFP.

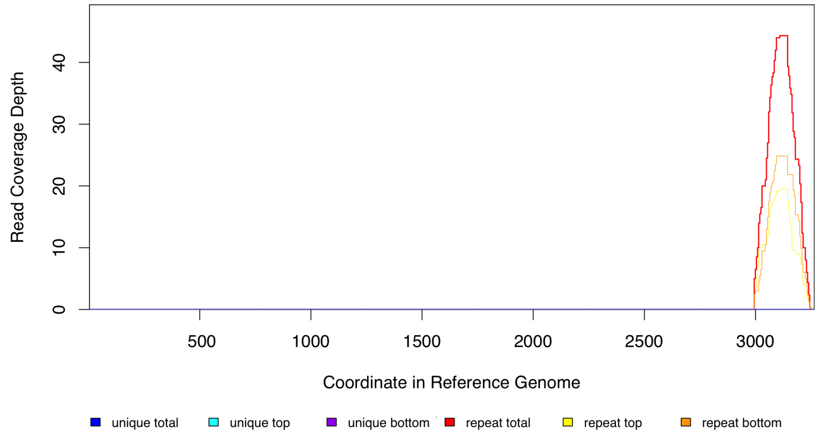

(c) Reference sequence: AY048743.

Figure S4: Breseq read coverage plots of all reference genomes used for sample AHG080. a) 'NC000913' is the MG1655 reference sequence from GenBank (NC_000913.3). b) The sequence for the sfGFP construct is included in the supplementary methods. c) AY048743 is the pDK4 plasmid sequence from GenBank (AY048743.1). Unique reads do not map to this sequence because there is no engineered gene knock-out (nor kanamyacin cassette) present.

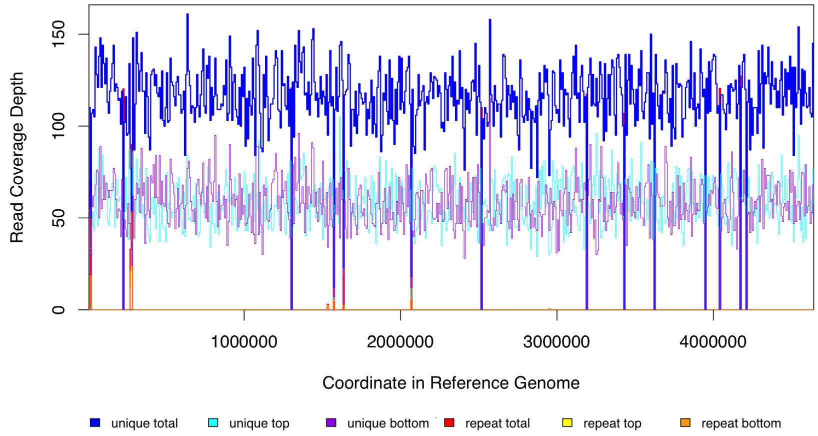

(a) Reference sequence: NC000913.

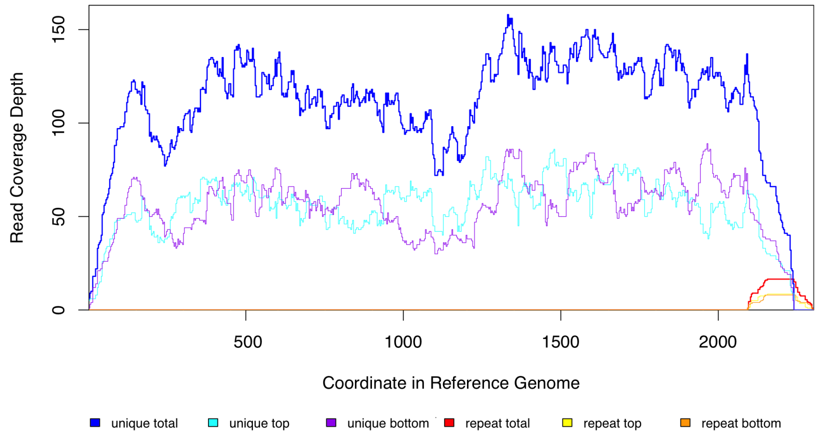

(b) Reference sequence: sfGFP.

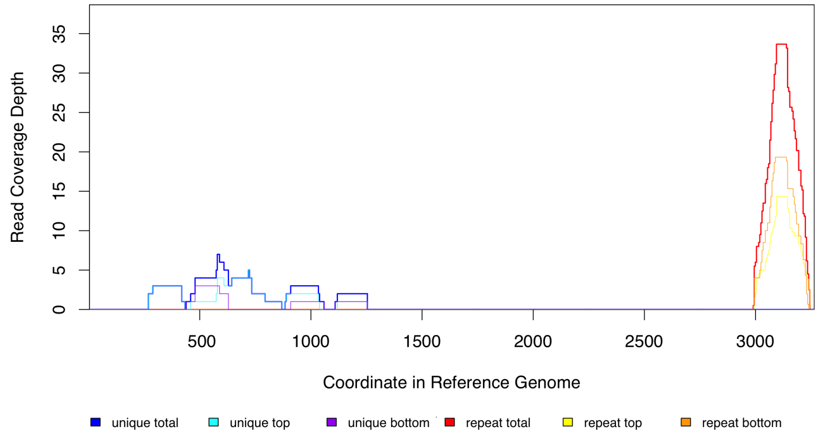

(c) Reference sequence: AY048743.

Figure S5: Breseq read coverage plots of all reference genomes used for sample AHG082. a) 'NC000913' is the MG1655 reference sequence from GenBank (NC_000913.3). b) The sequence for the sfGFP construct is included in the supplementary methods. c) AY048743 is the pDK4 plasmid sequence from GenBank (AY048743.1). Unique reads do not map to this sequence because there is no engineered gene knock-out (nor kanamyacin cassette) present.

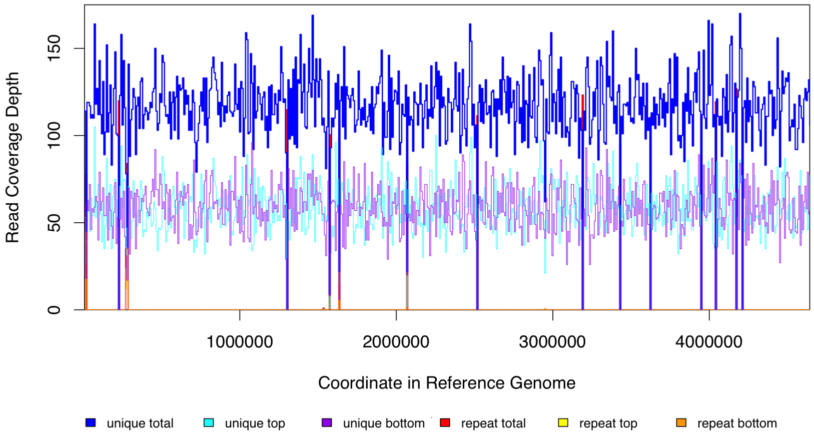

(a) Reference sequence: NC000913.

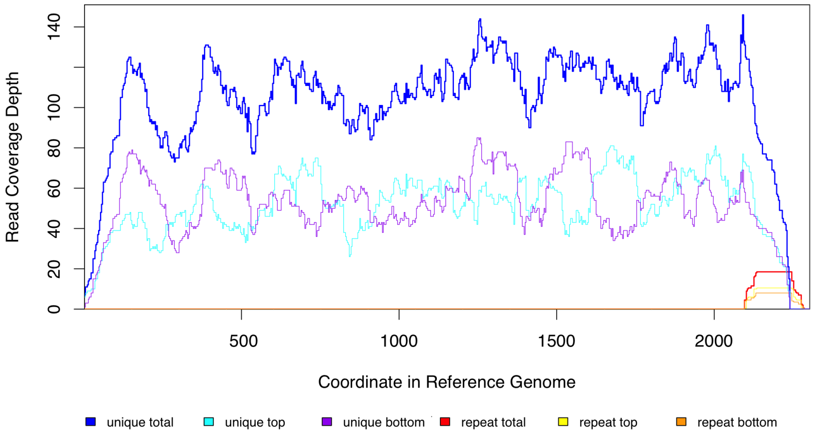

(b) Reference sequence: sfGFP.

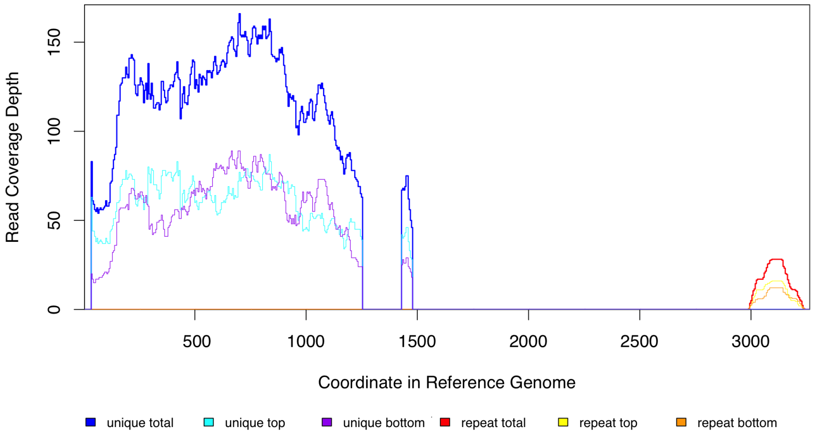

(c) Reference sequence: AY048743.

Figure S6: Breseq read coverage plots of all reference genomes used for sample AHG052. This sample is represenative of all coverage plots. a) 'NC000913' is the MG1655 reference sequence from GenBank (NC_000913.3). b) The sequence for the sfGFP construct is included in the supplementary methods. c) AY048743 is the pDK4 plasmid sequence from GenBank (AY048743.1). Reads map to this sequence because there is an engineered gene knock-out created by swapping the gene of interest with a kanamyacin cassette.

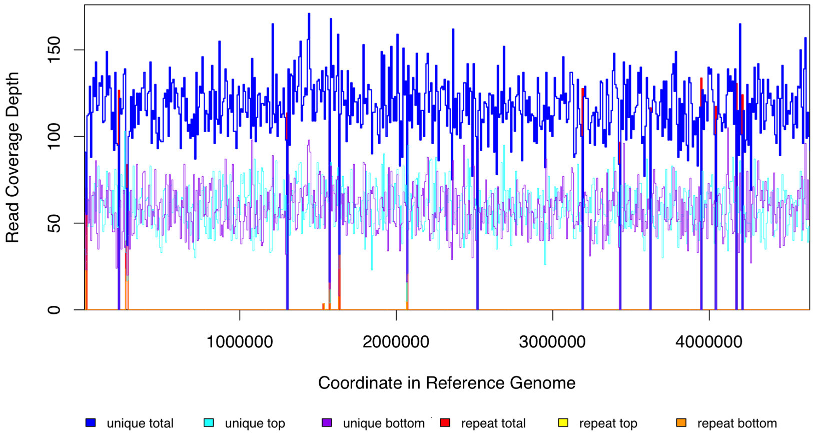

(a) Reference sequence: NC000913.

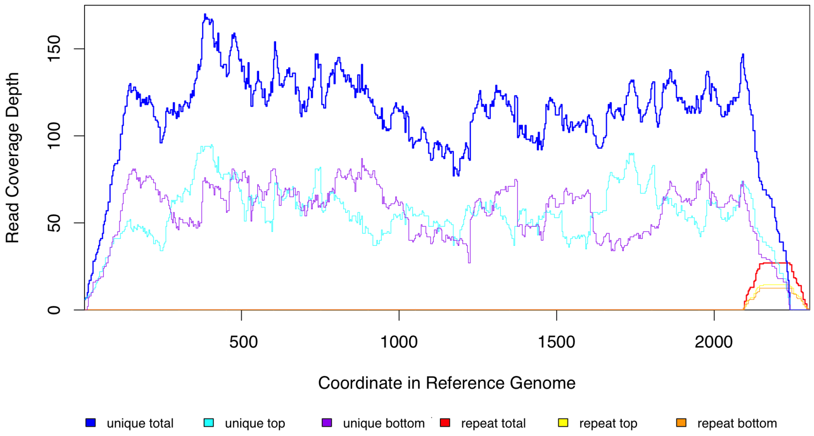

(b) Reference sequence: sfGFP.

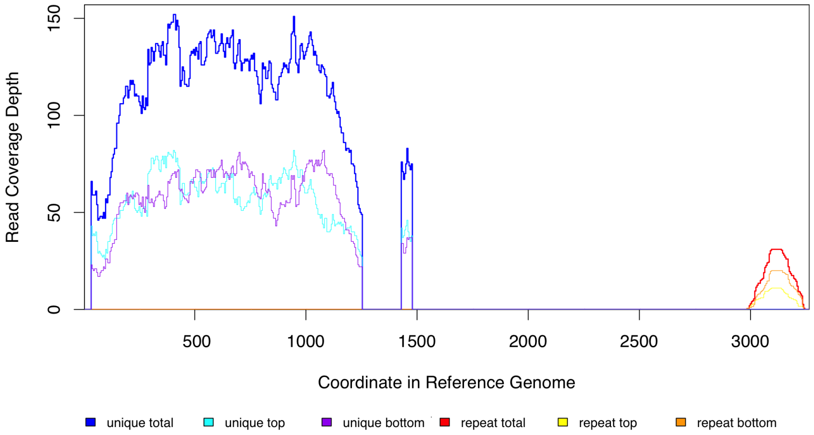

(c) Reference sequence: AY048743.

Figure S7: Breseq read coverage plots of all reference genomes used for sample AHG100. This sample is represenative of all coverage plots. a) 'NC000913' is the MG1655 reference sequence from GenBank (NC_000913.3). b) The sequence for the sfGFP construct is included in the supplementary methods. c) AY048743 is the pDK4 plasmid sequence from GenBank (AY048743.1). Reads map to this sequence because there is an engineered gene knock-out created by swapping the gene of interest with a kanamyacin cassette.

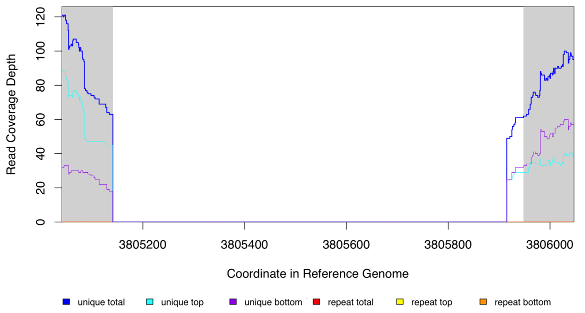

(a) Sample: AHG052.

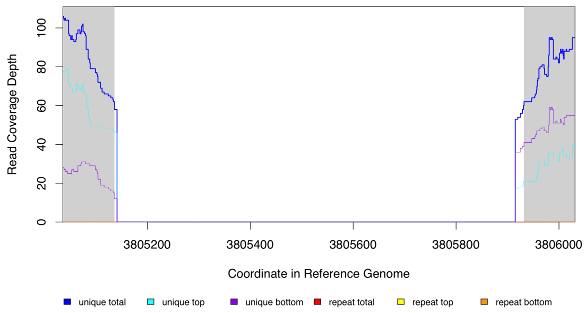

(b) Sample: AHG078.

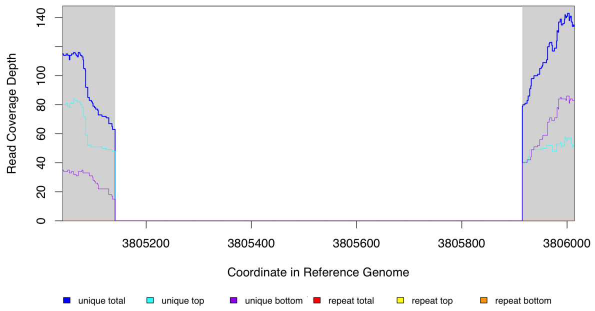

(c) Sample: AHG088.

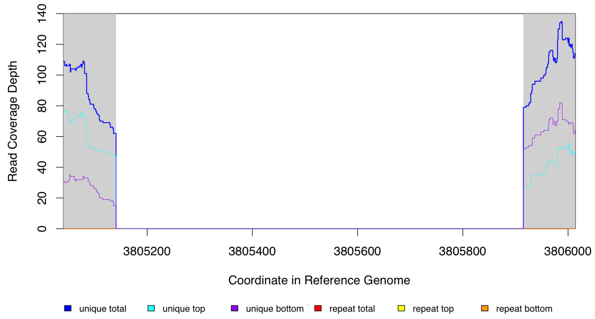

(d) Sample: AHG098.

Figure S8: Breseq read coverage plots for the region of the NC_000913.3 reference sequence around the gene *waaP* for the four samples engineered to have a knock-out at this gene: a) AHG052, b)AHG078, c) AHG088, and d) AHG098. The white background on the plot shows the region detected by breseq as having a reduced read coverage as compared to the regions with grey background.. No reads map to the *waaP* coding sequence because a knock-out has been engineered at this region by replacing the gene coding sequence with a kanamyacin cassette. Only samples with $\Delta waaP$ are included as representative since the coverage plots at the knock-out sites for the other genes and samples are similar.

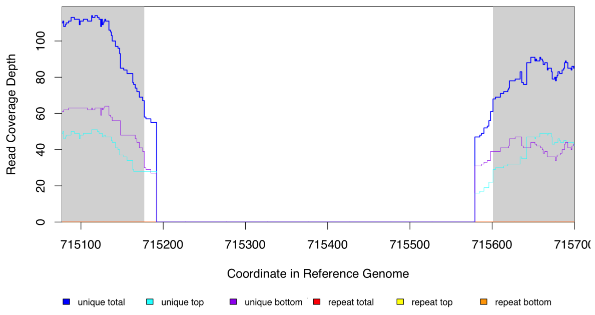

(a) Sample: AHG058.

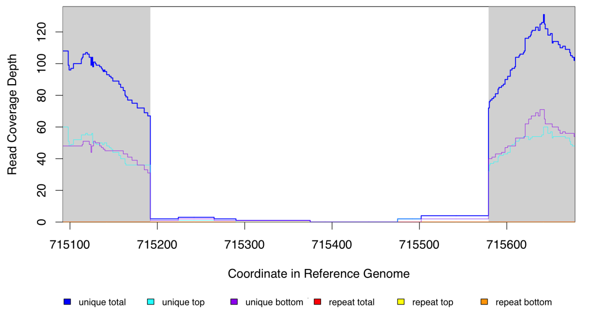

(b) Sample: AHG070.

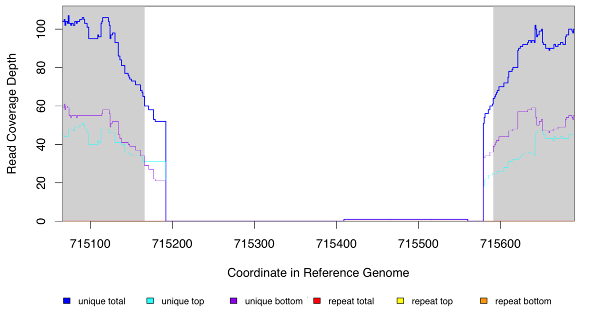

(c) Sample: AHG090.

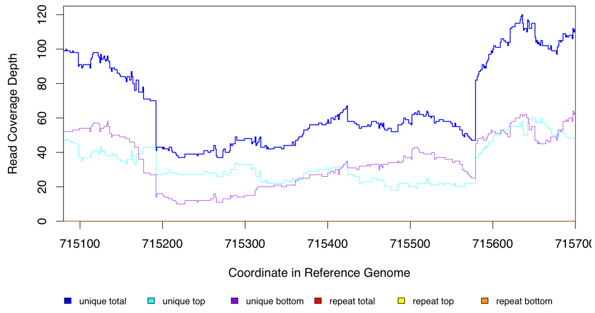

(d) Sample: AHG100.

Figure S9: Breseq read coverage plots for the region of the NC_000913.3 reference sequence around the pseudogene *ybfG* for the four samples engineered to have a knock-out at this locus: a) AHG058, b)AHG070, c) AHG090, and d) AHG100. In a-c), no reads map to the *ybfG* putative coding sequence because a knock-out has been engineered at this region by replacing the pseudogene sequence with a kanamyacin cassette. However, for d) there has likely been contamination by sample AHG098 ($\Delta$*waaP*, *rpoB* S512F) into sample AHG100 ($\Delta$*ybfG*, *rpoB* S512F) and so reads are mapping to the region around *ybfG* (see also read coverage plot in figure S10 and read junction evidence in figure S12). The white background on the plots for panels a-c) shows the region detected by breseq as having a reduced read coverage as compared to the regions with grey background. Panel d) was created manually at the *ybfG* locus using the breseq BAM2COV command and so there is no background shading to the plot.

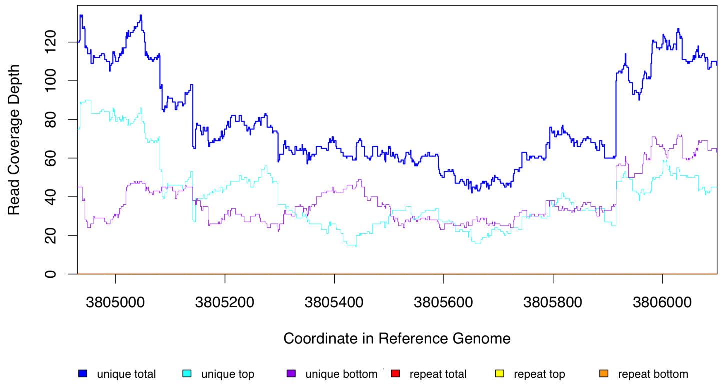

Figure S10: Breseq read coverage plots for the region of the NC_000913.3 reference sequence around the gene *waaP* (3805140 - 3805915) for the AHG100 samples that is **not** engineered to have a knock-out at this site. There has likely been contamination by sample AHG098 ($\Delta$*waaP*, *rpoB* S512F) into sample AHG100 ($\Delta$*ybfG*, *rpoB* S512F) and so there is decreased read coverage at the *waaP* locus as a result (see also read coverage plot in figure S9d and read junction evidence in figure S12).

New junction evidence

| | | seq id | position | reads (cov) | reads (cov) | score | skew | freq | annotation | gene | product |
|---|---|---|---|---|---|---|---|---|---|---|---|
| * | ? | AY048743 | 32 = | 0 (0.000) | 63 (0.550) | 20/298 | 0.2 | 100% | intergenic (–/–427) | *–/kanamycin resistance* | –/Tn5 neomycin phosphotransferase |
| | ? | NC_000913 | = 3805140 | 0 (0.000) | | | | | coding (778/798 nt) | *waaP* | lipopolysaccharide core heptose (I) kinase |

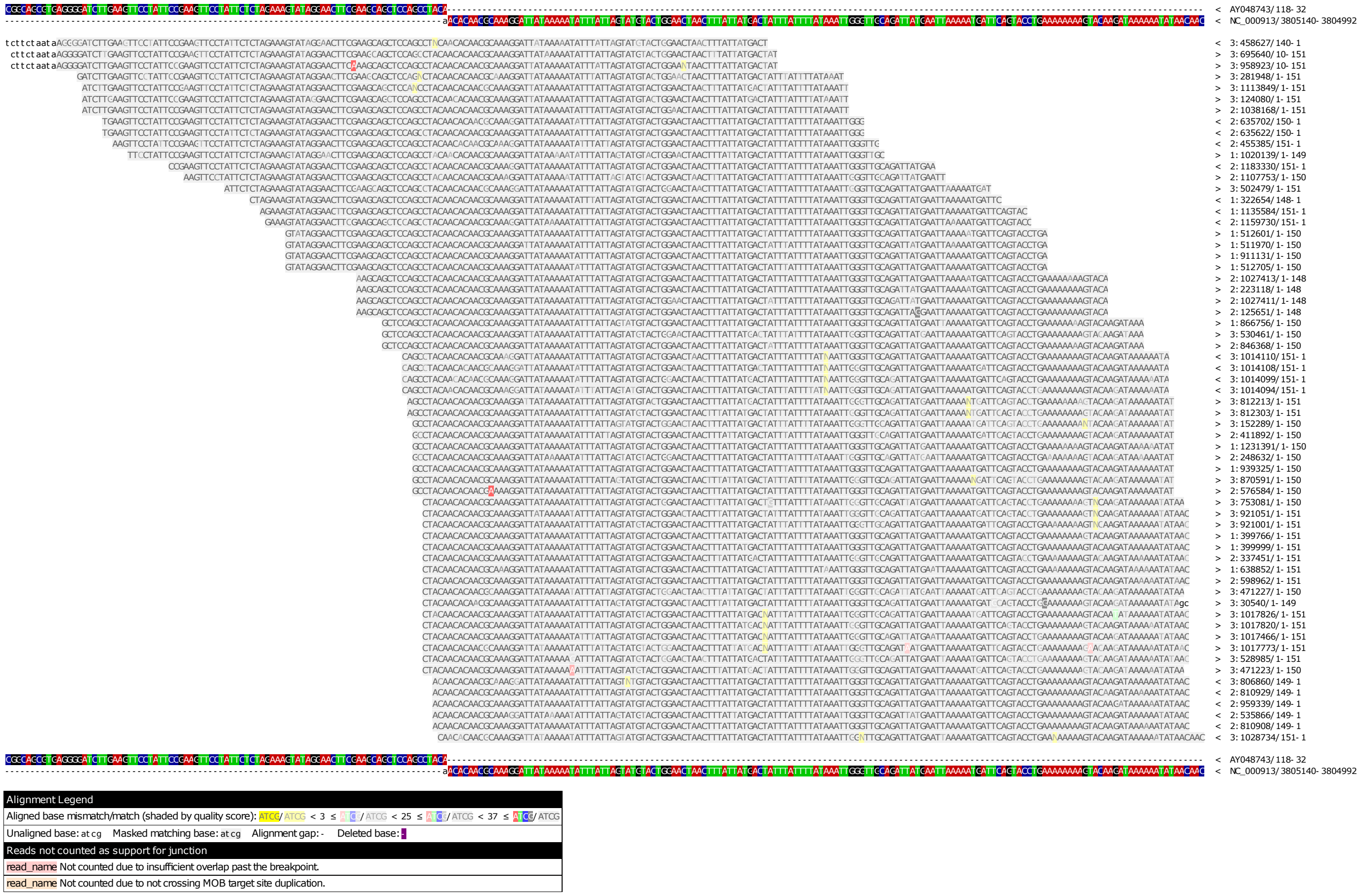

(a) New junction evidence upstream of the *waaP* gene for sample AHG052. (Figure continues on next page.)

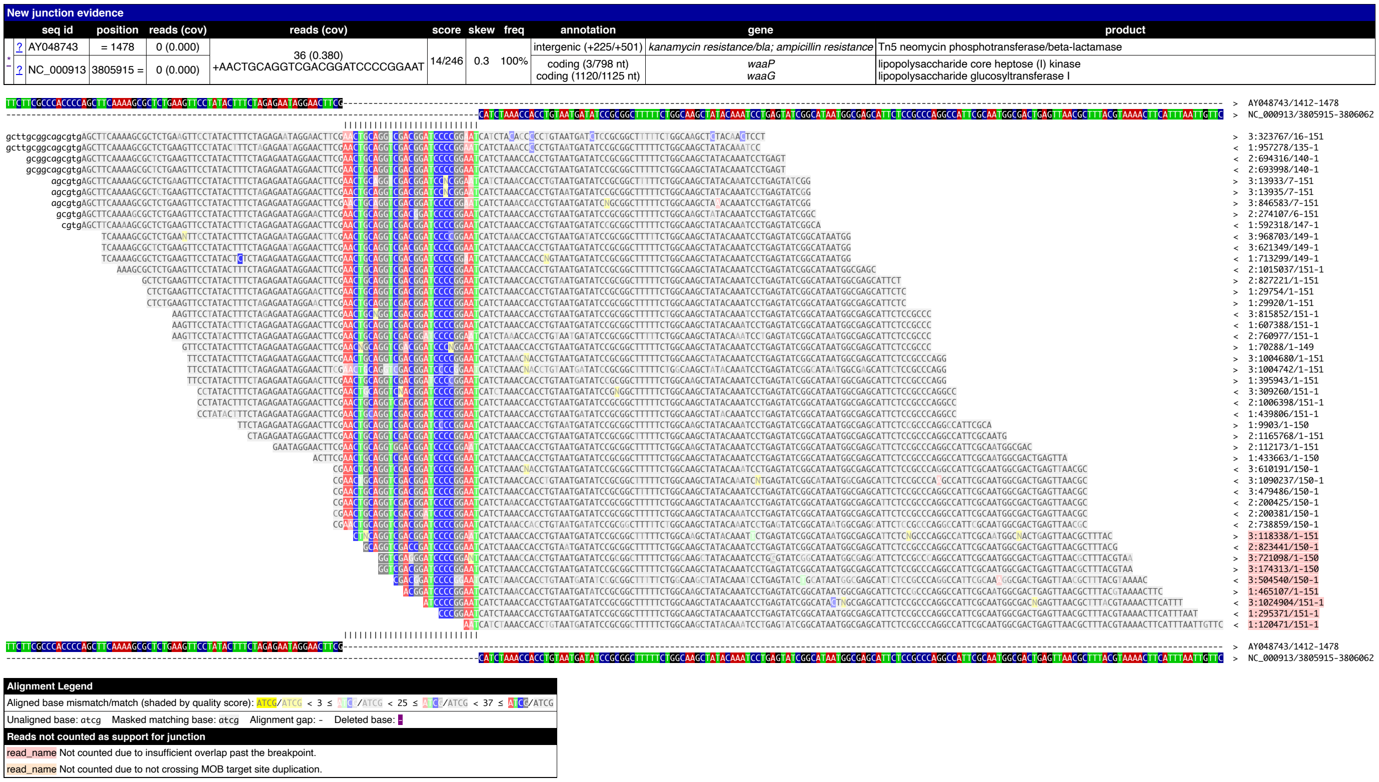

(b) New junction evidence downstream of the *waaP* gene for sample AHG052. (Figure continued from previous page.)

Figure S11: Breseq new junction evidence supporting the insertion of the kanamyacin cassette from the pKD4 plasmid (AY048743.1) at the region of the NC_000913.3 reference sequence around the *waaP* gene for sample AHG052. Panel a), which is on the previous page, shows evidence in support of a new junction at position 3 805 150 of the NC_000913.3 reference sequence (upstream of the *waaP* gene). Panel b) shows evidence of a second junction around position 3 805 915 (downstream of the *waaP* gene). These positions correspond well with the lack of read coverage for this sample at this region shown in figure S8a. Only the new junction evidence from AHG052 is shown as a representative since new junction evidence for other samples and knocked-out genes look similar. All plots are from Breseq.

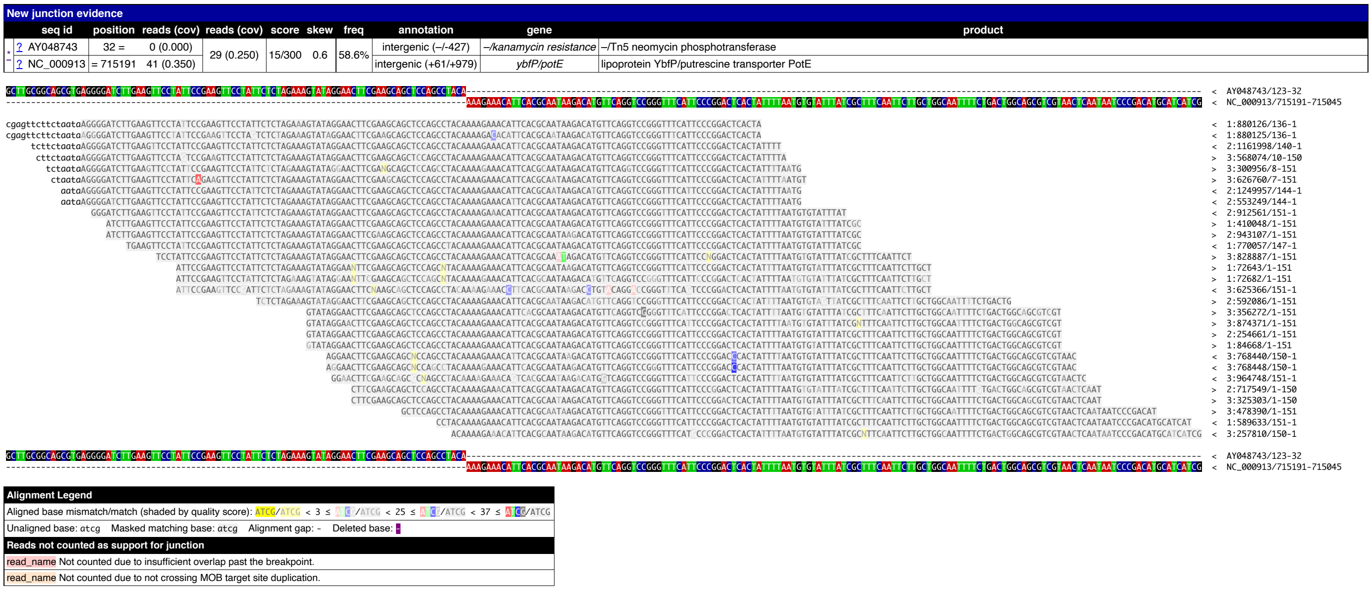

(a) New junction evidence upstream of the *ybfG* pseudogene for sample AHG100. (Figure continues on next page.)

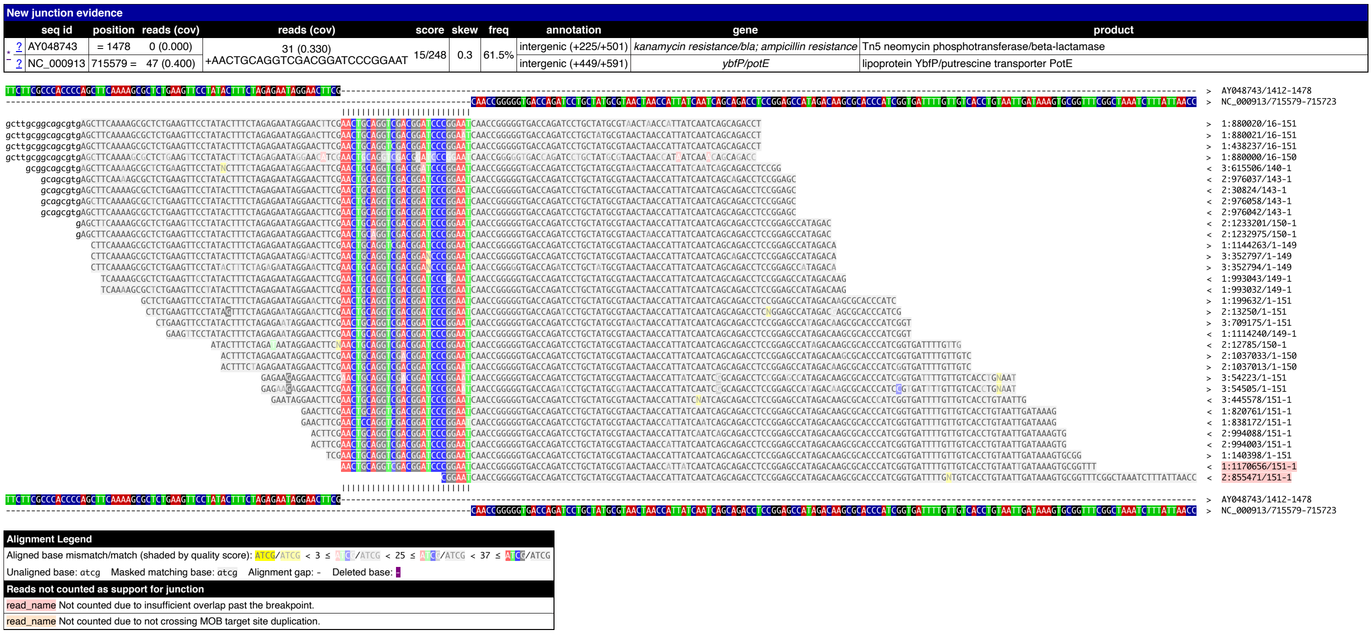

(b) New junction evidence downstream of the *ybfG* pseudogene for sample AHG100. (Figure continued from previous page and continues on next page.)

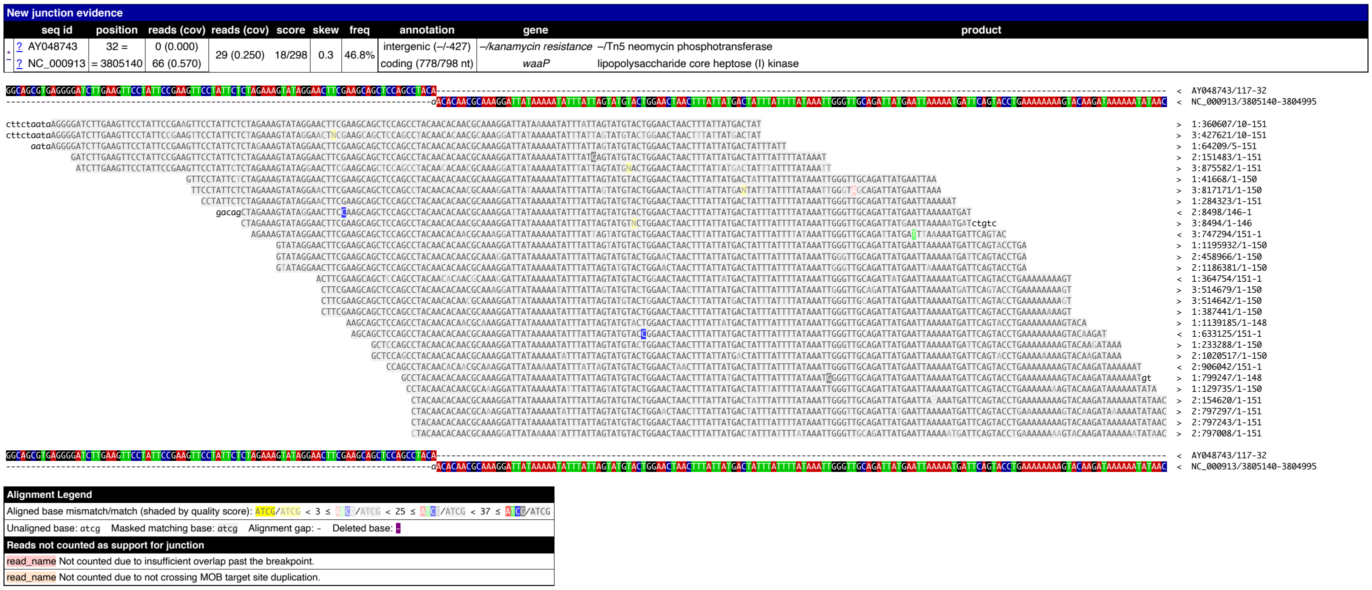

(c) New junction evidence upstream of the *waaP* gene for sample AHG100. (Figure continued from previous page and continues on next page.)

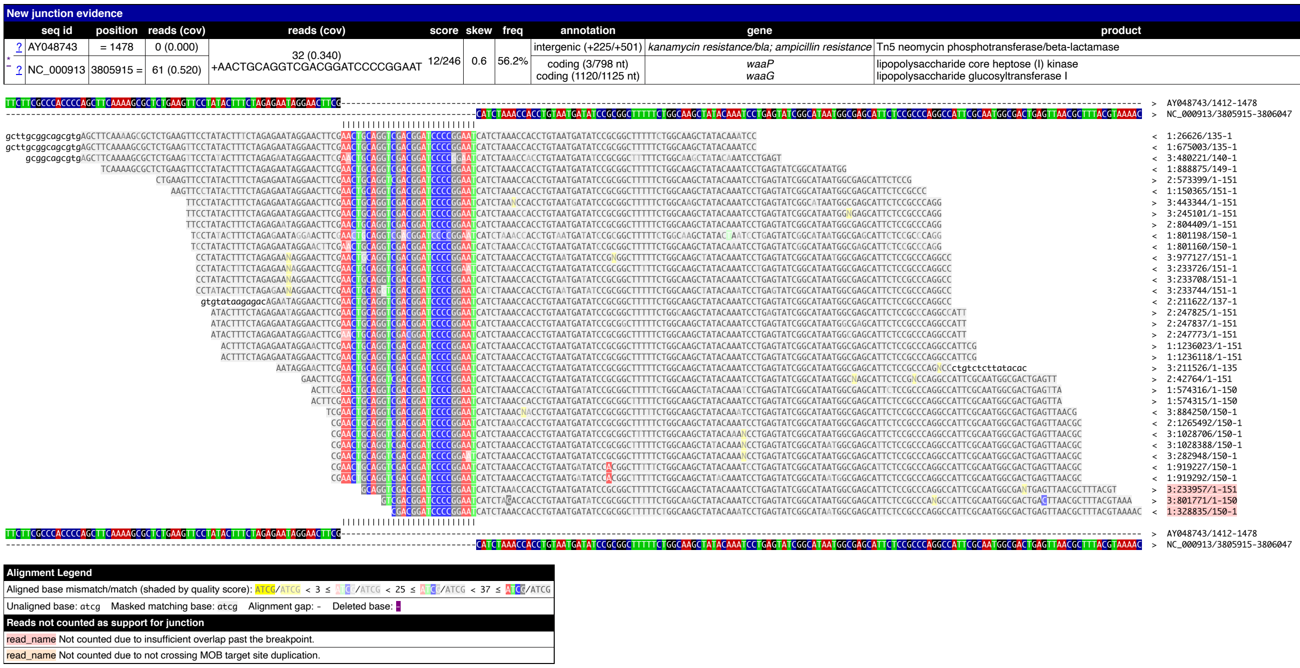

(d) New junction evidence downstream of the *waaP* gene for sample AHG100. (Figure continued from previous page.)

Figure S12: Breseq new junction evidence supporting the hypothesis of likely contamination by sample AHG098 (Δ*waaP*, *rpoB* S512F) into sample AHG100 (Δ*ybfG*, *rpoB* S512F); see also read coverage plots in figures S9d and S10. The new junction evidence for sample AHG100 shows existence of insertion of the kanamyacin cassette from the pKD4 plasmid (AY048743.1) at two different loci of the NC_000913.3 reference sequence. Panels a-b) show the new junctions around the *ybfG* pseudogene and panels c-d) show the new junctions around the *waaP* gene. All plots are from Breseq.

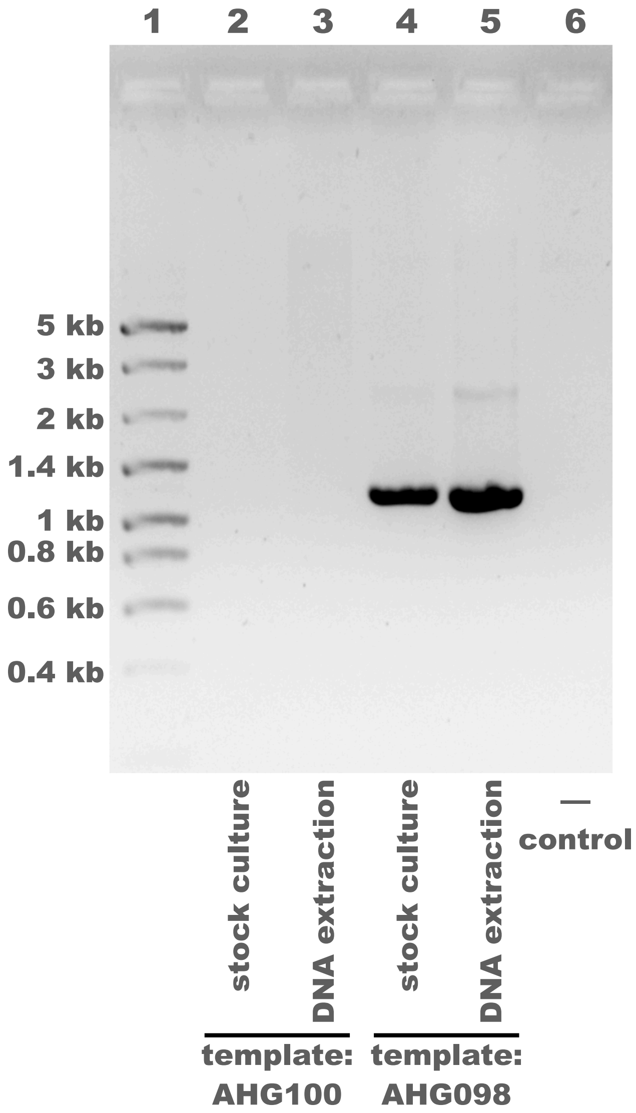

Figure S13: **PCR suggests that contamination of sample AHG100 (*rpoB* S512F $\Delta ybfG$) occurred after creation of the stock culture and after DNA extraction, perhaps during DNA sequencing.** PCR was done using a $\Delta waaP$-specific primer (waaP_76747_For in table S13) and the Datsenko-Wanner K1 primer specific to the inserted kanamyacin cassette. Therefore amplification is expected only when the $\Delta waaP$ knock-out DNA sequence is present in the PCR template. The test sample, AHG100: *rpoB* S512F $\Delta ybfG$, was used as a PCR template in lanes 2-3; lanes 4-5 show the positive control with AHG098: *rpoB* S512F $\Delta waaP$ and lane 6 shows the negative control without any PCR template. No amplification is observed for the test sample AHG100, this suggests that the sample was not contaminated prior to being sent for DNA sequencing. The frozen stock cultures (lanes 2, 4) and the DNA extractions that were sent for sequencing (lanes 3, 5) were used as PCR templates for both samples. For the stock cultures, 1mL of LB was inoculated directly with frozen stock, incubated for 1.5 hours at $37°C$, then $1\mu L$ was added to the PCR reaction. A $25\mu L$ total PCR reaction volume was used with DreamTaq DNA polymerase, annealing temperature of $55°C$, and 2 minute elongation time. Gel electrophoresis was done in 1% agarose. Lane 1 shows the NZYDNA Ladder VIII, with band sizes labeled at left.

Table S3: Complete list of mutations as identified by breseq for each of the 25 genotypes. (Engineered mutations listed in table S12.) Rows coloured in gray highlight mutations identified by breseq that correspond with the genetic constructs created for this study. Rows coloured in blue highlight non-synonymous nucleotide substitutions that were not engineered and differ from the wild-type strain (AHG015). All other types of mutations (synonymous nucleotide substitutions, intergenic nucleotide substitutions, small indels, and mobile element insertions) that were not engineered and mutations present on the wild-type strain are listed on white background. Sample AHG079 is labeled with "(Ref.)" to highlight that this was the reference genotype used in competitions.

| Sample | Mutation | Gene name | Gene product | Annotation | Reference | Position |
|---|---|---|---|---|---|---|
| AHG015 (WT) | large deletion | insB9, insA9, [crl] | insB9, insA9, [crl] | Δ776 bp | MG1655 (NC_000913.3) | 257908-258676 |
| AHG015 (WT) | small indel | ybjO | putative inner membrane protein | +33 bp coding (413-445/489 nt) | MG1655 (NC_000913.3) | 898401 |
| AHG015 (WT) | intergenic mobile element insertion | bamD/raiA | outer membrane protein assembly factor BamD/ribosome-associated inhibitor A | IS1 (+) +8 bp | MG1655 (NC_000913.3) | 2737118-2737125 |
| AHG015 (WT) | small indel | rrlA | 23S ribosomal RNA | Δ1 bp noncoding (207/2905 nt) | MG1655 (NC_000913.3) | 4037725 |
| AHG015 (WT) | small indel | gltP/yjcO | glutamate / aspartate : H(+) symporter GltP/Sel1 repeat-containing protein YjcO | +GC intergenic (+587/+55) | MG1655 (NC_000913.3) | 4296381 |
| AHG015 (WT) | SNP intergenic | sfGFP/cat | Superfolder GFP fluorophore / type A chloramphenicol O-acetyltransferase | T→C intergenic (+204/-99) | sfGFP construct (sequence) | 1248 |
| AHG015 (WT) | small indel | cat | type A chloramphenicol O-acetyltransferase | +A coding (609/639 nt) | sfGFP construct (sequence) | 1955 |
| AHG015 (WT) | small indel | cat | type A chloramphenicol O-acetyltransferase | +T coding (621/639 nt) | sfGFP construct (sequence) | 1967 |
| AHG058 | SNP synonymous | nptII | Tn5 neomycin phosphotransferase | S104S (A311G) | pKD4 plasmid (AY048743.1) | 770 |

Continued on next page

**Table S3 – continued from previous page**

| Sample | Mutation | Gene name | Gene product | Annotation | Reference | Position |
|---|---|---|---|---|---|---|
| AHG058 | gene knock-out | ybfP/potE | lipoprotein YbfP/putrescine transporter PotE | Missing coverage (figure S9a) & new junction evidence | MG1655 (NC_000913.3) | 715177-715600 |
| AHG060 | gene knock-out | marR | DNA-binding transcriptional repressor MarR | Missing coverage & new junction evidence | MG1655 (NC_000913.3) | 1618307-1618725 |
| AHG064 | mobile element insertion | yhhZ | putative endonuclease YhhZ | IS1 (+) +9 bp coding (666-674/1179 nt) | MG1655 (NC_000913.3) | 3582495-3582503 |
| AHG064 | gene knock-out | yidK | putative transporter YidK | Missing coverage & new junction evidence | MG1655 (NC_000913.3) | 3858389-3860101 |
| AHG066 | SNP synonymous | nptII | Tn5 neomycin phosphotransferase | V83V (T248C) | pKD4 plasmid (AY048743.1) | 707 |
| AHG066 | gene knock-out | nuoC | NADH:quinone oxidoreductase subunit CD | Missing coverage & new junction evidence | MG1655 (NC_000913.3) | 2401267-2403045 |
| AHG066 | SNP synonymous | kgtP | $\alpha$ keto-glutarate: H(+) symporter | V350V (C1049A) | MG1655 (NC_000913.3) | 2723888 |
| AHG052 | gene knock-out | waaP/waaG | lipopolysaccharide core heptose (I) kinase / lipopolysaccharide glucosyltransferase I | Missing coverage (figure S8a) & new junction evidence (figure S11) | MG1655 (NC_000913.3) | 3805141-3805947 |
| AHG052 | SNP intergenic | –/nptII | –/Tn5 neomycin phosphotransferase | C→T intergenic (–/-223) | pKD4 plasmid (AY048743.1) | 236 |
| AHG052 | SNP intergenic | –/nptII | –/Tn5 neomycin phosphotransferase | A→T intergenic (–/-88) | pKD4 plasmid (AY048743.1) | 371 |
| AHG079 (Ref.) | large deletion | ntpII | kanamycin resistance,bla; | $\Delta$3267 bp (no reads mapped to pKD4 plasmid, figure S2d) | pKD4 plasmid (AY048743.1) | 1-3267 |
| AHG079 (Ref.) | small indel | ybjO | putative inner membrane protein | +33 bp pseudogene (412/456 nt) | MG1655 (NC_000913.3) | 897624 |

Continued on next page

**Table S3 – continued from previous page**

| Sample | Mutation | Gene name | Gene product | Annotation | Reference | Position |
|---|---|---|---|---|---|---|
| AHG079 (Ref.) | SNP non-synonymous | *rpoB* | RNA polymerase subunit beta | H526Y (C1575T) | MG1655 (NC_000913.3) | 4182786 |
| AHG079 (Ref.) | large deletion | sfGFP | Superfolder GFP fluorophore | Missing coverage evidence, figure S2b | sfGFP construct (sequence) | 1-1227 |
| AHG079 (Ref.) | gene insertion | mCherry | mCherry fluorophore | Reads mapped to mCherry CDS, figure S2c | mCherry construct (sequence) | 325-1035 |
| AHG080 | large deletion | ntpII | kanamycin resistance,bla; | $\Delta$3267 bp (no reads mapped to pKD4 plasmid, figure S4c) | pKD4 plasmid (AY048743.1) | 1-3267 |
| AHG080 | SNP non-synonymous | *rpoB* | RNA polymerase subunit beta | H526Y (C1575T) | MG1655 (NC_000913.3) | 4182786 |
| AHG090 | SNP synonymous | nptII | Tn5 neomycin phosphotransferase | S104S (A311G) | pKD4 plasmid (AY048743.1) | 770 |
| AHG090 | gene knock-out | ybfP/potE | lipoprotein YbfP/putrescine transporter PotE | Missing coverage (figure S9c) & new junction evidence | MG1655 (NC_000913.3) | 715166-715590 |
| AHG090 | SNP non-synonymous | *rpoB* | RNA polymerase subunit beta | H526Y (C1575T) | MG1655 (NC_000913.3) | 4182786 |
| AHG092 | gene knock-out | marR | DNA-binding transcriptional repressor MarR | Missing coverage & new junction evidence | MG1655 (NC_000913.3) | 1618315-1618725 |
| AHG092 | SNP non-synonymous | *rpoB* | RNA polymerase subunit beta | H526Y (C1575T) | MG1655 (NC_000913.3) | 4182786 |
| AHG094 | gene knock-out | yidK | putative transporter YidK | Missing coverage & new junction evidence | MG1655 (NC_000913.3) | 3858389-3860089 |
| AHG094 | SNP non-synonymous | *rpoB* | RNA polymerase subunit beta | H526Y (C1575T) | MG1655 (NC_000913.3) | 4182786 |
| AHG096 | SNP synonymous | nptII | Tn5 neomycin phosphotransferase | V83V (T248C) | pKD4 plasmid (AY048743.1) | 707 |

Continued on next page

**Table S3 – continued from previous page**

| Sample | Mutation | Gene name | Gene product | Annotation | Reference | Position |
|---|---|---|---|---|---|---|
| AHG096 | gene knock-out | nuoC | NADH:quinone oxidoreductase subunit CD | Missing coverage & new junction evidence | MG1655 (NC_000913.3) | 2401261-2403045 |
| AHG096 | SNP non-synonymous | *rpoB* | RNA polymerase subunit beta | H526Y (C1575T) | MG1655 (NC_000913.3) | 4182786 |
| AHG088 | SNP intergenic | –/nptII | –/Tn5 neomycin phosphotransferase | C→T intergenic (–/-223) | pKD4 plasmid (AY048743.1) | 236 |
| AHG088 | SNP intergenic | –/nptII | –/Tn5 neomycin phosphotransferase | A→T intergenic (–/-88) | pKD4 plasmid (AY048743.1) | 371 |
| AHG088 | gene knock-out | waaP | lipopolysaccharide core heptose (I) kinase | Missing coverage (figure S8c) & new junction evidence | MG1655 (NC_000913.3) | 3805141-3805914 |
| AHG088 | SNP non-synonymous | *rpoB* | RNA polymerase subunit beta | H526Y (C1575T) | MG1655 (NC_000913.3) | 4182786 |
| AHG082 | large deletion | ntpII | kanamycin resistance,bla; | Δ3267 bp (no reads mapped to pKD4 plasmid, figure S5c) | pKD4 plasmid (AY048743.1) | 1-3267 |
| AHG082 | SNP non-synonymous | yeaC | DUF1315 domain-containing protein YeaC | A32G (G94C) | MG1655 (NC_000913.3) | 1861071 |
| AHG082 | SNP synonymous | zinT | metal-binding protein ZinT | S23S (G68A) | MG1655 (NC_000913.3) | 2040634 |
| AHG082 | SNP intergenic | insH6/yoeA | CP4-44 prophage; IS5 transposase and trans-activator/CP4-44 prophage; TonB-dependent receptor plug domain-containing protein YoeA | T→C intergenic (-1316/-34) | MG1655 (NC_000913.3) | 2067792 |
| AHG082 | SNP non-synonymous | *rpoB* | RNA polymerase subunit beta | S512F (C1534T) | MG1655 (NC_000913.3) | 4182745 |
| AHG100 | SNP synonymous | zinT | metal-binding protein ZinT | S23S (G68A) | MG1655 (NC_000913.3) | 2040634 |

Continued on next page

**Table S3 – continued from previous page**

| Sample | Mutation | Gene name | Gene product | Annotation | Reference | Position |
|---|---|---|---|---|---|---|
| AHG100 | SNP intergenic | insH6/yoeA | CP4-44 prophage; IS5 transposase and trans-activator/CP4-44 prophage; TonB-dependent receptor plug domain-containing protein YoeA | T→C intergenic (-1316/-34) | MG1655 (NC_000913.3) | 2067792 |
| AHG100 | SNP non-synonymous | *rpoB* | RNA polymerase subunit beta | S512F (C1534T) | MG1655 (NC_000913.3) | 4182745 |
| AHG102 | gene knock-out | marR | DNA-binding transcriptional repressor MarR | Missing coverage & new junction evidence | MG1655 (NC_000913.3) | 1618308-1618725 |
| AHG102 | SNP synonymous | zinT | metal-binding protein ZinT | S23S (G68A) | MG1655 (NC_000913.3) | 2040634 |
| AHG102 | SNP intergenic | insH6/yoeA | CP4-44 prophage; IS5 transposase and trans-activator/CP4-44 prophage; TonB-dependent receptor plug domain-containing protein YoeA | T→C intergenic (-1316/-34) | MG1655 (NC_000913.3) | 2067792 |
| AHG102 | SNP non-synonymous | *rpoB* | RNA polymerase subunit beta | S512F (C1534T) | MG1655 (NC_000913.3) | 4182745 |
| AHG104 | SNP synonymous | zinT | metal-binding protein ZinT | S23S (G68A) | MG1655 (NC_000913.3) | 2040634 |
| AHG104 | SNP intergenic | insH6/yoeA | CP4-44 prophage; IS5 transposase and trans-activator/CP4-44 prophage; TonB-dependent receptor plug domain-containing protein YoeA | T→C intergenic (-1316/-34) | MG1655 (NC_000913.3) | 2067792 |
| AHG104 | gene knock-out | yidK | putative transporter YidK | Missing coverage & new junction evidence | MG1655 (NC_000913.3) | 3858381-3860080 |

Continued on next page

**Table S3 – continued from previous page**

| Sample | Mutation | Gene name | Gene product | Annotation | Reference | Position |
|---|---|---|---|---|---|---|
| AHG104 | SNP non-synonymous | *rpoB* | RNA polymerase subunit beta | S512F (C1534T) | MG1655 (NC_000913.3) | 4182745 |
| AHG106 | SNP synonymous | nptII | Tn5 neomycin phosphotransferase | V83V (T248C) | pKD4 plasmid (AY048743.1) | 707 |
| AHG106 | SNP synonymous | zinT | metal-binding protein ZinT | S23S (G68A) | MG1655 (NC_000913.3) | 2040634 |
| AHG106 | SNP intergenic | insH6/yoeA | CP4-44 prophage; IS5 transposase and trans-activator/CP4-44 prophage; TonB-dependent receptor plug domain-containing protein YoeA | T→C intergenic (-1316/-34) | MG1655 (NC_000913.3) | 2067792 |
| AHG106 | gene knock-out | nuoC | NADH:quinone oxidoreductase subunit CD | Missing coverage & new junction evidence | MG1655 (NC_000913.3) | 2401267-2403045 |
| AHG106 | SNP non-synonymous | *rpoB* | RNA polymerase subunit beta | S512F (C1534T) | MG1655 (NC_000913.3) | 4182745 |
| AHG098 | SNP intergenic | –/nptII | –/Tn5 neomycin phosphotransferase | C→T intergenic (–/-223) | pKD4 plasmid (AY048743.1) | 236 |
| AHG098 | SNP intergenic | –/nptII | –/Tn5 neomycin phosphotransferase | A→T intergenic (–/-88) | pKD4 plasmid (AY048743.1) | 371 |
| AHG098 | SNP synonymous | zinT | metal-binding protein ZinT | S23S (G68A) | MG1655 (NC_000913.3) | 2040634 |
| AHG098 | SNP intergenic | insH6/yoeA | CP4-44 prophage; IS5 transposase and trans-activator/CP4-44 prophage; TonB-dependent receptor plug domain-containing protein YoeA | T→C intergenic (-1316/-34) | MG1655 (NC_000913.3) | 2067792 |
| AHG098 | SNP synonymous | gcvP | glycine decarboxylase | L822L (C2465T) | MG1655 (NC_000913.3) | 3046543 |

Continued on next page

**Table S3 – continued from previous page**

| Sample | Mutation | Gene name | Gene product | Annotation | Reference | Position |
|---|---|---|---|---|---|---|
| AHG098 | gene knock-out | waaP | lipopolysaccharide core heptose (I) kinase | Missing coverage (figure S8d) & new junction evidence | MG1655 (NC_000913.3) | 3805141-3805914 |
| AHG098 | SNP non-synonymous | *rpoB* | RNA polymerase subunit beta | S512F (C1534T) | MG1655 (NC_000913.3) | 4182745 |
| AHG098 | SNP synonymous | nrfA | cytochrome c552 nitrite reductase | A440A (C1319A) | MG1655 (NC_000913.3) | 4289049 |
| AHG068 | large deletion | ntpII | kanamycin resistance,bla; | Δ3267 bp (no reads mapped to pKD4 plasmid, figure S3c) | pKD4 plasmid (AY048743.1) | 1-3267 |
| AHG068 | SNP synonymous | ascB | 6-phospho-beta-glucosidase AscB | K366K (G1097A) | MG1655 (NC_000913.3) | 2842054 |
| AHG068 | SNP non-synonymous | *rpoB* | RNA polymerase subunit beta | I572S (T1714G) | MG1655 (NC_000913.3) | 4182925 |
| AHG070 | SNP synonymous | nptII | Tn5 neomycin phosphotransferase | S104S (A311G) | pKD4 plasmid (AY048743.1) | 770 |
| AHG070 | gene knock-out | ybfP/potE | lipoprotein YbfP/putrescine transporter PotE | Missing coverage (figure S9c) & new junction evidence | MG1655 (NC_000913.3) | 715192-715578 |
| AHG070 | SNP non-synonymous | *rpoB* | RNA polymerase subunit beta | I572S (T1714G) | MG1655 (NC_000913.3) | 4182925 |
| AHG072 | gene knock-out | marR | DNA-binding transcriptional repressor MarR | Missing coverage & new junction evidence | MG1655 (NC_000913.3) | 1618315-1618725 |
| AHG072 | SNP non-synonymous | *rpoB* | RNA polymerase subunit beta | I572S (T1714G) | MG1655 (NC_000913.3) | 4182925 |
| AHG074 | gene knock-out | yidK | putative transporter YidK | Missing coverage & new junction evidence | MG1655 (NC_000913.3) | 3858382-3860079 |
| AHG074 | SNP non-synonymous | *rpoB* | RNA polymerase subunit beta | I572S (T1714G) | MG1655 (NC_000913.3) | 4182925 |
| AHG076 | SNP synonymous | nptII | Tn5 neomycin phosphotransferase | V83V (T248C) | pKD4 plasmid (AY048743.1) | 707 |

Continued on next page

**Table S3 – continued from previous page**

| Sample | Mutation | Gene name | Gene product | Annotation | Reference | Position |
|---|---|---|---|---|---|---|
| AHG076 | SNP non-synonymous | sieB | Rac prophage; phage superinfection exclusion protein | N135K (C404A) | MG1655 (NC_000913.3) | 1418266 |
| AHG076 | gene knock-out | nuoC | NADH:quinone oxidoreductase subunit CD | Missing coverage & new junction evidence | MG1655 (NC_000913.3) | 2401265-2403045 |
| AHG076 | SNP non-synonymous | glpD | aerobic glycerol 3-phosphate dehydrogenase | D263Y (G786T) | MG1655 (NC_000913.3) | 3562766 |
| AHG076 | SNP non-synonymous | *rpoB* | RNA polymerase subunit beta | I572S (T1714G) | MG1655 (NC_000913.3) | 4182925 |
| AHG078 | SNP intergenic | –/nptII | –/Tn5 neomycin phosphotransferase | C→T intergenic (–/-223) | pKD4 plasmid (AY048743.1) | 236 |
| AHG078 | SNP intergenic | –/nptII | –/Tn5 neomycin phosphotransferase | A→T intergenic (–/-88) | pKD4 plasmid (AY048743.1) | 371 |
| AHG078 | gene knock-out | waaP/waaG | lipopolysaccharide core heptose (I) kinase / lipopolysaccharide glucosyltransferase I | Missing coverage (figure S8b) & new junction evidence | MG1655 (NC_000913.3) | 3805136-3805931 |
| AHG078 | SNP non-synonymous | *rpoB* | RNA polymerase subunit beta | I572S (T1714G) | MG1655 (NC_000913.3) | 4182925 |

#### 1.2 Antibiotic dose response curves of the wild-type genotype

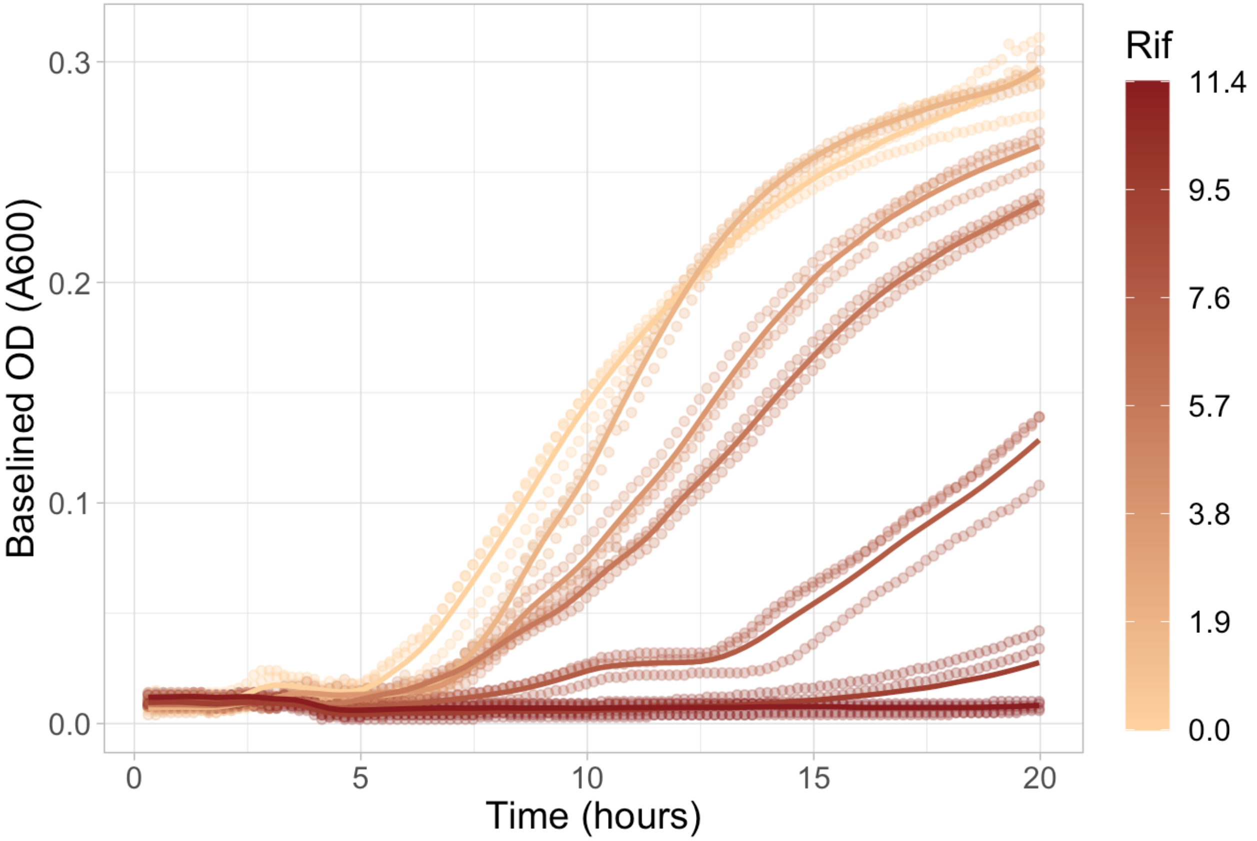

Figure S14: Dose response curves of the antibiotic-susceptible, wild-type genotype (sample AHG015) to different rifampicin concentrations. The y-axis shows the time in hours, the x-axis shows the baseline-subtracted optical density (OD) at 600nm, and the colours indicate different rifampicin concentrations with the legend on the right in units of $\mu g/mL$. Each line shows the average of three growth curve replicates (loess smoothing was used to interpolate) and points show the individual replicates. Baselining was done by subtracting the smallest observed OD value (including blank wells) at each time point.

#### 1.3 Ascertaining flow cytometry estimates of competitive fitness

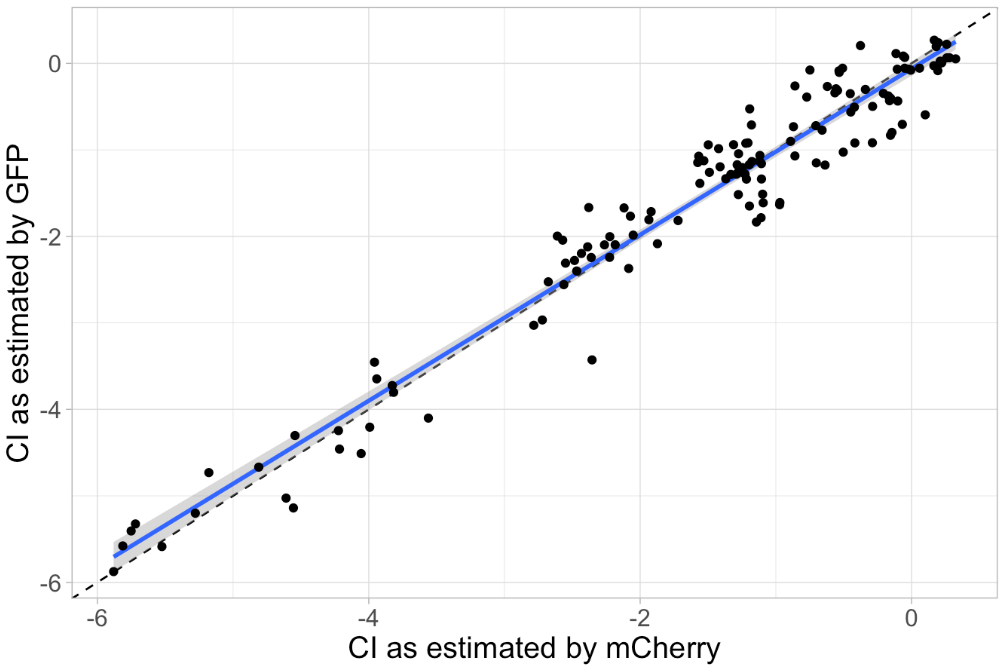

Figure S15: **Nearly identical $\hat{w}$ estimates were found when the fluorophores of the reference and competitor genotypes were swapped.** The competitive index ($\hat{w}$) as estimated by using mCherry-labeled competitor genotypes and a GFP-labeled reference genotype (shown in x-axis) corresponds well with the same metric estimated by using GFP-labeled competitor genotypes and an mCherry-labeled reference genotype (shown in y-axis). Data corresponds to a sample of genotypes representing all knock-outs and *rpoB* mutants as assayed once in all environments. The dashed, black line shows $y = x$. The blue line shows the mean and the blue shaded region shows the 95% confidence interval of the mean for the least-squares estimated linear regression. The coefficients of this regression are given in table S4; the overall regression is statistically significant and explains most of the variation in the response variable (adjusted $R^2 = 0.949$, $F(1, 138) = 2586$, $p < 10^{-15}$).

Table S4: Least-squares estimated coefficients of the linear regression shown in figure S15.

| **Coefficient** | **Estimate** | **Standard Error** | **t statistic** | **p-value** |
|---|---|---|---|---|
| Intercept (b) | -0.0619 | 0.0410 | -1.51 | 0.133 |
| CI as estimated by mCherry ($\beta$) | 0.960 | 0.0189 | 50.8 | $< 10^{-15}$ |

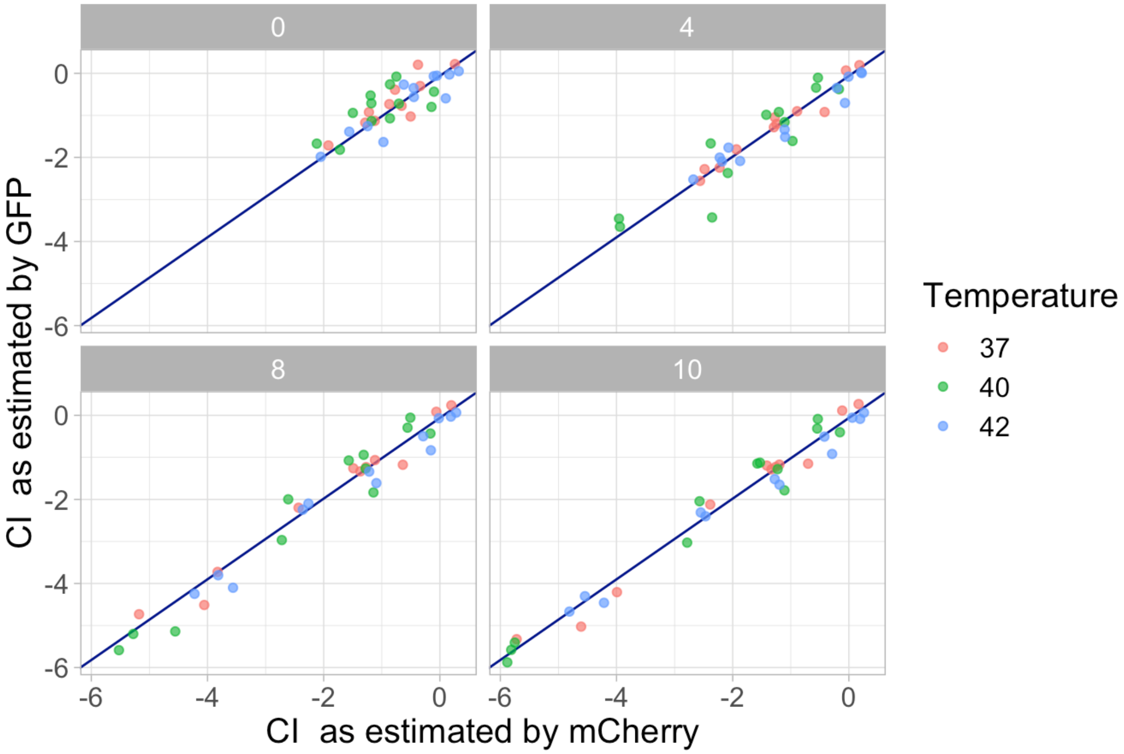

Figure S16: The plot shows environment-specific effects of swapping fluorophores. The same data is plotted as in figure S15 but here panels show the different antibiotic concentrations and points are coloured by temperature. The blue line shows the simple linear regression on CI as estimated by mCherry, and whose coefficient estimates are given in table S4. As shown in table S5, the effect of environment is not significant.

Table S5: Results of ANOVA show that the environment-specific effect of swapping fluorophores is not statistically significant.

| **Source of Variation** | **SS** | **df** | **MS** | **F statistic** | **p-value** |
|---|---|---|---|---|---|
| CI as estimated by mCherry | 300 | 1 | 300 | 2641 | $< 10^{-15}$ |
| Environment | 1.59 | 11 | 0.14 | 1.27 | 0.25 |
| Residuals | 14.4 | 127 | 0.11 | | |

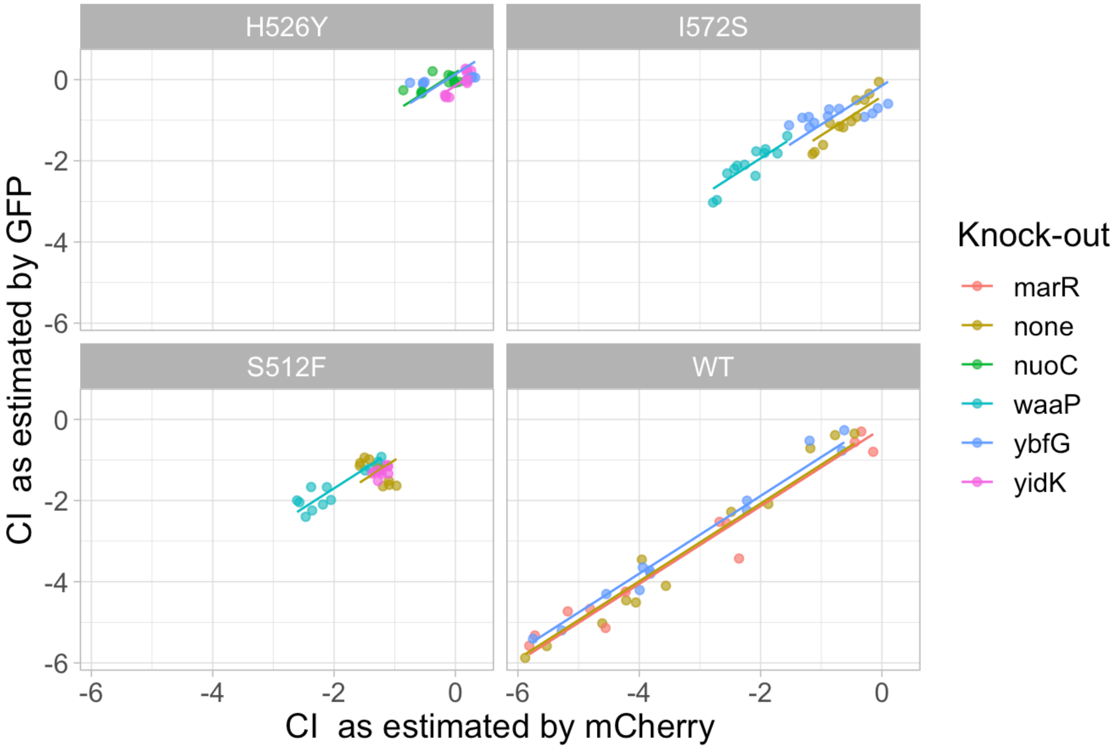

Figure S17: The plot shows genotype-specific effects of swapping fluorophores. The same data is plotted as in figure S15 but here panels show the different *rpoB* substitutions and points are coloured by genotype. Lines show the multiple regression on genotype as detailed in table S6. This effect is statistically significant; however it explains only about 1% more of the variation in the data (adjusted $R^2 = 0.9588$ for overall regression) as compared to not including this effect.

Table S6: Results of ANOVA show that the genotype-specific effect of swapping fluorophores is statistically significant but small.

| **Source of Variation** | **SS** | **df** | **MS** | **F statistic** | **p-value** |
|---|---|---|---|---|---|
| CI as estimated by mCherry | 300 | 1 | 300 | 3204 | $< 10^{-15}$ |
| Genotype | 4.12 | 11 | 0.37 | 4.00 | $< 10^{-4}$ |
| Residuals | 11.9 | 127 | 0.09 | | |

#### 1.4 Cost of resistance

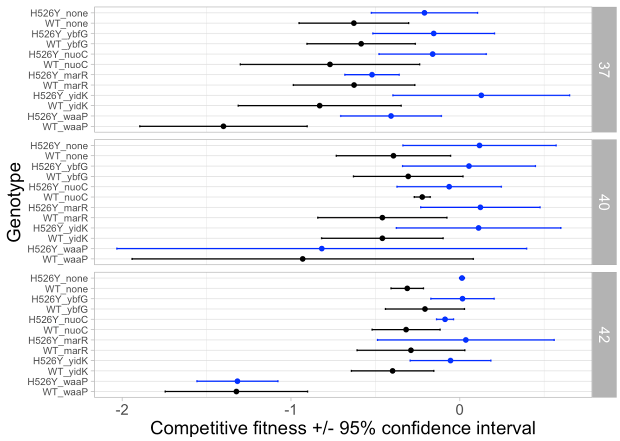

(a) ABR *rpoB* H526Y shown in blue.

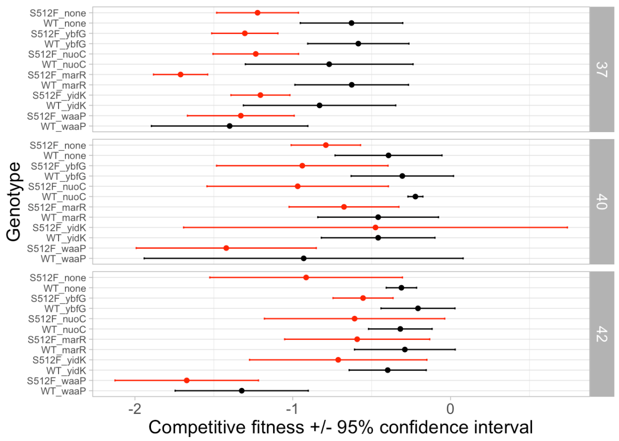

(b) ABR *rpoB* S512F shown in red.

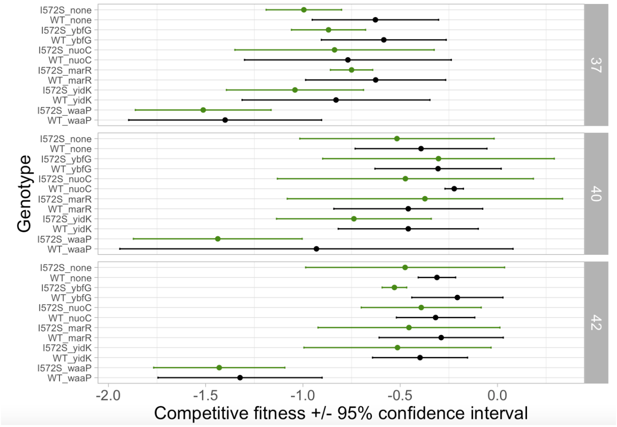

(c) ABR *rpoB* I572S shown in green.

Figure S18: Competitive fitness for ABR mutations in the absence of antibiotic ($0\mu g/mL$ rifampicin) indicates the cost of resistance. Each panel shows a different ABR *rpoB* mutation (colours) as compared to the wild-type *rpoB* (black). The x-axis indicates the competitive index, with points showing mean values and error bars showing the 95% confidence interval on the mean values. The y-axis shows the genotype, with first the *rpoB* mutation then the knock-out (e.g., "H526Y_nuoC" indicates the H526Y mutation at *rpoB* on the $\Delta nuoc$ background); "WT" refers to the wild-type *rpoB* sequence and "none" indicates that there is no knock-out. In each panel, the facets show the three different temperature environments. **(a)** Contrary to previous studies (1; 2), the H526Y mutation has no significant fitness cost – in fact, it exhibits a significantly higher fitness than wild-type at $42°C$ and has a fitness advantage on the $\Delta waaP$ background at $37°C$. **(b)** The S512F mutation has no significant fitness cost on most backgrounds except two at $37°C$ ($\Delta ybfG$ and $\Delta marR$, but not on the wild-type *rpoB* background) and one ($\Delta nuoC$) at $40°C$. This is contrary to previous studies that have found S512F to have a higher fitness than wild-type at $40°C$ (1). **(c)** The I572S mutation has no significant fitness costs, except on the $\Delta ybfG$ background at $42°C$.

#### 1.5 GxG summary statistics

##### 1.5.1 'Classical' pairwise epistasis

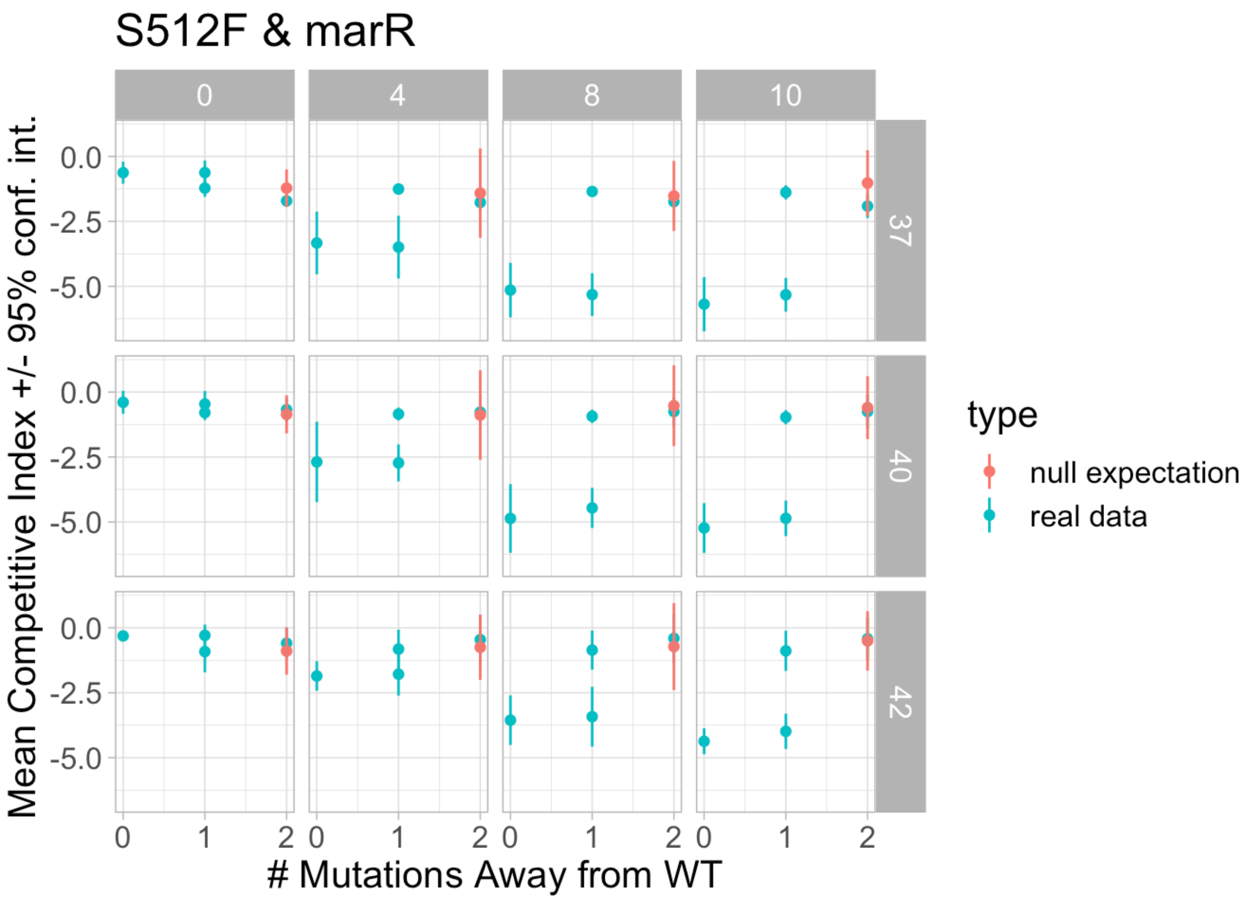

Figure S19: An illustration of how pairwise epistasis was calculated, in this case for the combination of the S512F *rpoB* ABR and the $\Delta marR$ knock-out mutations. The plot shows the observed mean competitive index (± the 95% confidence interval as determined from parametric boostrapping) in blue on the y-axis as a dependent-variable of the number of mutations away from wild-type (x-axis) in each of the 12 environments. Different antibiotic concentrations are shown in each column and different temperatures are shown on each row. The red points show the mean competitive index expected under the null hypothesis of no epistasis. In this example there are no instances where the epistasis is significantly different from zero as shown by the consistent overlap between the 95% confidence interval error bars and the points indicating mean values. Figure S21 summarizes the pairwise epistasis estimates for all genotypes combinations in all environments.

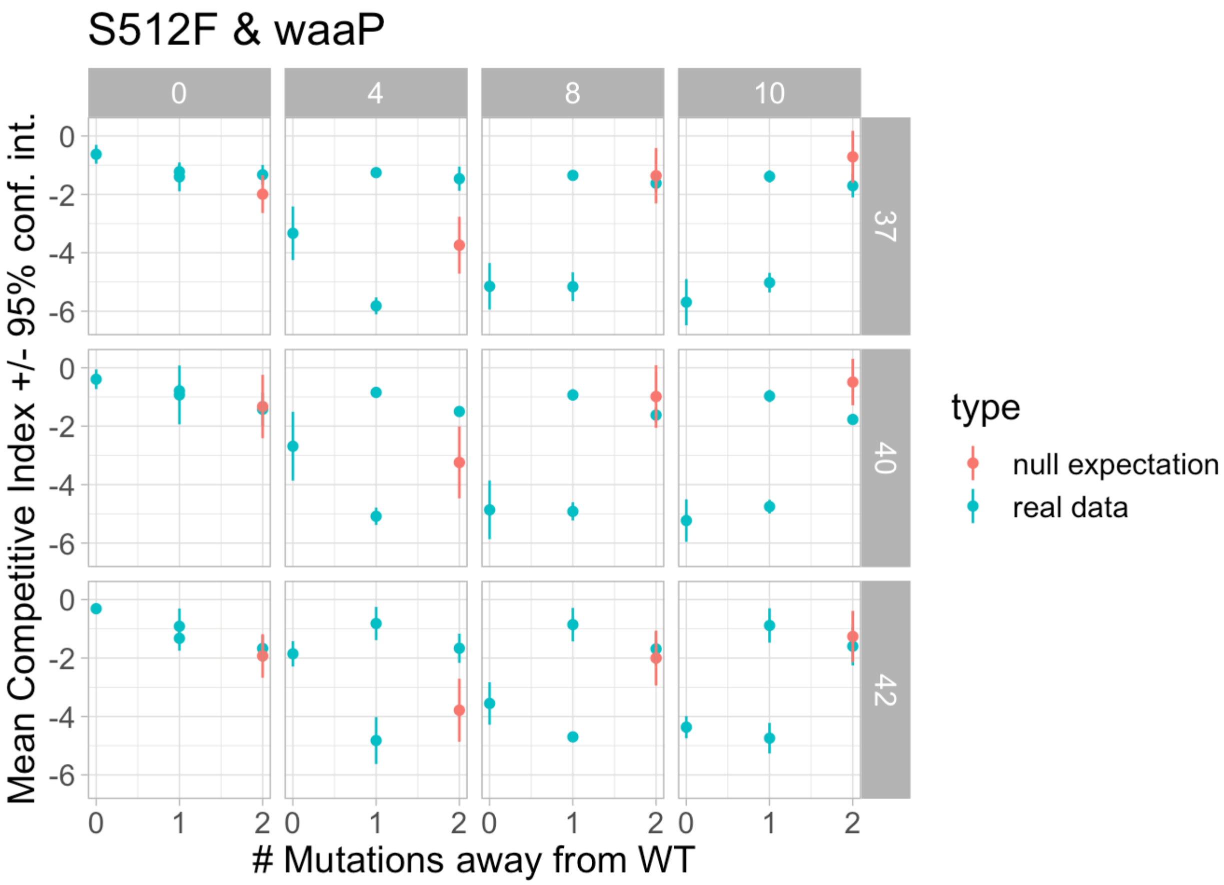

Figure S20: An example of significant pairwise epistasis, in this case for the combination of the S512F *rpoB* ABR and the $\Delta waaP$ knock-out mutations. The data is plotted in the same style as described in figure S19. Positive epistasis is apparent in all of the temperature environments with 4$\mu$g/mL rifampicin, as seen by the larger fitness values of the real data (blue points) as compared to the null expectation of no epistasis (red points). No overlap between the 95% confidence intervals and the mean indicates that the effect is statistically significant. The equivalent plots for $\Delta waaP$ in combination with the other ABR mutants, H526Y and I572S, show a similar trend of positive epistasis at 4$\mu$g/mL rifampicin, as summarized in figure S21. The 40°$C$ and 10$\mu$g/mL environment displays significant negative pairwise epistasis.

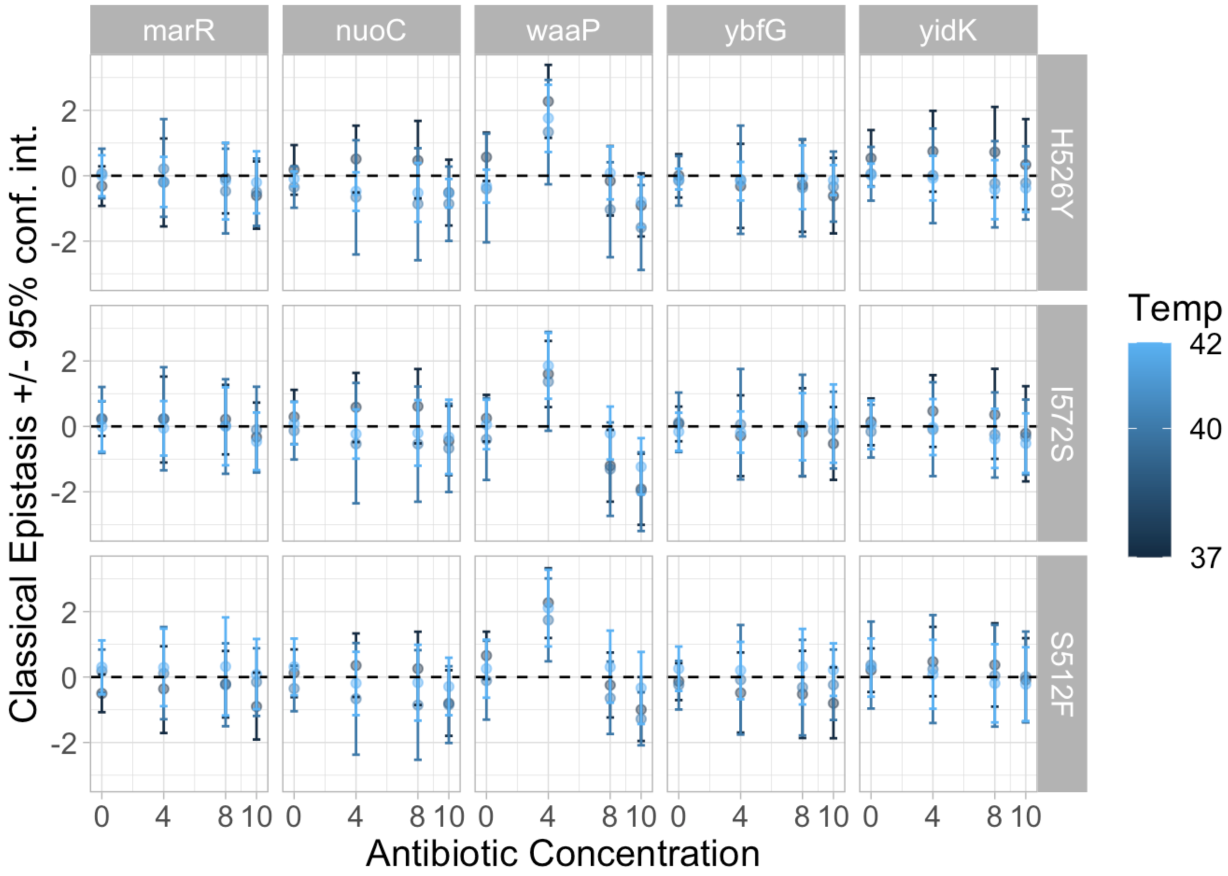

Figure S21: **Estimated pairwise epistasis across all genotypes and environments.** The plots are laid out in the same manner as figure 2 except that the y-axis here shows the mean estimated 'classical' pairwise epistasis and error bars show the 95% confidence interval as determined from parametric bootstrapping.

##### 1.5.2 Gamma epistasis

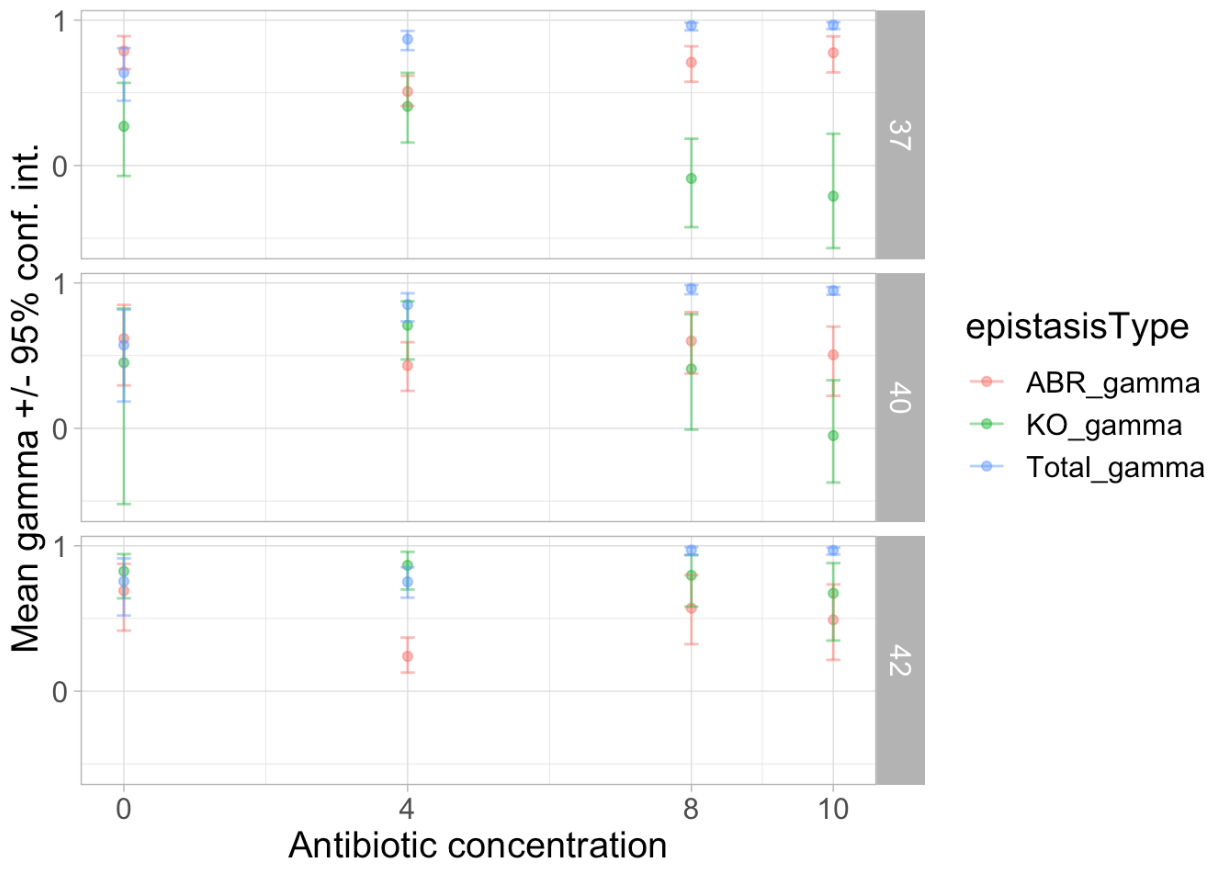

Figure S22: The amount of gamma epistasis is different for different gene classes and temperature environments. Points show the mean gamma epistasis for different gene classes as indicated by the colour (red for ABR mutations at *rpoB*, green for gene knock-out mutations, and blue for the total as already shown in figure 3a. The x-axis shows the rifampicin antibiotic concentration in $\mu g/mL$ and the rows show the three different temperature environments. The error bars show the 95% parametric bootstrap confidence interval. No regression lines are shown as the correlations are not significant at $\alpha = 0.01$.

##### 1.5.3 Fraction of sign epistasis

Table S7: List of the genotypes that exhibited reciprocal sign epistasis in more than 5% of the 10 000 parametric bootstrap samples. The counts of observed reciprocal sign epistasis for all bootstrap samples is given for each of the three temperature environments. Only the environment without anitbiotic is shown because the other environments did not exhibit reciprocal sign epistasis in more than 5% of bootstrap samples.

| **rpoB mutation** | **Knock-out mutation** | **Count at** $37°C$ | **Count at** $40°C$ | **Count at** $42°C$ |
|---|---|---|---|---|
| I572S | ΔnuoC | 3437 | 2514 | 2831 |
| I572S | ΔmarR | 2327 | 3538 | 2632 |
| I572S | ΔyidK | 1688 | 2331 | 2586 |
| I572S | ΔybfG | 698 | 2864 | 2242 |
| S512F | ΔyidK | 551 | 3236 | 1469 |
| H526Y | ΔybfG | 596 | 976 | 694 |
| H526Y | ΔmarR | 2480 | ( <500 ) | 1227 |
| S512F | ΔwaaP | 2357 | ( <500 ) | ( <500 ) |
| S512F | ΔmarR | ( <500 ) | 1831 | 1276 |
| H526Y | ΔnuoC | ( <500 ) | 1473 | ( <500 ) |
| H526Y | ΔwaaP | ( <500 ) | 1381 | ( <500 ) |
| I572S | ΔwaaP | ( <500 ) | 744 | ( <500 ) |
| H526Y | ΔyidK | ( <500 ) | 565 | ( <500 ) |
| S512F | ΔnuoC | ( <500 ) | ( <500 ) | 1268 |

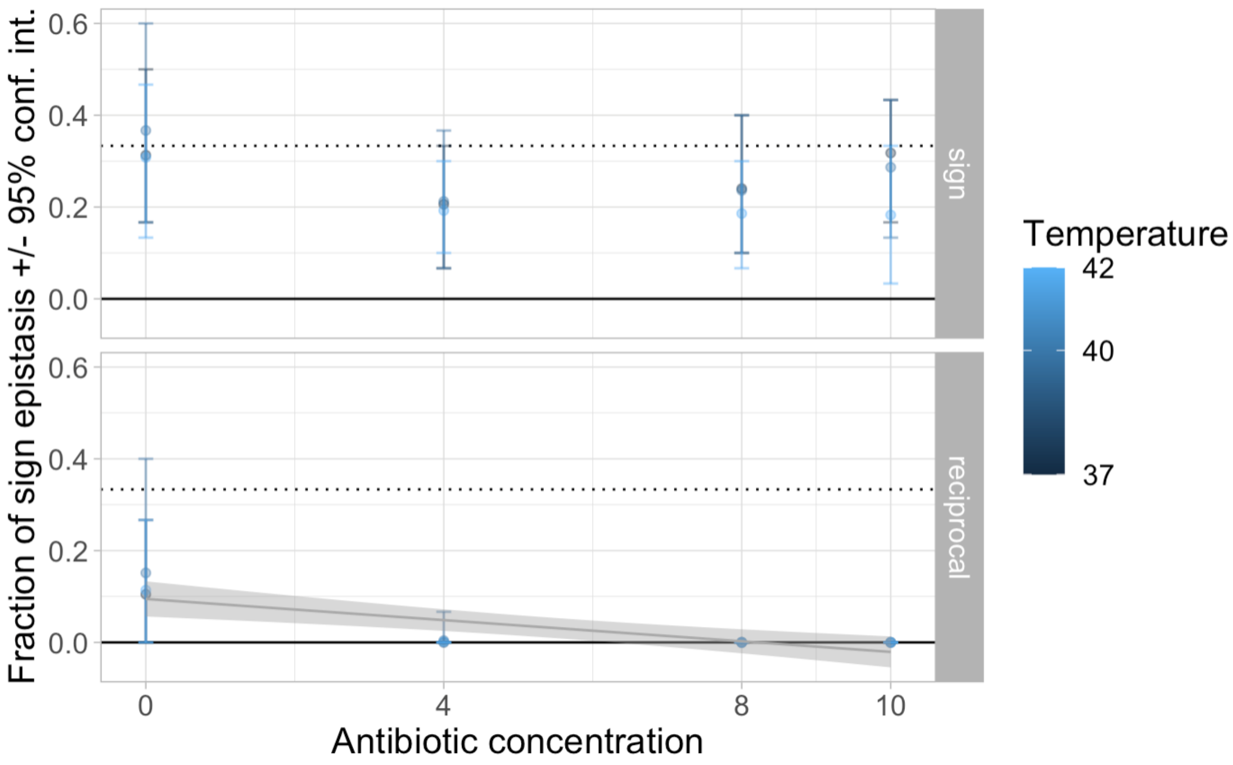

Figure S23: The fraction of simple and reciprocal sign epistasis for different environments. Reciprocal sign epistasis (bottom facet) is mostly observed in the absence of antibiotic while the fraction of simple sign epistasis (top facet) shows no apparent trend with different environments. The points show the mean fraction and the error bars show the 95% parametric bootstrap confidence interval. The x-axis shows the antibiotic concentration and the colour shows the temperature. The solid line at $y = 0$ highlights the fraction of simple and reciprocal sign epistasis expected in a purely additive landscape without any epistasis. The dotted line at $y = \frac{1}{3}$ highlights the fraction of simple and reciprocal sign epistasis expected in a completely epistatic house-of-cards landscape. There is no significant correlation between the antibiotic concentration and the fraction of simple sign epistasis ($F(1, 10) = 2.15$, $p = 0.17$). The grey trend-line shows the significant negative correlation of antibiotic concentration on the fraction of reciprocal sign epistasis ($F(1, 10) = 20.3$, $p < 0.005$, adjusted $R^2 = 0.637$) and the grey shaded region shows the 95% confidence interval of the mean for the least-squares estimated linear regression. Antibiotic concentration explains more of the variation in the fraction of reciprocal sign epistasis than environmental quality (figure S40).

###### 1.5.4 Roughness-to-slope ratio

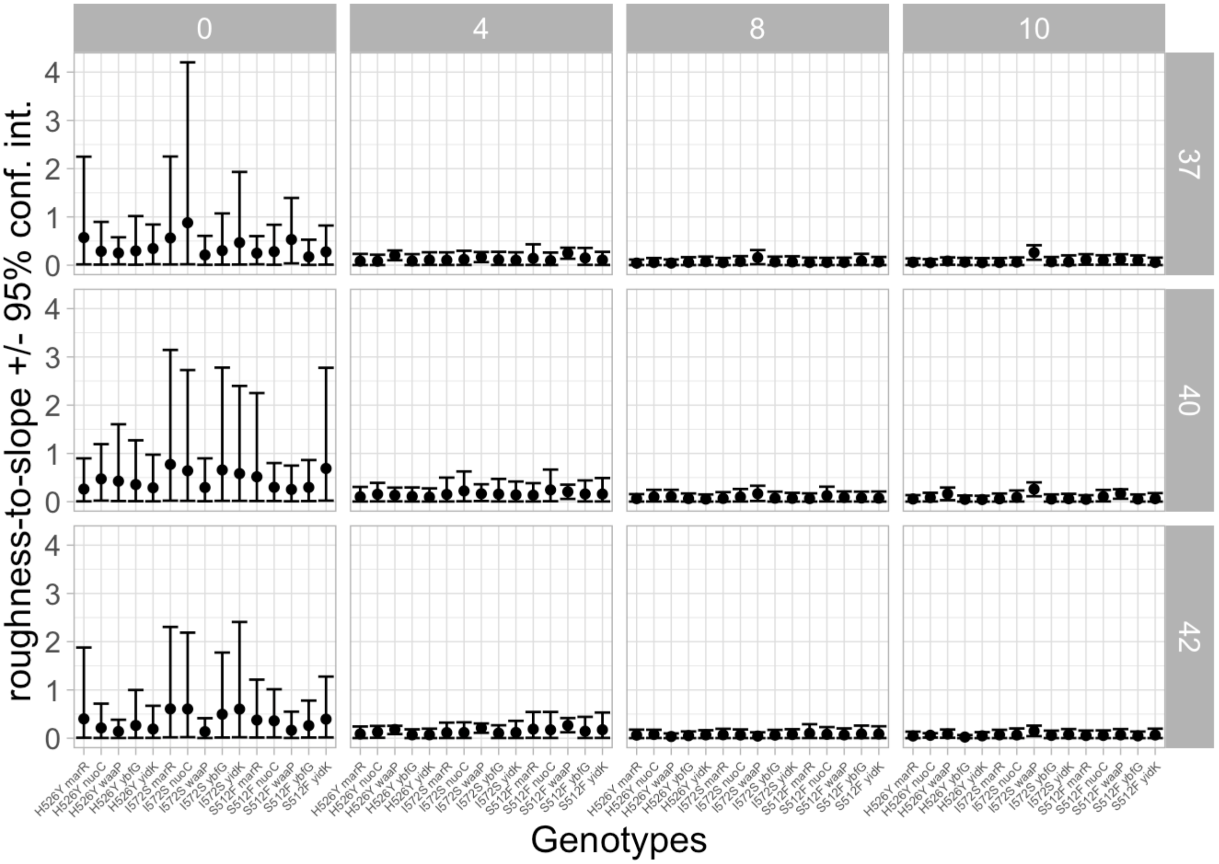

Figure S24: The roughness-to-slope ratio (y-axis) for each genotype (x-axis) in each environment. The dots show the mean roughness-to-slope ratios and the error bars show the 95% parametric bootstrap confidence interval. The columns show different antibiotic concentrations and the rows show different temperature environments.

##### 1.5.5 Diminishing-returns epistasis

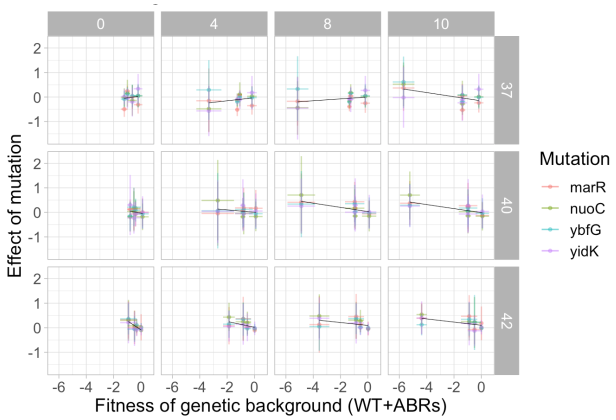

(a) knock-out mutations, excluding $\Delta waaP$

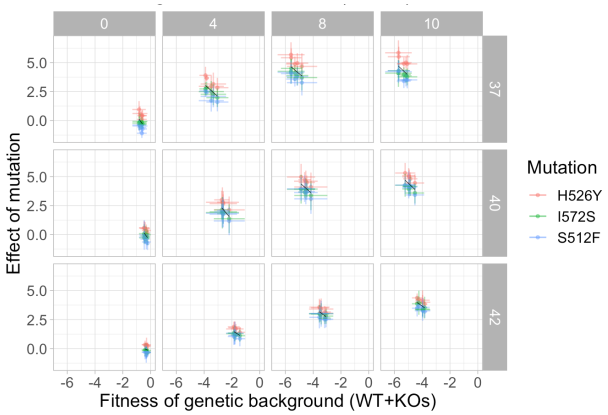

(b) ABR *rpoB* mutations, excluding $\Delta waaP$ background

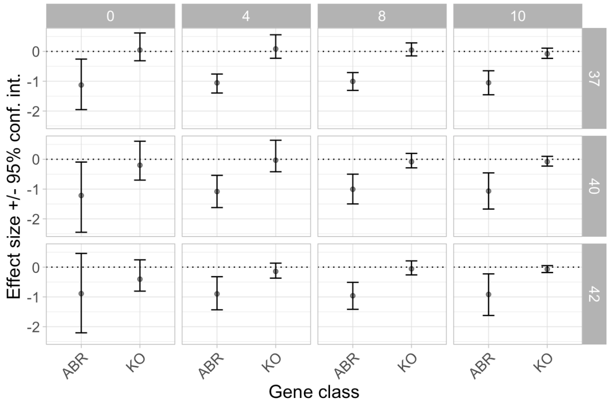

(c) Slopes of the diminishing-returns epistasis excluding $\Delta waaP$

Figure S25: **When the $\Delta waaP$ mutation and background are removed from the analysis, knock-outs continue to display no diminishing-returns epistasis and ABR *rpoB* mutations exhibit diminishing-returns epistasis in all but one of the environments.** The plots are in the same style as figure 4 except that the effect of the $\Delta waaP$ mutation and mutations on the $\Delta waaP$ background have been removed to confirm that $\Delta waaP$ is not responsible for the overall trend.

#### 1.6 Multiple regression for competitive fitness

##### 1.6.1 Additive ('null') model

Table S8: ANOVA shows that the additive effects of *rpoB* mutants, KO mutants, antibiotic concentration, and temperature are all significant.

| Source of Variation | SS | df | MS | F statistic | p-value |
|---|---|---|---|---|---|
| *rpoB* mutant | 1245 | 3 | 415.0 | 533.7 | $< 10^{-15}$ |
| KO mutant | 114.7 | 5 | 22.9 | 29.50 | $< 10^{-15}$ |
| Antibiotic concentration | 216.6 | 3 | 72.2 | 92.83 | $< 10^{-15}$ |
| Temperature | 66.7 | 2 | 33.3 | 42.86 | $< 10^{-15}$ |
| Residuals | 763.6 | 982 | 0.8 | | |

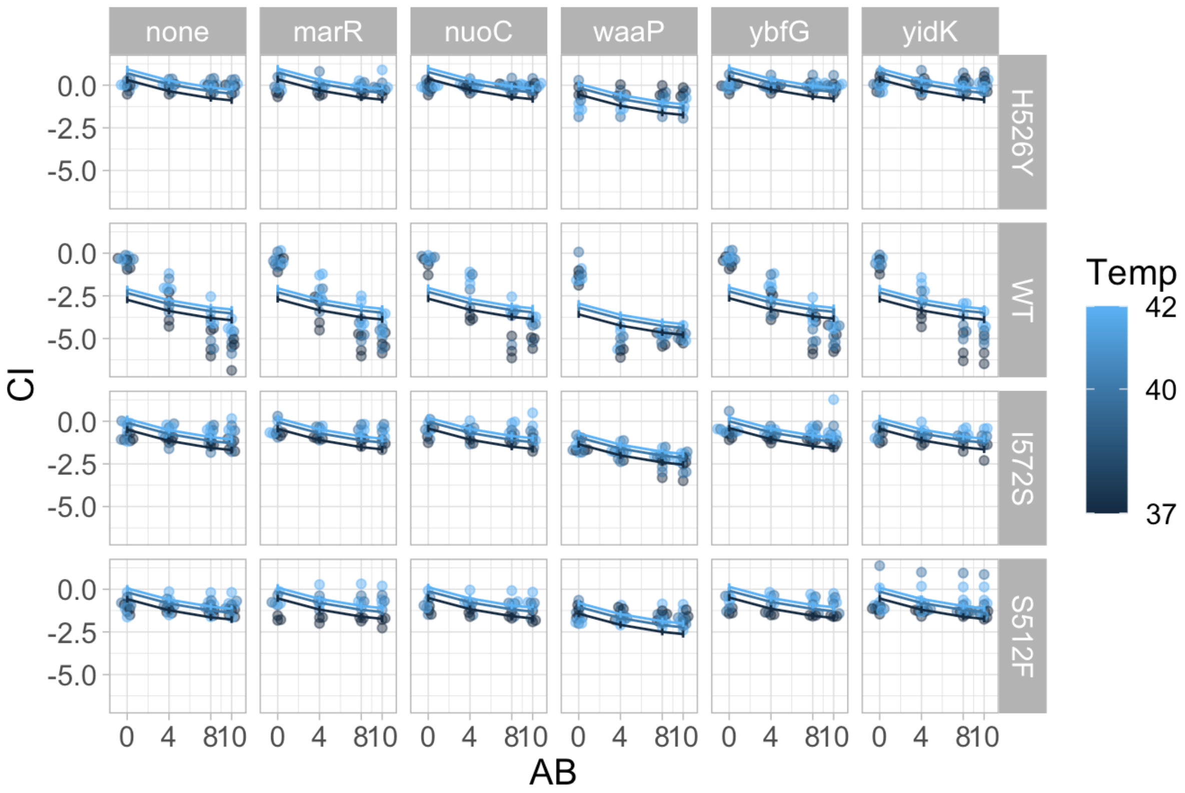

Figure S26: A linear regression with only additive effects explains almost 70% of the variation observed in the competition data. The x-axis shows the rifampicin antibiotic concentration (AB) in $\mu g/mL$ and the y-axis shows the $\hat{w}$ competitive index (CI). Points show the data and lines show the mean predictions with error-bars depicting the 95% confidence intervals for the means of the least-squares estimated linear regression. Therefore the space between the data points and the mean lines indicates the residuals of the model fitting. This plot shows us that much of the residual variation can be attributed to wild-type, antibiotic-susceptible genotypes (WT, second row) since this predictor has the biggest deviation between the data points and the model. This model was found to explain less of the variation than models with an interaction between *rpoB* genotypes and antibiotic environment.

##### 1.6.2 Models with first order interactions

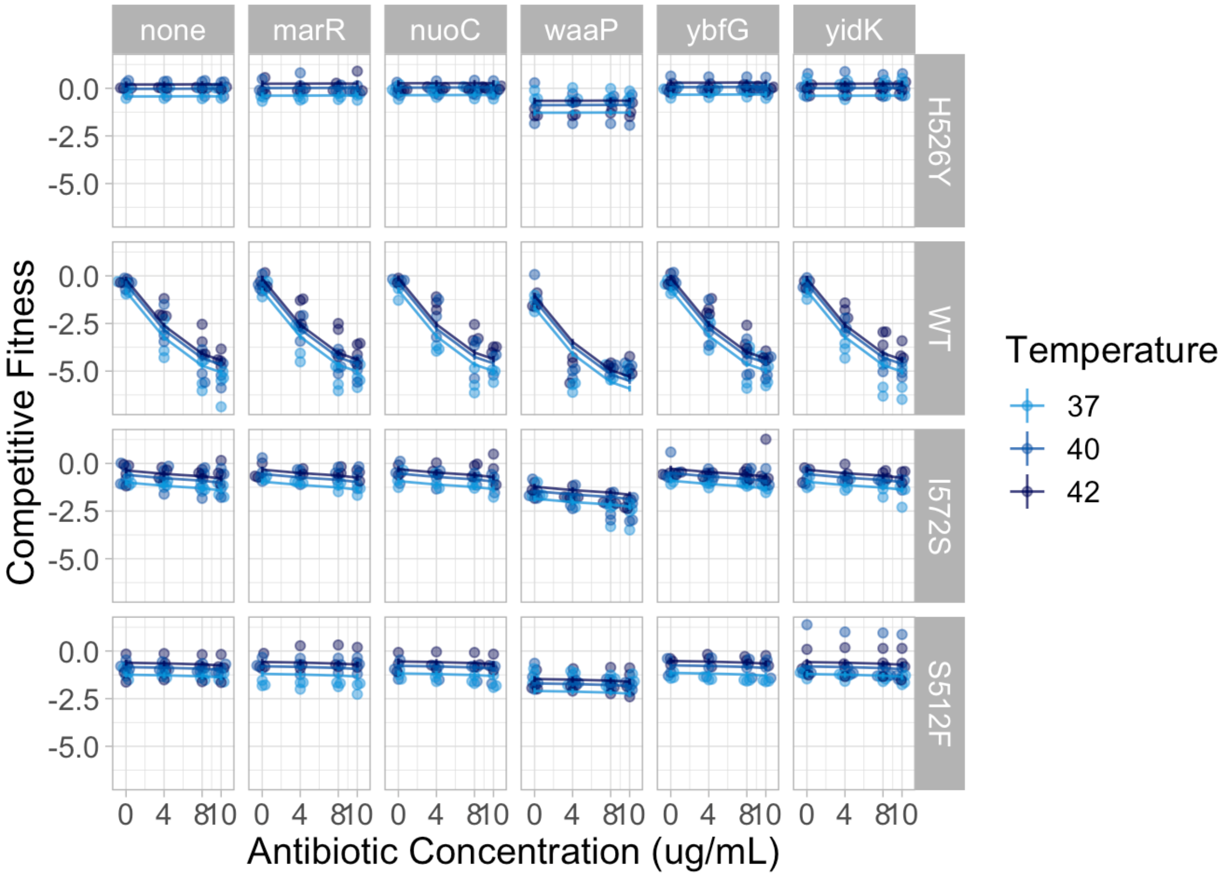

Figure S27: A linear regression with additive effects and $G_{rpoB} \times E_{\mathrm{AB}}$, an interaction between *rpoB* genotype and antibiotic environment, explains much of the variation observed in the competition data. The x-axis shows the rifampicin antibiotic concentration in $\mu g/mL$ and the y-axis shows the $\hat{w}$ competitive index. Points show the data and lines show the mean predictions with error-bars depicting the 95% confidence intervals for the means of the least-squares estimated linear regression. Therefore the space between the data points and the mean lines indicates the residuals of the model fitting. This plot shows that most of the systematic variation in the data is explained by this model.

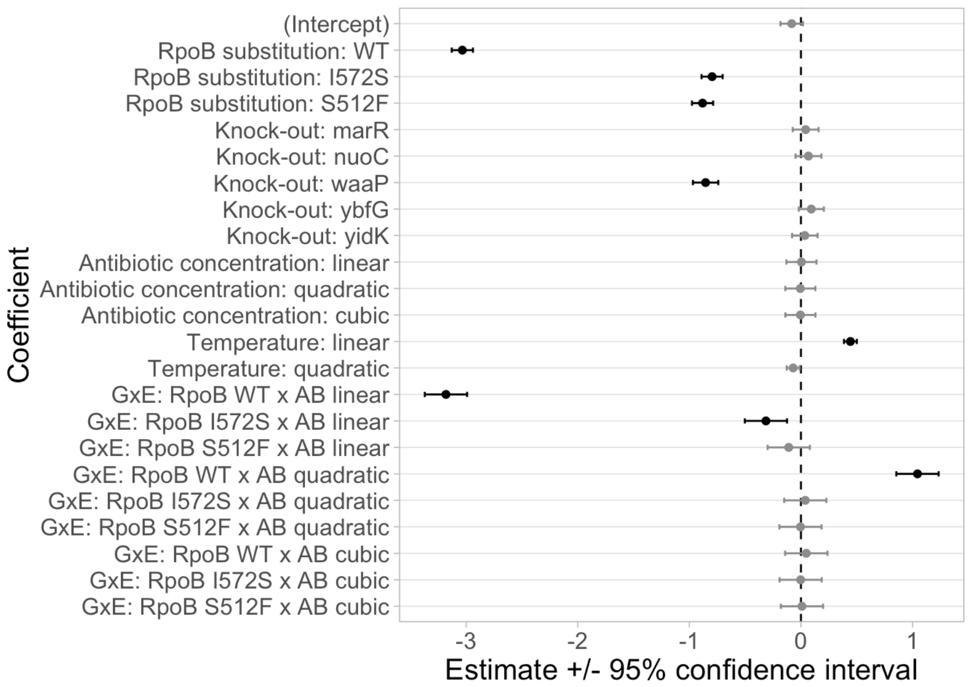

Figure S28: Estimated mean values for the coefficients in the regression model with additive effects and $G_{rpoB} \times E_{\mathrm{AB}}$, an interaction between *rpoB* genotype and antibiotic environment (error bars show 95% confidence intervals). The intercept is for *rpoB* with the H526Y substitution and without any knock-out mutation (i.e., the reference genotype for all competitions) as grown in the 37°C environment without any rifampicin antibiotic. Statistically significant coefficients ($\alpha = 0.05$) are shown in black and have confidence intervals that do not overlap with zero. Coefficients whose estimates are not statistically significant are shown in gray and have confidence intervals that overlap with zero (dashed vertical line).

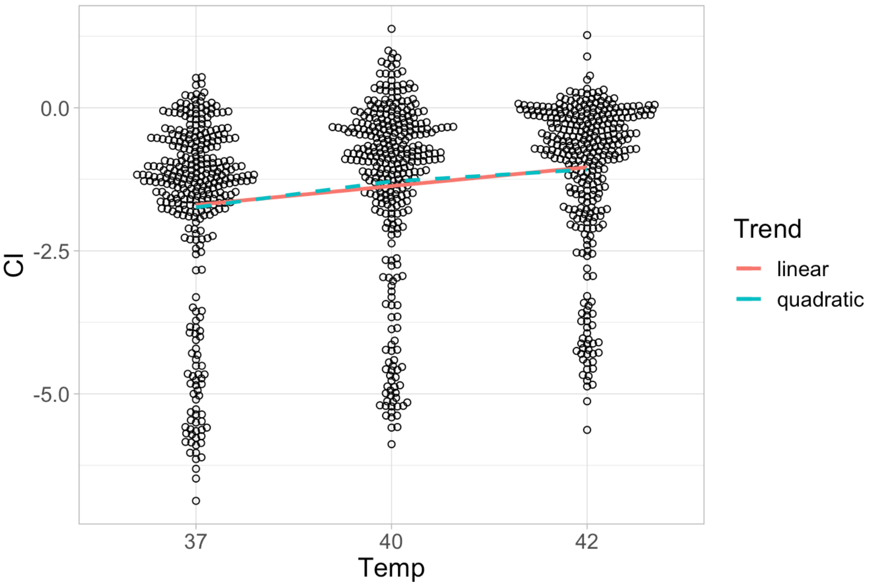

Figure S29: A beeswarm plot of the average effect of temperature (categorical x-axis) on estimated competitive index (quantitative y-axis) across all genotypes and antibiotic environments. Circles show individual estimates, but deviations along the x-axis away from the category labels are not quantitatively meaningful: it's merely to prevent overlapping of dense data points. The solid line shows the average linear (statistically significant) trend, and the dashed line shows the average quadratic (but **not** statistically significant) trend (see estimated coefficients in figures S28 and S31).

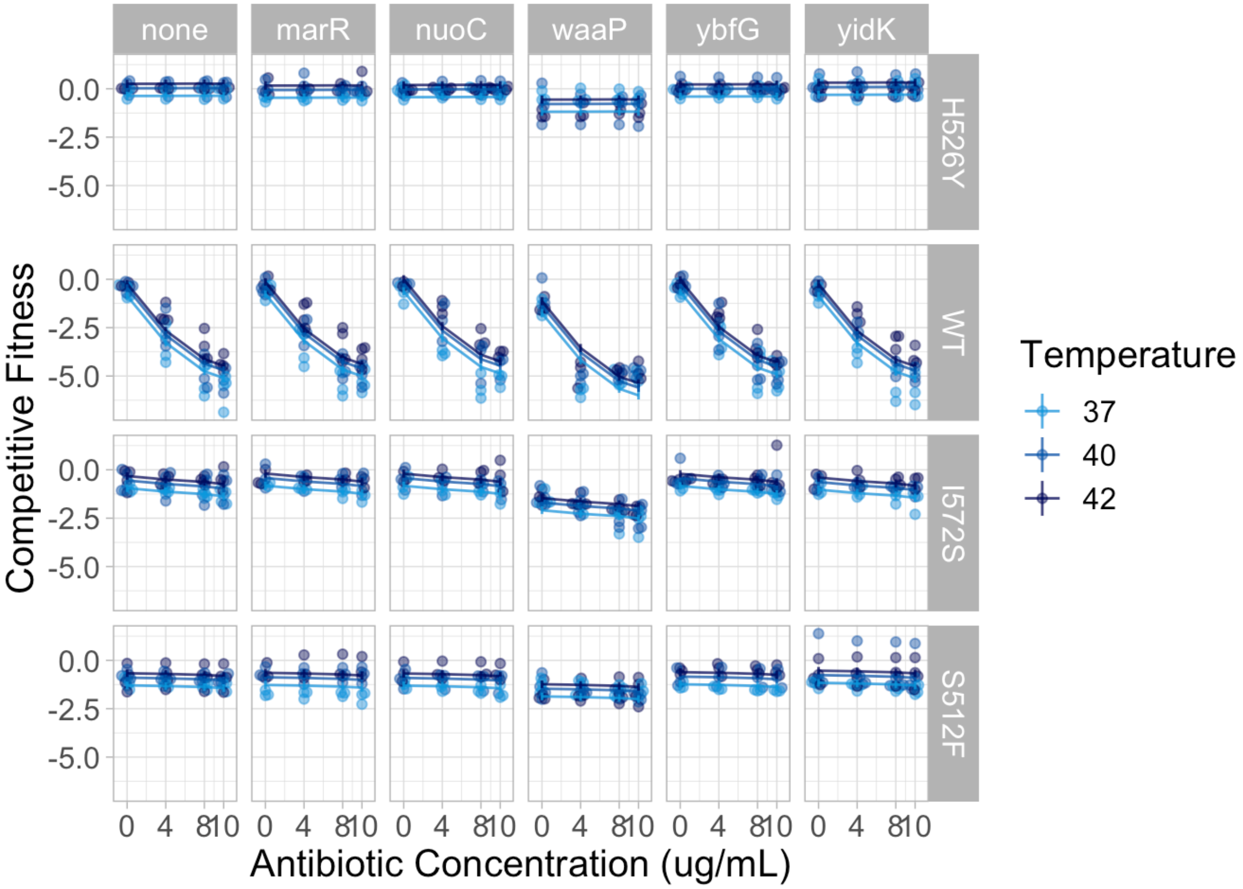

Figure S30: A linear regression with additive effects, an interaction between *rpoB* genotype and antibiotic environment ($G_{rpoB} \times E_{\mathrm{AB}}$), and a pairwise epistatic interaction between *rpoB* and knock-out genotypes ($G_{rpoB} \times G_{\mathrm{KO}}$) explains much of the variation observed in the competition data. The x-axis shows the rifampicin antibiotic concentration in $\mu g/mL$ and the y-axis shows the $\hat{w}$ competitive index. Points show the data and lines show the mean predictions with error-bars depicting the 95% confidence intervals for the means of the least-squares linear regression. Therefore the space between the data points and the mean lines indicates the residuals of the model fitting. This plot shows that most of the systematic variation in the data is explained by this model.

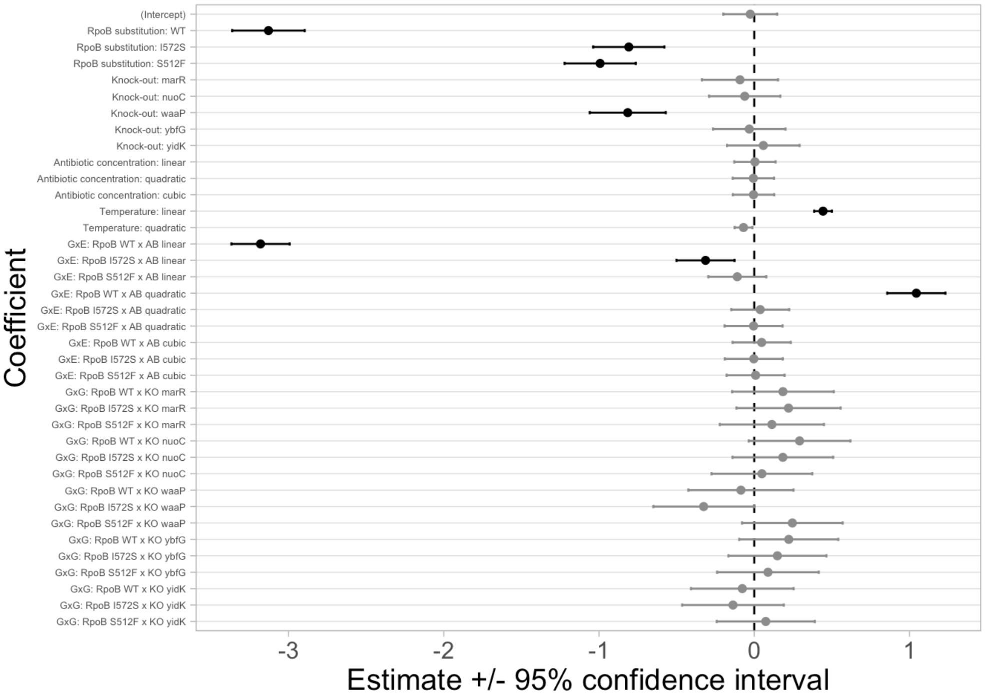

Figure S31: Estimated mean values for the coefficients in the regression model with additive effects and an interaction between *rpoB* genotype and antibiotic environment ($G_{rpoB} \times E_{\mathrm{AB}}$) as well as epistasis between *rpoB* and knock-out genotypes ($G_{rpoB} \times G_{\mathrm{KO}}$). Error bars show 95% confidence intervals. The intercept is for *rpoB* with the H526Y substitution and without any knock-out mutation (i.e., the reference genotype for all competitions) as grown in the 37°C environment without any rifampicin antibiotic. Statistically significant coefficients are shown in black and have confidence intervals that do not overlap with zero. Coefficients whose estimates are not statistically significant are shown in gray and have confidence intervals that overlap with zero (dashed vertical line). Notice that none of the coefficients associated with the epistatic interactions of *rpoB* and knock-out genotypes are significant. The interaction of *rpoB* I572S with the $\Delta waaP$ knock-out is barely significant ($p = 0.0492$).

##### 1.6.3 Models with second order interactions

We used multiple linear regression to look at the second-order interactions of the predictor variables on competitive fitness. In order to investigate second order interactions despite the risk of overfitting the data (n=996 whereas there are > 150 model parameters), we fitted the following complex model without reporting the p-values:

$$\hat{w} \sim \underbrace{G_{rpoB} + G_{\mathrm{KO}} + E_{\mathrm{AB}} + E_{\mathrm{T}}}_{\text{additve effects}} + \underbrace{G_{rpoB} \times E_{\mathrm{AB}} + E_{\mathrm{AB}} \times E_{\mathrm{T}}}_{1^{st}\text{ order interactions}} + \underbrace{G_{rpoB} \times G_{\mathrm{KO}} \times E_{\mathrm{AB}} + G_{rpoB} \times G_{\mathrm{KO}} \times E_{\mathrm{T}}}_{2^{nd}\text{ order interactions}},$$

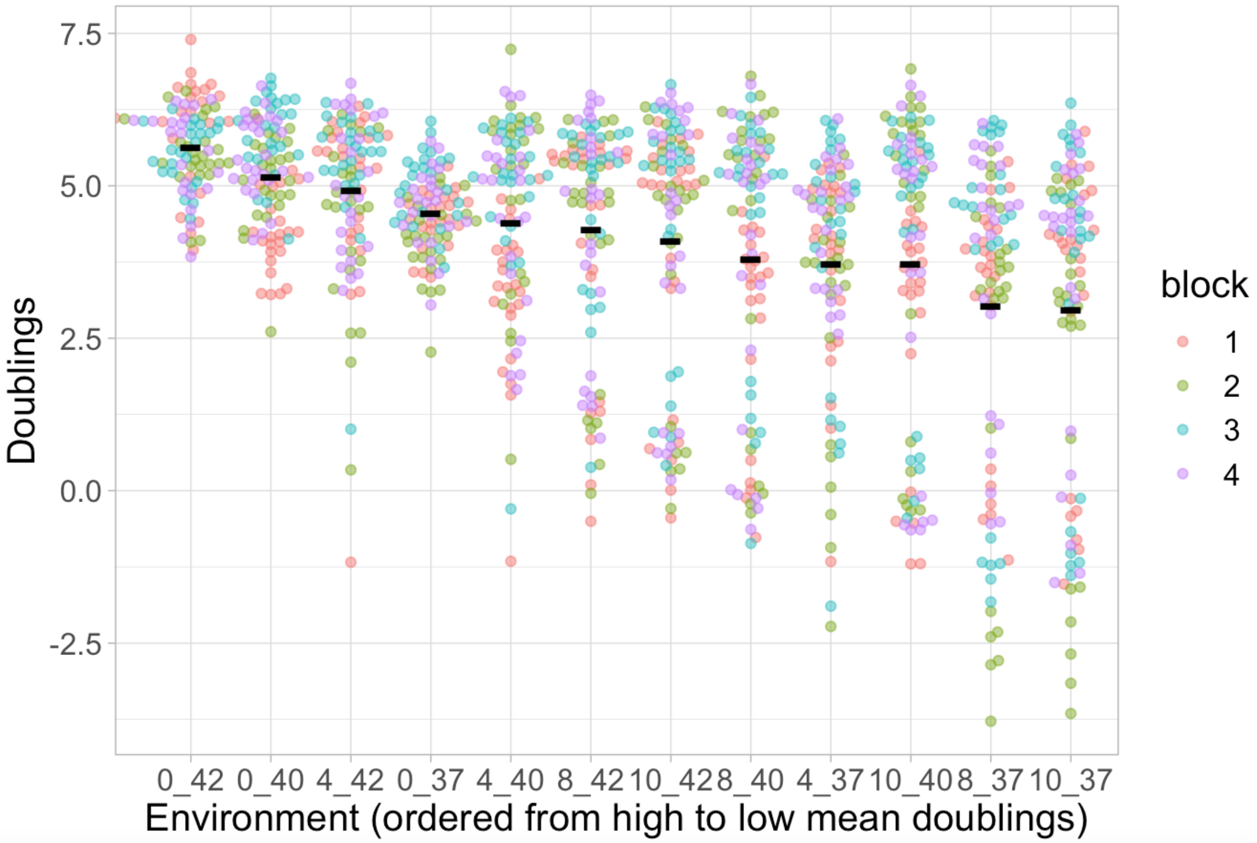

Figure S32: This plot shows that the environmental quality metric 'mean growth of all competitor strains in each environment' is able to distinguish between environments. The x-axis shows each environment in order from highest to lowest quality and the y-axis shows the total amount of growth estimated by the number of doublings observed between 0 and 20 hours (negative values suggest cell death). The black horizontal lines show the mean doublings across all replicates and genotypes: these values are used below as the metric of environmental quality. The coloured dots show the doublings of individual competitor genotypes and their replicates. Colours indicate the experimental replicate block and, ideally, should not show any trend. Both the environment and the replicate block explain a significant amount of the variation in the data (table S9). The genotypes that perform poorly in the low quality environments correspond to antibiotic susceptible *rpoB* wild-type genotypes as shown in figure S33.

Table S9: results of ANOVA for environmental quality metric: the mean growth of all competitor strains in each environment.

| **Source of Variation** | **SS** | **df** | **MS** | **F statistic** | **p-value** |
|---|---|---|---|---|---|
| Environment | 679 | 11 | 61.76 | 18.08 | $< 10^{-15}$ |
| Replicate block | 105 | 3 | 34.91 | 10.22 | $< 10^{-5}$ |
| Residuals | 3843 | 1125 | 3.42 | | |

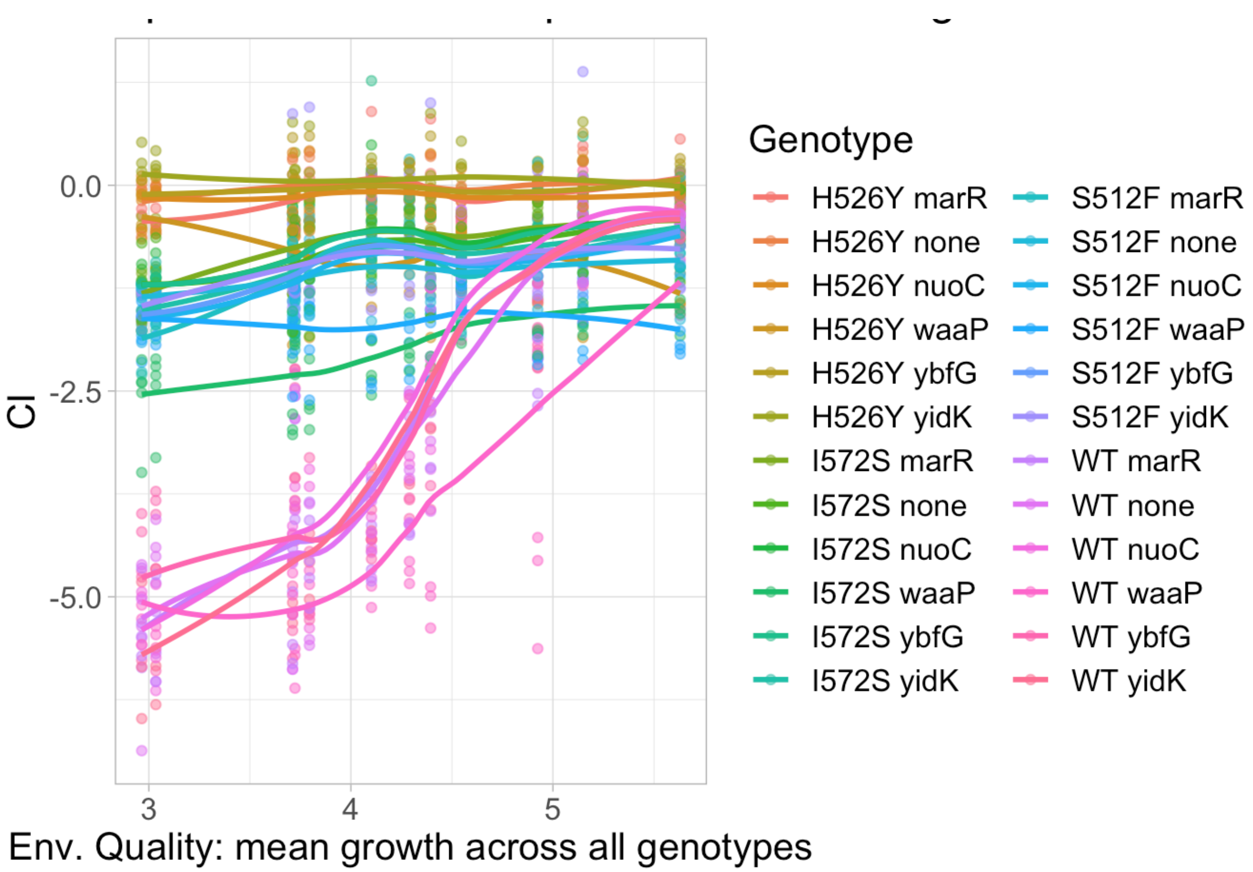

Figure S33: This plot shows that the environmental quality metric 'mean growth of all competitor strains in each environment' is able to distinguish specialist from generalist genotypes. The x-axis shows the 12 different abiotic environments as quantified by the environmental quality metric and ordered from lowest to highest quality; the y-axis shows the competitive index ($\hat{w}$) for each genotype, with values of individual replicates shown as dots ($n = 3{-}4$) and the mean value of each genotype shown as lines connected between environments. The colours indicate the genotype. All genotypes starting with "WT" are susceptible to rifampicin and therefore considered specialists to environments with no or low antibiotic concentrations. Conversely, all other genotypes are antibiotic resistant and can be considered generalists across all environments. Environments with higher antibiotic concentrations are grouped together as low quality and therefore susceptible genotypes perform poorly at lower values of the environmental quality metric (exhibiting a '/' or 'S' shape on the plot) and generalist genotypes perform about the same across all environments (exhibiting a straight line on the plot).

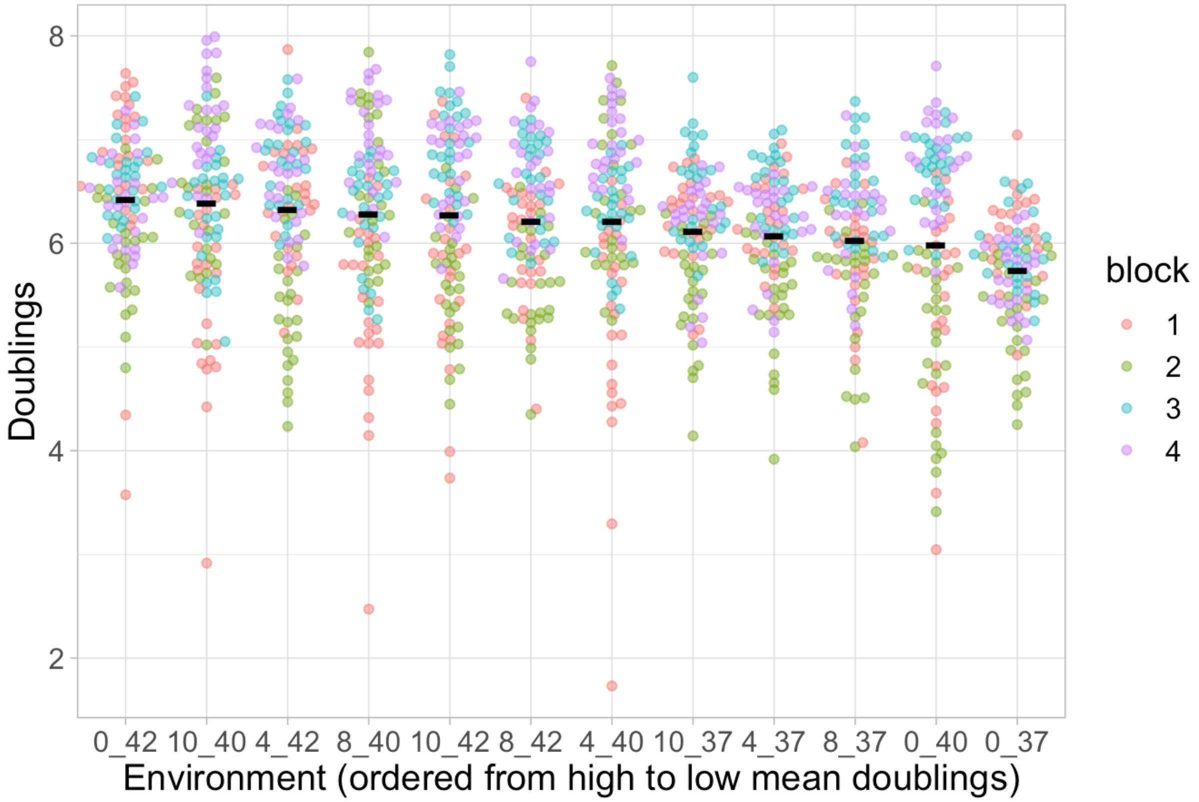

Figure S34: This plot shows that the environmental quality metric 'the mean growth of the reference strain, H526Y, in each environment' is not able to distinguish between environments. The data is plotted in the style of figure S32. However, unlike that figure, here there is a stronger effect of the replicate blocks as compared to environment (quantified in table S10).

Table S10: results of ANOVA for environmental quality metric: the mean growth of the reference strain, H526Y, in each environment.

| **Source of Variation** | **SS** | **df** | **MS** | **F statistic** | **p-value** |
|---|---|---|---|---|---|
| Environment | 40.6 | 11 | 3.69 | 8.01 | $< 10^{-12}$ |
| Replicate block | 151.7 | 3 | 50.56 | 109.83 | $< 10^{-15}$ |
| Residuals | 534.5 | 534.5 | 0.46 | | |

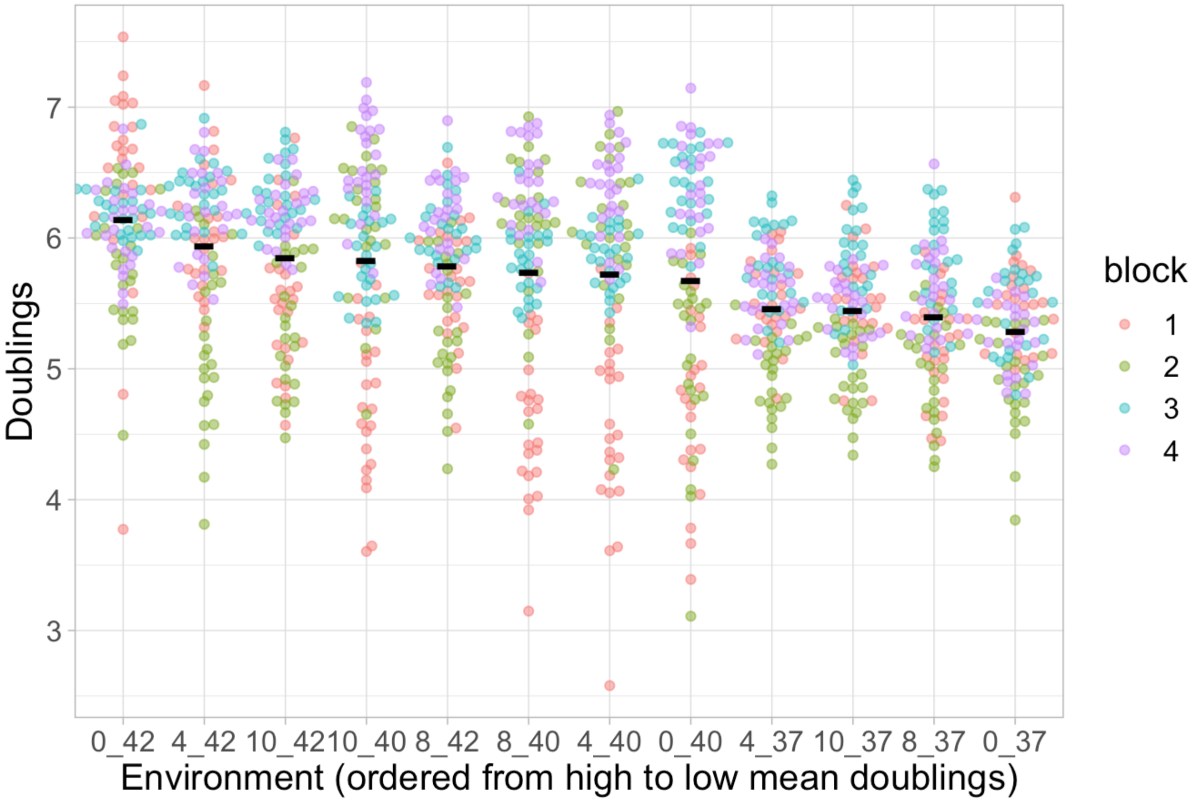

Figure S35: This plot shows that the environmental quality metric 'the mean total growth in the well (reference + competitor) in each environment' does not distinguish well between environments. The data is plotted in the style of figure S32. However, unlike that figure, here there is a stronger effect of the replicate blocks as compared to environment (quantified in table S11).

Table S11: results of ANOVA for environmental quality metric: the mean total growth in the well (reference + competitor) in each environment.

| Source of Variation | SS | df | MS | F statistic | p-value |
|---|---|---|---|---|---|
| Environment | 64.7 | 11 | 5.89 | 18.75 | $< 10^{-15}$ |
| Replicate block | 113.6 | 3 | 37.88 | 120.66 | $< 10^{-15}$ |
| Residuals | 353.2 | 1125 | 0.31 | | |

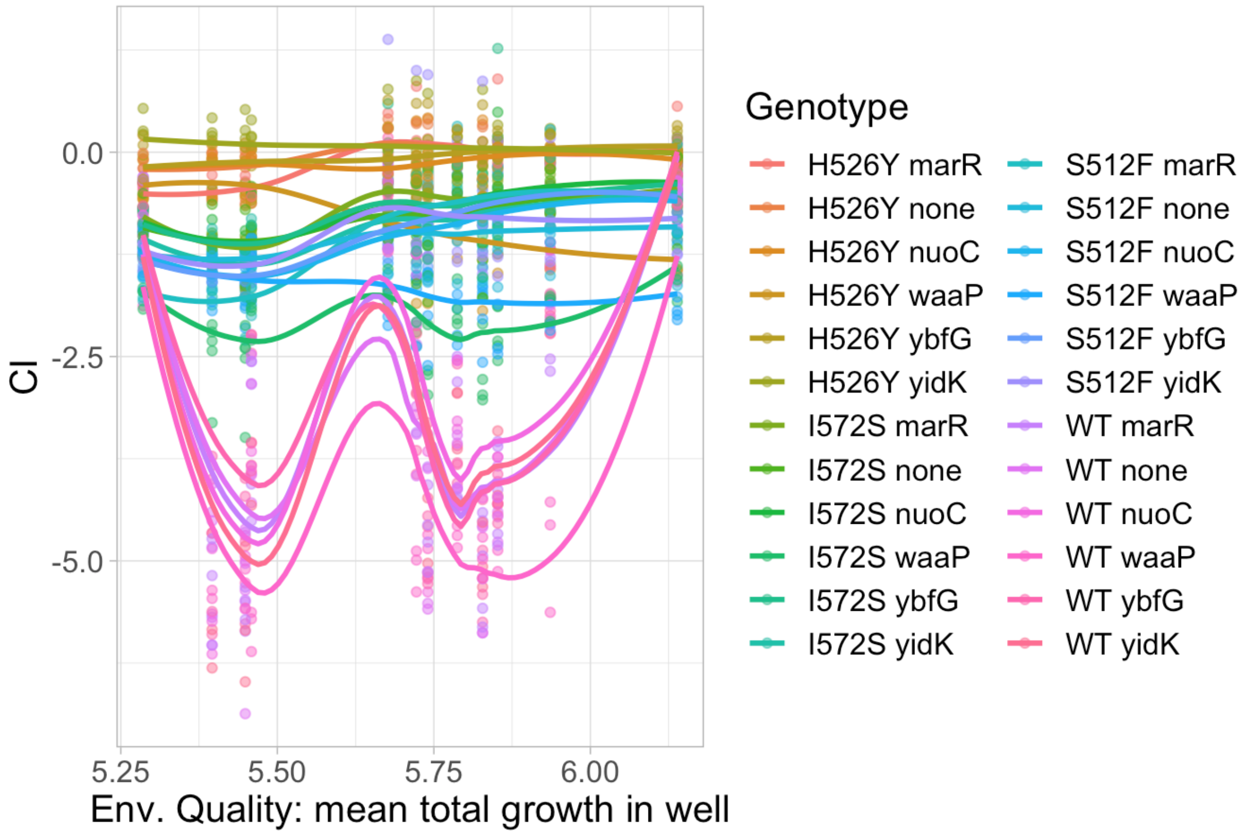

Figure S36: This plot shows that the environmental quality metric 'mean total growth in the well (reference + competitor) in each environment' is not able to distinguish specialist and generalist genotypes. The data is plotted in the style of figure S33. However, unlike that figure, here the antibiotic environments are not grouped together as low quality environments: that's why the specialist genotypes make a 'w' shape in the plot.

###### 1.7.2 Finlay-Wilkinson environmental quality regression for GxG summary statistics

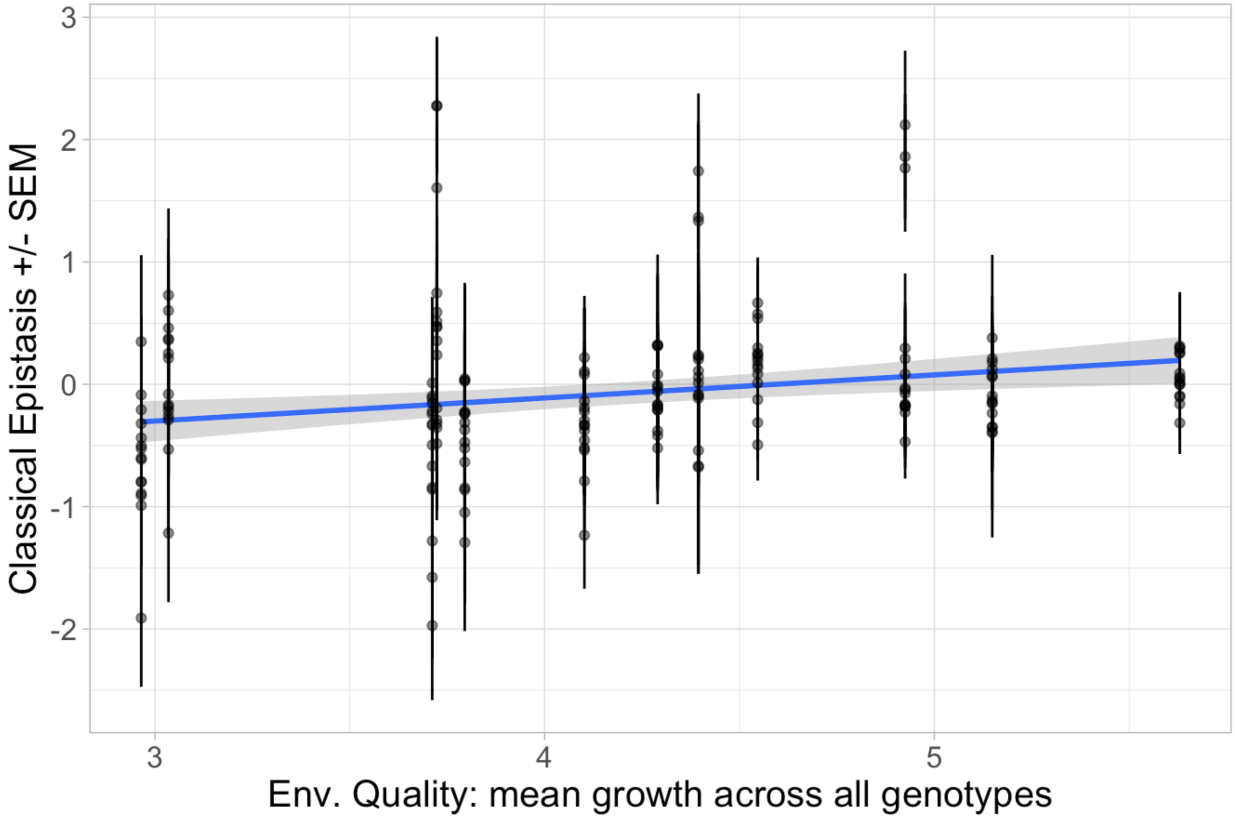

Figure S37: Plot of pairwise epistasis as a function of environmental quality across all genotypes and environments. The x-axis shows the environmental quality and the y-axis shows the mean pairwise epistasis for each of the 15 combinations of knock-outs and ABR *rpoB* mutations, with error bars for the standard error. The blue line shows the mean and the blue shaded region shows the 95% confidence interval of the least-squares estimated linear regression. The linear regression is significant ($F(1, 178) = 10.16$, $p < 0.005$) but does not explain much of the variation in the data (adjusted $R^2 = 0.0487$).

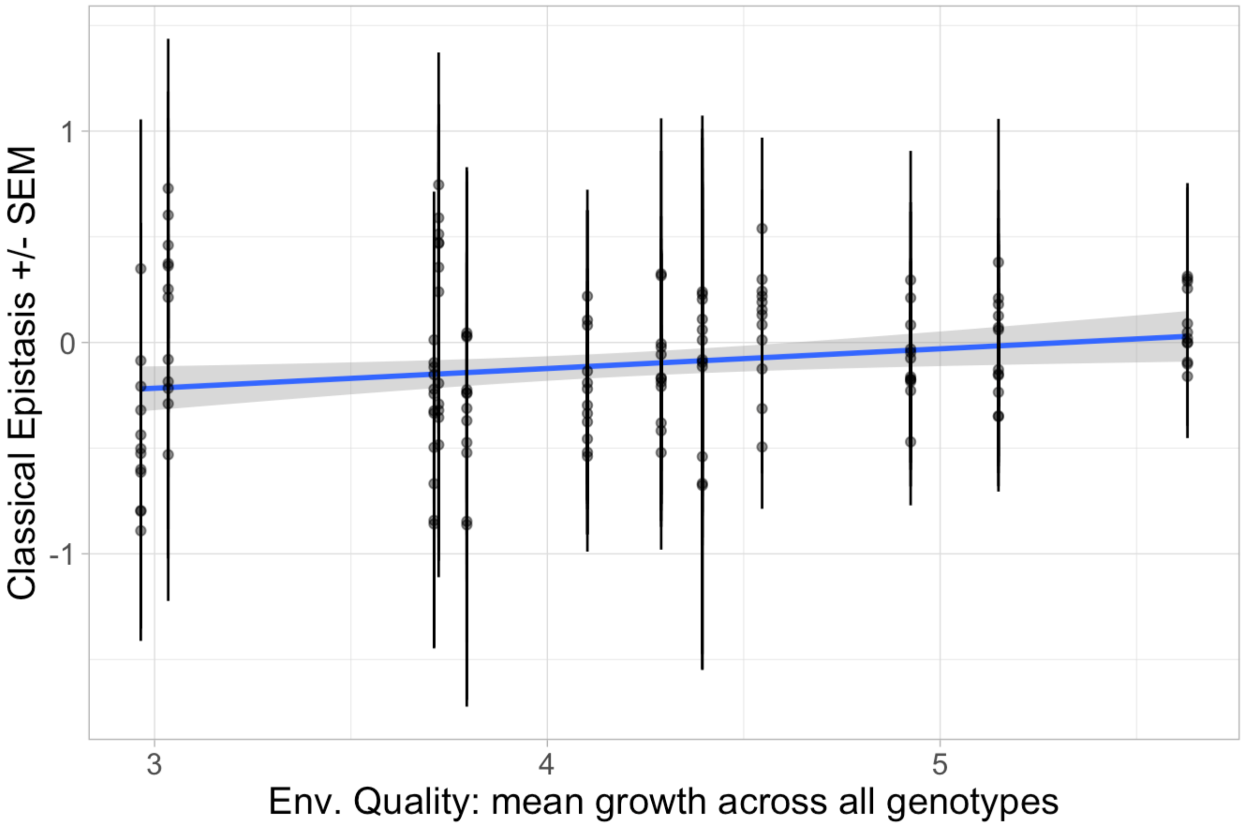

Figure S38: Plot of pairwise epistasis as a function of environmental quality for all gentoypes except the $\Delta waaP$ genotypes, as these have unusually high pairwise epistasis. This plot is similar to figure S37 except that only 12 points are shown (i.e., three $\Delta waaP$ genotypes were excluded) for each environment. The linear regression is marginally significant ($F(1,142) = 6.357$, $p = 0.0128$) but it does not explain much of the variation in the data (adjusted $R^2 = 0.0361$).

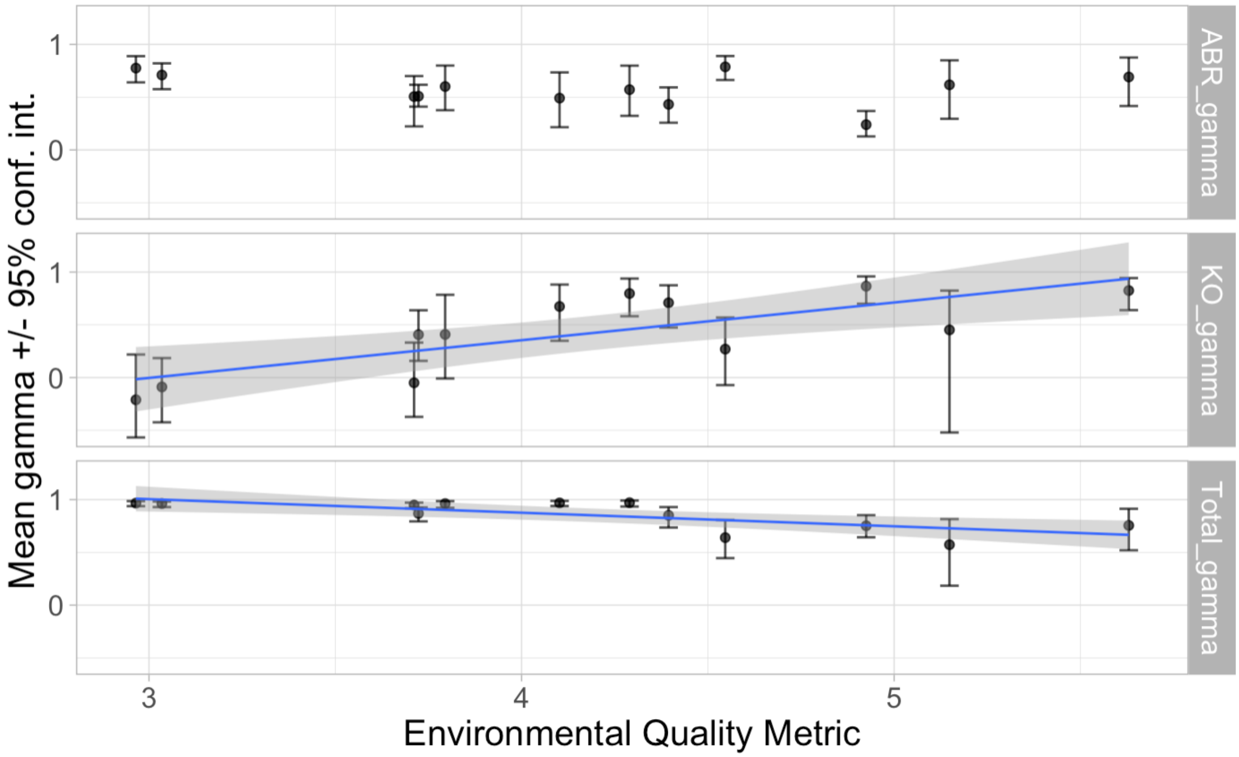

Figure S39: The mean gamma epistasis (y-axis) as a function of environmental quality (x-axis). The points show the mean gamma epistasis, with facets indicating the gene classes. The error bars show the 95% parametric bootstrap confidence intervals. If shown, the blue line indicates the mean of the significant, least-squares estimated linear regression and the grey shaded region shows its 95% confidence interval. Remember that gamma epistasis values near one indicate *low* G×G interactions and values near zero or negative one indicate *high* G×G interactions. **Top facet:** The gamma epistasis exhibited only by ABR *rpoB* mutations (ABR_gamma) is not correlated with either environmental quality ($F(1, 10) = 0.614$, $p = 0.45$) nor antibiotic concentration alone ($F(1, 10) = 0.0803$, $p = 0.78$; see figure S22). **Middle facet:** The gamma epistasis exhibited only by knock-out mutations (KO_gamma) is significantly correlated with environmental quality ($F(1, 10) = 14.3$, $p < 0.005$) and explains more of the variation in the data (adjusted $R^2 = 0.547$) than when regressed against antibiotic concentration alone (adjusted $R^2 = 0.086$; the regression is not significant: $F(1, 10) = 2.04$, $p = 0.184$, see figure S22). Note that this trend is opposite to that found for other measures: G×G is highest in the lowest quality environments and lowest in the highest quality environments. **Bottom facet:** For the total gamma epistasis, the regression on environmental quality is significant ($F(1, 10) = 12.0$, $p < 0.01$) but explains about half the variation in the data (adjusted $R^2 = 0.5$) as compared to $\sim \frac{4}{5}$ of the variation in the data for the regression on antibiotic concentration alone (see main text and figure 3a).

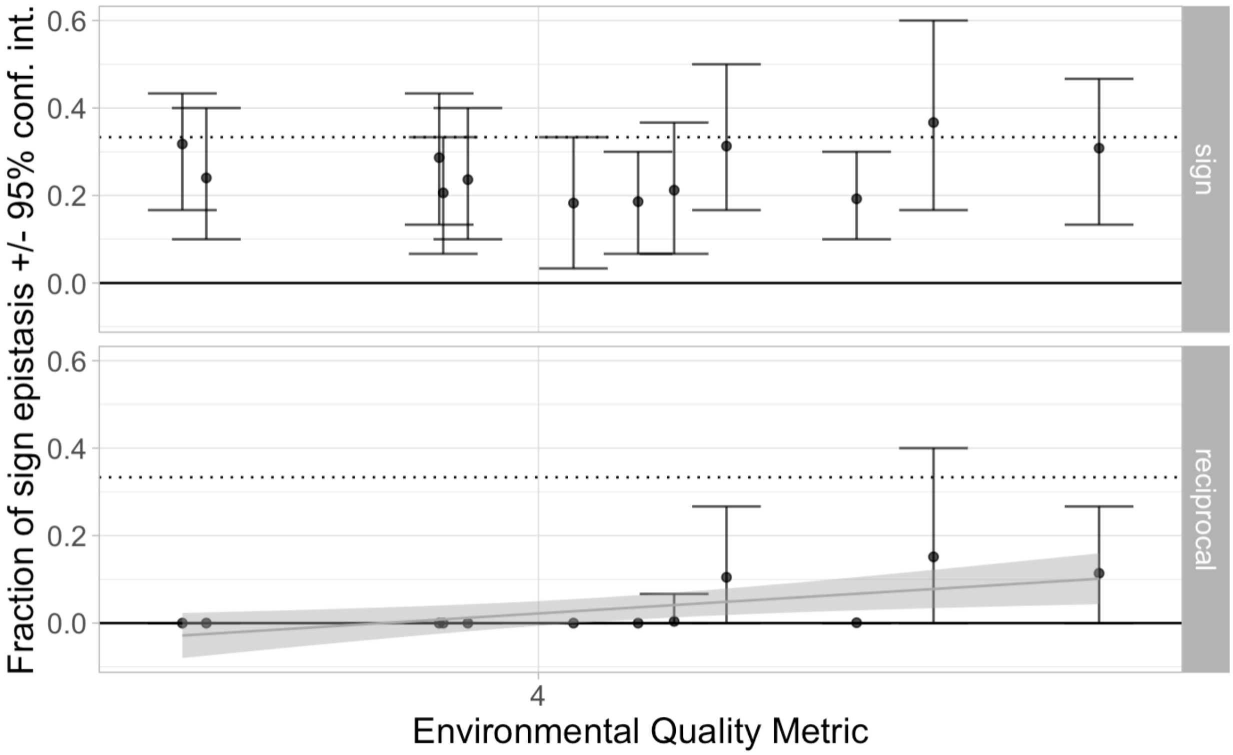

Figure S40: The mean fraction of simple sign (top facet) and reciprocal sign epistasis (bottom facet) as a function of environmental quality (x-axis) shows no specific trend that is an improvement over the effect of antibiotic concentration (figure S23). The points show the mean fraction and the error bars show the 95% parametric bootstrap confidence interval. There is no significant correlation between the environmental quality metric and the fraction of simple sign epistasis ($F(1, 10) = 0.375$, $p = 0.55$). For the reciprocal sign epistasis, the regression on environmental quality is marginally significant ($F(1, 10) = 9.30$, $p = 0.0122$, grey trend-line shows the mean and the shaded region shows the 95% confidence interval of the least-squares estimated linear regression). The regression of reciprocal sign epistasis explains less than half of the variation in the data (adjusted $R^2 = 0.430$) as compared to over 60% of the variation in the data for the regression on antibiotic concentration alone (see figures 3b & S23).

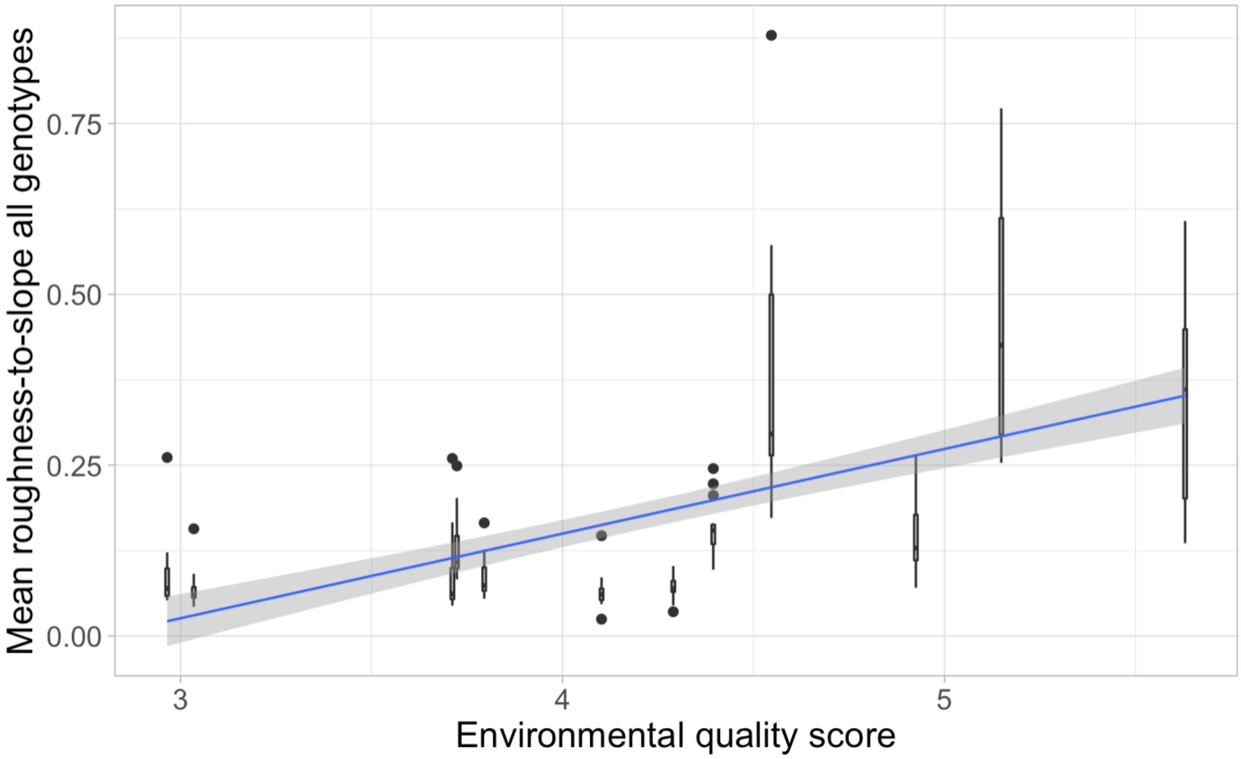

Figure S41: Boxplots of the roughness to slope ratio for all genes (y-axis) as a function of environmental quality (x-axis) shows a significant correlation ($F(1, 178) = 95.7$, $p < 10^{-15}$) but the amount of variation explained by the environmental quality score (adjusted $R^2 = 0.346$) is less than than explained by antibiotic concentration alone ($\sim 50\%$, see figure 3c).

### 2 Materials & Methods

#### 2.1 Constructing and confirming the genotypes

##### 2.1.1 Obtaining the ABR Mutations

*RpoB* H526Y and S512F mutants have strong resistance to rifampicin ($> 200\mu$g/mL) and were already available in the Gordo lab (3). *RpoB* I572N happened to evolve *de novo* by screening for spontaneous mutations with intermediate rifampicin resistance (resistant at 20 $\mu g/mL$ but susceptible at 100 $\mu g/mL$) in rich media (Luria broth) batch culture and then using Sanger sequencing to confirm the SNP location on the *rpoB* locus.

##### 2.1.2 Genetic engineering

P1*vir* phage transduction (4) with rifampicin selective plates was used to move each *rpoB* mutant onto two different fluorescently labeled backgrounds, one with Superfolder GFP and the other with mCherry, both with a chloramphenicol resistance cassette.

Knock-out single mutants were built onto the *rpoB* wild-type GFP-labeled and mCherry-labeled backgrounds by $\lambda$ Red-mediated recombination using PCR products (5). The homology primers listed in table S13 were used to PCR amplify the in-frame, single gene deletion mutants from the Keio collection samples, then these PCR products were used as templates for recombineering. P1 phage transduction with selection for the inserted kanamycin cassette was then used to clean-up any off-target mutations that may have been introduced by the $\lambda$ Red recombinase. However, we did not counterselect to eliminate the kanamycin resistance cassette from knock-outs – meaning that all knock-outs have the gene-of-interest replaced with a kanamycin cassette.

Double-mutants were built by moving the knock-outs onto the *rpoB* mutant backgrounds, again using P1 phage transduction. All sample codes and their corresponding engineered mutations are listed in table S12.

All genotypes were phenotypically screened for their respective markers: resistance to chloramphenicol, susceptibility/resistance to rifampicin and kanamycin, and GFP or mCherry fluorescence. Fluorescence was screened using a BioTek H1 microplate reader with GFP (ex: 469 nm / em: 525 nm) and RFP (ex: 531 nm / em: 593 nm) filters.

Table S12: Sample names and their corresponding engineered mutations.

| Sample | rpoB mutation | Knock-out | Fluorophore | Antibiotic resistance(s) |
|---|---|---|---|---|
| AHG015 (WT) | - | - | sfGFP | chloramphenicol |
| AHG056 | - | $\Delta$ybfG | sfGFP | chloramphenicol, kanamyacin |
| AHG060 | - | $\Delta$marR | sfGFP | chloramphenicol, kanamyacin |
| AHG064 | - | $\Delta$yidK | sfGFP | chloramphenicol, kanamyacin |
| AHG066 | - | $\Delta$nuoC | sfGFP | chloramphenicol, kanamyacin |
| AHG052 | - | $\Delta$waaP | sfGFP | chloramphenicol, kanamyacin |
| AHG079 (Ref) | H526Y | - | mCherry | chloramphenicol, rifampicin |
| AHG080 | H526Y | - | sfGFP | chloramphenicol, rifampicin |
| AHG090 | H526Y | $\Delta$ybfG | sfGFP | chloramphenicol, rifampicin, kanamyacin |
| AHG092 | H526Y | $\Delta$marR | sfGFP | chloramphenicol, rifampicin, kanamyacin |
| AHG094 | H526Y | $\Delta$yidK | sfGFP | chloramphenicol, rifampicin, kanamyacin |
| AHG096 | H526Y | $\Delta$nuoC | sfGFP | chloramphenicol, rifampicin, kanamyacin |
| AHG088 | H526Y | $\Delta$waaP | sfGFP | chloramphenicol, rifampicin, kanamyacin |
| AHG082 | S512F | - | sfGFP | chloramphenicol, rifampicin |
| AHG100 | S512F | $\Delta$ybfG | sfGFP | chloramphenicol, rifampicin, kanamyacin |
| AHG102 | S512F | $\Delta$marR | sfGFP | chloramphenicol, rifampicin, kanamyacin |
| AHG104 | S512F | $\Delta$yidK | sfGFP | chloramphenicol, rifampicin, kanamyacin |

Continued on next page

**Table S12 – continued from previous page**

| Sample | rpoB mutation | Knock-out | Fluorophore | Antibiotic resistance(s) |
|---|---|---|---|---|
| AHG106 | S512F | ΔnuoC | sfGFP | chloramphenicol, rifampicin, kanamyacin |
| AHG098 | S512F | ΔwaaP | sfGFP | chloramphenicol, rifampicin, kanamyacin |
| AHG068 | I572S | - | sfGFP | chloramphenicol, rifampicin |
| AHG070 | I572S | ΔybfG | sfGFP | chloramphenicol, rifampicin, kanamyacin |
| AHG072 | I572S | ΔmarR | sfGFP | chloramphenicol, rifampicin, kanamyacin |
| AHG074 | I572S | ΔyidK | sfGFP | chloramphenicol, rifampicin, kanamyacin |
| AHG076 | I572S | ΔnuoC | sfGFP | chloramphenicol, rifampicin, kanamyacin |
| AHG078 | I572S | ΔwaaP | sfGFP | chloramphenicol, rifampicin, kanamyacin |

Table S13: The sequences of the homology primers used to PCR amplify the in-frame, single gene deletion mutants from the listed Keio collection samples, respectively. Product lengths are given for when the gene is present in MG1655 and for when the gene is knocked-out in the Keio sample. The names of the primers indicate their positions in the MG1655 reference genome (GenBank accession CP032667.1). PCR for the Datsenko-Wanner recombineering was done with New England Biolabs Phusion High-Fidelity DNA Polymerase (M0530) by following the manufacturer's guidelines with a 2.5 minute elongation at 67°C.

| **ΔmarR** | |
|---|---|
| Primer name | Primer sequence |
| marR_2262520_For | AAGTAACAACTGGCTGCGTG |
| marR_2264206_Rev | CCACAAAATAACCGCCACCA |
| Product length in MG1655: | 1687 bp |
| Product length in Keio JW5248: | 2637 bp |
| **ΔnuoC** | |
| Primer name | Primer sequence |
| nuoC_1478699_For | GAGCGTATGAACCCGGAAAC |
| nuoC_1482106_Rev | GGCTCCATTTCATCGGCATT |
| Product length in MG1655: | 3408 bp |
| Product length in Keio JW5375: | 2933 bp |
| **ΔpstA** | |
| Primer name | Primer sequence |
| pstA_4613126_For | AATATCCCGATTGTTGGCGC |
| pstA_4615306_Rev | TCCTGCTTCAGTTCGGTGAT |
| Product length in MG1655: | 2181 bp |
| Product length in Keio JW3704: | 2618 bp |
| **ΔsurA** | |
| Primer name | Primer sequence |
| surA_3826005_For | CGTCTCTGCTGCAATCTGAC |
| surA_3828504_Rev | GGAATGCCAGCGTCGTTAA |
| Product length in MG1655: | 2500 bp |
| Product length in Keio JW0052: | 2560 bp |
| **ΔwaaP** | |
| Primer name | Primer sequence |
| waaP_76747_For | AATCGCCGATTTCCAGAAGC |
| waaP_78757_Rev | CGCCCTTTATCACGACCAAG |
| Product length in MG1655: | 2011 bp |
| Product length in Keio JW3605: | 2541 bp |
| **ΔwcaE** | |
| Primer name | Primer sequence |
| wcaE_1752899_For | AATTATCCTCACCGGTGGCA |
| wcaE_1755232_Rev | GAATTTCGGGTTGCAGGTGT |
| Product length in MG1655: | 2334 bp |
| Product length in Keio JW2040: | 2915 bp |
| **ΔybfG** | |
| Primer name | Primer sequence |
| ybfG_3164439_For | CAGAAGGTCGCTAATGTGCC |
| ybfG_3166129_Rev | TACCGCCACATTACTGCTGA |
| Product length in MG1655: | 1691 bp |
| Product length in Keio JW5094: | 2656 bp |

Continued on next page

**Table S13 – continued from previous page**

| **ΔyhhS** | |
|---|---|
| Primer name | Primer sequence |
| yhhS_271090_For | TAGCGCTGTGGTGAAGGTAA |
| yhhS_273647_Rev | ATGGCTGAACACTGGATGGA |
| Product length in MG1655: | 2558 bp |
| Product length in Keio JW5945: | 2635 bp |
| **ΔyidK** | |
| Primer name | Primer sequence |
| yidK_22529_For | TTTTCAATTTGCCGCCGTTG |
| yidK_25340_Rev | AACCAGTAATCAGCGTCCCA |
| Product length in MG1655: | 2812 bp |
| Product length in Keio JW3655: | 2424 bp |
| **ΔyzgL** | |
| Primer name | Primer sequence |
| yzgL_318532_For | GGTACCGCGCGTGAATATTT |
| yzgL_320142_Rev | CGTTCCTGACTGAATCGCTG |
| Product length in MG1655: | 1611 bp |
| Product length in Keio JW3390: | 2657 bp |

**Sequence data for Superfolder GFP inserted construct (in GenBank format):**

```
LOCUS       sfGFP                   2301 bp    DNA
DEFINITION  Template plasmid RB300, partial sequence.
ACCESSION   .
VERSION     .
SOURCE      Template plasmid RB300 made by Roberto Balbontin
  ORGANISM  Template plasmid RB300
            other sequences; artificial sequences; vectors.
REFERENCE   1  (bases 1 to 2301)
  AUTHORS   Balbontin, R.
FEATURES             Location/Qualifiers
     source          1..2301
                     /organism="synthetic construct"
                     /mol_type="other DNA"
     misc_feature    1..3
                     /note="yiaS"
     misc_feature    35..66
                     /note="ysaC"
     misc_feature    211..297
                     /note="PLTetO-1 promoter"
     misc_feature    298..324
                     /note="RBSpZA31-luc"
     CDS             325..1044
                     /note="Superfolder GFP fluorophore"
                     /codon_start=1
                     /transl_table=11
                     /product="sfGFP"
                     /protein_id="QBQ65835"
                     /translation="MASKGEELFTGVVPILVELDGDVNGHKFSVRGEGEGDATNGKLT
                     LKFICTTGKLPVPWPTLVTTLTYGVQCFSRYPDHMKQHDFFKSAMPEGYVQERTISFK
                     DDGTYKTRAEVKFEGDTLVNRIELKGIDFKEDGNILGHKLEYNFNSHNVYITADKQKN
                     GIKANFKIRHNVEDGSVQLADHYQQNTPIGDGPVLLPDNHYLSTQSVLSKDPNEKRDH
                     MVLLEFVTAAGITHGMDELYK-"
     misc_feature    1093..1138
                     /note="FRT site"
     CDS             1347..1985
                     /note="chloramphenicol resistance"
                     /codon_start=1
                     /transl_table=11
                     /product="type A chloramphenicol O-acetyltransferase"
                     /protein_id="WP_015420186 "
                     /translation="MEKKITGYTTVDISQWHRKEHFEAFQSVAQCTYNQTVQLDITAF
                     LKTVKKNKHKFYPAFIHILARLMNAHPELRMAMKDGELVIWDSVHPCYTVFHEQTETF
                     SSLWSEYHDDFRQFLHIYSQDVACYGENLAYFPKGFIENMFFVSANPWVSFTSFDLNV
                     ANMDNFFAPVFTMGKYYTQGDKVLMPLAIQVHHAVCDGFHVGRCLMNTTVLR"
     misc_feature    2023..2068
                     /note="FRT site"
     misc_feature    2120..2280
                     /note="ysaD"
     misc_feature    2299..2301
                     /note="yiaT"
ORIGIN
        1 taatccctca cgccggggct tcatcgcccc ggcactacga attgatatgt tccttgctgt
```

```
       61 aacgccgctt ccacgctgct ggcgttaaac cagtatgttt ctgaaaaatc tgccggaaat
      121 agccaacgtc attaaagata tcgacgtcta agaaaccatt attatcatga cattaaccta
      181 taaaaatagg cgtatcacga ggccctttcg tcttcacctc gagtccctat cagtgataga
      241 gattgacatc cctatcagtg atagagatac tgagcacatc agcaggacgc actgaccgaa
      301 ttcattaaag aggagaaagg taccatggct tctaaaggtg aagaactgtt caccggtgtt
      361 gttccgatcc tggttgaact ggatggtgat gttaacggcc acaaattctc tgttcgtggt
      421 gaaggtgaag gtgatgcaac caacggtaaa ctgaccctga aattcatctg cactaccggt
      481 aaactgccgg ttccatggcc gactctggtg actaccctga cctatggtgt tcagtgtttt
      541 tctcgttacc cggatcacat gaagcagcat gatttcttca aatctgcaat gccggaaggt
      601 tatgtacagg agcgcaccat ttctttcaaa gacgatggca cctacaaaac ccgtgcagag
      661 gttaaatttg aaggtgatac tctggtgaac cgtattgaac tgaaaggcat tgatttcaaa
      721 gaggacggca acatcctggg ccacaaactg gaatataact tcaactccca taacgtttac
      781 atcaccgcag acaaacagaa gaacggtatc aaagctaact tcaaaattcg ccataacgtt
      841 gaagacggta gcgtacagct ggcggaccac taccagcaga acactccgat cggtgatggt
      901 ccggttctgc tgccggataa ccactacctg tccacccagt ctgttctgtc caaagacccg
      961 aacgaaaagc gcgaccacat ggtgctgctg gagttcgtta ctgcagcagg tatcacgcac
     1021 ggcatggatg agctctacaa ataatgaatt cgagcattta aatctagagg catccatatg
     1081 aatatcctcc ttagttccta ttccgaagtt cctattctct agaaagtata ggaacttcgg
     1141 cgcgcctacc tgtgacggaa gatcacttcg cagaataaat aaatcctggt gtccctgttg
     1201 ataccgggaa gccctgggcc aacttttggc gaaaatgaga cgttgattgg cacgtaagag
     1261 gttccaactt tcaccataat gaaataagat cactaccggg cgtatttttt gagttgtcga
     1321 gattttcagg agctaaggaa gctaaaatgg agaaaaaaat cactggatat accaccgttg
     1381 atatatccca atggcatcgt aaagaacatt ttgaggcatt tcagtcagtt gctcaatgta
     1441 cctataacca gaccgttcag ctggatatta cggccttttt aaagaccgta aagaaaaata
     1501 agcacaagtt ttatccggcc tttattcaca ttcttgcccg cctgatgaat gctcatccgg
     1561 aattacgtat ggcaatgaaa gacggtgagc tggtgatatg ggatagtgtt cacccttgtt
     1621 acaccgtttt ccatgagcaa actgaaacgt tttcatcgct ctggagtgaa taccacgacg
     1681 atttccggca gtttctacac atatattcgc aagatgtggc gtgttacggt gaaaacctgg
     1741 cctatttccc taaagggttt attgagaata tgtttttcgt ctcagccaat ccctgggtga
     1801 gtttcaccag ttttgattta aacgtggcca atatggacaa cttcttcgcc cccgttttca
     1861 ccatgggcaa atattatacg caaggcgaca aggtgctgat gccgctggcg attcaggttc
     1921 atcatgccgt ttgtgatggc ttccatgtcg gcagatgctt aatgaataca acagtactgc
     1981 gatgagtggc agggcggggc gtaaggcgcg ccatttaaat gaagttccta ttccgaagtt
     2041 cctattctct agaaagtata ggaacttcga agcagctcca gcctacacta ttgcatggag
     2101 caggccgaca acgtcgagtc actgcataac ctgttgctgc ccgccgatgc gccatttacg
     2161 acatttgaag gcaagggatt gttcagccat aagatcctga ttcagacgcc aggttcccgg
     2221 tatgaacaga gcaatccaga gcaacaacct ttgtggcatt acgacggagc accagccgca
     2281 ccgacatcca ccggtgaatt a
//
```

**Sequence data for mCherry inserted construct (in GenBank format):**

```
LOCUS       mCherry                   2292 bp    DNA
DEFINITION  Template plasmid RB296, partial sequence.
ACCESSION   .
VERSION     .
SOURCE      Template plasmid RB296 made by Roberto Balbontin
  ORGANISM  Template plasmid RB296
            other sequences; artificial sequences; vectors.
REFERENCE   1  (bases 1 to 2292)
  AUTHORS   Balbontin, R.
FEATURES             Location/Qualifiers
     source          1..2292
                     /organism="synthetic construct"
                     /mol_type="other DNA"
     misc_feature    1..3
                     /note="yiaS"
     misc_feature    35..66
                     /note="ysaC"
     misc_feature    211..297
                     /note="PLTetO-1 promoter"
     misc_feature    298..324
                     /note="RBSpZA31-luc"
     CDS             325..1035
                     /note="mCherry fluorophore"
                     /codon_start=1
                     /transl_table=11
                     /product="mCherry"
                     /protein_id="AZQ25025.1"
                     /translation="MVSKGEEDNMAIIKEFMRFKVHMEGSVNGHEFEIEGEGEGRPYE
                     GTQTAKLKVTKGGPLPFAWDILSPQFMYGSKAYVKHPADIPDYLKLSFPEGFKWERVM
                     NFEDGGVVTVTQDSSLQDGEFIYKVKLRGTNFPSDGPVMQKKTMGWEASSERMYPEDG
                     ALKGEIKQRLKLKDGGHYDAEVKTTYKAKKPVQLPGAYNVNIKLDITSHNEDYTIVEQ
                     YERAEGRHSTGGMDELYK"
     misc_feature    1084..1129
                     /note="FRT site"
     CDS             1338..1976
                     /note="chloramphenicol resistance"
                     /codon_start=1
                     /transl_table=11
                     /product="type A chloramphenicol O-acetyltransferase"
                     /protein_id="WP_015420186 "
                     /translation="MEKKITGYTTVDISQWHRKEHFEAFQSVAQCTYNQTVQLDITAF
                     LKTVKKNKHKFYPAFIHILARLMNAHPELRMAMKDGELVIWDSVHPCYTVFHEQTETF
                     SSLWSEYHDDFRQFLHIYSQDVACYGENLAYFPKGFIENMFFVSANPWVSFTSFDLNV
                     ANMDNFFAPVFTMGKYYTQGDKVLMPLAIQVHHAVCDGFHVGRCLMNTTVLR"
     misc_feature    2014..2059
                     /note="FRT site"
     misc_feature    2111..2271
                     /note="ysaD"
     misc_feature    2290..2292
                     /note="yiaT"
ORIGIN
        1 taatccctca cgccggggct tcatcgcccc ggcactacga attgatatgt tccttgctgt
       61 aacgccgctt ccacgctgct ggcgttaaac cagtatgttt ctgaaaaatc tgccggaaat
```

```
     121 agccaacgtc attaaagata tcgacgtcta agaaaccatt attatcatga cattaaccta
     181 taaaaatagg cgtatcacga ggccctttcg tcttcacctc gagtccctat cagtgataga
     241 gattgacatc cctatcagtg atagagatac tgagcacatc agcaggacgc actgaccgaa
     301 ttcattaaag aggagaaagg taccatggtg agcaagggcg aggaggataa catggccatc
     361 atcaaggagt tcatgcgctt caaggtgcac atggagggct ccgtgaacgg ccacgagttc
     421 gagatcgagg gcgagggcga gggccgcccc tacgagggca cccagaccgc caagctgaag
     481 gtgaccaagg gtggccccct gcccttcgcc tgggacatcc tgtcccctca gttcatgtac
     541 ggctccaagg cctacgtgaa gcaccccgcc gacatccccg actacttgaa gctgtccttc
     601 cccgagggct tcaagtggga gcgcgtgatg aacttcgagg acggcggcgt ggtgaccgtg
     661 acccaggact cctccctgca ggacggcgag ttcatctaca aggtgaagct gcgcggcacc
     721 aacttcccct ccgacggccc cgtaatgcag aagaagacca tgggctggga ggcctcctcc
     781 gagcggatgt accccgagga cggcgccctg aagggcgaga tcaagcagag gctgaagctg
     841 aaggacggcg gccactacga cgctgaggtc aagaccacct acaaggccaa gaagcccgtg
     901 cagctgcccg gcgcctacaa cgtcaacatc aagttggaca tcacctccca caacgaggac
     961 tacaccatcg tggaacagta cgaacgcgcc gagggccgcc actccaccgg cggcatggac
    1021 gagctgtaca agtaatgaat tcgagcattt aaatctagag gcatccatat gaatatcctc
    1081 cttagttcct attccgaagt tcctattctc tagaaagtat aggaacttcg gcgcgcctac
    1141 ctgtgacgga agatcacttc gcagaataaa taaatcctgg tgtccctgtt gataccggga
    1201 agccctgggc caacttttgg cgaaaatgag acgttgattg gcacgtaaga ggttccaact
    1261 ttcaccataa tgaaataaga tcactaccgg gcgtattttt tgagttgtcg agattttcag
    1321 gagctaagga agctaaaatg gagaaaaaaa tcactggata taccaccgtt gatatatccc
    1381 aatggcatcg taaagaacat tttgaggcat ttcagtcagt tgctcaatgt acctataacc
    1441 agaccgttca gctggatatt acggcctttt taaagaccgt aaagaaaaat aagcacaagt
    1501 tttatccggc ctttattcac attcttgccc gcctgatgaa tgctcatccg gaattacgta
    1561 tggcaatgaa agacggtgag ctggtgatat gggatagtgt tcacccttgt tacaccgttt
    1621 tccatgagca aactgaaacg ttttcatcgc tctggagtga ataccacgac gatttccggc
    1681 agtttctaca catatattcg caagatgtgg cgtgttacgg tgaaaacctg gcctatttcc
    1741 ctaaagggtt tattgagaat atgtttttcg tctcagccaa tccctgggtg agtttcacca
    1801 gttttgattt aaacgtggcc aatatggaca acttcttcgc ccccgttttc accatgggca
    1861 aatattatac gcaaggcgac aaggtgctga tgccgctggc gattcaggtt catcatgccg
    1921 tttgtgatgg cttccatgtc ggcagatgct taatgaatac aacagtactg cgatgagtgg
    1981 cagggcgggg cgtaaggcgc gccatttaaa tgaagttcct attccgaagt tcctattctc
    2041 tagaaagtat aggaacttcg aagcagctcc agcctacact attgcatgga gcaggccgac
    2101 aacgtcgagt cactgcataa cctgttgctg cccgccgatg cgccatttac gacatttgaa
    2161 ggcaagggat tgttcagcca taagatcctg attcagacgc caggttcccg gtatgaacag
    2221 agcaatccag agcaacaacc tttgtggcat tacgacggag caccagccgc accgacatcc
    2281 accggtgaat ta
//
```

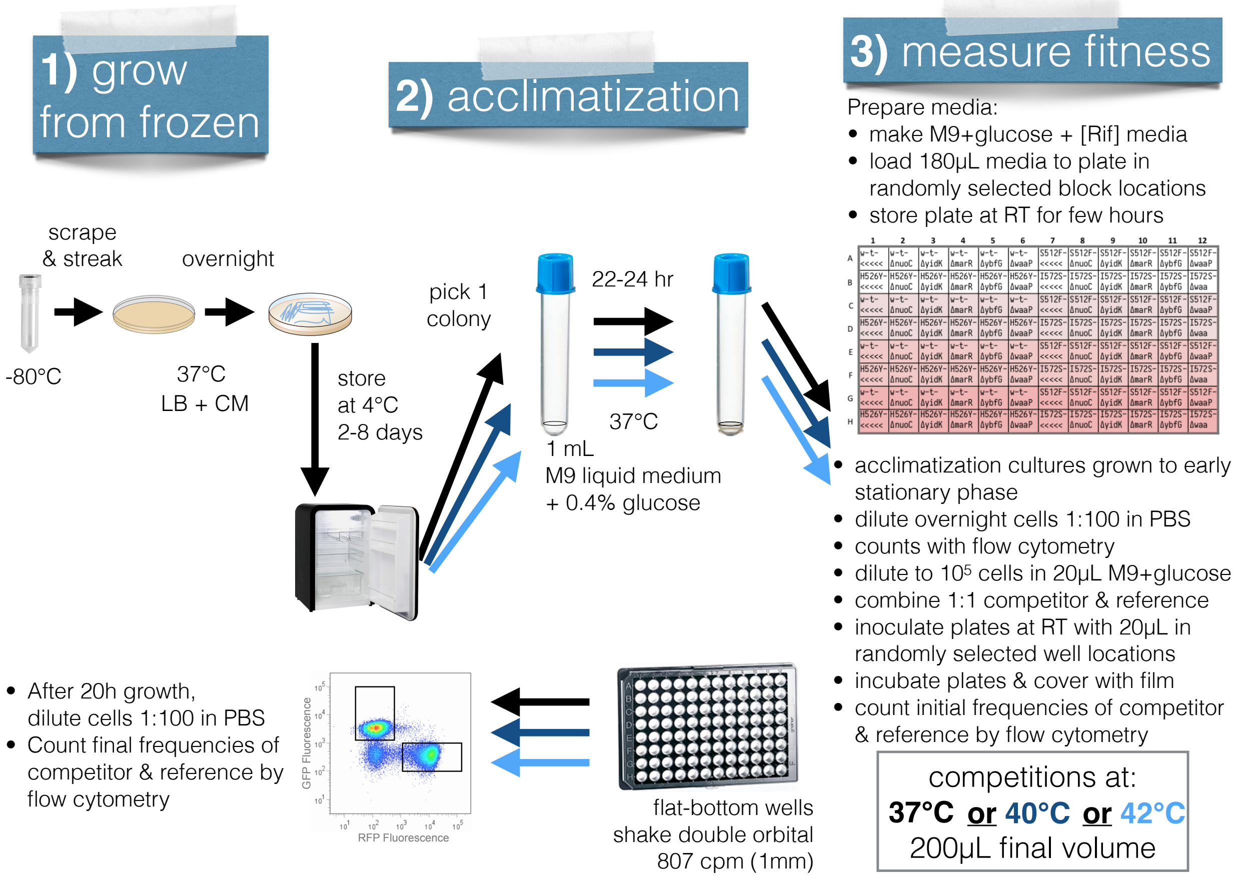

Figure S42: **Summary of the protocol used to estimate fitness in different antibiotic concentration and temperature environments. 1)** To ameliorate the possibility of *de novo* mutations, frozen glycerol stocks were bottle-necked to single colonies by streaking on agar plates. **2)** After streaking and growth on agar plates, all cells experienced the same temperature acclimitization of at least 40 hours and at most 8 days at 4°C followed by batch culture growth at 37°C for 22-24 hours. **3)** Batch culture competitions took place in 96-well plates with randomized positions of the four blocks of antibiotic concentrations and the 24 samples. Only one plate was incubated at a time, therefore triple arrows show steps in the protocol that had to be repeated for each temperature environment in order to sample a complete block of all samples in all environments. Regardless of the temperature environment where competition occurred, batch culture acclimitization always occurred at 37°C.

##### 2.2.3 Estimating competitive index

We used `R` (4.1.1) as run using `RStudio` (2021.09.0+351) to calculate the competitive index using the equation indicated in the results. The GitLab repository for this manuscript, https://gitlab.com/evoldynamics/epistasis-decreases-with-increasing-antibiotic-pressure, contains the competitive index estimates and `R` code for calculating the competitive index from the cell counts data, `calculating_selection.Rmd`.

#### 2.3 Analyses

Parametric bootstrapping (10 000 bootstrap replicates) was used to determine the 95% confidence intervals on all of the summary statistics of epistasis. For the diminishing-returns epistasis, linear regressions were run on each of the 10 000 bootstrap replicates (x = fitness of the background, y = fitness effect of the mutation) and the mean slope value was retained in order to determine the 95% confidence intervals across all linear regressions.

We fitted a linear model in `R` using the equation $\hat{w} \sim G_{rpoB} + G_{\mathrm{KO}} + E_{\mathrm{T}} + E_{\mathrm{AB}}$ and the `lm` command from the `stats` package (version 4.1.1). The genotypic effects of *rpoB* ($G_{rpoB}$) and knock-out ($G_{\mathrm{KO}}$) mutations were treated as categorical variables (in `R`: unordered factors), while the environmental effects of temperature ($E_{\mathrm{T}}$) and antibiotic concentration ($E_{\mathrm{AB}}$) were treated as semi-quantitative variables (in `R`: ordered factors) using polynomial contrasts.

All analyses are publicly available on GitLab at https://gitlab.com/evoldynamics/epistasis-decreases-with-increasing-antibiotic-pressure. The epistasis calculations can be found in `estimating_epistasis.Rmd`, the linear model fitting can be found in `ANOVA_etal.Rmd`, and Finlay-Wilkinson regression can be found in `FinleyWilkinson.Rmd`.
